# Population structure and evolution of *Salmonella enterica* serotype Typhi in Zimbabwe before a typhoid conjugate vaccine immunization campaign

**DOI:** 10.1101/2022.09.01.506167

**Authors:** Gaetan Thilliez, Tapfumanei Mashe, Blessmore V. Chaibva, Valerie Robertson, Matt Bawn, Andrew Tarupiwa, Faustinos Tatenda Takawira, Marleen M. Kock, Stanley Midzi, Lusubilo W. Mwamakamba, Jorge Matheu, Robert A. Kingsley, Marthie M. Ehlers

## Abstract

**Background:** The continued emergence of *Salmonella enterica* serovar Typhi (*S*. Typhi) with ever increasing antimicrobial resistance (AMR), necessitates the use of vaccines in endemic countries. A typhoid fever outbreak in Harare, Zimbabwe in 2018 from a multidrug resistant *S*. Typhi with additional resistance to ciprofloxacin was the catalyst for the introduction of a typhoid conjugate vaccine program. To investigate the historic emergence and evolution of AMR of endemic *S*. Typhi in Zimbabwe and determined the population structure, gene flux and sequence polymorphisms of strains isolated prior to mass typhoid vaccination to provide a baseline for future evaluation of the effect of the vaccination program.

**Methods:** We determined the population structure, gene flux and sequence polymorphisms and reconstructed the evolution of AMR. The *S*. Typhi population structure was investigated in the context the genome sequence of 1904 strains isolated from 65 countries to reconstruct spread of endemic strains into Zimbabwe.

**Findings:** The population structure of *S*. Typhi in Zimbabwe is dominated by multidrug resistant genotype 4.3.1.1 (H58) that spread to Zimbabwe from neighboring countries around 2009. Evolution of AMR within Zimbabwe included acquisition of an IncN plasmid carrying a *qnrS* gene and a mutation in the quinolone resistance determining region of *gyrA* gene, both implicated in resistance to quinolone antibiotics. A minority population of antimicrobial susceptible *S*. Typhi genotype 3.3.1 strains was detected in typhoid cases.

**Interpretation:** The currently dominant *S*. Typhi population is genotype 4.3.1.1 that spread to Zimbabwe and acquired additional AMR though acquisition of a plasmid and mutation of the *gyrA* gene. This study provides a baseline for future evaluation of the impact of the Typhoid Conjugate Vaccine program in Harare.

**Funding:** RAK and GT were supported by Bill and Melinda Gates Foundation project OPP1217121 and the BBSRC Institute Strategic Programme BB/R012504/1 and its constituent project BBS/E/F/000PR10348.

## Introduction

Typhoid fever is a systemic disease caused by *Salmonella enterica* serotype Typhi (*S*. Typhi) that remains an important cause of morbidity and mortality in low-resource settings (1, 2). The current global burden of disease is estimated at 11 to 18 million infections resulting in 135 900 deaths annually, with the majority recorded in South Asia and Africa (3-5). Prior to the use of antimicrobial therapy for management of typhoid fever, case fatality rates exceeded 20% due to complications such as intestinal perforation (6). Timely access to effective antimicrobial therapy is central to preventing complications such as intestinal perforation and death (7). Fluoroquinolones are generally used in resource-limited countries as the primary therapy for typhoid for decades (6). Recent emergence of fluoroquinolone and cephalosporin resistant strains of *S*. Typhi has resulted in an increased reliance on azithromycin and carbapenems which are expensive and often inaccessible in resource-limited settings where typhoid is most common (6, 8). The emergence and escalating antimicrobial resistance throughout the world have focused increasing attention on the use of typhoid vaccines (6, 8, 9).

Multiple outbreaks of typhoid fever have been reported in Zimbabwe since 2009, with most beginning during the rainy season (10, 11). A typhoid outbreak caused by a ciprofloxacin-resistant strain of *S*. Typhi was detected in Harare, Zimbabwe, in September 2018 (12). Analysis of a small number of strains of *S*. Typhi isolated from typhoid fever cases between 2012 and 2019 revealed that most isolates during this period were H58 encoding resistance to aminoglycoside, β-lactam, phenicol, sulphonamide, tetracycline and fluoroquinolone antibiotics (13). In response, an emergency reactive vaccination campaign using Typhoid Conjugate Vaccine (TCV) was implemented from February to March 2019, targeting more than 323 000 persons who were at high risk for typhoid infection in Harare. Initial reports suggested that the vaccine provided moderate protection against typhoid fever, with an adjusted vaccine effectiveness of up to 67% (11). Further epidemiological investigation of the effect of the vaccine program on *S*. Typhi incidence and population structure of the pathogen are needed to fully evaluate the outcome (14). The aim of this study was to investigate the population structure of *S*. Typhi isolates from urban areas of Harare to establish the history of spread of the current endemic clones in the context of the global *S*. Typhi population and the understand the molecular basis and evolution of antimicrobial resistance in Zimbabwe. We focused on the population structure of *S*. Typhi prior to the TCV vaccination program to provide a baseline for future evaluation of potential effects of the program on endemic *S*. Typhi in Harare.

## Methods

### Bacterial strains and data sources

Epidemiological data was obtained from the Harare city department health reports and clinical records. The extracted information included details regarding demographic indicators, age, sex, suburb and clinical presentation of the disease. Ethics approval for the study was granted by the University of Pretoria, South Africa (779/2018) and Medical Research Council of Zimbabwe (MRCZ/A/2369). A total of 95 *S*. Typhi isolates from Zimbabwe were investigated in the context of 1,904 isolates collected between 1905 and 2019 and originated from 65 countries spanning six continents in this study (Asia, Africa, North and South America, Europe, and Australia and Oceania) (Table S1). Of the 95 isolates 38 *S*. Typhi from Zimbabwe were previously sequenced by Mashe *et al*. (12) and Ingle *et al*. (15) while 57 were sequenced for this study. This isolate set is a convenience sample of strains from Zimbabwe from stool or blood. *S*. Typhi strains were identified using biochemical and slide agglutination as described previously (16, 17). Susceptibility of 68 *S*. Typhi (2018) was determined using disc diffusion tests (Kirby-Bauer) (18) with concentrations of antibiotics as follows: ampicillin (10 µg), chloramphenicol (30 µg), trimethoprim/sulfamethoxazole (1.25/23.75 µg), ceftriaxone (30 µg), azithromycin (15 µg), ciprofloxacin (5 µg) and tetracycline (30 µg) (Mast, Hampshire, UK) (18). Zone diameters were measured and interpreted using CLSI guidelines (19). Accession number and associated metadata are provided in Supplementary Table 1.

### Whole-genome sequencing and quality control

Isolates were cultured on MacConkey agar for 18–20 hours at 37°C. A single colony was used to inoculate LB Broth and genomic DNA extracted from 1 mL using a Promega Wizard kit according to the manufacturer’s instructions (Promega, USA). DNA was quantified using the Qubit 3 and Nanodrop (Thermofisher, UK). Library preparation of genomic DNA was done using the LITE pipeline as described previously (20) and sequencing was performed using Nextseq (Illumina) with a Mid Output Flowcell (NSQ® 500 Mid Output KT v2). Read quality was assessed with fastp (21) and summarized with multiqc (22). Bracken (23) was used to assess the level of contamination. Sequences with a theoretical read depth below 20x, or with less than 80% of *Salmonella* reads were excluded from further analysis.

### Phylogenetic reconstruction

Maximum-likelihood phylogenetic trees were constructed from the core single-nucleotide polymorphism (SNP) alignment with reference to *S* Typhi strain CT18 (24) using snippy version 4.3.6 as previously described (25). The root node of trees was identified by including outgroups that were removed from final version of the tree (Supplementary table 2). RAxML (version 8.2.10) (26) was used to construct maximum likelihood phylogenetic trees from the core alignment, with the generalized time-reversible model and a Gamma distribution (GTR+Γ substitution GTRGAMMA in RAxML) to model site-specific rate variation. Support for the maximum-likelihood phylogeny was assessed with rapid bootstraps based on the MRE_IGN Bootstrapping criterion. For time-scaled phylogenetic trees, the 4.3.1.1EA1 subtree was extracted using the tree_subset function from treeio (27) and dating of nodes was performed using bactDating (28) using the root from the subtree. Strict gamma, relaxed gamma, mixed gamma, arc, carc and mixedcarc clock models were tested and compared using the BactDating modelcompare function. The arc model was used for the analysis as it showed the lowest deviance information criterion (DIC) (28).

### Identification of sequence polymorphisms, pangenome analysis and annotation

Antimicrobial resistance genes and plasmid replicons were identified using ARIBA version 2.14.6 (29) with the Plasmidfinder (version 1.2) (30) and ResFinder (version 3.1) (31) databases. Mutations in the *gyrA, gyrB* and *parC* chromosomal genes were detected using resistance gene identifier (RGI; version 5.1.1) (29). Genome assembly was carried out using SPAdes version 3.13.0 (32) with default parameters. The quality of the assembly was assessed with the quality assessment tool for genome assemblies QUAST version 5.0.2 (33). Assemblies larger than 5.5 MB were excluded from further analysis. Gene models and annotation was carried out using Prokka version 1.14.5 and Bandage (34, 35). For determination of pangenome and accessory genome, assembled and annotated genome sequences were used as input for Roary version 3.11.2 (36) to identify gene families and their distribution within *S*. Typhi isolates from Zimbabwe. The gene presence absence matrix was filtered to focus on genes present in at least 3 isolates (∼3%) and at most 90 isolates (∼95%).

Regions of interest were extracted from the relevant genomes for further analysis. Prophage annotation was done using a combination of Prokka to generate the gene model and by manual curation using the output from BLASTp derived annotation of the ORF against nr database (37). Nucleotide sequence BLAST results of prophage ZIM331 against the P88 reference (NC_026014) was visualized using genoPlotR (38).

## Results

### Two genotypes were circulating in Zimbabwe in 2018 with ongoing evolution of AMR in the dominant 4.3.1 (H58) genotype

Typhoid fever is a reportable disease in Zimbabwe with many of the cases in high density populations of urban Harare situated in north-eastern Zimbabwe (Figure 1A). To establish a convenience sample of *S*. Typhi strains prior to TCV vaccination, all available data regarding typhoid in Harare city health reporting systems between January 2018 and December 2018 were reviewed. In total, 3,946 suspected typhoid fever cases were reported in Zimbabwe (Figure 1B). An increase in suspected cases were reported from the first week of 2018 that peaked in the fourth week followed by a gradual decline from week 10 to week 18 (Figure 1C). An increase in overall daily cases was observed from week 45 to week 52 (Figure 1C). Of the 3,946 suspected typhoid fever cases 128 were confirmed by culture tests and 57 were randomly selected for whole genome sequencing and analysis.

**Figure 1.**
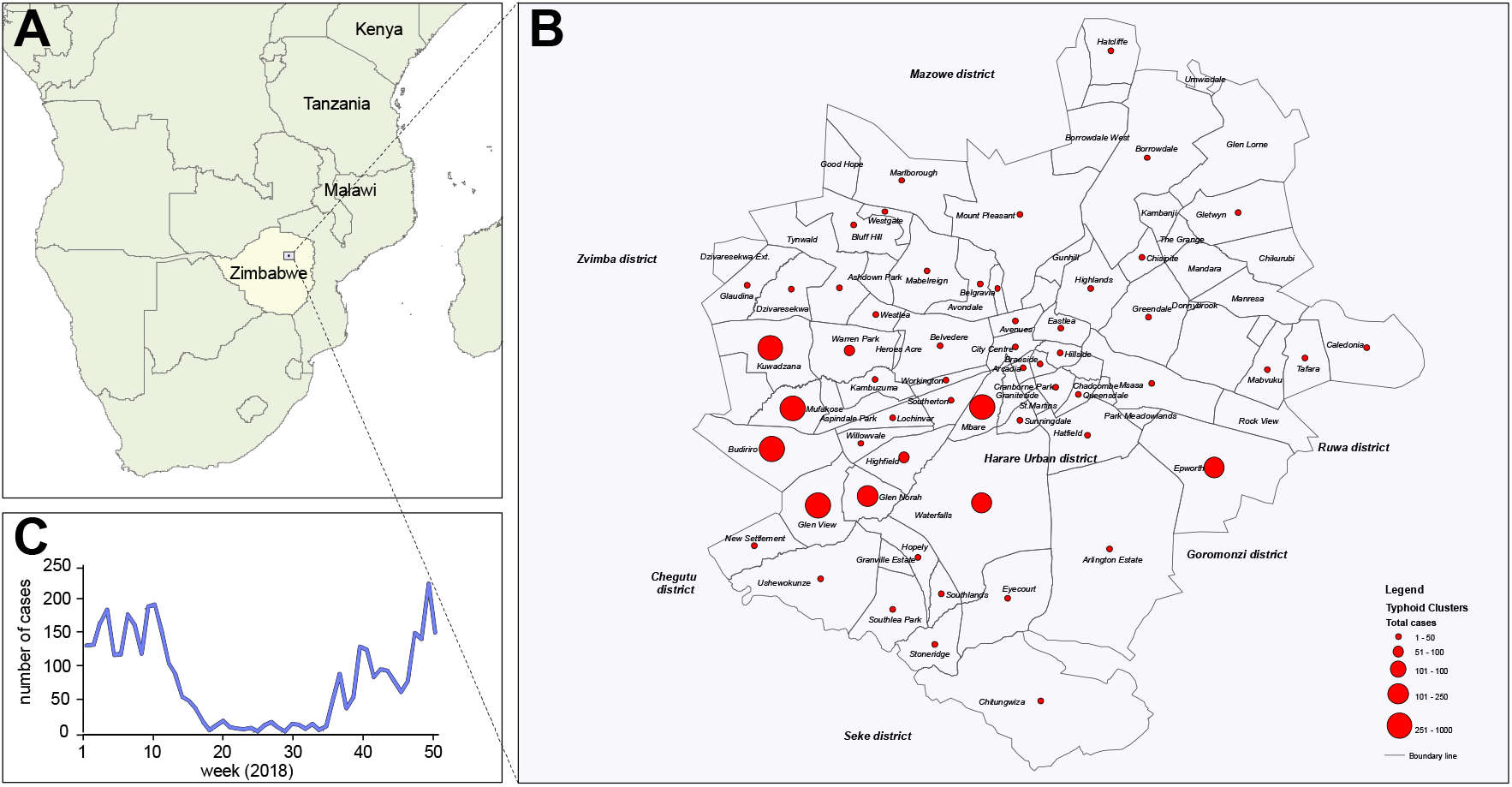
Epidemiology of typhoid fever in Harare, Zimbabwe in 2018. (A) Map of southern Africa showing the location of Zimbabwe and Harare. (B) Geographic distribution of suspected and confirmed cases of typhoid fever in Harare in 2018. Suburbs are indicated and the number of cases indicated (red circles) as indicated in the key (inset) (C) Daily number of suspected and confirmed typhoid cases (seven-day average) during the year 2018.

To investigate the phylogenetic relationship of *S*. Typhi isolated from clinical cases of typhoid fever in Zimbabwe, the whole genome sequence from a total of 95 isolates from typhoid fever infections in Zimbabwe between 2012 and 2019 were analyzed. These included 85 from infections in Zimbabwe (12) and 10 from clinical infections in the UK that were associated with travel to Zimbabwe (15). A maximum likelihood phylogenetic tree based on variation in the core genome sequence revealed a population structure for *S*. Typhi isolated from Zimbabwe consisting of two subclades corresponding to genotypes 4.3.1 (H58) (88/95, 93%) and 3.3.1 (7/95, 7%) (Figure 2). The *S*. Typhi isolates from the 2018 outbreak were present throughout the tree including genotypes of both 4.3.1.1 and 3.3.1 and were closely related to isolates from previous years and five isolates from 2019 (Figure 2). *Salmonella* Typhi isolates of genotype 4.3.1.1 (H58) encoded between six and ten AMR genes, while resistance genes were not detected in any isolates of genotype 3.3.1.

**Figure 2.**
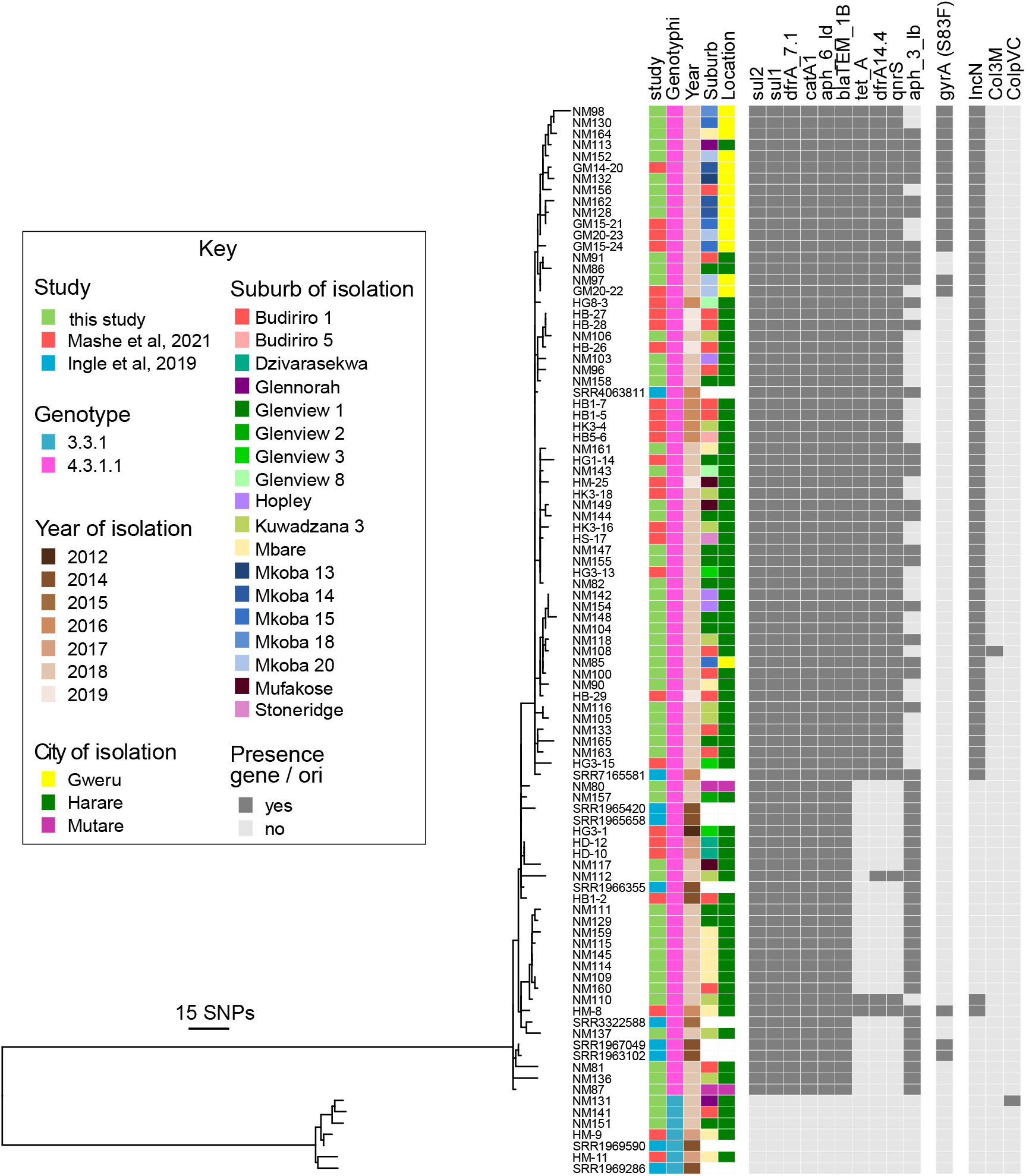
Phylogenetic relationship and genomic characteristics of 95 *S*. Typhi strains isolated from Harare and Gweru from 2012 to 2019. A maximum likelihood phylogenetic tree constructed using nucleotide sequence variation in the shared genome of 95 *S*. Typhi strains with reference to *S* Typhi CT18 whole genome sequence assembly (24) and rooted to the reference as an outgroup. Source of the sequence data (study), the genotype (genotyphi), year of isolation (year), and the city (location) and city suburb (suburb) are indicated by colors indicated in the key (inset). The approximate number of SNPs are indicated (bar).

All isolates of 4.3.1.1 (H58) had *aph-6, bla*_TEM-1B_, *dfr*A7.1, *catA1, sul1* and *sul2* genes conferring resistance to aminoglycosides, penicillin and older cephalosporins, trimethoprim, phenicols, and sulphonamides, respectively (Figure 2). A total of 62 isolates had three additional AMR genes, *tet*A, *dfr*A14 and *qnrS*, whose presence coincided with the detection of sequence from an IncN plasmid (subtype PST3). Most strains (60/62) with *tet*A, *dfr*A14 and *qnrS* genes were present in a single distal sub-clade within the Zimbabwe 4.3.1.1 population structure (blue sub-clade in Figure 2), with two isolates with this AMR profile situated in a more-basal rooted clade. All of the isolates with the *tet*A, *dfr*A14 and *qnrS* genes were isolated from 2016 to 2019. The *aph*3lb gene has a complex distribution within the 4.3.1.1 (H58) population in Zimbabwe consistent with multiple acquisitions or losses. While the *aph*3lb was present in all but one isolate in the basal-rooted clade, it was sporadically distributed within the distal clade containing the IncN plasmid. A total of 18 isolates (20%) contained a mutation in the *gyrA* gene predicted to result in a S83F substitution in GyrA, known to result in increased minimum inhibitory concentration for fluroquinolone antibiotics (39). GyrA S83F was present in two clusters of thirteen and two isolates within the distal clade containing the IncN plasmid, and three isolates from the basally rooted clade, all of which except two also had the *qnrS* gene.

### Analysis of the accessory genome of Zimbabwe endemic *S*. Typhi indicates clade specific clusters of genes

To investigate the clade-specific gene content of *S*. Typhi circulating in Zimbabwe we determined the pangenome of 97 isolates by comparing all predicted protein sequences using Roary software. Three large clusters of genes co-occurred at a similar frequency and phylogenetic distribution in either all genotype 4.3.1.1EA1 strains, a subset of 4.3.1.1EA1 or only 3.3.1, and were designated groups 1, 2 and 3, respectively (Figure 3). Group 1 contained genes present within a transposon containing the *aph-6, bla*_TEM-1B_, *dfr*A7.1 and *catA1* resistance genes, described previously (40, 41). Group 2 contained genes consistent with a plasmid including the IncN replicon and the *tetA, dfrA14* and *qnrS* genes. Alignment of the nucleotide sequence of group 3 genes to sequences in the NCBI database using BLAST identified multiple prophage genes (Figure 4). A putative prophage that we designated ZIM331 was most closely related to prophage P88 and consisted of 47 predicted coding sequences of which 36 had similarity to genes with functions associated with prophage functions and seven genes encoding hypothetical proteins of unknown function. A cluster of four genes were putative cargo genes and exhibited sequence similarity to *hxsDBCA*, a super-family of genes that encode proteins with diverse activity in metabolic processes (42).

**Figure 3.**
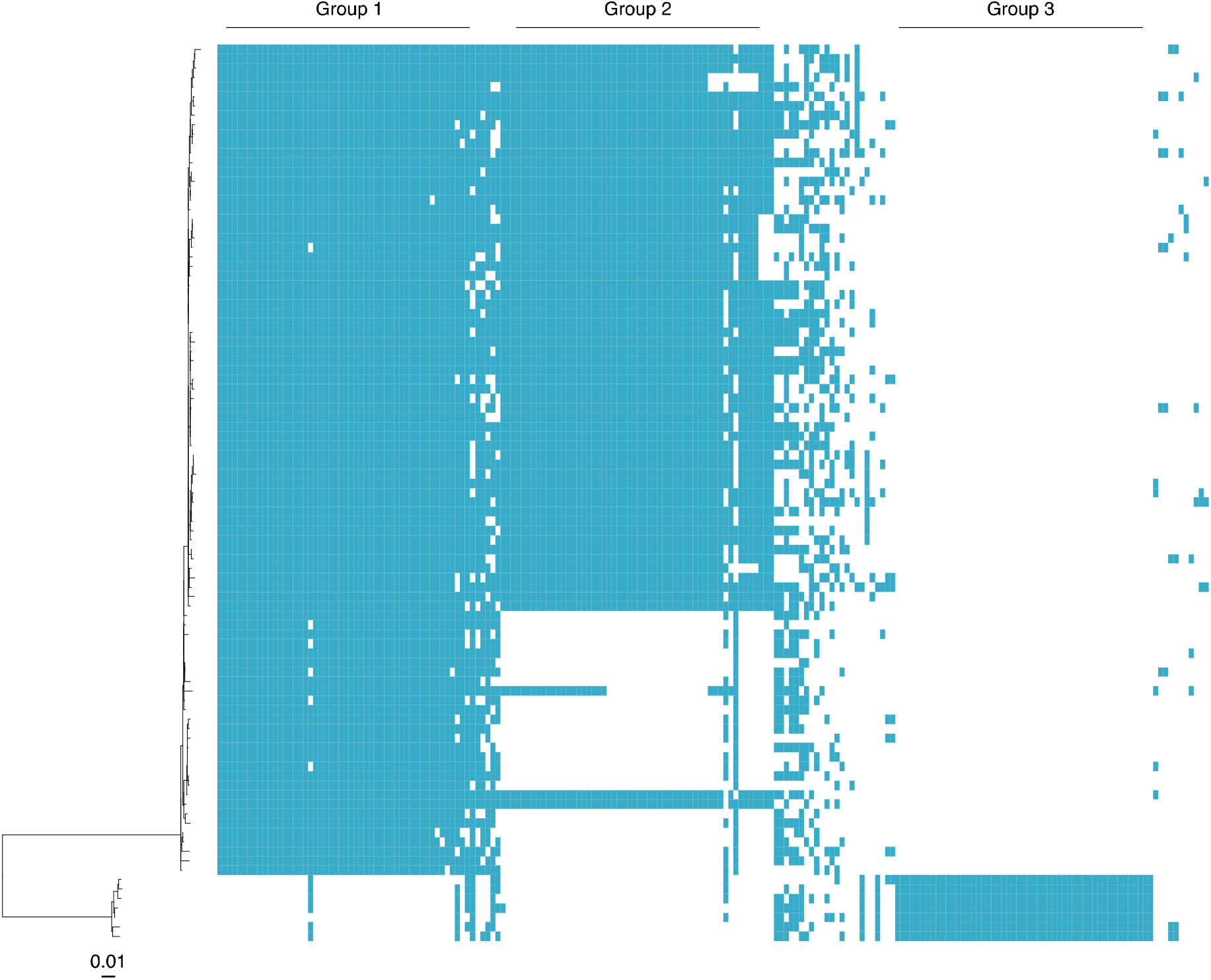
Accessory genome of 95 *S*. Typhi isolates from typhoid fever cases linked to Zimbabwe. Gene families (columns) present in each genome (blue) are arranged by frequency at which they occur in 95 genomes of *S*. Typhi strains isolated in Zimbabwe (n=85) or in the UK and associated with travel to Zimbabwe (n=10). Only genes present in greater than two isolates (∼3%) and less than 91 isolates (∼95%) are shown. Genes with a similar frequency and phylogenetic distribution were classified as group 1, 2 and 3 that correlate with genes present on a composite transposon, IncN plasmid and prophage element, respectively.

**Figure 4.**
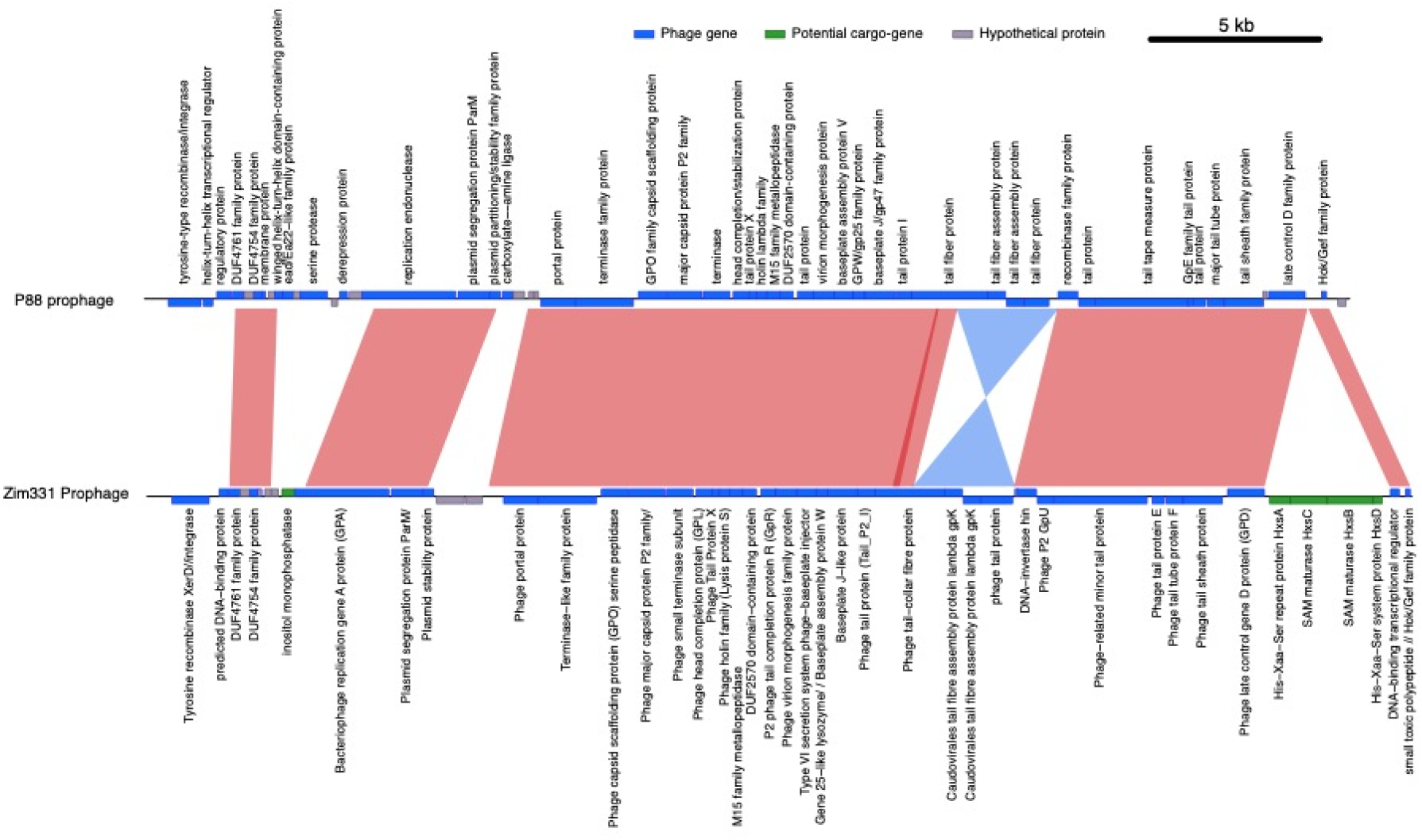
Comparison of prophage Zim331 with prophage P88. A gene model predicted using prokka show genes with predicted phage functions (blue bars), potential cargo genes (green bars) and hypothetical proteins with no known function (grey bars) based on sequence alignment in the NCBI database, are indicated for prophage P88 and prophage Zim331. Predicted function for proteins encoded by genes are indicated and regions exhibiting >90% sequence identity in direct alignment (red) or reverse and complement alignment (blue) are indicated.

### The 4.3.1.1 clone emerged in Zimbabwe around 2009 and acquired additional AMR genes on an IncN plasmid around 2012

To investigate the phylogenetic relationship of 95 *S*. Typhi isolates from Zimbabwe in the context of the global *S*. Typhi population structure we constructed a maximum likelihood tree including 1,904 *S*. Typhi isolates from 65 countries, described previously (43, 44) (Figure 5). A cluster of seven genotype 3.3.1 were closely related to other isolates from East and Southern African countries (Supplemental figure 1). The majority of the isolates belonged to clade 4 and in particular subclade 4.3 that include H58 (Figure 5). To further resolve the phylogenetic structure of isolates in clade 4.3 Zimbabwe and the global collection, a phylogenetic tree was constructed based on variation in the core genome sequence of clade 4.3 only (Figure 6). Genotype 4.3.1.1 isolates from Zimbabwe were present on a distal lineage within a subclade formed by isolates from East Africa and Southern Africa. The ladder topology of this part of the phylogenetic tree was consistent with multiple transmission events in a southernly direction from Kenya to Tanzania, Malawi and into Zimbabwe, followed by local spread.

**Figure 5.**
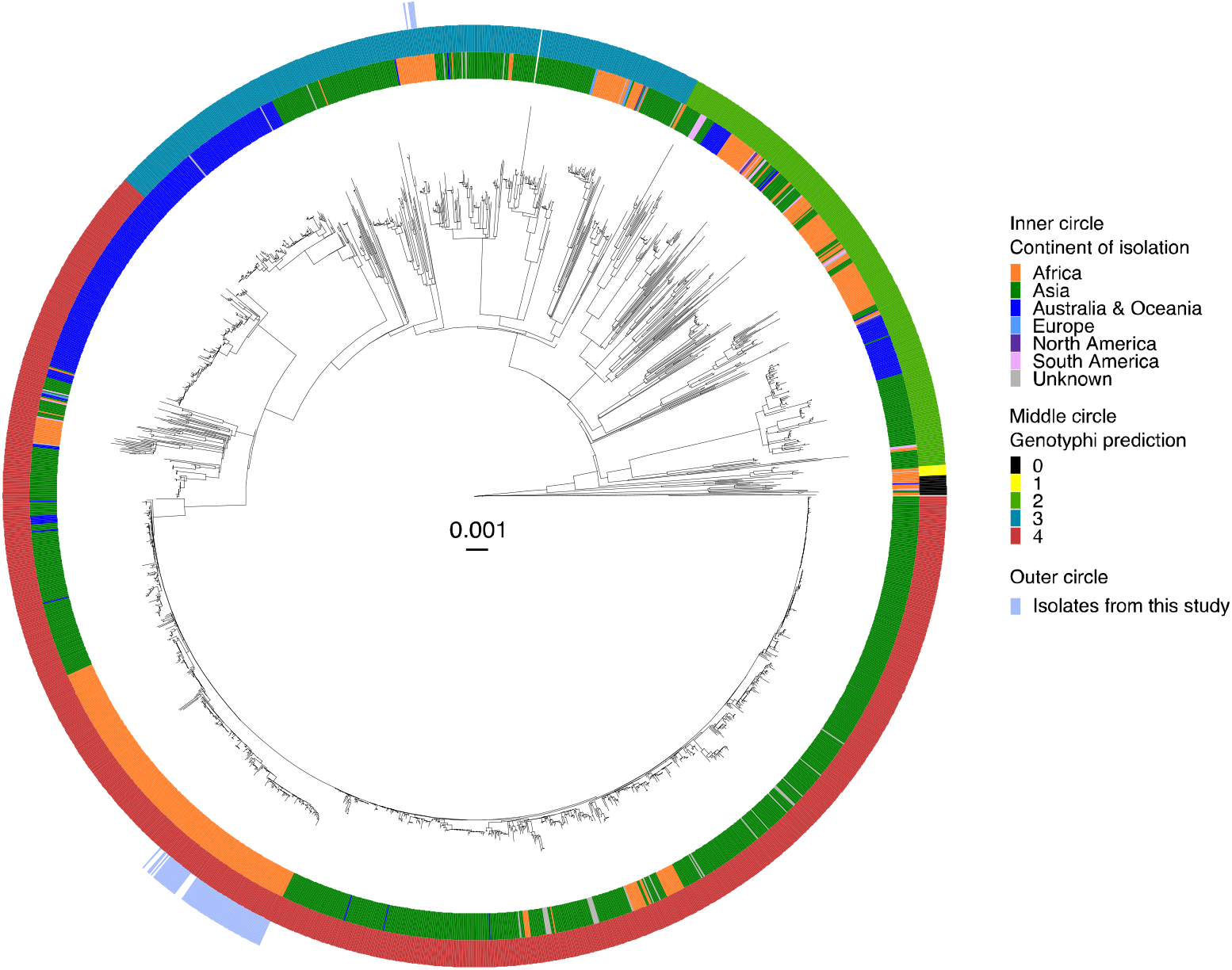
Phylogenetic relationship of 95 *S*. Typhi strains isolates from Zimbabwe in the context of 1904 *S*. Typhi strains isolated from globally dispersed locations. Maximum likelihood phylogenetic tree constructed based on variation in shared nucleotide sequence with reference to *S* Typhi CT18 whole genome sequence assembly (24). Continent of isolation (inner circle), genotype based on TyphiNET designation (middle circle) and isolates reported in this study (outer circle) are color coded as indicated in the key (inset).

**Figure 6.**
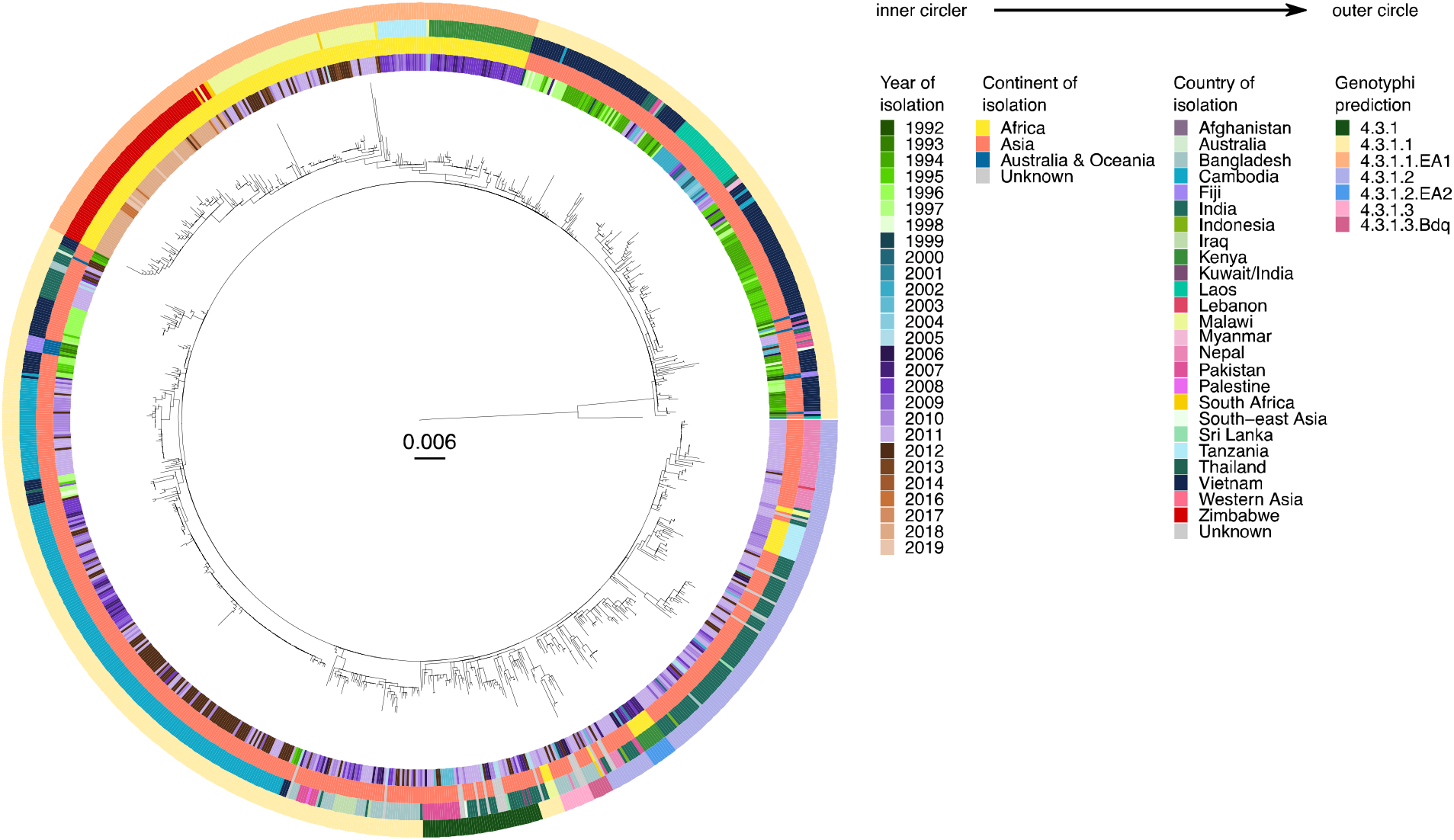
Phylogenetic relationship of genotype 4.3.1 *S*. Typhi strains isolates from Zimbabwe and globally dispersed locations. Maximum likelihood phylogenetic tree constructed based on variation in shared nucleotide sequence with reference to *S* Typhi CT18 whole genome sequence assembly (24). Year of isolation, continent of isolation, country of isolation and genotype based on TyphiNET designation are indicated in concentric circles color coded as indicated in the key (inset).

To estimate the time of key spread and evolutionary events associated with AMR in the emergence of the 4.3.1.1 Zimbabwe endemic clone, a subtree containing genomes from the 4.3.1.1EA1 subclade was extracted from the genotype 4.3 clade maximum likelihood tree. Linear regression analysis indicated a strong temporal signal for the accumulation of SNPs in the 4.3.1.1EA1 subtree that was absent if date of isolation was randomly assigned to taxa. (Supplemental Figure 2). A time-scaled tree constructed by Bayesian inference using BactDating (28) indicated that the common ancestor of the 4.3.1.1EA1 clade existed around 1987.236 [95% confidence interval 1977.5 - 1994.0] (Figure 7). Most of the deeply rooted isolates, that were from Kenya, had IncH1 replicon genes that correlated with the presence of *aph*6ld, *bla*_TEM_, *sul2, aph*3lb, *sul1, cat*A1, *dfrA*7.1 and *tetB* AMR genes. Isolates from Tanzania, Malawi, South Africa and Zimbabwe lacked the IncH1 replicon genes but most had *aph*6ld, *bla*_TEM_, *sul1, sul2, aph*3lb, *cat*A1, *dfrA*7.1 but not *tetB*. Fourteen of 20 isolates from Tanzania had IncFIB replicon genes and had lost *sul1, cat*A1, *dfrA*7.1 and *tetB*, but gained a *dfrA*14.4 gene. This was consistent with acquisition of the IncHI1 plasmid in Kenya followed by sporadic losses. Most isolates from Malawi, South Africa and Zimbabwe had the *aph*6ld, *bla*_TEM_, *sul2, aph*3lb, *sul1, cat*A1, *dfrA*7.1 and *tetB* AMR genes, despite lacking the IncHI1 replicon genes that coincided in isolates from Kenya. The exception was the sporadic apparent loss of the *aph*3lb gene from 30 of the 95 isolates from Zimbabwe, an event not observed in isolates from Kenya, Tanzania, Malawi or South Africa.

**Figure 7.**
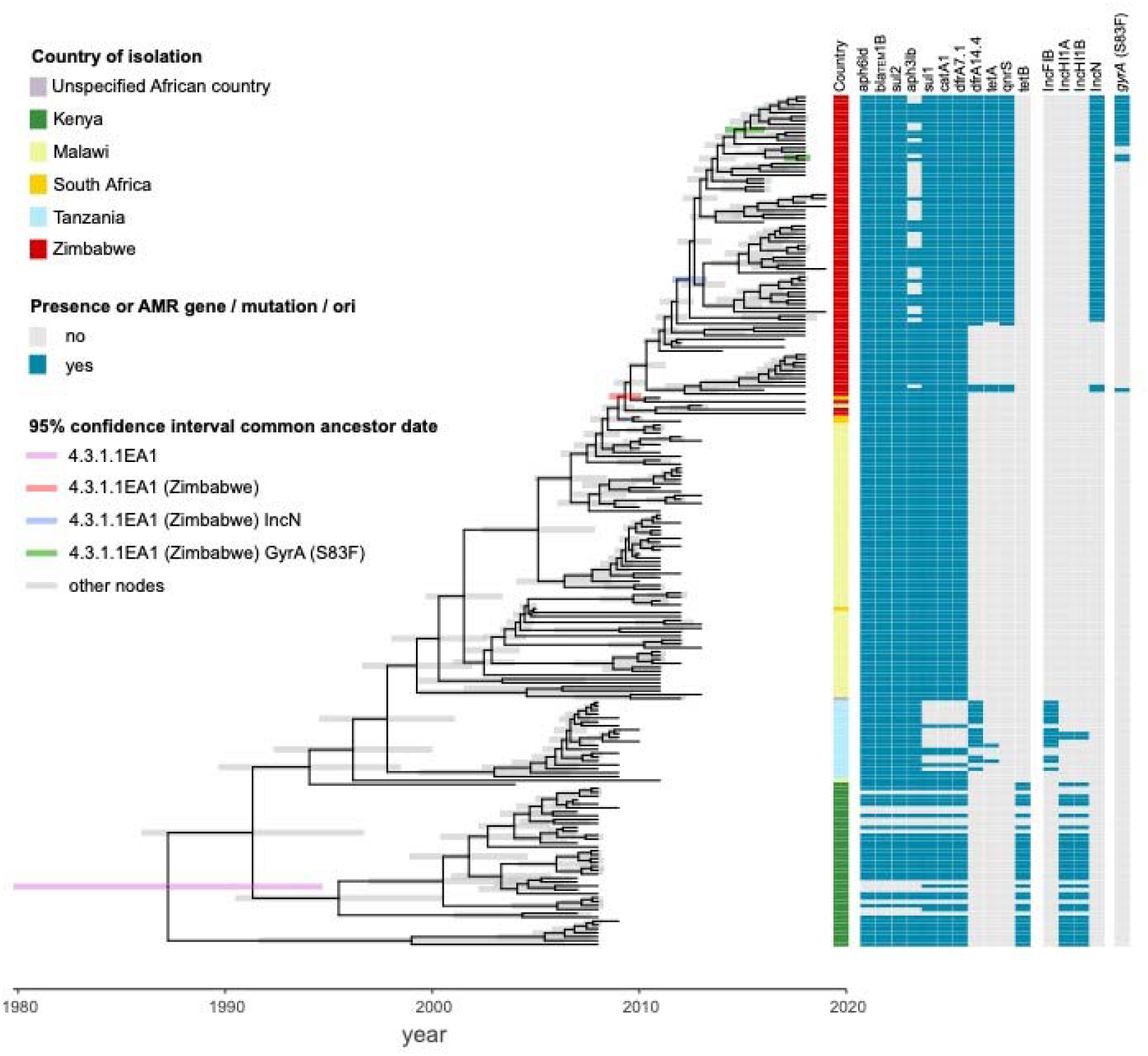
Time-scaled phylogenetic tree of genotype 4.3.1.1EA1 *S*. Typhi isolates. Terminal branch lengths are constrained to date of isolation and the 95% credibility interval is indicated by shaded bar color-coded to identify nodes corresponding to the common ancestor of the 4.3.1.1EA1 clade, 4.3.1.1EA1 isolated from Zimbabwe, 4.3.1.1EA1 from Zimbabwe carrying an IncN plasmid and 4.3.1.1EA1 from Zimbabwe with mutations in the QRDR of *gryA* as indicated in the key (inset). Country of isolation and presence absence of AMR and replicon are indicated in the key (inset).

The common ancestor of all *S*. Typhi 4.3.1.1EA1 isolates from Zimbabwe was estimated to have existed around 2009 [95% confidence interval: 2008.5 – 2010.0], consistent with epidemiological records indicating increased outbreaks of typhoid fever from this time (11, 13). Additional evolution of AMR was also exclusively observed in isolates from Zimbabwe with the apparent acquisition of *dfrA*14, *qnrS* and *tetA* AMR genes whose presence coincided with the presence of IncN replicon genes. The common ancestor of the IncN-containing isolates was around 2012 [95% confidence interval: 2011.5 – 2013.3]. Isolates with a *gyrA* mutation resulting in the S83F substitution associated with fluoroquinolone resistance shared a common ancestor or were isolated since around 2015.

## Discussion

Recurrent outbreaks of typhoid fever have been recorded in Zimbabwe since 2009 (13, 45). We found that during the period 2012 to 2019 strains of genotype 4.3.1.1, also known as H58, and genotype 3.3.1 were endemic. Both were likely to have been endemic during this time since they each formed clusters of closely related strains, 3.3.1 between 2014 and 2018 and genotype 4.3.1.1 throughout (13). The absence of 3.3.1 in 2012 and 2019 was likely due to the relatively small number of isolates investigated in these years. The vast majority of *S*. Typhi isolates from Zimbabwe in this study (88/95) were of the globally distributed genotype 4.3.1.1 that is characterized by multidrug resistance encoded on a transposon on an IncHI1 plasmid or incorporated into the chromosome (46). In contrast, we found that isolates of genotype 3.3.1 lacked AMR genes entirely.

All the *S*. Typhi genotype 4.3.1.1 isolates from Zimbabwe formed a discrete cluster within a subclade designated genotype 4.3.1.1EA1, composed entirely of isolates from east and southern African, or travel to this region (47). 4.3.1.1EA1 was in turn rooted within isolates from Southern Asia consistent with initial introduction from Southern Asia into Kenya and Tanzania and subsequent spread south into Malawi, as reported previously (44, 48). Our analyses confirm further transmission of this clone to Zimbabwe. The strong association of isolates from each country into distinct subclades within the genotype 4.3.1.1EA1 population structure suggests that spread resulted from a single transmission event into each country followed by local transmission of a clone. Multiple transmission events would be expected to result in a greater degree of mixing of isolates from each country in the phylogenetic tree, although additional analysis of more recent isolates from Kenya, Tanzania and Malawi may reveal other transmission. The common ancestor of all Zimbabwe isolates was around 2009 marking the earliest date for introduction of genotype 4.3.1.1EA1 into Zimbabwe. This coincides with reports of renewed outbreaks in Zimbabwe from this time for unknown reasons (45), but may be due to the arrival of this new genotype.

To date, genotype 3.3.1 isolates have garnered little attention compared to 4.3.1.1 as they are relatively rare and susceptible to antibiotics and consequently their global spread remains to be determined. A total of 34 isolates of genotype 3.3.1 were present in the global strain collection of 1904 whole genome sequences used in this study, while only 60 were available on TyphiNET out of 5,327 genomes (accessed August 2022) (49), suggesting that this genotype remains relatively rare globally, or under sampled. Nonetheless, the 34 genotype 3.3.1 isolates were from 10 different countries, with over 90% from East and Southern African (n=21) and Asian countries (n=10). Notably, in common with genotype 4.3.1.1EA1 isolates, genotype 3.3.1 isolates largely clustered based on the continent and the country of origin, consistent with international spread and subsequent domestic transmission of local clones. In contrast to genotype 4.3.1.1EA1, the topology of genotype 3.3.1 phylogeny consisted of country-specific clades with deeply rooted branches consistent with rapid initial spread globally and little current evidence of spread since. Additional genomes sequences are needed to investigate the time and phylogeographic spread in detail.

The evolution of *S*. Typhi strains with ever greater resistance to antimicrobials through acquisition of AMR genes or mutations in drug targets or efflux pumps is continuously reducing the options for therapy (46, 50). Our analysis of the evolution of the 4.3.1.1EA1 clade highlighted a concerning trend of increased resistance in Zimbabwe. Deeply rooted 4.3.1.1EA1 clades containing strains isolated before 2010 in Kenya were multidrug resistant due to AMR genes on an IncHI1 plasmid typical of genotype 4.3.1.1 isolates from South Asia (44). As 4.3.1.1EA1 spread south through Tanzania, isolates appear to have lost the IncHI1 plasmid but retained the AMR genes, likely due to their incorporation into the chromosome as previously described (44). The ancestral strain that spread to Zimbabwe around 2009 was of this genotype, but within 3 years an ancestor to nearly two thirds of isolates in the present study, had gained an IncN plasmid containing additional genes including the *qnrS* gene conferring resistance to quinolone antibiotics. The IncN plasmid is predicted to contain around 50 genes and it is possible that the energy cost of maintaining this plasmid may only have been favorable following the loss of the IncHI1 plasmid that contained up to 225 genes (51), but this remains to be investigated. Fluroquinolone resistance in *S*. Typhi is normally associated with mutations in the quinolone resistance determining region (QRDR) of GyrA and ParC. A previous study reported that QRDR mutations emerged independently on at least 94 occasions globally but almost exclusively in South Asia (48). We detected at least four independent acquisitions of QRDR mutations in the *gyrA* gene. Notably, two of the mutation events that accounted for 15 of 18 isolates also contained the *qnrS* gene on the IncN plasmid, suggesting that accumulation additional QRDR mutations may further increase fluoroquinolone.

In response to the 2018 ciprofloxacin-resistant typhoid outbreak, Zimbabwe carried out a mass typhoid vaccination campaign from February to March 2019 in nine suburbs of Harare with TCV. Over 318,000 doses were administered targeting children aged between 6 months and 15 years in affected communities. Previously, whole genome sequencing was retrospectively used to investigate the effect on the *S*. Typhi population in Thailand following a national immunization program in 1977 in response to a large outbreak (52). *S*. Typhi isolates from after the immunization program were found to be travel associated cases from neighboring countries. Our study provides a detailed insight into the emergence and baseline population structure of *S*. Typhi in Zimbabwe prior to the recent immunization program to enable assessment of the impact this program on the population structure of *S*. Typhi in the future.

## Author Contributions

GT, TM, RAK, MME designed the study. TM, BVC, VR, AT, TT, MMK, SM, LWM, JM acquired data. GT, TM, MB carried out analysis. GT, TM, RAK, MME interpreted the analysis. GT, TM, RAK drafted the manuscript. All authors critically reviewed the manuscript and approved the final version of the manuscript.

## Declaration of Interests

The authors declare no conflicts of interests arising from financial or personal relationships with other people or organizations.

**Supplemental Figure 1.**
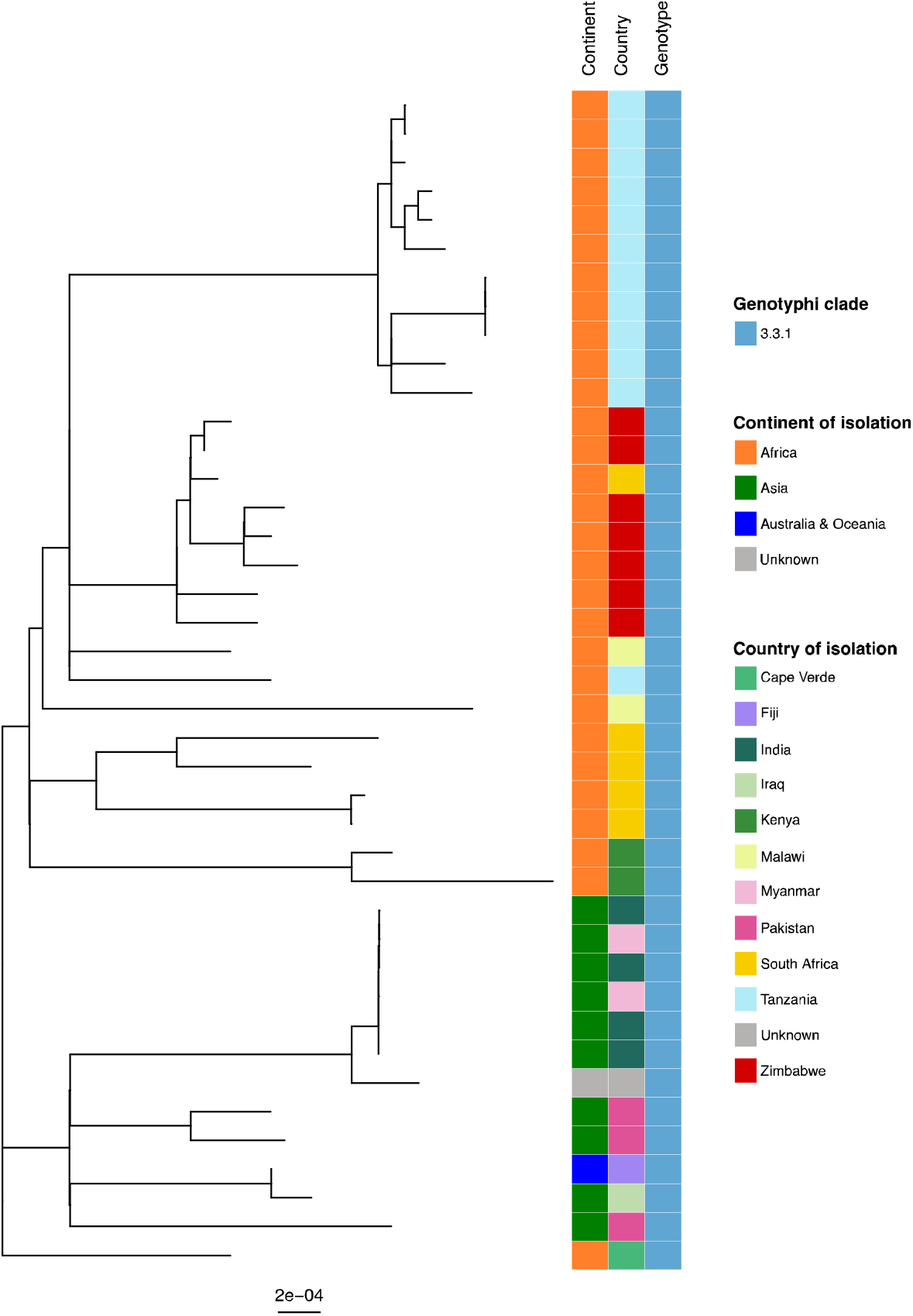
Phylogeny of the subclade 3.3.1 extracted from the global tree. Continent and Country of isolation are represented on a colour coded on as indicated on the key

**Supplemental Figure 2.**
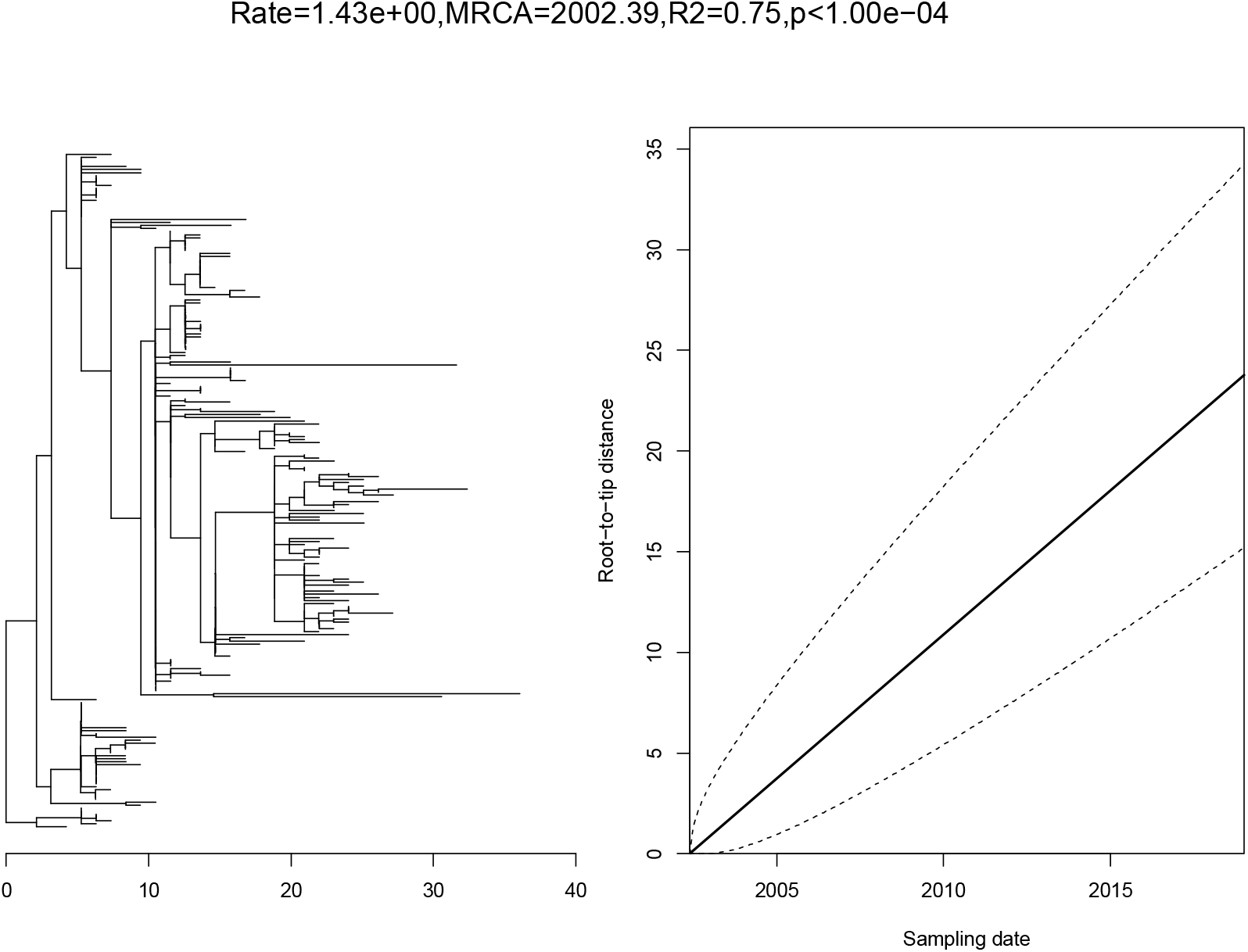
Root to tip regression analysis indicating temporal signal for the accumulation of SNPs within the 4.3.1.1EA1 clade.

## References

1. James SL, Abate D, Abate KH, Abay SM, Abbafati C, Abbasi N, et al. Global, regional, and national incidence, prevalence, and years lived with disability for 354 diseases and injuries for 195 countries and territories, 1990–2017: a systematic analysis for the Global Burden of Disease Study 2017. The Lancet. 2018;392(10159):1789–858.

2. Roth GA, Abate D, Abate KH, Abay SM, Abbafati C, Abbasi N, et al. Global, regional, and national age-sex-specific mortality for 282 causes of death in 195 countries and territories, 1980–2017: a systematic analysis for the Global Burden of Disease Study 2017. The Lancet. 2018;392(10159):1736–88.

3. Antillón M, Warren JL, Crawford FW, Weinberger DM, Kürüm E, Pak GD, et al. The burden of typhoid fever in low-and middle-income countries: a meta-regression approach. PLoS neglected tropical diseases. 2017;11(2):e0005376.

4. Mogasale V, Maskery B, Ochiai RL, Lee JS, Mogasale VV, Ramani E, et al. Burden of typhoid fever in low-income and middle-income countries: a systematic, literature-based update with risk-factor adjustment. The Lancet Global health. 2014;2(10):e570–e80.

5. Stanaway JD, Reiner RC, Blacker BF, Goldberg EM, Khalil IA, Troeger CE, et al. The global burden of typhoid and paratyphoid fevers: a systematic analysis for the Global Burden of Disease Study 2017. The Lancet Infectious Diseases. 2019;19(4):369–81.

6. Andrews JR, Qamar FN, Charles RC, Ryan ET. Extensively drug-resistant typhoid— are conjugate vaccines arriving just in time? New England Journal of Medicine. 2018;379(16):1493–5.

7. Marchello CS, Hong CY, Crump JA. Global Typhoid Fever Incidence: A Systematic Review and Meta-analysis. Clinical Infectious Diseases. 2019;68(Supplement_2):S105–S16.

8. Klemm EJ, Shakoor S, Page AJ, Qamar FN, Judge K, Saeed DK, et al. Emergence of an extensively drug-resistant Salmonella enterica serovar Typhi clone harboring a promiscuous plasmid encoding resistance to fluoroquinolones and third-generation cephalosporins. MBio. 2018;9(1):e00105–18.

9. Vekemans J, Hasso-Agopsowicz M, Kang G, Hausdorff WP, Fiore A, Tayler E, et al. Leveraging Vaccines to Reduce Antibiotic Use and Prevent Antimicrobial Resistance: A World Health Organization Action Framework. Clin Infect Dis. 2021;73(4):e1011–e7.

10. N’Cho H S, Masunda KPE, Mukeredzi I, Manangazira P, Govore E, Duri C, et al. Notes from the Field: Typhoid Fever Outbreak - Harare, Zimbabwe, October 2017-February 2018. MMWR Morb Mortal Wkly Rep. 2019;68(2):44–5.

11. Lightowler MS, Manangazira P, Nackers F, Van Herp M, Phiri I, Kuwenyi K, et al. Effectiveness of typhoid conjugate vaccine in Zimbabwe used in response to an outbreak among children and young adults: A matched case control study. Vaccine. 2022;40(31):4199–210.

12. Mashe T, Leekitcharoenphon P, Mtapuri-Zinyowera S, Kingsley RA, Robertson V, Tarupiwa A, et al. Salmonella enterica serovar Typhi H58 clone has been endemic in Zimbabwe from 2012 to 2019. Journal of Antimicrobial Chemotherapy. 2020.

13. Mashe T, Leekitcharoenphon P, Mtapuri-Zinyowera S, Kingsley RA, Robertson V, Tarupiwa A, et al. Salmonella enterica serovar Typhi H58 clone has been endemic in Zimbabwe from 2012 to 2019. J Antimicrob Chemother. 2021;76(5):1160–7.

14. Yap K-P, Ho WS, Gan HM, Chai LC, Thong KL. Global MLST of Salmonella Typhi Revisited in Post-genomic Era: Genetic Conservation, Population Structure, and Comparative Genomics of Rare Sequence Types. Frontiers in Microbiology. 2016;7(270).

15. Ingle DJ, Nair S, Hartman H, Ashton PM, Dyson ZA, Day M, et al. Informal genomic surveillance of regional distribution of Salmonella Typhi genotypes and antimicrobial resistance via returning travellers. PLoS neglected tropical diseases. 2019;13(9):e0007620.

16. Afema JA, Byarugaba DK, Shah DH, Atukwase E, Nambi M, Sischo WM. Potential sources and transmission of Salmonella and antimicrobial resistance in Kampala, Uganda. PLoS One. 2016;11(3):e0152130.

17. Holt KE, Parkhill J, Mazzoni CJ, Roumagnac P, Weill FX, Goodhead I, et al. High-throughput sequencing provides insights into genome variation and evolution in Salmonella Typhi. Nat Genet. 2008;40(8):987–93.

18. Hudzicki J. Kirby-Bauer disk diffusion susceptibility test protocol. 2009.

19. Wayne P. Clinical and Laboratory Standards Institute (CLSI); 2010. Performance standards for antimicrobial susceptibility testing. 2010;20.

20. Kirkwood M, Vohra P, Bawn M, Thilliez G, Pye H, Tanner J, et al. Ecological niche adaptation of Salmonella Typhimurium U288 is associated with altered pathogenicity and reduced zoonotic potential. Commun Biol. 2021;4(1):498.

21. Chen S, Zhou Y, Chen Y, Gu J. fastp: an ultra-fast all-in-one FASTQ preprocessor. Bioinformatics. 2018;34(17):i884–i90.

22. Ewels P, Magnusson M, Lundin S, Käller M. MultiQC: summarize analysis results for multiple tools and samples in a single report. Bioinformatics. 2016;32(19):3047–8.

23. Lu J, Breitwieser FP, Thielen P, Salzberg SL. Bracken: estimating species abundance in metagenomics data. PeerJ Computer Science. 2017;3:e104.

24. Parkhill J, Dougan G, James K, Thomson N, Pickard D, Wain J, et al. Complete genome sequence of a multiple drug resistant Salmonella enterica serovar Typhi CT18. Nature. 2001;413(6858):848.

25. Branchu P, Charity OJ, Bawn M, Thilliez G, Dallman TJ, Petrovska L, et al. SGI-4 in Monophasic Salmonella Typhimurium ST34 Is a Novel ICE That Enhances Resistance to Copper. Frontiers in microbiology. 2019; 10:[1118 p.].

26. Stamatakis A. RAxML-VI-HPC: maximum likelihood-based phylogenetic analyses with thousands of taxa and mixed models. Bioinformatics. 2006;22(21):2688–90.

27. Yu G, Smith DK, Zhu H, Guan Y, Lam TT-Y, McInerny G. ggtree: anrpackage for visualization and annotation of phylogenetic trees with their covariates and other associated data. Methods in Ecology and Evolution. 2017;8(1):28–36.

28. Didelot X, Croucher NJ, Bentley SD, Harris SR, Wilson DJ. Bayesian inference of ancestral dates on bacterial phylogenetic trees. Nucleic acids research. 2018;46(22):e134–e.

29. Hunt M, Mather AE, Sánchez-Busó L, Page AJ, Parkhill J, Keane JA, et al. ARIBA: rapid antimicrobial resistance genotyping directly from sequencing reads. Microbial genomics. 2017;3(10):e000131–e.

30. Carattoli A, Zankari E, García-Fernández A, Larsen MV, Lund O, Villa L, et al. In Silico Detection and Typing of Plasmids using PlasmidFinder and Plasmid Multilocus Sequence Typing. Antimicrobial Agents and Chemotherapy. 2014;58(7):3895–903.

31. Zankari E, Hasman H, Cosentino S, Vestergaard M, Rasmussen S, Lund O, et al. Identification of acquired antimicrobial resistance genes. Journal of antimicrobial chemotherapy. 2012;67(11):2640–4.

32. Bankevich A, Nurk S, Antipov D, Gurevich AA, Dvorkin M, Kulikov AS, et al. SPAdes: a new genome assembly algorithm and its applications to single-cell sequencing. Journal of computational biology. 2012;19(5):455–77.

33. Gurevich A, Saveliev V, Vyahhi N, Tesler G. QUAST: quality assessment tool for genome assemblies. Bioinformatics. 2013;29(8):1072–5.

34. Wick RR, Schultz MB, Zobel J, Holt KE. Bandage: interactive visualization of de novo genome assemblies. Bioinformatics. 2015;31(20):3350–2.

35. Seemann T. Prokka: rapid prokaryotic genome annotation. Bioinformatics. 2014;30(14):2068–9.

36. Page AJ, Cummins CA, Hunt M, Wong VK, Reuter S, Holden MTG, et al. Roary: rapid large-scale prokaryote pan genome analysis. Bioinformatics. 2015;31(22):3691–3.

37. Camacho C, Coulouris G, Avagyan V, Ma N, Papadopoulos J, Bealer K, et al. BLAST+: architecture and applications. BMC Bioinformatics. 2009;10:421.

38. Guy L, Kultima JR, Andersson SG. genoPlotR: comparative gene and genome visualization in R. Bioinformatics. 2010;26(18):2334–5.

39. Turner AK, Nair S, Wain J. The acquisition of full fluoroquinolone resistance in Salmonella Typhi by accumulation of point mutations in the topoisomerase targets. Journal of Antimicrobial Chemotherapy. 2006;58(4):733–40.

40. Chiou CS, Lauderdale TL, Phung DC, Watanabe H, Kuo JC, Wang PJ, et al. Antimicrobial resistance in Salmonella enterica Serovar Typhi isolates from Bangladesh, Indonesia, Taiwan, and Vietnam. Antimicrob Agents Chemother. 2014;58(11):6501–7.

41. Hendriksen RS, Leekitcharoenphon P, Lukjancenko O, Lukwesa-Musyani C, Tambatamba B, Mwaba J, et al. Genomic signature of multidrug-resistant Salmonella enterica serovar typhi isolates related to a massive outbreak in Zambia between 2010 and 2012. J Clin Microbiol. 2015;53(1):262–72.

42. Sofia HJ, Chen G, Hetzler BG, Reyes-Spindola JF, Miller NE. Radical SAM, a novel protein superfamily linking unresolved steps in familiar biosynthetic pathways with radical mechanisms: functional characterization using new analysis and information visualization methods. Nucleic Acids Res. 2001;29(5):1097–106.

43. Wong VK, Baker S, Pickard DJ, Parkhill J, Page AJ, Feasey NA, et al. Phylogeographical analysis of the dominant multidrug-resistant H58 clade of Salmonella Typhi identifies inter-and intracontinental transmission events. Nature genetics. 2015;47(6):632.

44. Wong VK, Baker S, Pickard DJ, Parkhill J, Page AJ, Feasey NA, et al. Phylogeographical analysis of the dominant multidrug-resistant H58 clade of Salmonella Typhi identifies inter- and intracontinental transmission events. Nat Genet. 2015;47(6):632–9.

45. Polonsky JA, Martinez-Pino I, Nackers F, Chonzi P, Manangazira P, Van Herp M, et al. Descriptive epidemiology of typhoid fever during an epidemic in Harare, Zimbabwe, 2012. PLoS One. 2014;9(12):e114702.

46. Klemm EJ, Shakoor S, Page AJ, Qamar FN, Judge K, Saeed DK, et al. Emergence of an Extensively Drug-Resistant Salmonella enterica Serovar Typhi Clone Harboring a Promiscuous Plasmid Encoding Resistance to Fluoroquinolones and Third-Generation Cephalosporins. mBio. 2018;9(1).

47. Kariuki S, Dyson ZA, Mbae C, Ngetich R, Kavai SM, Wairimu C, et al. Multiple introductions of multidrug-resistant typhoid associated with acute infection and asymptomatic carriage, Kenya. Elife. 2021;10.

48. da Silva KE, Tanmoy AM, Pragasam AK, Iqbal J, Sajib MSI, Mutreja A, et al. The international and intercontinental spread and expansion of antimicrobial-resistant Salmonella Typhi: a genomic epidemiology study. Lancet Microbe. 2022.

49. Argimon S, Yeats CA, Goater RJ, Abudahab K, Taylor B, Underwood A, et al. A global resource for genomic predictions of antimicrobial resistance and surveillance of Salmonella Typhi at pathogenwatch. Nat Commun. 2021;12(1):2879.

50. Dyson ZA, Klemm EJ, Palmer S, Dougan G. Antibiotic Resistance and Typhoid. Clin Infect Dis. 2019;68(Suppl 2):S165–S70.

51. Holt KE, Thomson NR, Wain J, Phan MD, Nair S, Hasan R, et al. Multidrug-resistant Salmonella enterica serovar paratyphi A harbors IncHI1 plasmids similar to those found in serovar typhi. J Bacteriol. 2007;189(11):4257–64.

52. Dyson ZA, Thanh DP, Bodhidatta L, Mason CJ, Srijan A, Rabaa MA, et al. Whole Genome Sequence Analysis of Salmonella Typhi Isolated in Thailand before and after the Introduction of a National Immunization Program. PLOS Neglected Tropical Diseases. 2017;11(1):e0005274.

