## Supplementary Table 1 for "Population structure and evolution of *Salmonella enterica* serotype Typhi in Zimbabwe before a typhoid conjugate vaccine immunization campaign"

| SRA_samp_id | Read_ID | Sample_id | Genotyphi | Age | Sex | Specimen | Year | Continent | Region | Country | Town | Suburb | reference |
|---|---|---|---|---|---|---|---|---|---|---|---|---|---|
| QIB-NM100 | SRR20666403 | NM100 | 4.3.1.1.EA1 | 21 | M | Blood | 2018 | Africa | Southern Africa | Zimbabwe | Harare | Budiriro 1 | This Study |
| QIB-NM105 | SRR20666402 | NM105 | 4.3.1.1.EA1 | 22 | M | Stool | 2018 | Africa | Southern Africa | Zimbabwe | Harare | Kuwadzana 3 | This Study |
| QIB-NM106 | SRR20666391 | NM106 | 4.3.1.1.EA1 | 56 | F | Stool | 2018 | Africa | Southern Africa | Zimbabwe | Harare | Kuwadzana 3 | This Study |
| QIB-NM108 | SRR20666380 | NM108 | 4.3.1.1.EA1 | 12 | F | Blood | 2018 | Africa | Southern Africa | Zimbabwe | Harare | Budiriro 1 | This Study |
| QIB-NM113 | SRR20666369 | NM113 | 4.3.1.1.EA1 | 11 | M | Blood | 2018 | Africa | Southern Africa | Zimbabwe | Harare | Glennorah | This Study |
| QIB-NM114 | SRR20666358 | NM114 | 4.3.1.1.EA1 | 21 | F | Blood | 2018 | Africa | Southern Africa | Zimbabwe | Harare | Mbare | This Study |
| QIB-NM115 | SRR20666350 | NM115 | 4.3.1.1.EA1 | 43 | M | Stool | 2018 | Africa | Southern Africa | Zimbabwe | Harare | Mbare | This Study |
| QIB-NM116 | SRR20666349 | NM116 | 4.3.1.1.EA1 | 21 | M | Blood | 2018 | Africa | Southern Africa | Zimbabwe | Harare | Kuwadzana 3 | This Study |
| QIB-NM118 | SRR20666348 | NM118 | 4.3.1.1.EA1 | 33 | F | Blood | 2018 | Africa | Southern Africa | Zimbabwe | Harare | Kuwadzana 3 | This Study |
| QIB-NM98 | SRR20666347 | NM98 | 4.3.1.1.EA1 | 36 | F | Blood | 2018 | Africa | Southern Africa | Zimbabwe | Gweru | Mkoba 18 | This Study |
| QIB-NM80 | SRR20666401 | NM80 | 4.3.1.1.EA1 | 35 | M | Stool | 2018 | Africa | Southern Africa | Zimbabwe | Mutare | Mutare | This Study |
| QIB-NM81 | SRR20666400 | NM81 | 4.3.1.1.EA1 | 17 | M | Blood | 2018 | Africa | Southern Africa | Zimbabwe | Harare | Budiriro 1 | This Study |
| QIB-NM82 | SRR20666399 | NM82 | 4.3.1.1.EA1 | 9 | F | Stool | 2018 | Africa | Southern Africa | Zimbabwe | Harare | Glenview 1 | This Study |
| QIB-NM85 | SRR20666398 | NM85 | 4.3.1.1.EA1 | 25 | F | Blood | 2018 | Africa | Southern Africa | Zimbabwe | Gweru | Mkoba 15 | This Study |
| QIB-NM87 | SRR20666397 | NM87 | 4.3.1.1.EA1 | 11 | M | Stool | 2018 | Africa | Southern Africa | Zimbabwe | Mutare | Mutare | This Study |
| QIB-NM90 | SRR20666396 | NM90 | 4.3.1.1.EA1 | 34 | M | Stool | 2018 | Africa | Southern Africa | Zimbabwe | Harare | Mbare | This Study |
| QIB-NM91 | SRR20666395 | NM91 | 4.3.1.1.EA1 | 70 | F | Blood | 2018 | Africa | Southern Africa | Zimbabwe | Harare | Budiriro 1 | This Study |
| QIB-NM96 | SRR20666394 | NM96 | 4.3.1.1.EA1 | 21 | M | Blood | 2018 | Africa | Southern Africa | Zimbabwe | Harare | Budiriro 1 | This Study |
| QIB-NM97 | SRR20666393 | NM97 | 4.3.1.1.EA1 | 35 | M | Blood | 2018 | Africa | Southern Africa | Zimbabwe | Gweru | Mkoba 20 | This Study |
| QIB-NM103 | SRR20666392 | NM103 | 4.3.1.1.EA1 | 16 | M | Blood | 2018 | Africa | Southern Africa | Zimbabwe | Harare | Hopley | This Study |
| QIB-NM104 | SRR20666390 | NM104 | 4.3.1.1.EA1 | 45 | F | Blood | 2018 | Africa | Southern Africa | Zimbabwe | Harare | Glenview 1 | This Study |
| QIB-NM109 | SRR20666389 | NM109 | 4.3.1.1.EA1 | 21 | M | Blood | 2018 | Africa | Southern Africa | Zimbabwe | Harare | Mbare | This Study |
| QIB-NM110 | SRR20666388 | NM110 | 4.3.1.1.EA1 | 11 | F | Stool | 2018 | Africa | Southern Africa | Zimbabwe | Harare | Kuwadzana 3 | This Study |
| QIB-NM111 | SRR20666387 | NM111 | 4.3.1.1.EA1 | 22 | M | Stool | 2018 | Africa | Southern Africa | Zimbabwe | Harare | Glenview 1 | This Study |
| QIB-NM112 | SRR20666386 | NM112 | 4.3.1.1.EA1 | 40 | F | Blood | 2018 | Africa | Southern Africa | Zimbabwe | Harare | Kuwadzana 3 | This Study |
| QIB-NM117 | SRR20666385 | NM117 | 4.3.1.1.EA1 | 55 | F | Blood | 2018 | Africa | Southern Africa | Zimbabwe | Harare | Mufakose | This Study |
| QIB-NM128 | SRR20666384 | NM128 | 4.3.1.1.EA1 | 17 | M | Stool | 2018 | Africa | Southern Africa | Zimbabwe | Gweru | Mkoba 14 | This Study |
| QIB-NM129 | SRR20666383 | NM129 | 4.3.1.1.EA1 | 28 | F | Blood | 2018 | Africa | Southern Africa | Zimbabwe | Harare | Glenview 1 | This Study |
| QIB-NM130 | SRR20666382 | NM130 | 4.3.1.1.EA1 | 21 | M | Blood | 2018 | Africa | Southern Africa | Zimbabwe | Gweru | Mkoba 15 | This Study |
| QIB-NM132 | SRR20666381 | NM132 | 4.3.1.1.EA1 | 26 | F | Blood | 2018 | Africa | Southern Africa | Zimbabwe | Gweru | Mkoba 13 | This Study |
| QIB-NM133 | SRR20666379 | NM133 | 4.3.1.1.EA1 | 31 | M | Blood | 2018 | Africa | Southern Africa | Zimbabwe | Harare | Budiriro 1 | This Study |
| QIB-NM136 | SRR20666378 | NM136 | 4.3.1.1.EA1 | 29 | F | Blood | 2018 | Africa | Southern Africa | Zimbabwe | Harare | Kuwadzana 3 | This Study |
| QIB-NM137 | SRR20666377 | NM137 | 4.3.1.1.EA1 | 27 | M | Blood | 2018 | Africa | Southern Africa | Zimbabwe | Harare | Kuwadzana 3 | This Study |
| QIB-NM142 | SRR20666376 | NM142 | 4.3.1.1.EA1 | 23 | F | Blood | 2018 | Africa | Southern Africa | Zimbabwe | Harare | Hopley | This Study |
| QIB-NM143 | SRR20666375 | NM143 | 4.3.1.1.EA1 | 12 | M | Blood | 2018 | Africa | Southern Africa | Zimbabwe | Harare | Glenview 8 | This Study |
| QIB-NM144 | SRR20666374 | NM144 | 4.3.1.1.EA1 | 36 | F | Stool | 2018 | Africa | Southern Africa | Zimbabwe | Harare | Glenview 1 | This Study |
| QIB-NM145 | SRR20666373 | NM145 | 4.3.1.1.EA1 | 21 | M | Blood | 2018 | Africa | Southern Africa | Zimbabwe | Harare | Mbare | This Study |
| QIB-NM147 | SRR20666372 | NM147 | 4.3.1.1.EA1 | 49 | F | Blood | 2018 | Africa | Southern Africa | Zimbabwe | Harare | Glenview 1 | This Study |
| QIB-NM148 | SRR20666371 | NM148 | 4.3.1.1.EA1 | 37 | F | Blood | 2018 | Africa | Southern Africa | Zimbabwe | Harare | Glenview 1 | This Study |
| QIB-NM149 | SRR20666370 | NM149 | 4.3.1.1.EA1 | 39 | F | Blood | 2018 | Africa | Southern Africa | Zimbabwe | Harare | Mufakose | This Study |
| QIB-NM152 | SRR20666368 | NM152 | 4.3.1.1.EA1 | 27 | M | Stool | 2018 | Africa | Southern Africa | Zimbabwe | Gweru | Mkoba 20 | This Study |
| QIB-NM154 | SRR20666367 | NM154 | 4.3.1.1.EA1 | 20 | F | Blood | 2018 | Africa | Southern Africa | Zimbabwe | Harare | Hopley | This Study |
| QIB-NM155 | SRR20666366 | NM155 | 4.3.1.1.EA1 | 19 | M | Blood | 2018 | Africa | Southern Africa | Zimbabwe | Harare | Glenview 1 | This Study |
| QIB-NM156 | SRR20666365 | NM156 | 4.3.1.1.EA1 | 14 | M | Stool | 2018 | Africa | Southern Africa | Zimbabwe | Gweru | Budiriro 1 | This Study |
| QIB-NM157 | SRR20666364 | NM157 | 4.3.1.1.EA1 | 14 | M | Stool | 2018 | Africa | Southern Africa | Zimbabwe | Harare | Glenview 2 | This Study |
| QIB-NM158 | SRR20666363 | NM158 | 4.3.1.1.EA1 | 36 | F | Blood | 2018 | Africa | Southern Africa | Zimbabwe | Harare | Glenview 1 | This Study |
| QIB-NM159 | SRR20666362 | NM159 | 4.3.1.1.EA1 | 59 | F | Stool | 2018 | Africa | Southern Africa | Zimbabwe | Harare | Mbare | This Study |
| QIB-NM160 | SRR20666361 | NM160 | 4.3.1.1.EA1 | 38 | M | Stool | 2018 | Africa | Southern Africa | Zimbabwe | Harare | Budiriro 1 | This Study |
| QIB-NM161 | SRR20666360 | NM161 | 4.3.1.1.EA1 | 50 | F | Blood | 2018 | Africa | Southern Africa | Zimbabwe | Harare | Mbare | This Study |

| | | | | | | | | | | | | | |
|---|---|---|---|---|---|---|---|---|---|---|---|---|---|
| QIB-NM162 | SRR20666359 | NM162 | 4.3.1.1.EA1 | 16 | M | Blood | 2018 | Africa | Southern Africa | Zimbabwe | Gweru | Mkoba 14 | This Study |
| QIB-NM163 | SRR20666357 | NM163 | 4.3.1.1.EA1 | 25 | M | Blood | 2018 | Africa | Southern Africa | Zimbabwe | Harare | Budiriro 1 | This Study |
| QIB-NM164 | SRR20666356 | NM164 | 4.3.1.1.EA1 | 44 | M | Blood | 2018 | Africa | Southern Africa | Zimbabwe | Gweru | Mbare | This Study |
| QIB-NM165 | SRR20666355 | NM165 | 4.3.1.1.EA1 | 26 | M | Blood | 2018 | Africa | Southern Africa | Zimbabwe | Harare | Glenview 1 | This Study |
| QIB-NM86 | SRR20666354 | NM86 | 4.3.1.1.EA1 | 22 | M | Blood | 2018 | Africa | Southern Africa | Zimbabwe | Harare | Glenview 1 | This Study |
| QIB-NM131 | SRR20666353 | NM131 | 3.3.1 | 56 | F | Blood | 2018 | Africa | Southern Africa | Zimbabwe | Harare | Glennorah | This Study |
| QIB-NM141 | SRR20666352 | NM141 | 3.3.1 | 29 | M | Blood | 2018 | Africa | Southern Africa | Zimbabwe | Harare | Budiriro 1 | This Study |
| QIB-NM151 | SRR20666351 | NM151 | 3.3.1 | 30 | F | Stool | 2018 | Africa | Southern Africa | Zimbabwe | Harare | Glenview 1 | This Study |
| HG3-1 | ERR4870975 | HG3-1 | 4.3.1.1.EA1 | 34 | F | Stool | 2012 | Africa | Southern Africa | Zimbabwe | Harare | Glenview 3 | Mashe et al 2021 |
| HB1-2 | ERR4870976 | HB1-2 | 4.3.1.1.EA1 | 8 | M | Blood | 2014 | Africa | Southern Africa | Zimbabwe | Harare | Budiriro 1 | Mashe et al 2021 |
| HG8-3 | ERR4870977 | HG8-3 | 4.3.1.1.EA1 | 26 | F | Blood | 2016 | Africa | Southern Africa | Zimbabwe | Harare | Glenview 8 | Mashe et al 2021 |
| HK3-4 | ERR4870978 | HK3-4 | 4.3.1.1.EA1 | 7 | F | Blood | 2016 | Africa | Southern Africa | Zimbabwe | Harare | Kuwadzana 3 | Mashe et al 2021 |
| HB1-5 | ERR4870979 | HB1-5 | 4.3.1.1.EA1 | 3 | M | Blood | 2016 | Africa | Southern Africa | Zimbabwe | Harare | Budiriro 1 | Mashe et al 2021 |
| HB5-6 | ERR4870980 | HB5-6 | 4.3.1.1.EA1 | 21 | M | Blood | 2016 | Africa | Southern Africa | Zimbabwe | Harare | Budiriro 5 | Mashe et al 2021 |
| HB1-7 | ERR4870981 | HB1-7 | 4.3.1.1.EA1 | 21 | M | Blood | 2016 | Africa | Southern Africa | Zimbabwe | Harare | Budiriro 1 | Mashe et al 2021 |
| HM-8 | ERR4870982 | HM-8 | 4.3.1.1.EA1 | 5 | F | Blood | 2016 | Africa | Southern Africa | Zimbabwe | Harare | Mbare | Mashe et al 2021 |
| HM-9 | ERR4870983 | HM-9 | 3.3.1 | 20 | M | Stool | 2017 | Africa | Southern Africa | Zimbabwe | Harare | Dzivarasekwa | Mashe et al 2021 |
| HD-10 | ERR4870984 | HD-10 | 4.3.1.1.EA1 | 7 | F | Blood | 2017 | Africa | Southern Africa | Zimbabwe | Harare | Dzivarasekwa | Mashe et al 2021 |
| HM-11 | ERR4870985 | HM-11 | 3.3.1 | 10 | F | Blood | 2017 | Africa | Southern Africa | Zimbabwe | Harare | Glenview 3 | Mashe et al 2021 |
| HD-12 | ERR4870986 | HD-12 | 4.3.1.1.EA1 | 13 | F | Blood | 2017 | Africa | Southern Africa | Zimbabwe | Harare | Glenview 1 | Mashe et al 2021 |
| HG3-13 | ERR4870987 | HG3-13 | 4.3.1.1.EA1 | 16 | F | Stool | 2018 | Africa | Southern Africa | Zimbabwe | Harare | Glenview 3 | Mashe et al 2021 |
| HG1-14 | ERR4870988 | HG1-14 | 4.3.1.1.EA1 | 6 | M | Stool | 2018 | Africa | Southern Africa | Zimbabwe | Harare | Kuwadzana 3 | Mashe et al 2021 |
| HG3-15 | ERR4870989 | HG3-15 | 4.3.1.1.EA1 | 18 | M | Blood | 2018 | Africa | Southern Africa | Zimbabwe | Harare | Stoneridge | Mashe et al 2021 |
| HK3-16 | ERR4870990 | HK3-16 | 4.3.1.1.EA1 | 19 | M | Blood | 2018 | Africa | Southern Africa | Zimbabwe | Harare | Kuwadzana 3 | Mashe et al 2021 |
| HS-17 | ERR4870991 | HS-17 | 4.3.1.1.EA1 | 64 | F | Blood | 2018 | Africa | Southern Africa | Zimbabwe | Harare | Hopley | Mashe et al 2021 |
| HK3-18 | ERR4870992 | HK3-18 | 4.3.1.1.EA1 | 23 | F | Blood | 2018 | Africa | Southern Africa | Zimbabwe | Harare | Mkoba 14 | Mashe et al 2021 |
| HH-19 | ERR4870993 | HH-19 | 4.3.1.1.EA1 | 21 | M | Stool | 2018 | Africa | Southern Africa | Zimbabwe | Harare | Mkoba 15 | Mashe et al 2021 |
| GM14-20 | ERR4870994 | GM14-20 | 4.3.1.1.EA1 | 9 | F | Blood | 2018 | Africa | Southern Africa | Zimbabwe | Gweru | Mkoba 20 | Mashe et al 2021 |
| GM15-21 | ERR4870995 | GM15-21 | 4.3.1.1.EA1 | 48 | M | Blood | 2018 | Africa | Southern Africa | Zimbabwe | Gweru | Mkoba 20 | Mashe et al 2021 |
| GM20-22 | ERR4870996 | GM20-22 | 4.3.1.1.EA1 | 44 | M | Blood | 2018 | Africa | Southern Africa | Zimbabwe | Gweru | Mkoba 15 | Mashe et al 2021 |
| GM20-23 | ERR4870997 | GM20-23 | 4.3.1.1.EA1 | 17 | M | Blood | 2018 | Africa | Southern Africa | Zimbabwe | Gweru | Mufakose | Mashe et al 2021 |
| GM15-24 | ERR4870998 | GM15-24 | 4.3.1.1.EA1 | 34 | M | Blood | 2018 | Africa | Southern Africa | Zimbabwe | Gweru | Budiriro | Mashe et al 2021 |
| HM-25 | ERR4870999 | HM-25 | 4.3.1.1.EA1 | 15 | F | Blood | 2019 | Africa | Southern Africa | Zimbabwe | Harare | Budiriro | Mashe et al 2021 |
| HB-26 | ERR4871000 | HB-26 | 4.3.1.1.EA1 | 13 | F | Blood | 2019 | Africa | Southern Africa | Zimbabwe | Harare | Budiriro | Mashe et al 2021 |
| HB-27 | ERR4871001 | HB-27 | 4.3.1.1.EA1 | 13 | F | Blood | 2019 | Africa | Southern Africa | Zimbabwe | Harare | Budiriro | Mashe et al 2021 |
| HB-28 | ERR4871002 | HB-28 | 4.3.1.1.EA1 | 6 | F | Blood | 2019 | Africa | Southern Africa | Zimbabwe | Harare | Mbare | Mashe et al 2021 |
| HB-29 | ERR4871003 | HB-29 | 4.3.1.1.EA1 | 14 | M | Blood | 2019 | Africa | Southern Africa | Zimbabwe | Harare | Mbare | Mashe et al 2021 |
| unknown | SRR1963102 | SRR1963102 | 4.3.1.1.EA1 | unknown | unknown | unknown | 2014 | Africa (UK travel) | Southern Africa (UK travel) | Zimbabwe (UK travel) | unknown | unknown | Ingle et al 2019 |
| unknown | SRR1965420 | SRR1965420 | 4.3.1.1.EA1 | unknown | unknown | unknown | 2014 | Africa (UK travel) | Southern Africa (UK travel) | Zimbabwe (UK travel) | unknown | unknown | Ingle et al 2019 |
| unknown | SRR1965658 | SRR1965658 | 4.3.1.1.EA1 | unknown | unknown | unknown | 2014 | Africa (UK travel) | Southern Africa (UK travel) | Zimbabwe (UK travel) | unknown | unknown | Ingle et al 2019 |
| unknown | SRR1966355 | SRR1966355 | 4.3.1.1.EA1 | unknown | unknown | unknown | 2014 | Africa (UK travel) | Southern Africa (UK travel) | Zimbabwe (UK travel) | unknown | unknown | Ingle et al 2019 |
| unknown | SRR1967049 | SRR1967049 | 4.3.1.1.EA1 | unknown | unknown | unknown | 2014 | Africa (UK travel) | Southern Africa (UK travel) | Zimbabwe (UK travel) | unknown | unknown | Ingle et al 2019 |
| unknown | SRR1969286 | SRR1969286 | 3.3.1 | unknown | unknown | unknown | 2014 | Africa (UK travel) | Southern Africa (UK travel) | Zimbabwe (UK travel) | unknown | unknown | Ingle et al 2019 |
| unknown | SRR1969590 | SRR1969590 | 3.3.1 | unknown | unknown | unknown | 2014 | Africa (UK travel) | Southern Africa (UK travel) | Zimbabwe (UK travel) | unknown | unknown | Ingle et al 2019 |
| unknown | SRR3322588 | SRR3322588 | 4.3.1.1.EA1 | unknown | unknown | unknown | 2015 | Africa (UK travel) | Southern Africa (UK travel) | Zimbabwe (UK travel) | unknown | unknown | Ingle et al 2019 |
| unknown | SRR4063811 | SRR4063811 | 4.3.1.1.EA1 | unknown | unknown | unknown | 2016 | Africa (UK travel) | Southern Africa (UK travel) | Zimbabwe (UK travel) | unknown | unknown | Ingle et al 2019 |
| unknown | SRR7165581 | SRR7165581 | 4.3.1.1.EA1 | unknown | unknown | unknown | 2016 | Africa (UK travel) | Southern Africa (UK travel) | Zimbabwe (UK travel) | unknown | unknown | Ingle et al 2019 |
| H05118260 | ERR1017039 | N/A | 4.3.1.1 | unknown | unknown | Blood | 2005 | Unknown | Unknown | Unknown | unknown | unknown | Wong et al, 2016 |
| H05196407 | ERR1017040 | N/A | 4.3.1 | unknown | unknown | Blood | 2005 | Unknown | Unknown | Unknown | unknown | unknown | Wong et al, 2016 |
| H05272442 | ERR1017041 | N/A | 3.1.1 | unknown | unknown | Blood | 2005 | Unknown | Unknown | Unknown | unknown | unknown | Wong et al, 2016 |

| | | | | | | | | | | | | | |
|---|---|---|---|---|---|---|---|---|---|---|---|---|---|
| H05406403 | ERR1017042 | N/A | 4.3.1.1 | unknown | unknown | Blood | 2005 | Unknown | Unknown | Unknown | unknown | unknown | Wong et al, 2016 |
| H06016481 | ERR1017045 | N/A | 4.3.1.1 | unknown | unknown | Blood | 2005 | Unknown | Unknown | Unknown | unknown | unknown | Wong et al, 2016 |
| H06156550 | ERR1017048 | N/A | 3.0.1 | unknown | unknown | Blood | 2006 | Unknown | Unknown | Unknown | unknown | unknown | Wong et al, 2016 |
| H06384614 | ERR1017052 | N/A | 4.3.1 | unknown | unknown | Blood | 2006 | Asia | South Asia | Pakistan | unknown | unknown | Wong et al, 2016 |
| H06394364 | ERR1017054 | N/A | 4.3.1.1 | unknown | unknown | Blood | 2006 | Asia | South Asia | Bangladesh | unknown | unknown | Wong et al, 2016 |
| H06414501 | ERR1017055 | N/A | 3.3.2 | unknown | unknown | Blood | 2006 | Asia | South Asia | Bangladesh | unknown | unknown | Wong et al, 2016 |
| H06434426 | ERR1017060 | N/A | 2.0.2 | unknown | unknown | Blood | 2006 | Unknown | Unknown | Unknown | unknown | unknown | Wong et al, 2016 |
| H07014193 | ERR1017063 | N/A | 2.2 | unknown | unknown | Blood | 2006 | Asia | South Asia | Pakistan | unknown | unknown | Wong et al, 2016 |
| H07044209 | ERR1017064 | N/A | 4.3.1.2 | unknown | unknown | Blood | 2007 | Asia | South Asia | India | unknown | unknown | Wong et al, 2016 |
| H07288307 | ERR1017067 | N/A | 4.3.1.2 | unknown | unknown | Blood | 2007 | Asia | South Asia | Nepal | unknown | unknown | Wong et al, 2016 |
| H07288308 | ERR1017068 | N/A | 4.3.1 | unknown | unknown | Blood | 2007 | Asia | South Asia | India | unknown | unknown | Wong et al, 2016 |
| H07324312 | ERR1017069 | N/A | 4.3.1.1 | unknown | unknown | Blood | 2007 | Asia | South Asia | Pakistan | unknown | unknown | Wong et al, 2016 |
| H07336300 | ERR1017070 | N/A | 4.3.1.2 | unknown | unknown | Blood | 2007 | Asia | South Asia | Pakistan | unknown | unknown | Wong et al, 2016 |
| H07336301 | ERR1017071 | N/A | 4.3.1.1 | unknown | unknown | Blood | 2007 | Asia | South Asia | Bangladesh | unknown | unknown | Wong et al, 2016 |
| H07364324 | ERR1017072 | N/A | 4.3.1.1 | unknown | unknown | Blood | 2007 | Asia | South Asia | Bangladesh | unknown | unknown | Wong et al, 2016 |
| H07384494 | ERR1017074 | N/A | 4.3.1.2 | unknown | unknown | Blood | 2007 | Asia | South Asia | India | unknown | unknown | Wong et al, 2016 |
| H07394454 | ERR1017075 | N/A | 4.3.1 | unknown | unknown | Blood | 2007 | Asia | South Asia | India | unknown | unknown | Wong et al, 2016 |
| H07402281 | ERR1017077 | N/A | 4.3.1.3 | unknown | unknown | Blood | 2007 | Asia | South Asia | Bangladesh | unknown | unknown | Wong et al, 2016 |
| H07404418 | ERR1017078 | N/A | 3.1.1 | unknown | unknown | Blood | 2007 | Africa | West Africa | Ghana | unknown | unknown | Wong et al, 2016 |
| H08038152 | ERR1017081 | N/A | 2.2.2 | unknown | unknown | Blood | 2008 | Asia | South Asia | India | unknown | unknown | Wong et al, 2016 |
| H08082282 | ERR1017083 | N/A | 4.3.1.2 | unknown | unknown | Blood | 2008 | Asia | South Asia | India | unknown | unknown | Wong et al, 2016 |
| H08294356 | ERR1017087 | N/A | 4.3.1.1 | unknown | unknown | Blood | 2008 | Asia | South Asia | Bangladesh | unknown | unknown | Wong et al, 2016 |
| H08302395 | ERR1017088 | N/A | 3.2.2 | unknown | unknown | Blood | 2008 | Asia | South Asia | Bangladesh | unknown | unknown | Wong et al, 2016 |
| H08302396 | ERR1017089 | N/A | 4.3.1.1 | unknown | unknown | Blood | 2008 | Asia | South Asia | Bangladesh | unknown | unknown | Wong et al, 2016 |
| H08328499 | ERR1017090 | N/A | 4.3.1 | unknown | unknown | Blood | 2008 | Asia | W Asia/S Asia | Kuwait/India | unknown | unknown | Wong et al, 2016 |
| H08384665 | ERR1017091 | N/A | 3.3 | unknown | unknown | Blood | 2008 | Asia | South Asia | Pakistan | unknown | unknown | Wong et al, 2016 |
| H08398180 | ERR1017092 | N/A | 4.3.1 | unknown | unknown | Blood | 2008 | Asia | W Asia/S Asia | Kuwait/India | unknown | unknown | Wong et al, 2016 |
| H08428259 | ERR1017094 | N/A | 4.3.1.2 | unknown | unknown | Blood | 2008 | Asia | South Asia | India | unknown | unknown | Wong et al, 2016 |
| H08434321 | ERR1017095 | N/A | 3.3 | unknown | unknown | Blood | 2008 | Asia | South Asia | India/Pakistan | unknown | unknown | Wong et al, 2016 |
| H09044141 | ERR1017096 | N/A | 3.1.1 | unknown | unknown | Blood | 2009 | Africa | West Africa | Nigeria | unknown | unknown | Wong et al, 2016 |
| H0905277 | ERR1017097 | N/A | 4.3.1.1 | unknown | unknown | Blood | 2009 | Asia | South Asia | Pakistan | unknown | unknown | Wong et al, 2016 |
| H09132110 | ERR1017098 | N/A | 3.3.1 | unknown | unknown | Blood | 2009 | Asia | South Asia | Pakistan | unknown | unknown | Wong et al, 2016 |
| IB5335 | ERR108648 | N/A | 4.3.1.1.EA1 | unknown | unknown | Blood | 2009 | Africa | East Africa | Tanzania | unknown | unknown | Wong et al, 2015 |
| IB5336 | ERR108649 | N/A | 2.5 | unknown | unknown | Blood | 2009 | Africa | East Africa | Tanzania | unknown | unknown | Wong et al, 2015 |
| IB5339 | ERR108652 | N/A | 3.3.1 | unknown | unknown | Blood | 2009 | Africa | East Africa | Tanzania | unknown | unknown | Wong et al, 2015 |
| IB5340 | ERR108653 | N/A | 3.3.1 | unknown | unknown | Blood | 2009 | Africa | East Africa | Tanzania | unknown | unknown | Wong et al, 2015 |
| IB5341 | ERR108654 | N/A | 4.3.1.2 | unknown | unknown | Blood | 2009 | Africa | East Africa | Tanzania | unknown | unknown | Wong et al, 2015 |
| IB5342 | ERR108655 | N/A | 3.3.1 | unknown | unknown | Blood | 2009 | Africa | East Africa | Tanzania | unknown | unknown | Wong et al, 2015 |
| IB5343 | ERR108656 | N/A | 4.3.1.1.EA1 | unknown | unknown | Blood | 2009 | Africa | East Africa | Tanzania | unknown | unknown | Wong et al, 2015 |
| IB5345 | ERR108658 | N/A | 4.3.1.1.EA1 | unknown | unknown | Blood | 2009 | Africa | East Africa | Tanzania | unknown | unknown | Wong et al, 2015 |
| IB5347 | ERR108659 | N/A | 4.3.1.1.EA1 | unknown | unknown | Blood | 2009 | Africa | East Africa | Tanzania | unknown | unknown | Wong et al, 2015 |
| IB5348 | ERR108660 | N/A | 4.3.1.1.EA1 | unknown | unknown | Blood | 2009 | Africa | East Africa | Tanzania | unknown | unknown | Wong et al, 2015 |
| IB5351 | ERR108663 | N/A | 4.3.1.2 | unknown | unknown | Blood | 2010 | Africa | East Africa | Tanzania | unknown | unknown | Wong et al, 2015 |
| IB5352 | ERR108664 | N/A | 4.3.1.1.EA1 | unknown | unknown | Blood | 2010 | Africa | East Africa | Tanzania | unknown | unknown | Wong et al, 2015 |
| IB5353 | ERR108665 | N/A | 4.3.1.2 | unknown | unknown | Blood | 2010 | Africa | East Africa | Tanzania | unknown | unknown | Wong et al, 2015 |
| IB5354 | ERR108666 | N/A | 4.3.1.2 | unknown | unknown | Blood | 2010 | Africa | East Africa | Tanzania | unknown | unknown | Wong et al, 2015 |
| IB5355 | ERR108667 | N/A | 4.3.1.2 | unknown | unknown | Blood | 2010 | Africa | East Africa | Tanzania | unknown | unknown | Wong et al, 2015 |
| IB5356 | ERR108668 | N/A | 4.3.1.2 | unknown | unknown | Blood | 2010 | Africa | East Africa | Tanzania | unknown | unknown | Wong et al, 2015 |
| IB5357 | ERR108669 | N/A | 4.3.1.2 | unknown | unknown | Blood | 2010 | Africa | East Africa | Tanzania | unknown | unknown | Wong et al, 2015 |
| IB5358 | ERR108670 | N/A | 4.3.1.2 | unknown | unknown | Blood | 2010 | Africa | East Africa | Tanzania | unknown | unknown | Wong et al, 2015 |

| | | | | | | | | | | | | | |
|---|---|---|---|---|---|---|---|---|---|---|---|---|---|
| IB5359 | ERR108671 | N/A | 3.3.1 | unknown | unknown | Blood | 2010 | Africa | East Africa | Tanzania | unknown | unknown | Wong et al, 2015 |
| IB5360 | ERR108672 | N/A | 3.3.1 | unknown | unknown | Blood | 2009 | Africa | East Africa | Tanzania | unknown | unknown | Wong et al, 2015 |
| IB5366 | ERR108676 | N/A | 4.3.1.1.EA1 | unknown | unknown | Blood | 2009 | Africa | East Africa | Tanzania | unknown | unknown | Wong et al, 2015 |
| IB5367 | ERR108677 | N/A | 3.3.1 | unknown | unknown | Blood | 2009 | Africa | East Africa | Tanzania | unknown | unknown | Wong et al, 2015 |
| IB5368 | ERR108678 | N/A | 2.5.1 | unknown | unknown | Blood | 2009 | Africa | East Africa | Tanzania | unknown | unknown | Wong et al, 2015 |
| IB5369 | ERR108679 | N/A | 4.3.1.1.EA1 | unknown | unknown | Blood | 2009 | Africa | East Africa | Tanzania | unknown | unknown | Wong et al, 2015 |
| IB5370 | ERR108680 | N/A | 3.3.1 | unknown | unknown | Blood | 2010 | Africa | East Africa | Tanzania | unknown | unknown | Wong et al, 2015 |
| IB5371 | ERR108681 | N/A | 3.3.1 | unknown | unknown | Blood | 2010 | Africa | East Africa | Tanzania | unknown | unknown | Wong et al, 2015 |
| IB5374 | ERR108684 | N/A | 3.3.1 | unknown | unknown | Blood | 2010 | Africa | East Africa | Tanzania | unknown | unknown | Wong et al, 2015 |
| IB5375 | ERR108685 | N/A | 4.3.1.2 | unknown | unknown | Blood | 2010 | Africa | East Africa | Tanzania | unknown | unknown | Wong et al, 2015 |
| IB5376 | ERR108686 | N/A | 4.3.1.2 | unknown | unknown | Blood | 2010 | Africa | East Africa | Tanzania | unknown | unknown | Wong et al, 2015 |
| IB5378 | ERR108687 | N/A | 4.3.1.2 | unknown | unknown | Blood | 2010 | Africa | East Africa | Tanzania | unknown | unknown | Wong et al, 2015 |
| IB5379 | ERR108688 | N/A | 4.3.1.2 | unknown | unknown | Blood | 2010 | Africa | East Africa | Tanzania | unknown | unknown | Wong et al, 2015 |
| IB5380 | ERR108689 | N/A | 4.3.1.2 | unknown | unknown | Blood | 2010 | Africa | East Africa | Tanzania | unknown | unknown | Wong et al, 2015 |
| IB5381 | ERR108690 | N/A | 3.3.1 | unknown | unknown | Blood | 2010 | Africa | East Africa | Tanzania | unknown | unknown | Wong et al, 2015 |
| IB5382 | ERR108691 | N/A | 4.3.1.2 | unknown | unknown | Blood | 2010 | Africa | East Africa | Tanzania | unknown | unknown | Wong et al, 2015 |
| IB5383 | ERR108692 | N/A | 3.3.1 | unknown | unknown | Blood | 2009 | Africa | East Africa | Tanzania | unknown | unknown | Wong et al, 2015 |
| IB5384 | ERR108693 | N/A | 4.3.1.1.EA1 | unknown | unknown | Blood | 2009 | Africa | East Africa | Tanzania | unknown | unknown | Wong et al, 2015 |
| IB5385 | ERR108694 | N/A | 4.3.1.1.EA1 | unknown | unknown | Blood | 2010 | Africa | East Africa | Tanzania | unknown | unknown | Wong et al, 2015 |
| MDUST73 | ERR204244 | N/A | 4.2.1 | unknown | unknown | Unknown | 2008 | Australia & Oceania | Oceania | Fiji | unknown | unknown | Wong et al, 2015 |
| MDUST74 | ERR204245 | N/A | 4.2.1 | unknown | unknown | Unknown | 2008 | Australia & Oceania | Oceania | Fiji | unknown | unknown | Wong et al, 2015 |
| MDUST75 | ERR204246 | N/A | 4.2.2 | unknown | unknown | Unknown | 2008 | Australia & Oceania | Oceania | Fiji | unknown | unknown | Wong et al, 2015 |
| MDUST76 | ERR204247 | N/A | 4.2.2 | unknown | unknown | Unknown | 2008 | Australia & Oceania | Oceania | Fiji | unknown | unknown | Wong et al, 2015 |
| MDUST77 | ERR204248 | N/A | 4.2.1 | unknown | unknown | Unknown | 2008 | Australia & Oceania | Oceania | Fiji | unknown | unknown | Wong et al, 2015 |
| MDUST78 | ERR204249 | N/A | 4.2.1 | unknown | unknown | Unknown | 2008 | Australia & Oceania | Oceania | Fiji | unknown | unknown | Wong et al, 2015 |
| MDUST79 | ERR204250 | N/A | 4.2.1 | unknown | unknown | Unknown | 2008 | Australia & Oceania | Oceania | Fiji | unknown | unknown | Wong et al, 2015 |
| MDUST80 | ERR204251 | N/A | 4.2.2 | unknown | unknown | Unknown | 2008 | Australia & Oceania | Oceania | Fiji | unknown | unknown | Wong et al, 2015 |
| MDUST81 | ERR204252 | N/A | 4.2.1 | unknown | unknown | Unknown | 2008 | Australia & Oceania | Oceania | Fiji | unknown | unknown | Wong et al, 2015 |
| MDUST82 | ERR204253 | N/A | 4.2.1 | unknown | unknown | Unknown | 2008 | Australia & Oceania | Oceania | Fiji | unknown | unknown | Wong et al, 2015 |
| MDUST83 | ERR204254 | N/A | 4.2.2 | unknown | unknown | Unknown | 2008 | Australia & Oceania | Oceania | Fiji | unknown | unknown | Wong et al, 2015 |
| MDUST84 | ERR204255 | N/A | 4.2.1 | unknown | unknown | Unknown | 2008 | Australia & Oceania | Oceania | Fiji | unknown | unknown | Wong et al, 2015 |
| MDUST85 | ERR204256 | N/A | 4.2.1 | unknown | unknown | Unknown | 2008 | Australia & Oceania | Oceania | Fiji | unknown | unknown | Wong et al, 2015 |
| MDUST86 | ERR204257 | N/A | 4.2.1 | unknown | unknown | Unknown | 2008 | Australia & Oceania | Oceania | Fiji | unknown | unknown | Wong et al, 2015 |
| MDUST87 | ERR204258 | N/A | 4.2.1 | unknown | unknown | Unknown | 2008 | Australia & Oceania | Oceania | Fiji | unknown | unknown | Wong et al, 2015 |
| MDUST88 | ERR204259 | N/A | 4.2.1 | unknown | unknown | Unknown | 2008 | Australia & Oceania | Oceania | Fiji | unknown | unknown | Wong et al, 2015 |
| MDUST89 | ERR204260 | N/A | 4.2.2 | unknown | unknown | Unknown | 2008 | Australia & Oceania | Oceania | Fiji | unknown | unknown | Wong et al, 2015 |
| MDUST90 | ERR204261 | N/A | 4.2.2 | unknown | unknown | Unknown | 2008 | Australia & Oceania | Oceania | Fiji | unknown | unknown | Wong et al, 2015 |
| MDUST91 | ERR204262 | N/A | 4.2.2 | unknown | unknown | Unknown | 2008 | Australia & Oceania | Oceania | Fiji | unknown | unknown | Wong et al, 2015 |
| MDUST92 | ERR204263 | N/A | 4.2.2 | unknown | unknown | Unknown | 2008 | Australia & Oceania | Oceania | Fiji | unknown | unknown | Wong et al, 2015 |
| MDUST93 | ERR204264 | N/A | 4.2.2 | unknown | unknown | Unknown | 2008 | Australia & Oceania | Oceania | Fiji | unknown | unknown | Wong et al, 2015 |
| MDUST94 | ERR204265 | N/A | 4.2.2 | unknown | unknown | Unknown | 2008 | Australia & Oceania | Oceania | Fiji | unknown | unknown | Wong et al, 2015 |
| MDUST95 | ERR204266 | N/A | 4.2.2 | unknown | unknown | Unknown | 2008 | Australia & Oceania | Oceania | Fiji | unknown | unknown | Wong et al, 2015 |
| MDUST96 | ERR204267 | N/A | 4.2.2 | unknown | unknown | Unknown | 2008 | Australia & Oceania | Oceania | Fiji | unknown | unknown | Wong et al, 2015 |
| MDUST98 | ERR204269 | N/A | 4.2.2 | unknown | unknown | Unknown | 2008 | Australia & Oceania | Oceania | Fiji | unknown | unknown | Wong et al, 2015 |
| MDUST99 | ERR204270 | N/A | 4.2.1 | unknown | unknown | Unknown | 2008 | Australia & Oceania | Oceania | Fiji | unknown | unknown | Wong et al, 2015 |
| MDUST100 | ERR204271 | N/A | 4.2.1 | unknown | unknown | Unknown | 2008 | Australia & Oceania | Oceania | Fiji | unknown | unknown | Wong et al, 2015 |
| MDUST101 | ERR204272 | N/A | 4.2.2 | unknown | unknown | Unknown | 2008 | Australia & Oceania | Oceania | Fiji | unknown | unknown | Wong et al, 2015 |
| MDUST102 | ERR204273 | N/A | 4.2.2 | unknown | unknown | Unknown | 2008 | Australia & Oceania | Oceania | Fiji | unknown | unknown | Wong et al, 2015 |
| MDUST103 | ERR204274 | N/A | 4.2.2 | unknown | unknown | Unknown | 2008 | Australia & Oceania | Oceania | Fiji | unknown | unknown | Wong et al, 2015 |
| MDUST104 | ERR204275 | N/A | 4.2.2 | unknown | unknown | Unknown | 2008 | Australia & Oceania | Oceania | Fiji | unknown | unknown | Wong et al, 2015 |

| | | | | | | | | | | | | | |
|---|---|---|---|---|---|---|---|---|---|---|---|---|---|
| MDUST105 | ERR204276 | N/A | 4.2.2 | unknown | unknown | Unknown | 2008 | Australia & Oceania | Oceania | Fiji | unknown | unknown | Wong et al, 2015 |
| Ke378 | ERR212564 | N/A | 4.3.1.1.EA1 | unknown | unknown | Blood | 2008 | Africa | East Africa | Kenya | unknown | unknown | Wong et al, 2015 |
| Ke451 | ERR212636 | N/A | 4.3.1.1.EA1 | unknown | unknown | Blood | 2008 | Africa | East Africa | Kenya | unknown | unknown | Wong et al, 2015 |
| Ke452 | ERR212637 | N/A | 4.3.1.2.EA2 | unknown | unknown | Blood | 2008 | Africa | East Africa | Kenya | unknown | unknown | Wong et al, 2015 |
| Ke453 | ERR212638 | N/A | 4.3.1.1.EA1 | unknown | unknown | Blood | 2008 | Africa | East Africa | Kenya | unknown | unknown | Wong et al, 2015 |
| Ke454 | ERR212639 | N/A | 4.3.1.1.EA1 | unknown | unknown | Blood | 2008 | Africa | East Africa | Kenya | unknown | unknown | Wong et al, 2015 |
| Ke455 | ERR212640 | N/A | 4.3.1.1.EA1 | unknown | unknown | Blood | 2008 | Africa | East Africa | Kenya | unknown | unknown | Wong et al, 2015 |
| Ke458 | ERR212643 | N/A | 4.3.1.1.EA1 | unknown | unknown | Blood | 2008 | Africa | East Africa | Kenya | unknown | unknown | Wong et al, 2015 |
| Ke461 | ERR212646 | N/A | 4.3.1.1.EA1 | unknown | unknown | Blood | 2008 | Africa | East Africa | Kenya | unknown | unknown | Wong et al, 2015 |
| Ke462 | ERR212647 | N/A | 2.5 | unknown | unknown | Blood | 2008 | Africa | East Africa | Kenya | unknown | unknown | Wong et al, 2015 |
| Ke463 | ERR212648 | N/A | 4.3.1.1.EA1 | unknown | unknown | Blood | 2008 | Africa | East Africa | Kenya | unknown | unknown | Wong et al, 2015 |
| Ke466 | ERR212651 | N/A | 4.3.1.1.EA1 | unknown | unknown | Blood | 2008 | Africa | East Africa | Kenya | unknown | unknown | Wong et al, 2015 |
| Ke467 | ERR212652 | N/A | 4.3.1.1.EA1 | unknown | unknown | Blood | 2008 | Africa | East Africa | Kenya | unknown | unknown | Wong et al, 2015 |
| Ke468 | ERR212653 | N/A | 4.3.1.1.EA1 | unknown | unknown | Blood | 2008 | Africa | East Africa | Kenya | unknown | unknown | Wong et al, 2015 |
| Ke469 | ERR212654 | N/A | 4.3.1.1.EA1 | unknown | unknown | Blood | 2008 | Africa | East Africa | Kenya | unknown | unknown | Wong et al, 2015 |
| Ke470 | ERR212655 | N/A | 4.3.1.1.EA1 | unknown | unknown | Blood | 2008 | Africa | East Africa | Kenya | unknown | unknown | Wong et al, 2015 |
| Ke472 | ERR212657 | N/A | 4.3.1.1.EA1 | unknown | unknown | Blood | 2008 | Africa | East Africa | Kenya | unknown | unknown | Wong et al, 2015 |
| Ke474 | ERR212659 | N/A | 4.3.1.1.EA1 | unknown | unknown | Blood | 2008 | Africa | East Africa | Kenya | unknown | unknown | Wong et al, 2015 |
| Ke475 | ERR212660 | N/A | 4.3.1.1.EA1 | unknown | unknown | Blood | 2008 | Africa | East Africa | Kenya | unknown | unknown | Wong et al, 2015 |
| Ke476 | ERR212661 | N/A | 4.3.1.1.EA1 | unknown | unknown | Blood | 2008 | Africa | East Africa | Kenya | unknown | unknown | Wong et al, 2015 |
| Ke477 | ERR212662 | N/A | 4.3.1.1.EA1 | unknown | unknown | Blood | 2008 | Africa | East Africa | Kenya | unknown | unknown | Wong et al, 2015 |
| Ke479 | ERR212664 | N/A | 4.3.1.1.EA1 | unknown | unknown | Blood | 2008 | Africa | East Africa | Kenya | unknown | unknown | Wong et al, 2015 |
| Ke480 | ERR212665 | N/A | 4.3.1.1.EA1 | unknown | unknown | Blood | 2008 | Africa | East Africa | Kenya | unknown | unknown | Wong et al, 2015 |
| Ke481 | ERR212666 | N/A | 4.3.1.1.EA1 | unknown | unknown | Blood | 2008 | Africa | East Africa | Kenya | unknown | unknown | Wong et al, 2015 |
| Ke482 | ERR212667 | N/A | 4.3.1.1.EA1 | unknown | unknown | Blood | 2008 | Africa | East Africa | Kenya | unknown | unknown | Wong et al, 2015 |
| Ke492 | ERR212677 | N/A | 4.3.1.2.EA2 | unknown | unknown | Blood | 2001 | Africa | East Africa | Kenya | unknown | unknown | Wong et al, 2015 |
| MDUST1 | ERR213219 | N/A | 4.2.2 | unknown | unknown | Unknown | 2011 | Australia & Oceania | Oceania | Fiji | unknown | unknown | Wong et al, 2015 |
| MDUST2 | ERR213220 | N/A | 4.2.2 | unknown | unknown | Unknown | 2011 | Australia & Oceania | Oceania | Fiji | unknown | unknown | Wong et al, 2015 |
| MDUST3 | ERR213221 | N/A | 4.2.2 | unknown | unknown | Unknown | 2011 | Australia & Oceania | Oceania | Fiji | unknown | unknown | Wong et al, 2015 |
| MDUST4 | ERR213222 | N/A | 4.2.2 | unknown | unknown | Unknown | 2011 | Australia & Oceania | Oceania | Fiji | unknown | unknown | Wong et al, 2015 |
| MDUST5 | ERR213223 | N/A | 4.2.2 | unknown | unknown | Unknown | 2011 | Australia & Oceania | Oceania | Fiji | unknown | unknown | Wong et al, 2015 |
| MDUST6 | ERR213224 | N/A | 4.2.2 | unknown | unknown | Unknown | 2011 | Australia & Oceania | Oceania | Fiji | unknown | unknown | Wong et al, 2015 |
| MDUST7 | ERR213225 | N/A | 4.2.2 | unknown | unknown | Unknown | 2011 | Australia & Oceania | Oceania | Fiji | unknown | unknown | Wong et al, 2015 |
| MDUST8 | ERR213226 | N/A | 4.2.2 | unknown | unknown | Unknown | 2011 | Australia & Oceania | Oceania | Fiji | unknown | unknown | Wong et al, 2015 |
| MDUST9 | ERR213227 | N/A | 4.2.2 | unknown | unknown | Unknown | 2011 | Australia & Oceania | Oceania | Fiji | unknown | unknown | Wong et al, 2015 |
| MDUST10 | ERR213228 | N/A | 4.2.2 | unknown | unknown | Unknown | 2011 | Australia & Oceania | Oceania | Fiji | unknown | unknown | Wong et al, 2015 |
| MDUST11 | ERR213229 | N/A | 4.2.2 | unknown | unknown | Unknown | 2011 | Australia & Oceania | Oceania | Fiji | unknown | unknown | Wong et al, 2015 |
| MDUST12 | ERR213230 | N/A | 4.2.2 | unknown | unknown | Unknown | 2011 | Australia & Oceania | Oceania | Fiji | unknown | unknown | Wong et al, 2015 |
| MDUST13 | ERR213231 | N/A | 4.2.2 | unknown | unknown | Unknown | 2011 | Australia & Oceania | Oceania | Fiji | unknown | unknown | Wong et al, 2015 |
| MDUST14 | ERR213232 | N/A | 4.2.2 | unknown | unknown | Unknown | 2011 | Australia & Oceania | Oceania | Fiji | unknown | unknown | Wong et al, 2015 |
| MDUST15 | ERR213233 | N/A | 4.2.1 | unknown | unknown | Unknown | 2011 | Australia & Oceania | Oceania | Fiji | unknown | unknown | Wong et al, 2015 |
| MDUST16 | ERR213234 | N/A | 4.2.2 | unknown | unknown | Unknown | 2011 | Australia & Oceania | Oceania | Fiji | unknown | unknown | Wong et al, 2015 |
| MDUST17 | ERR213235 | N/A | 4.2.2 | unknown | unknown | Unknown | 2011 | Australia & Oceania | Oceania | Fiji | unknown | unknown | Wong et al, 2015 |
| MDUST18 | ERR213236 | N/A | 4.2.2 | unknown | unknown | Unknown | 2011 | Australia & Oceania | Oceania | Fiji | unknown | unknown | Wong et al, 2015 |
| MDUST19 | ERR213237 | N/A | 4.2.2 | unknown | unknown | Unknown | 2011 | Australia & Oceania | Oceania | Fiji | unknown | unknown | Wong et al, 2015 |
| MDUST20 | ERR213238 | N/A | 4.2.2 | unknown | unknown | Unknown | 2011 | Australia & Oceania | Oceania | Fiji | unknown | unknown | Wong et al, 2015 |
| MDUST21 | ERR213239 | N/A | 4.2.2 | unknown | unknown | Unknown | 2011 | Australia & Oceania | Oceania | Fiji | unknown | unknown | Wong et al, 2015 |
| MDUST22 | ERR213240 | N/A | 4.2.2 | unknown | unknown | Unknown | 2011 | Australia & Oceania | Oceania | Fiji | unknown | unknown | Wong et al, 2015 |
| MDUST23 | ERR213241 | N/A | 4.2.2 | unknown | unknown | Unknown | 2011 | Australia & Oceania | Oceania | Fiji | unknown | unknown | Wong et al, 2015 |
| MDUST24 | ERR213242 | N/A | 4.2.2 | unknown | unknown | Unknown | 2011 | Australia & Oceania | Oceania | Fiji | unknown | unknown | Wong et al, 2015 |

| | | | | | | | | | | | | | |
|---|---|---|---|---|---|---|---|---|---|---|---|---|---|
| MDUST25 | ERR213243 | N/A | 4.2.2 | unknown | unknown | Unknown | 2011 | Australia & Oceania | Oceania | Fiji | unknown | unknown | Wong et al, 2015 |
| MDUST26 | ERR213244 | N/A | 4.2.2 | unknown | unknown | Unknown | 2011 | Australia & Oceania | Oceania | Fiji | unknown | unknown | Wong et al, 2015 |
| MDUST27 | ERR213245 | N/A | 4.2.2 | unknown | unknown | Unknown | 2011 | Australia & Oceania | Oceania | Fiji | unknown | unknown | Wong et al, 2015 |
| MDUST28 | ERR213246 | N/A | 4.2.2 | unknown | unknown | Unknown | 2011 | Australia & Oceania | Oceania | Fiji | unknown | unknown | Wong et al, 2015 |
| MDUST29 | ERR213247 | N/A | 4.2.2 | unknown | unknown | Unknown | 2011 | Australia & Oceania | Oceania | Fiji | unknown | unknown | Wong et al, 2015 |
| MDUST30 | ERR213248 | N/A | 4.2.2 | unknown | unknown | Stool | 2009 | Australia & Oceania | Oceania | Fiji | unknown | unknown | Wong et al, 2015 |
| MDUST31 | ERR213249 | N/A | 4.2.1 | unknown | unknown | Blood | 2000 | Australia & Oceania | Oceania | Fiji | unknown | unknown | Wong et al, 2015 |
| MDUST32 | ERR213250 | N/A | 4.2.1 | unknown | unknown | Blood | 2000 | Australia & Oceania | Oceania | Fiji | unknown | unknown | Wong et al, 2015 |
| MDUST33 | ERR213251 | N/A | 4.2.1 | unknown | unknown | Stool | 1997 | Australia & Oceania | Oceania | Fiji | unknown | unknown | Wong et al, 2015 |
| MDUST34 | ERR213252 | N/A | 4.2.1 | unknown | unknown | Pus | 1997 | Australia & Oceania | Oceania | Fiji | unknown | unknown | Wong et al, 2015 |
| MDUST35 | ERR213253 | N/A | 4.2 | unknown | unknown | Blood | 1996 | Australia & Oceania | Oceania | Fiji | unknown | unknown | Wong et al, 2015 |
| MDUST36 | ERR213254 | N/A | 4.2.1 | unknown | unknown | Stool | 1996 | Australia & Oceania | Oceania | Fiji | unknown | unknown | Wong et al, 2015 |
| MDUST37 | ERR213255 | N/A | 4.2.1 | unknown | unknown | Stool | 1996 | Australia & Oceania | Oceania | Fiji | unknown | unknown | Wong et al, 2015 |
| MDUST38 | ERR213256 | N/A | 4.2.1 | unknown | unknown | Stool | 1996 | Australia & Oceania | Oceania | Fiji | unknown | unknown | Wong et al, 2015 |
| MDUST39 | ERR213257 | N/A | 4.2.1 | unknown | unknown | Stool | 1996 | Australia & Oceania | Oceania | Fiji | unknown | unknown | Wong et al, 2015 |
| MDUST40 | ERR213258 | N/A | 4.2.1 | unknown | unknown | Stool | 1996 | Australia & Oceania | Oceania | Fiji | unknown | unknown | Wong et al, 2015 |
| MDUST41 | ERR213259 | N/A | 4.2.1 | unknown | unknown | Blood | 1996 | Australia & Oceania | Oceania | Fiji | unknown | unknown | Wong et al, 2015 |
| MDUST42 | ERR213260 | N/A | 4.2.3 | unknown | unknown | Blood | 1996 | Australia & Oceania | Oceania | Fiji | unknown | unknown | Wong et al, 2015 |
| MDUST43 | ERR213261 | N/A | 4.2.3 | unknown | unknown | Blood | 1996 | Australia & Oceania | Oceania | Fiji | unknown | unknown | Wong et al, 2015 |
| MDUST44 | ERR213262 | N/A | 3.5 | unknown | unknown | Blood | 1996 | Australia & Oceania | Oceania | Fiji | unknown | unknown | Wong et al, 2015 |
| MDUST45 | ERR213263 | N/A | 0.1 | unknown | unknown | Blood | 1996 | Australia & Oceania | Oceania | Fiji | unknown | unknown | Wong et al, 2015 |
| MDUST46 | ERR213264 | N/A | 2 | unknown | unknown | Blood | 1996 | Australia & Oceania | Oceania | Fiji | unknown | unknown | Wong et al, 2015 |
| MDUST47 | ERR213265 | N/A | 4.2.3 | unknown | unknown | Blood | 1996 | Australia & Oceania | Oceania | Fiji | unknown | unknown | Wong et al, 2015 |
| MDUST48 | ERR213266 | N/A | 2.3.5 | unknown | unknown | Stool | 1996 | Australia & Oceania | Oceania | Fiji | unknown | unknown | Wong et al, 2015 |
| MDUST49 | ERR213267 | N/A | 4.2 | unknown | unknown | Stool | 1996 | Australia & Oceania | Oceania | Fiji | unknown | unknown | Wong et al, 2015 |
| MDUST50 | ERR213268 | N/A | 4.2 | unknown | unknown | Stool | 1996 | Australia & Oceania | Oceania | Fiji | unknown | unknown | Wong et al, 2015 |
| MDUST51 | ERR213269 | N/A | 4.2.3 | unknown | unknown | Stool | 1995 | Australia & Oceania | Oceania | Fiji | unknown | unknown | Wong et al, 2015 |
| MDUST52 | ERR213270 | N/A | 4.2 | unknown | unknown | Stool | 1995 | Australia & Oceania | Oceania | Fiji | unknown | unknown | Wong et al, 2015 |
| MDUST53 | ERR213271 | N/A | 4.2.3 | unknown | unknown | Stool | 1995 | Australia & Oceania | Oceania | Fiji | unknown | unknown | Wong et al, 2015 |
| MDUST54 | ERR213272 | N/A | 4.2.3 | unknown | unknown | Stool | 1995 | Australia & Oceania | Oceania | Fiji | unknown | unknown | Wong et al, 2015 |
| MDUST55 | ERR213273 | N/A | 4.3.1.1 | unknown | unknown | Blood | 1995 | Australia & Oceania | Oceania | Fiji | unknown | unknown | Wong et al, 2015 |
| MDUST56 | ERR213274 | N/A | 2.3.5 | unknown | unknown | Stool | 1994 | Australia & Oceania | Oceania | Fiji | unknown | unknown | Wong et al, 2015 |
| MDUST57 | ERR213275 | N/A | 2.3.5 | unknown | unknown | Blood | 1994 | Australia & Oceania | Oceania | Fiji | unknown | unknown | Wong et al, 2015 |
| MDUST58 | ERR213276 | N/A | 4.2.3 | unknown | unknown | Stool | 1994 | Australia & Oceania | Oceania | Fiji | unknown | unknown | Wong et al, 2015 |
| MDUST59 | ERR213277 | N/A | 4.2.3 | unknown | unknown | Stool | 1994 | Australia & Oceania | Oceania | Fiji | unknown | unknown | Wong et al, 2015 |
| MDUST60 | ERR213278 | N/A | 4.2.3 | unknown | unknown | Stool | 1994 | Australia & Oceania | Oceania | Fiji | unknown | unknown | Wong et al, 2015 |
| MDUST61 | ERR213279 | N/A | 4.2.3 | unknown | unknown | Stool | 1994 | Australia & Oceania | Oceania | Fiji | unknown | unknown | Wong et al, 2015 |
| MDUST62 | ERR213280 | N/A | 4.2.1 | unknown | unknown | Blood | 1994 | Australia & Oceania | Oceania | Fiji | unknown | unknown | Wong et al, 2015 |
| MDUST63 | ERR213281 | N/A | 2.3.5 | unknown | unknown | Stool | 1994 | Australia & Oceania | Oceania | Fiji | unknown | unknown | Wong et al, 2015 |
| MDUST64 | ERR213282 | N/A | 2.3.5 | unknown | unknown | Stool & Blood | 1994 | Australia & Oceania | Oceania | Fiji | unknown | unknown | Wong et al, 2015 |
| MDUST65 | ERR213283 | N/A | 2.3.5 | unknown | unknown | Stool | 1994 | Australia & Oceania | Oceania | Fiji | unknown | unknown | Wong et al, 2015 |
| MDUST66 | ERR213284 | N/A | 2.3.5 | unknown | unknown | Stool | 1994 | Australia & Oceania | Oceania | Fiji | unknown | unknown | Wong et al, 2015 |
| MDUST67 | ERR213285 | N/A | 4.3.1.1 | unknown | unknown | Stool | 1994 | Australia & Oceania | Oceania | Fiji | unknown | unknown | Wong et al, 2015 |
| MDUST68 | ERR213286 | N/A | 2.3.5 | unknown | unknown | Stool | 1994 | Australia & Oceania | Oceania | Fiji | unknown | unknown | Wong et al, 2015 |
| MDUST69 | ERR213287 | N/A | 4.2.3 | unknown | unknown | Stool | 1996 | Australia & Oceania | Oceania | Fiji | unknown | unknown | Wong et al, 2015 |
| MDUST70 | ERR213288 | N/A | 4.2.2 | unknown | unknown | Unknown | 2008 | Australia & Oceania | Oceania | Fiji | unknown | unknown | Wong et al, 2015 |
| MDUST71 | ERR213289 | N/A | 4.2.2 | unknown | unknown | Unknown | 2008 | Australia & Oceania | Oceania | Fiji | unknown | unknown | Wong et al, 2015 |
| MDUST72 | ERR213290 | N/A | 4.2.2 | unknown | unknown | Unknown | 2008 | Australia & Oceania | Oceania | Fiji | unknown | unknown | Wong et al, 2015 |
| Ke062 | ERR217406 | N/A | 4.3.1.1.EA1 | unknown | unknown | Blood | 2004 | Africa | East Africa | Kenya | unknown | unknown | Wong et al, 2015 |
| 1010996 | ERR279098 | N/A | 2.2 | unknown | unknown | Blood | 2011 | Africa | Southern Africa | Malawi | unknown | unknown | Wong et al, 2015 |

| | | | | | | | | | | | | | |
|---|---|---|---|---|---|---|---|---|---|---|---|---|---|
| 1014063 | ERR279099 | N/A | 4.3.1.1.EA1 | unknown | unknown | Blood | 2011 | Africa | Southern Africa | Malawi | unknown | unknown | Wong et al, 2015 |
| 1019695 | ERR279100 | N/A | 4.3.1.1.EA1 | unknown | unknown | Blood | 2011 | Africa | Southern Africa | Malawi | unknown | unknown | Wong et al, 2015 |
| 1019515 | ERR279101 | N/A | 4.3.1.1.EA1 | unknown | unknown | Blood | 2011 | Africa | Southern Africa | Malawi | unknown | unknown | Wong et al, 2015 |
| 1021678 | ERR279102 | N/A | 4.3.1.1.EA1 | unknown | unknown | Blood | 2011 | Africa | Southern Africa | Malawi | unknown | unknown | Wong et al, 2015 |
| 1025912 | ERR279103 | N/A | 4.3.1.1.EA1 | unknown | unknown | Blood | 2011 | Africa | Southern Africa | Malawi | unknown | unknown | Wong et al, 2015 |
| 1030208 | ERR279105 | N/A | 4.3.1.1.EA1 | unknown | unknown | Blood | 2012 | Africa | Southern Africa | Malawi | unknown | unknown | Wong et al, 2015 |
| 1008634 | ERR279106 | N/A | 4.3.1.1.EA1 | unknown | unknown | Blood | 2011 | Africa | Southern Africa | Malawi | unknown | unknown | Wong et al, 2015 |
| 1013486 | ERR279107 | N/A | 4.1.1 | unknown | unknown | Blood | 2011 | Africa | Southern Africa | Malawi | unknown | unknown | Wong et al, 2015 |
| 1021106 | ERR279109 | N/A | 4.3.1.1.EA1 | unknown | unknown | Blood | 2011 | Africa | Southern Africa | Malawi | unknown | unknown | Wong et al, 2015 |
| 1021701 | ERR279110 | N/A | 2.4.1 | unknown | unknown | Blood | 2011 | Africa | Southern Africa | Malawi | unknown | unknown | Wong et al, 2015 |
| 1025691 | ERR279111 | N/A | 2.4.1 | unknown | unknown | Blood | 2011 | Africa | Southern Africa | Malawi | unknown | unknown | Wong et al, 2015 |
| 1031809 | ERR279113 | N/A | 4.3.1.1.EA1 | unknown | unknown | Blood | 2012 | Africa | Southern Africa | Malawi | unknown | unknown | Wong et al, 2015 |
| 1007914 | ERR279114 | N/A | 4.3.1.1.EA1 | unknown | unknown | Blood | 2011 | Africa | Southern Africa | Malawi | unknown | unknown | Wong et al, 2015 |
| 1013305 | ERR279115 | N/A | 4.3.1.1 | unknown | unknown | Blood | 2011 | Africa | Southern Africa | Malawi | unknown | unknown | Wong et al, 2015 |
| 1016889 | ERR279116 | N/A | 4.3.1.1.EA1 | unknown | unknown | Blood | 2011 | Africa | Southern Africa | Malawi | unknown | unknown | Wong et al, 2015 |
| 1018216 | ERR279117 | N/A | 4.1.1 | unknown | unknown | Blood | 2011 | Africa | Southern Africa | Malawi | unknown | unknown | Wong et al, 2015 |
| 1023407 | ERR279118 | N/A | 4.3.1.1.EA1 | unknown | unknown | Blood | 2011 | Africa | Southern Africa | Malawi | unknown | unknown | Wong et al, 2015 |
| 1031785 | ERR279121 | N/A | 4.3.1.1.EA1 | unknown | unknown | Blood | 2012 | Africa | Southern Africa | Malawi | unknown | unknown | Wong et al, 2015 |
| 1007917 | ERR279122 | N/A | 4.3.1.1.EA1 | unknown | unknown | Blood | 2011 | Africa | Southern Africa | Malawi | unknown | unknown | Wong et al, 2015 |
| 1012911 | ERR279123 | N/A | 4.3.1.1.EA1 | unknown | unknown | Blood | 2011 | Africa | Southern Africa | Malawi | unknown | unknown | Wong et al, 2015 |
| 1016653 | ERR279124 | N/A | 4.3.1.1.EA1 | unknown | unknown | Blood | 2011 | Africa | Southern Africa | Malawi | unknown | unknown | Wong et al, 2015 |
| 1020791 | ERR279125 | N/A | 4.3.1.1.EA1 | unknown | unknown | Blood | 2011 | Africa | Southern Africa | Malawi | unknown | unknown | Wong et al, 2015 |
| 1022981 | ERR279126 | N/A | 4.1.1 | unknown | unknown | Blood | 2011 | Africa | Southern Africa | Malawi | unknown | unknown | Wong et al, 2015 |
| 1022553 | ERR279128 | N/A | 4.3.1.1.EA1 | unknown | unknown | Blood | 2011 | Africa | Southern Africa | Malawi | unknown | unknown | Wong et al, 2015 |
| 1030630 | ERR279129 | N/A | 4.3.1.1.EA1 | unknown | unknown | Blood | 2012 | Africa | Southern Africa | Malawi | unknown | unknown | Wong et al, 2015 |
| 1014451 | ERR279130 | N/A | 4.3.1.1.EA1 | unknown | unknown | Blood | 2011 | Africa | Southern Africa | Malawi | unknown | unknown | Wong et al, 2015 |
| 1018211 | ERR279131 | N/A | 4.3.1.1.EA1 | unknown | unknown | Blood | 2011 | Africa | Southern Africa | Malawi | unknown | unknown | Wong et al, 2015 |
| 1016605 | ERR279132 | N/A | 4.3.1.1.EA1 | unknown | unknown | Blood | 2011 | Africa | Southern Africa | Malawi | unknown | unknown | Wong et al, 2015 |
| 1021854 | ERR279133 | N/A | 4.3.1.1.EA1 | unknown | unknown | Blood | 2011 | Africa | Southern Africa | Malawi | unknown | unknown | Wong et al, 2015 |
| 1022075 | ERR279134 | N/A | 4.1.1 | unknown | unknown | Blood | 2011 | Africa | Southern Africa | Malawi | unknown | unknown | Wong et al, 2015 |
| 1032927 | ERR279136 | N/A | 4.3.1.1.EA1 | unknown | unknown | Blood | 2012 | Africa | Southern Africa | Malawi | unknown | unknown | Wong et al, 2015 |
| 1014383 | ERR279138 | N/A | 4.3.1.1.EA1 | unknown | unknown | Blood | 2011 | Africa | Southern Africa | Malawi | unknown | unknown | Wong et al, 2015 |
| 1017142 | ERR279139 | N/A | 4.1.1 | unknown | unknown | Blood | 2011 | Africa | Southern Africa | Malawi | unknown | unknown | Wong et al, 2015 |
| 1020184 | ERR279140 | N/A | 3.3.1 | unknown | unknown | Blood | 2011 | Africa | Southern Africa | Malawi | unknown | unknown | Wong et al, 2015 |
| 1023267 | ERR279141 | N/A | 2.4.1 | unknown | unknown | Blood | 2011 | Africa | Southern Africa | Malawi | unknown | unknown | Wong et al, 2015 |
| 1021624 | ERR279142 | N/A | 4.3.1.1.EA1 | unknown | unknown | Blood | 2011 | Africa | Southern Africa | Malawi | unknown | unknown | Wong et al, 2015 |
| 1024341 | ERR279143 | N/A | 4.3.1.1 | unknown | unknown | Blood | 2011 | Africa | Southern Africa | Malawi | unknown | unknown | Wong et al, 2015 |
| 1032463 | ERR279144 | N/A | 4.3.1.1.EA1 | unknown | unknown | Blood | 2012 | Africa | Southern Africa | Malawi | unknown | unknown | Wong et al, 2015 |
| A53009 | ERR279145 | N/A | 2.4.1 | unknown | unknown | Blood | 2008 | Africa | Southern Africa | Malawi | unknown | unknown | Wong et al, 2015 |
| 1013803 | ERR279146 | N/A | 4.3.1.1 | unknown | unknown | Blood | 2011 | Africa | Southern Africa | Malawi | unknown | unknown | Wong et al, 2015 |
| 1017137 | ERR279147 | N/A | 4.3.1.1.EA1 | unknown | unknown | Blood | 2011 | Africa | Southern Africa | Malawi | unknown | unknown | Wong et al, 2015 |
| 1019839 | ERR279148 | N/A | 4.3.1.1.EA1 | unknown | unknown | Blood | 2011 | Africa | Southern Africa | Malawi | unknown | unknown | Wong et al, 2015 |
| 1022401 | ERR279149 | N/A | 4.3.1.1.EA1 | unknown | unknown | Blood | 2011 | Africa | Southern Africa | Malawi | unknown | unknown | Wong et al, 2015 |
| 1020976 | ERR279150 | N/A | 4.3.1.1.EA1 | unknown | unknown | Blood | 2011 | Africa | Southern Africa | Malawi | unknown | unknown | Wong et al, 2015 |
| 1023999 | ERR279151 | N/A | 4.3.1.1.EA1 | unknown | unknown | Blood | 2011 | Africa | Southern Africa | Malawi | unknown | unknown | Wong et al, 2015 |
| 1032281 | ERR279152 | N/A | 4.3.1.1.EA1 | unknown | unknown | Blood | 2012 | Africa | Southern Africa | Malawi | unknown | unknown | Wong et al, 2015 |
| A53789 | ERR279153 | N/A | 4.3.1 | unknown | unknown | Blood | 2009 | Africa | Southern Africa | Malawi | unknown | unknown | Wong et al, 2015 |
| 1012752 | ERR279154 | N/A | 2.4.1 | unknown | unknown | Blood | 2011 | Africa | Southern Africa | Malawi | unknown | unknown | Wong et al, 2015 |
| 1020366 | ERR279155 | N/A | 4.3.1.1.EA1 | unknown | unknown | Blood | 2011 | Africa | Southern Africa | Malawi | unknown | unknown | Wong et al, 2015 |
| 1019833 | ERR279156 | N/A | 4.3.1.1.EA1 | unknown | unknown | Blood | 2011 | Africa | Southern Africa | Malawi | unknown | unknown | Wong et al, 2015 |

| | | | | | | | | | | | | | |
|---|---|---|---|---|---|---|---|---|---|---|---|---|---|
| 1029975 | ERR279158 | N/A | 4.3.1.1.EA1 | unknown | unknown | Blood | 2012 | Africa | Southern Africa | Malawi | unknown | unknown | Wong et al, 2015 |
| 1028143 | ERR279159 | N/A | 4.3.1.1.EA1 | unknown | unknown | Blood | 2012 | Africa | Southern Africa | Malawi | unknown | unknown | Wong et al, 2015 |
| 1032249 | ERR279160 | N/A | 4.3.1.1.EA1 | unknown | unknown | Blood | 2012 | Africa | Southern Africa | Malawi | unknown | unknown | Wong et al, 2015 |
| D50739 | ERR279161 | N/A | 2.4.1 | unknown | unknown | Blood | 2009 | Africa | Southern Africa | Malawi | unknown | unknown | Wong et al, 2015 |
| 1012727 | ERR279162 | N/A | 4.3.1.1.EA1 | unknown | unknown | Blood | 2011 | Africa | Southern Africa | Malawi | unknown | unknown | Wong et al, 2015 |
| 1020059 | ERR279163 | N/A | 4.3.1.1.EA1 | unknown | unknown | Blood | 2011 | Africa | Southern Africa | Malawi | unknown | unknown | Wong et al, 2015 |
| 1019806 | ERR279164 | N/A | 4.3.1.1.EA1 | unknown | unknown | Blood | 2011 | Africa | Southern Africa | Malawi | unknown | unknown | Wong et al, 2015 |
| 1018617 | ERR279165 | N/A | 4.3.1.1.EA1 | unknown | unknown | Blood | 2011 | Africa | Southern Africa | Malawi | unknown | unknown | Wong et al, 2015 |
| 1029451 | ERR279166 | N/A | 4.3.1.1.EA1 | unknown | unknown | Blood | 2012 | Africa | Southern Africa | Malawi | unknown | unknown | Wong et al, 2015 |
| 1032168 | ERR279168 | N/A | 4.3.1.1.EA1 | unknown | unknown | Blood | 2012 | Africa | Southern Africa | Malawi | unknown | unknown | Wong et al, 2015 |
| A55865 | ERR279169 | N/A | 2.2 | unknown | unknown | Blood | 2009 | Africa | Southern Africa | Malawi | unknown | unknown | Wong et al, 2015 |
| 1014152 | ERR279170 | N/A | 4.3.1.1.EA1 | unknown | unknown | Blood | 2011 | Africa | Southern Africa | Malawi | unknown | unknown | Wong et al, 2015 |
| 1021788 | ERR279173 | N/A | 4.3.1.1.EA1 | unknown | unknown | Blood | 2011 | Africa | Southern Africa | Malawi | unknown | unknown | Wong et al, 2015 |
| 1028963 | ERR279174 | N/A | 4.3.1.1.EA1 | unknown | unknown | Blood | 2012 | Africa | Southern Africa | Malawi | unknown | unknown | Wong et al, 2015 |
| 1026460 | ERR279175 | N/A | 4.3.1.1.EA1 | unknown | unknown | Blood | 2012 | Africa | Southern Africa | Malawi | unknown | unknown | Wong et al, 2015 |
| 1031581 | ERR279176 | N/A | 4.3.1.1.EA1 | unknown | unknown | Blood | 2012 | Africa | Southern Africa | Malawi | unknown | unknown | Wong et al, 2015 |
| A32420 | ERR279177 | N/A | 2.2 | unknown | unknown | Blood | 2004 | Africa | Southern Africa | Malawi | unknown | unknown | Wong et al, 2015 |
| A33112 | ERR279178 | N/A | 2 | unknown | unknown | Blood | 2005 | Africa | Southern Africa | Malawi | unknown | unknown | Wong et al, 2015 |
| A40201 | ERR279182 | N/A | 2.4.1 | unknown | unknown | Blood | 2006 | Africa | Southern Africa | Malawi | unknown | unknown | Wong et al, 2015 |
| D28995 | ERR279187 | N/A | 2.4.1 | unknown | unknown | Blood | 2005 | Africa | Southern Africa | Malawi | unknown | unknown | Wong et al, 2015 |
| A39483 | ERR279189 | N/A | 4.1.1 | unknown | unknown | Blood | 2006 | Africa | Southern Africa | Malawi | unknown | unknown | Wong et al, 2015 |
| A40566 | ERR279190 | N/A | 2.4.1 | unknown | unknown | Blood | 2006 | Africa | Southern Africa | Malawi | unknown | unknown | Wong et al, 2015 |
| D41342 | ERR279191 | N/A | 4.1.1 | unknown | unknown | Blood | 2007 | Africa | Southern Africa | Malawi | unknown | unknown | Wong et al, 2015 |
| D7558 | ERR279344 | N/A | 4.3.1 | unknown | unknown | Not provided | 2006 | Asia | South Asia | India | unknown | unknown | Wong et al, 2015 |
| D7649 | ERR279345 | N/A | 4.3.1 | unknown | unknown | Not provided | 2007 | Asia | South Asia | India | unknown | unknown | Wong et al, 2015 |
| C3551 | ERR279346 | N/A | 4.3.1.2 | unknown | unknown | Not provided | 2005 | Asia | South Asia | India | unknown | unknown | Wong et al, 2015 |
| C3891 | ERR279347 | N/A | 4.3.1.2 | unknown | unknown | Not provided | 2005 | Asia | South Asia | India | unknown | unknown | Wong et al, 2015 |
| C3495 | ERR279348 | N/A | 4.3.1.1 | unknown | unknown | Not provided | 2005 | Asia | South Asia | India | unknown | unknown | Wong et al, 2015 |
| C3634 | ERR279349 | N/A | 4.3.1.1 | unknown | unknown | Not provided | 2005 | Asia | South Asia | India | unknown | unknown | Wong et al, 2015 |
| E2889 | ERR279350 | N/A | 4.3.1.2 | unknown | unknown | Not provided | 2006 | Asia | South Asia | India | unknown | unknown | Wong et al, 2015 |
| E2990 | ERR279351 | N/A | 4.3.1.2 | unknown | unknown | Not provided | 2006 | Asia | South Asia | India | unknown | unknown | Wong et al, 2015 |
| E1240 | ERR279352 | N/A | 2.2.4 | unknown | unknown | Not provided | 2004 | Asia | South Asia | India | unknown | unknown | Wong et al, 2015 |
| E1303 | ERR279353 | N/A | 2.2.4 | unknown | unknown | Not provided | 2004 | Asia | South Asia | India | unknown | unknown | Wong et al, 2015 |
| Quailes | ERR294858 | N/A | 3.1 | unknown | unknown | Gallbladder fluid | 1958 | North America | North America | Jnited States of Americ | unknown | unknown | Wong et al, 2015 |
| 2008-001909 | ERR319402 | N/A | 4.3.1.1 | unknown | unknown | Blood | 2008 | Asia | Southeast Asia | Cambodia | unknown | unknown | Wong et al, 2015 |
| 2008-002399 | ERR319403 | N/A | 4.3.1.1 | unknown | unknown | Blood | 2008 | Asia | Southeast Asia | Cambodia | unknown | unknown | Wong et al, 2015 |
| 2008-003956 | ERR319404 | N/A | 4.3.1.1 | unknown | unknown | Blood | 2008 | Asia | Southeast Asia | Cambodia | unknown | unknown | Wong et al, 2015 |
| 2008-004254 | ERR319405 | N/A | 4.3.1.1 | unknown | unknown | Blood | 2008 | Asia | Southeast Asia | Cambodia | unknown | unknown | Wong et al, 2015 |
| 2002-216507 | ERR319406 | N/A | 4.3.1.1 | unknown | unknown | Blood | 2008 | Asia | Southeast Asia | Cambodia | unknown | unknown | Wong et al, 2015 |
| 2003-016955 | ERR319408 | N/A | 4.3.1.1 | unknown | unknown | Blood | 2008 | Asia | Southeast Asia | Cambodia | unknown | unknown | Wong et al, 2015 |
| 2008-006312 | ERR319409 | N/A | 4.3.1.1 | unknown | unknown | Blood | 2008 | Asia | Southeast Asia | Cambodia | unknown | unknown | Wong et al, 2015 |
| 2005-008425 | ERR319410 | N/A | 4.3.1.1 | unknown | unknown | Blood | 2008 | Asia | Southeast Asia | Cambodia | unknown | unknown | Wong et al, 2015 |
| 2008-006300 | ERR319411 | N/A | 4.3.1.1 | unknown | unknown | Blood | 2008 | Asia | Southeast Asia | Cambodia | unknown | unknown | Wong et al, 2015 |
| 2009-000156 | ERR319412 | N/A | 4.3.1.1 | unknown | unknown | Blood | 2009 | Asia | Southeast Asia | Cambodia | unknown | unknown | Wong et al, 2015 |
| 2009-000892 | ERR319413 | N/A | 4.3.1.1 | unknown | unknown | Blood | 2009 | Asia | Southeast Asia | Cambodia | unknown | unknown | Wong et al, 2015 |
| 2008-002960 | ERR319414 | N/A | 4.3.1.1 | unknown | unknown | Blood | 2009 | Asia | Southeast Asia | Cambodia | unknown | unknown | Wong et al, 2015 |
| 2009-004002 | ERR319415 | N/A | 4.3.1.1 | unknown | unknown | Blood | 2009 | Asia | Southeast Asia | Cambodia | unknown | unknown | Wong et al, 2015 |
| 2009-006046 | ERR319416 | N/A | 4.3.1.1 | unknown | unknown | Blood | 2009 | Asia | Southeast Asia | Cambodia | unknown | unknown | Wong et al, 2015 |
| 2009-007146 | ERR319417 | N/A | 4.3.1.1 | unknown | unknown | Blood | 2009 | Asia | Southeast Asia | Cambodia | unknown | unknown | Wong et al, 2015 |
| 2005-010587 | ERR319418 | N/A | 3.2.1 | unknown | unknown | Blood | 2009 | Asia | Southeast Asia | Cambodia | unknown | unknown | Wong et al, 2015 |

| | | | | | | | | | | | | | |
|---|---|---|---|---|---|---|---|---|---|---|---|---|---|
| 2003-008924 | ERR319419 | N/A | 4.3.1.1 | unknown | unknown | Blood | 2009 | Asia | Southeast Asia | Cambodia | unknown | unknown | Wong et al, 2015 |
| 2009-011935 | ERR319420 | N/A | 4.3.1.1 | unknown | unknown | Blood | 2009 | Asia | Southeast Asia | Cambodia | unknown | unknown | Wong et al, 2015 |
| 2009-012898 | ERR319421 | N/A | 4.3.1.1 | unknown | unknown | Blood | 2009 | Asia | Southeast Asia | Cambodia | unknown | unknown | Wong et al, 2015 |
| 2010-005196 | ERR319422 | N/A | 4.3.1.1 | unknown | unknown | Blood | 2010 | Asia | Southeast Asia | Cambodia | unknown | unknown | Wong et al, 2015 |
| 2010-011361 | ERR319423 | N/A | 4.3.1.1 | unknown | unknown | Blood | 2010 | Asia | Southeast Asia | Cambodia | unknown | unknown | Wong et al, 2015 |
| 2010-005872 | ERR319424 | N/A | 4.3.1.1 | unknown | unknown | Blood | 2010 | Asia | Southeast Asia | Cambodia | unknown | unknown | Wong et al, 2015 |
| 2006-011021 | ERR319425 | N/A | 4.3.1.1 | unknown | unknown | Blood | 2010 | Asia | Southeast Asia | Cambodia | unknown | unknown | Wong et al, 2015 |
| 2010-006551 | ERR319426 | N/A | 4.3.1.1 | unknown | unknown | Blood | 2010 | Asia | Southeast Asia | Cambodia | unknown | unknown | Wong et al, 2015 |
| 2007-013365 | ERR319427 | N/A | 4.3.1.1 | unknown | unknown | Blood | 2010 | Asia | Southeast Asia | Cambodia | unknown | unknown | Wong et al, 2015 |
| 2010-006866 | ERR319428 | N/A | 4.3.1.1 | unknown | unknown | Blood | 2010 | Asia | Southeast Asia | Cambodia | unknown | unknown | Wong et al, 2015 |
| 2007-012990 | ERR319429 | N/A | 4.3.1.1 | unknown | unknown | Blood | 2010 | Asia | Southeast Asia | Cambodia | unknown | unknown | Wong et al, 2015 |
| 2001-100510 | ERR319431 | N/A | 4.3.1.1 | unknown | unknown | Blood | 2010 | Asia | Southeast Asia | Cambodia | unknown | unknown | Wong et al, 2015 |
| 2010-007898 | ERR319432 | N/A | 4.3.1.1 | unknown | unknown | Blood | 2010 | Asia | Southeast Asia | Cambodia | unknown | unknown | Wong et al, 2015 |
| 2010-008154 | ERR319433 | N/A | 4.3.1.1 | unknown | unknown | Blood | 2010 | Asia | Southeast Asia | Cambodia | unknown | unknown | Wong et al, 2015 |
| 2010-007462 | ERR319434 | N/A | 4.3.1.1 | unknown | unknown | Blood | 2010 | Asia | Southeast Asia | Cambodia | unknown | unknown | Wong et al, 2015 |
| 2010-002605 | ERR319435 | N/A | 4.3.1.1 | unknown | unknown | Blood | 2010 | Asia | Southeast Asia | Cambodia | unknown | unknown | Wong et al, 2015 |
| 2010-009987 | ERR319436 | N/A | 4.3.1.1 | unknown | unknown | Blood | 2010 | Asia | Southeast Asia | Cambodia | unknown | unknown | Wong et al, 2015 |
| 2010-010162 | ERR319437 | N/A | 4.3.1.1 | unknown | unknown | Blood | 2010 | Asia | Southeast Asia | Cambodia | unknown | unknown | Wong et al, 2015 |
| 2010-002948 | ERR319438 | N/A | 4.3.1.1 | unknown | unknown | Blood | 2010 | Asia | Southeast Asia | Cambodia | unknown | unknown | Wong et al, 2015 |
| 2010-010972 | ERR319439 | N/A | 4.3.1.1 | unknown | unknown | Blood | 2010 | Asia | Southeast Asia | Cambodia | unknown | unknown | Wong et al, 2015 |
| 2010-011187 | ERR319440 | N/A | 4.3.1.1 | unknown | unknown | Blood | 2010 | Asia | Southeast Asia | Cambodia | unknown | unknown | Wong et al, 2015 |
| 2010-011712 | ERR319441 | N/A | 4.3.1.1 | unknown | unknown | Blood | 2010 | Asia | Southeast Asia | Cambodia | unknown | unknown | Wong et al, 2015 |
| 01-2011-000112 | ERR319442 | N/A | 4.3.1.1 | unknown | unknown | Blood | 2011 | Asia | Southeast Asia | Cambodia | unknown | unknown | Wong et al, 2015 |
| 01-2011-000483 | ERR319443 | N/A | 4.3.1.1 | unknown | unknown | Blood | 2011 | Asia | Southeast Asia | Cambodia | unknown | unknown | Wong et al, 2015 |
| 2005-009945 | ERR319444 | N/A | 4.3.1.1 | unknown | unknown | Blood | 2011 | Asia | Southeast Asia | Cambodia | unknown | unknown | Wong et al, 2015 |
| 2007-021515 | ERR319445 | N/A | 4.3.1.1 | unknown | unknown | Blood | 2011 | Asia | Southeast Asia | Cambodia | unknown | unknown | Wong et al, 2015 |
| 2006-007642 | ERR319446 | N/A | 4.3.1.1 | unknown | unknown | Blood | 2011 | Asia | Southeast Asia | Cambodia | unknown | unknown | Wong et al, 2015 |
| 2002-216622 | ERR319447 | N/A | 4.3.1.1 | unknown | unknown | Blood | 2011 | Asia | Southeast Asia | Cambodia | unknown | unknown | Wong et al, 2015 |
| 2010-004004 | ERR319448 | N/A | 4.3.1.1 | unknown | unknown | Blood | 2011 | Asia | Southeast Asia | Cambodia | unknown | unknown | Wong et al, 2015 |
| 01-2010-004103 | ERR319449 | N/A | 4.3.1.1 | unknown | unknown | Blood | 2011 | Asia | Southeast Asia | Cambodia | unknown | unknown | Wong et al, 2015 |
| 01-2011-000113 | ERR319450 | N/A | 4.3.1.1 | unknown | unknown | Blood | 2011 | Asia | Southeast Asia | Cambodia | unknown | unknown | Wong et al, 2015 |
| 01-2011-000309 | ERR319451 | N/A | 4.3.1.1 | unknown | unknown | Blood | 2011 | Asia | Southeast Asia | Cambodia | unknown | unknown | Wong et al, 2015 |
| 2010-021566 | ERR319452 | N/A | 4.3.1.1 | unknown | unknown | Blood | 2011 | Asia | Southeast Asia | Cambodia | unknown | unknown | Wong et al, 2015 |
| 2011-007445 | ERR319453 | N/A | 4.3.1.1 | unknown | unknown | Blood | 2011 | Asia | Southeast Asia | Cambodia | unknown | unknown | Wong et al, 2015 |
| 2005-011215 | ERR319454 | N/A | 4.3.1.1 | unknown | unknown | Blood | 2011 | Asia | Southeast Asia | Cambodia | unknown | unknown | Wong et al, 2015 |
| 2009-000374 | ERR319455 | N/A | 4.3.1.1 | unknown | unknown | Blood | 2011 | Asia | Southeast Asia | Cambodia | unknown | unknown | Wong et al, 2015 |
| 01-2011-001522 | ERR319456 | N/A | 4.3.1.1 | unknown | unknown | Blood | 2011 | Asia | Southeast Asia | Cambodia | unknown | unknown | Wong et al, 2015 |
| 2011-007660 | ERR319457 | N/A | 4.3.1.1 | unknown | unknown | Blood | 2011 | Asia | Southeast Asia | Cambodia | unknown | unknown | Wong et al, 2015 |
| 2011-007792 | ERR319458 | N/A | 4.3.1.1 | unknown | unknown | Blood | 2011 | Asia | Southeast Asia | Cambodia | unknown | unknown | Wong et al, 2015 |
| 2011-007930 | ERR319459 | N/A | 4.3.1.1 | unknown | unknown | Blood | 2011 | Asia | Southeast Asia | Cambodia | unknown | unknown | Wong et al, 2015 |
| 2006-008616 | ERR319460 | N/A | 4.3.1.1 | unknown | unknown | Blood | 2011 | Asia | Southeast Asia | Cambodia | unknown | unknown | Wong et al, 2015 |
| 2007-015726 | ERR319461 | N/A | 4.3.1.1 | unknown | unknown | Blood | 2011 | Asia | Southeast Asia | Cambodia | unknown | unknown | Wong et al, 2015 |
| BC2011-1325 | ERR319462 | N/A | 3.2.1 | unknown | unknown | Blood | 2011 | Asia | Southeast Asia | Cambodia | unknown | unknown | Wong et al, 2015 |
| BC2011-1311 | ERR319463 | N/A | 4.3.1.1 | unknown | unknown | Blood | 2011 | Asia | Southeast Asia | Cambodia | unknown | unknown | Wong et al, 2015 |
| BC2011-1307 | ERR319464 | N/A | 4.3.1.1 | unknown | unknown | Blood | 2011 | Asia | Southeast Asia | Cambodia | unknown | unknown | Wong et al, 2015 |
| BC2011-1305 | ERR319465 | N/A | 4.3.1.1 | unknown | unknown | Blood | 2011 | Asia | Southeast Asia | Cambodia | unknown | unknown | Wong et al, 2015 |
| BC20111457 | ERR319466 | N/A | 4.3.1.1 | unknown | unknown | Blood | 2011 | Asia | Southeast Asia | Cambodia | unknown | unknown | Wong et al, 2015 |
| BC2011-1461 | ERR319467 | N/A | 4.3.1.1 | unknown | unknown | Blood | 2011 | Asia | Southeast Asia | Cambodia | unknown | unknown | Wong et al, 2015 |
| BC2011-1534 | ERR319468 | N/A | 4.3.1.1 | unknown | unknown | Blood | 2011 | Asia | Southeast Asia | Cambodia | unknown | unknown | Wong et al, 2015 |
| BC2011-1612 | ERR319469 | N/A | 4.3.1.1 | unknown | unknown | Blood | 2011 | Asia | Southeast Asia | Cambodia | unknown | unknown | Wong et al, 2015 |

| | | | | | | | | | | | | | |
|---|---|---|---|---|---|---|---|---|---|---|---|---|---|
| BC2011-1683 | ERR319470 | N/A | 4.3.1.1 | unknown | unknown | Blood | 2011 | Asia | Southeast Asia | Cambodia | unknown | unknown | Wong et al, 2015 |
| BC2011-1739 | ERR319471 | N/A | 4.3.1.1 | unknown | unknown | Blood | 2011 | Asia | Southeast Asia | Cambodia | unknown | unknown | Wong et al, 2015 |
| 2011-018938 | ERR319472 | N/A | 4.3.1.1 | unknown | unknown | Blood | 2012 | Asia | Southeast Asia | Cambodia | unknown | unknown | Wong et al, 2015 |
| 2012-002759 | ERR319473 | N/A | 4.3.1.1 | unknown | unknown | Stool | 2012 | Asia | Southeast Asia | Cambodia | unknown | unknown | Wong et al, 2015 |
| 01-2010-002043 | ERR319474 | N/A | 4.3.1.1 | unknown | unknown | Blood | 2012 | Asia | Southeast Asia | Cambodia | unknown | unknown | Wong et al, 2015 |
| 01-2012-001846 | ERR319475 | N/A | 4.3.1.1 | unknown | unknown | Blood | 2012 | Asia | Southeast Asia | Cambodia | unknown | unknown | Wong et al, 2015 |
| 2012-007197 | ERR319476 | N/A | 4.3.1.1 | unknown | unknown | Blood | 2012 | Asia | Southeast Asia | Cambodia | unknown | unknown | Wong et al, 2015 |
| 01-2012-002175 | ERR319477 | N/A | 4.3.1.1 | unknown | unknown | Blood | 2012 | Asia | Southeast Asia | Cambodia | unknown | unknown | Wong et al, 2015 |
| 2012-008606 | ERR319478 | N/A | 4.3.1.1 | unknown | unknown | Blood | 2012 | Asia | Southeast Asia | Cambodia | unknown | unknown | Wong et al, 2015 |
| 2012-009077 | ERR319479 | N/A | 4.3.1.1 | unknown | unknown | Blood | 2012 | Asia | Southeast Asia | Cambodia | unknown | unknown | Wong et al, 2015 |
| 01-2012-002473 | ERR319480 | N/A | 4.3.1.1 | unknown | unknown | Blood | 2012 | Asia | Southeast Asia | Cambodia | unknown | unknown | Wong et al, 2015 |
| 01-2010-005183 | ERR319481 | N/A | 4.3.1.1 | unknown | unknown | Blood | 2012 | Asia | Southeast Asia | Cambodia | unknown | unknown | Wong et al, 2015 |
| 01-2012-002529 | ERR319482 | N/A | 4.3.1.1 | unknown | unknown | Blood | 2012 | Asia | Southeast Asia | Cambodia | unknown | unknown | Wong et al, 2015 |
| 2006-014301 | ERR319483 | N/A | 4.3.1.1 | unknown | unknown | Blood | 2012 | Asia | Southeast Asia | Cambodia | unknown | unknown | Wong et al, 2015 |
| 2012-009618 | ERR319484 | N/A | 4.3.1.1 | unknown | unknown | Blood | 2012 | Asia | Southeast Asia | Cambodia | unknown | unknown | Wong et al, 2015 |
| 2008-017452 | ERR319485 | N/A | 4.3.1.1 | unknown | unknown | Blood | 2012 | Asia | Southeast Asia | Cambodia | unknown | unknown | Wong et al, 2015 |
| 2012-010059 | ERR319486 | N/A | 4.3.1.1 | unknown | unknown | Blood | 2012 | Asia | Southeast Asia | Cambodia | unknown | unknown | Wong et al, 2015 |
| 2012-010060 | ERR319487 | N/A | 4.3.1.1 | unknown | unknown | Blood | 2012 | Asia | Southeast Asia | Cambodia | unknown | unknown | Wong et al, 2015 |
| 01-2012-002669 | ERR319488 | N/A | 4.3.1.1 | unknown | unknown | Blood | 2012 | Asia | Southeast Asia | Cambodia | unknown | unknown | Wong et al, 2015 |
| 2010-006045 | ERR319489 | N/A | 4.3.1.1 | unknown | unknown | Blood | 2012 | Asia | Southeast Asia | Cambodia | unknown | unknown | Wong et al, 2015 |
| 01-2012-002666 | ERR319490 | N/A | 4.3.1.1 | unknown | unknown | Blood | 2012 | Asia | Southeast Asia | Cambodia | unknown | unknown | Wong et al, 2015 |
| 2012-009913 | ERR319492 | N/A | 4.3.1.1 | unknown | unknown | Blood | 2012 | Asia | Southeast Asia | Cambodia | unknown | unknown | Wong et al, 2015 |
| 2005-010317 | ERR319494 | N/A | 4.3.1.1 | unknown | unknown | Blood | 2012 | Asia | Southeast Asia | Cambodia | unknown | unknown | Wong et al, 2015 |
| 2012-010406 | ERR319495 | N/A | 4.3.1.1 | unknown | unknown | Blood | 2012 | Asia | Southeast Asia | Cambodia | unknown | unknown | Wong et al, 2015 |
| A140 | ERR326598 | N/A | 2.1.8 | unknown | unknown | Blood | 2010 | Asia | Southeast Asia | Indonesia | unknown | unknown | Wong et al, 2015 |
| A268 | ERR326599 | N/A | 4.1 | unknown | unknown | Blood | 2009 | Asia | Southeast Asia | Indonesia | unknown | unknown | Wong et al, 2015 |
| A270 | ERR326600 | N/A | 1 | unknown | unknown | Blood | 2010 | Asia | Southeast Asia | Indonesia | unknown | unknown | Wong et al, 2015 |
| A271 | ERR326601 | N/A | 4.1 | unknown | unknown | Blood | 2009 | Asia | Southeast Asia | Indonesia | unknown | unknown | Wong et al, 2015 |
| A272 | ERR326602 | N/A | 4.1 | unknown | unknown | Blood | 2010 | Asia | Southeast Asia | Indonesia | unknown | unknown | Wong et al, 2015 |
| A273 | ERR326603 | N/A | 4.1 | unknown | unknown | Blood | 2009 | Asia | Southeast Asia | Indonesia | unknown | unknown | Wong et al, 2015 |
| A274 | ERR326604 | N/A | 3 | unknown | unknown | Blood | 2006 | Asia | Southeast Asia | Indonesia | unknown | unknown | Wong et al, 2015 |
| A275 | ERR326605 | N/A | 4.1 | unknown | unknown | Blood | 2009 | Asia | Southeast Asia | Indonesia | unknown | unknown | Wong et al, 2015 |
| A276 | ERR326606 | N/A | 4.1 | unknown | unknown | Blood | 2008 | Asia | Southeast Asia | Indonesia | unknown | unknown | Wong et al, 2015 |
| A277 | ERR326607 | N/A | 4.1 | unknown | unknown | Blood | 2009 | Asia | Southeast Asia | Indonesia | unknown | unknown | Wong et al, 2015 |
| A278 | ERR326608 | N/A | 4.1 | unknown | unknown | Blood | 2009 | Asia | Southeast Asia | Indonesia | unknown | unknown | Wong et al, 2015 |
| A281 | ERR326610 | N/A | 3 | unknown | unknown | Blood | 2008 | Asia | Southeast Asia | Indonesia | unknown | unknown | Wong et al, 2015 |
| A282 | ERR326611 | N/A | 3.1.2 | unknown | unknown | Blood | 2010 | Asia | Southeast Asia | Indonesia | unknown | unknown | Wong et al, 2015 |
| A283 | ERR326612 | N/A | 4.1 | unknown | unknown | Blood | 2010 | Asia | Southeast Asia | Indonesia | unknown | unknown | Wong et al, 2015 |
| A284 | ERR326613 | N/A | 3 | unknown | unknown | Blood | 2011 | Asia | Southeast Asia | Indonesia | unknown | unknown | Wong et al, 2015 |
| A285 | ERR326614 | N/A | 4.1 | unknown | unknown | Blood | 2010 | Asia | Southeast Asia | Indonesia | unknown | unknown | Wong et al, 2015 |
| A286 | ERR326615 | N/A | 2.1.6 | unknown | unknown | Blood | 2010 | Asia | Southeast Asia | Indonesia | unknown | unknown | Wong et al, 2015 |
| A287 | ERR326616 | N/A | 3.1.2 | unknown | unknown | Blood | 2008 | Asia | Southeast Asia | Indonesia | unknown | unknown | Wong et al, 2015 |
| A340 | ERR326618 | N/A | 2.1.9 | unknown | unknown | Blood | 2010 | Asia | Southeast Asia | Indonesia | unknown | unknown | Wong et al, 2015 |
| A341 | ERR326619 | N/A | 3.1.2 | unknown | unknown | Blood | 2008 | Asia | Southeast Asia | Indonesia | unknown | unknown | Wong et al, 2015 |
| A344-1st | ERR326622 | N/A | 4.1 | unknown | unknown | Blood | 2010 | Asia | Southeast Asia | Indonesia | unknown | unknown | Wong et al, 2015 |
| A347 | ERR326623 | N/A | 2.1.8 | unknown | unknown | Blood | 2007 | Asia | Southeast Asia | Indonesia | unknown | unknown | Wong et al, 2015 |
| A348 | ERR326624 | N/A | 2.1.9 | unknown | unknown | Blood | 2012 | Asia | Southeast Asia | Indonesia | unknown | unknown | Wong et al, 2015 |
| A350 | ERR326625 | N/A | 2.1.6 | unknown | unknown | Blood | 2010 | Asia | Southeast Asia | Indonesia | unknown | unknown | Wong et al, 2015 |
| A352 | ERR326626 | N/A | 3 | unknown | unknown | Blood | 2010 | Asia | Southeast Asia | Indonesia | unknown | unknown | Wong et al, 2015 |
| A354 | ERR326627 | N/A | 3 | unknown | unknown | Blood | 2011 | Asia | Southeast Asia | Indonesia | unknown | unknown | Wong et al, 2015 |

| | | | | | | | | | | | | | |
|---|---|---|---|---|---|---|---|---|---|---|---|---|---|
| A356 | ERR326629 | N/A | 2.1.9 | unknown | unknown | Blood | 2010 | Asia | Southeast Asia | Indonesia | unknown | unknown | Wong et al, 2015 |
| A357 | ERR326630 | N/A | 3.1.2 | unknown | unknown | Blood | 2009 | Asia | Southeast Asia | Indonesia | unknown | unknown | Wong et al, 2015 |
| A358 | ERR326631 | N/A | 4.1 | unknown | unknown | Blood | 2009 | Asia | Southeast Asia | Indonesia | unknown | unknown | Wong et al, 2015 |
| A359 | ERR326632 | N/A | 3.1.2 | unknown | unknown | Blood | 2010 | Asia | Southeast Asia | Indonesia | unknown | unknown | Wong et al, 2015 |
| A360 | ERR326633 | N/A | 4.1 | unknown | unknown | Blood | 2009 | Asia | Southeast Asia | Indonesia | unknown | unknown | Wong et al, 2015 |
| A363 | ERR326634 | N/A | 3.1.2 | unknown | unknown | Blood | 2009 | Asia | Southeast Asia | Indonesia | unknown | unknown | Wong et al, 2015 |
| A365 | ERR326635 | N/A | 2.1.9 | unknown | unknown | Blood | 2007 | Asia | Southeast Asia | Indonesia | unknown | unknown | Wong et al, 2015 |
| A366 | ERR326636 | N/A | 2.1.6 | unknown | unknown | Blood | 2009 | Asia | Southeast Asia | Indonesia | unknown | unknown | Wong et al, 2015 |
| A368 | ERR326637 | N/A | 2.1.9 | unknown | unknown | Blood | 2009 | Asia | Southeast Asia | Indonesia | unknown | unknown | Wong et al, 2015 |
| A369 | ERR326638 | N/A | 4.1 | unknown | unknown | Blood | 2008 | Asia | Southeast Asia | Indonesia | unknown | unknown | Wong et al, 2015 |
| A370 | ERR326639 | N/A | 3.1.2 | unknown | unknown | Blood | 2010 | Asia | Southeast Asia | Indonesia | unknown | unknown | Wong et al, 2015 |
| A373 | ERR326640 | N/A | 2.1.8 | unknown | unknown | Blood | 2008 | Asia | Southeast Asia | Indonesia | unknown | unknown | Wong et al, 2015 |
| A374 | ERR326641 | N/A | 4.1 | unknown | unknown | Blood | 2008 | Asia | Southeast Asia | Indonesia | unknown | unknown | Wong et al, 2015 |
| A375 | ERR326642 | N/A | 2.1.9 | unknown | unknown | Blood | 2005 | Asia | Southeast Asia | Indonesia | unknown | unknown | Wong et al, 2015 |
| A377 | ERR326644 | N/A | 2.1.9 | unknown | unknown | Blood | 2010 | Asia | Southeast Asia | Indonesia | unknown | unknown | Wong et al, 2015 |
| A378 | ERR326645 | N/A | 2.1.6 | unknown | unknown | Blood | 2008 | Asia | Southeast Asia | Indonesia | unknown | unknown | Wong et al, 2015 |
| A379 | ERR326646 | N/A | 4.1 | unknown | unknown | Blood | 2010 | Asia | Southeast Asia | Indonesia | unknown | unknown | Wong et al, 2015 |
| A380 | ERR326647 | N/A | 3 | unknown | unknown | Blood | 2006 | Asia | Southeast Asia | Indonesia | unknown | unknown | Wong et al, 2015 |
| A382 | ERR326648 | N/A | 2.1.9 | unknown | unknown | Blood | 2009 | Asia | Southeast Asia | Indonesia | unknown | unknown | Wong et al, 2015 |
| A385 | ERR326649 | N/A | 4.1 | unknown | unknown | Blood | 2012 | Asia | Southeast Asia | Indonesia | unknown | unknown | Wong et al, 2015 |
| A389 | ERR326650 | N/A | 2.1.8 | unknown | unknown | Blood | 2006 | Asia | Southeast Asia | Indonesia | unknown | unknown | Wong et al, 2015 |
| A390 | ERR326651 | N/A | 4.1 | unknown | unknown | Blood | 2007 | Asia | Southeast Asia | Indonesia | unknown | unknown | Wong et al, 2015 |
| A392 | ERR326652 | N/A | 4.1 | unknown | unknown | Blood | 2010 | Asia | Southeast Asia | Indonesia | unknown | unknown | Wong et al, 2015 |
| A394 | ERR326653 | N/A | 2.1.9 | unknown | unknown | Blood | 2007 | Asia | Southeast Asia | Indonesia | unknown | unknown | Wong et al, 2015 |
| A395 | ERR326654 | N/A | 4.1 | unknown | unknown | Blood | 2009 | Asia | Southeast Asia | Indonesia | unknown | unknown | Wong et al, 2015 |
| A396 | ERR326655 | N/A | 4.1 | unknown | unknown | Blood | 2009 | Asia | Southeast Asia | Indonesia | unknown | unknown | Wong et al, 2015 |
| A399 | ERR326656 | N/A | 3.1.2 | unknown | unknown | Blood | 2009 | Asia | Southeast Asia | Indonesia | unknown | unknown | Wong et al, 2015 |
| KM1646 | ERR326657 | N/A | 3.4 | unknown | unknown | Blood | 2010 | Asia | Southeast Asia | Laos | unknown | unknown | Wong et al, 2015 |
| KM1647 | ERR326658 | N/A | 3.4 | unknown | unknown | Blood | 2010 | Asia | Southeast Asia | Laos | unknown | unknown | Wong et al, 2015 |
| KM1648 | ERR326659 | N/A | 3.4 | unknown | unknown | Blood | 2010 | Asia | Southeast Asia | Laos | unknown | unknown | Wong et al, 2015 |
| LNT1197 | ERR326660 | N/A | 2.3.4 | unknown | unknown | Blood | 2009 | Asia | Southeast Asia | Laos | unknown | unknown | Wong et al, 2015 |
| LNT12 | ERR326661 | N/A | 3.4 | unknown | unknown | Blood | 2007 | Asia | Southeast Asia | Laos | unknown | unknown | Wong et al, 2015 |
| LNT13 | ERR326662 | N/A | 3.4 | unknown | unknown | Blood | 2007 | Asia | Southeast Asia | Laos | unknown | unknown | Wong et al, 2015 |
| LNT1339 | ERR326663 | N/A | 3.4 | unknown | unknown | Blood | 2010 | Asia | Southeast Asia | Laos | unknown | unknown | Wong et al, 2015 |
| LNT1360 | ERR326664 | N/A | 2.4 | unknown | unknown | Blood | 2010 | Asia | Southeast Asia | Laos | unknown | unknown | Wong et al, 2015 |
| LNT1365 | ERR326665 | N/A | 2.3.4 | unknown | unknown | Blood | 2010 | Asia | Southeast Asia | Laos | unknown | unknown | Wong et al, 2015 |
| LNT1367-3 | ERR326666 | N/A | 2.3.4 | unknown | unknown | Blood | 2010 | Asia | Southeast Asia | Laos | unknown | unknown | Wong et al, 2015 |
| LNT1374 | ERR326667 | N/A | 3.4 | unknown | unknown | Blood | 2010 | Asia | Southeast Asia | Laos | unknown | unknown | Wong et al, 2015 |
| LNT1377 | ERR326668 | N/A | 2.3.4 | unknown | unknown | Blood | 2010 | Asia | Southeast Asia | Laos | unknown | unknown | Wong et al, 2015 |
| LNT1378 | ERR326669 | N/A | 3.4 | unknown | unknown | Blood | 2010 | Asia | Southeast Asia | Laos | unknown | unknown | Wong et al, 2015 |
| LNT1426 | ERR326670 | N/A | 3.4 | unknown | unknown | Blood | 2010 | Asia | Southeast Asia | Laos | unknown | unknown | Wong et al, 2015 |
| LNT1497 | ERR326671 | N/A | 3.4 | unknown | unknown | Blood | 2010 | Asia | Southeast Asia | Laos | unknown | unknown | Wong et al, 2015 |
| LNT1480 | ERR326672 | N/A | 2.2.3,2.2.2 | unknown | unknown | Blood | 2010 | Asia | Southeast Asia | Laos | unknown | unknown | Wong et al, 2015 |
| LNT1516 | ERR326673 | N/A | 3.4 | unknown | unknown | Blood | 2010 | Asia | Southeast Asia | Laos | unknown | unknown | Wong et al, 2015 |
| LNT1542 | ERR326674 | N/A | 3.4 | unknown | unknown | Blood | 2010 | Asia | Southeast Asia | Laos | unknown | unknown | Wong et al, 2015 |
| LNT266 | ERR326676 | N/A | 3.4 | unknown | unknown | Blood | 2008 | Asia | Southeast Asia | Laos | unknown | unknown | Wong et al, 2015 |
| LNT279 | ERR326677 | N/A | 3.4 | unknown | unknown | Blood | 2008 | Asia | Southeast Asia | Laos | unknown | unknown | Wong et al, 2015 |
| LNT330 | ERR326678 | N/A | 2.4 | unknown | unknown | Blood | 2008 | Asia | Southeast Asia | Laos | unknown | unknown | Wong et al, 2015 |
| LNT366 | ERR326679 | N/A | 3.4 | unknown | unknown | Blood | 2008 | Asia | Southeast Asia | Laos | unknown | unknown | Wong et al, 2015 |
| LNT375 | ERR326680 | N/A | 3.4 | unknown | unknown | Blood | 2008 | Asia | Southeast Asia | Laos | unknown | unknown | Wong et al, 2015 |

| | | | | | | | | | | | | | |
|---|---|---|---|---|---|---|---|---|---|---|---|---|---|
| LNT565 | ERR326681 | N/A | 3.4 | unknown | unknown | Blood | 2008 | Asia | Southeast Asia | Laos | unknown | unknown | Wong et al, 2015 |
| LNT609 | ERR326682 | N/A | 3.4 | unknown | unknown | Blood | 2008 | Asia | Southeast Asia | Laos | unknown | unknown | Wong et al, 2015 |
| LNT659 | ERR326683 | N/A | 3.4 | unknown | unknown | Blood | 2008 | Asia | Southeast Asia | Laos | unknown | unknown | Wong et al, 2015 |
| LNT662 | ERR326684 | N/A | 2.3.4 | unknown | unknown | Blood | 2008 | Asia | Southeast Asia | Laos | unknown | unknown | Wong et al, 2015 |
| LNT666 | ERR326685 | N/A | 3.4 | unknown | unknown | Blood | 2008 | Asia | Southeast Asia | Laos | unknown | unknown | Wong et al, 2015 |
| LNT670 | ERR326686 | N/A | 3.4 | unknown | unknown | Blood | 2008 | Asia | Southeast Asia | Laos | unknown | unknown | Wong et al, 2015 |
| LNT705 | ERR326687 | N/A | 4.1 | unknown | unknown | Blood | 2008 | Asia | Southeast Asia | Laos | unknown | unknown | Wong et al, 2015 |
| LNT72 | ERR326688 | N/A | 2.3.4 | unknown | unknown | Blood | 2007 | Asia | Southeast Asia | Laos | unknown | unknown | Wong et al, 2015 |
| LNT722 | ERR326689 | N/A | 3.4 | unknown | unknown | Blood | 2008 | Asia | Southeast Asia | Laos | unknown | unknown | Wong et al, 2015 |
| 2008-003680 | ERR331205 | N/A | 4.3.1.1 | unknown | unknown | Blood | 2012 | Asia | Southeast Asia | Cambodia | unknown | unknown | Wong et al, 2015 |
| 01-2011-002432 | ERR331206 | N/A | 4.3.1.1 | unknown | unknown | Blood | 2012 | Asia | Southeast Asia | Cambodia | unknown | unknown | Wong et al, 2015 |
| 2007-019648 | ERR331207 | N/A | 4.3.1.1 | unknown | unknown | Blood | 2012 | Asia | Southeast Asia | Cambodia | unknown | unknown | Wong et al, 2015 |
| 2012-010389 | ERR331208 | N/A | 4.3.1.1 | unknown | unknown | Blood | 2012 | Asia | Southeast Asia | Cambodia | unknown | unknown | Wong et al, 2015 |
| A143 | ERR331212 | N/A | 2.1.3 | unknown | unknown | Blood | 2008 | Asia | Southeast Asia | Indonesia | unknown | unknown | Wong et al, 2015 |
| A149 | ERR331214 | N/A | 2.1.1 | unknown | unknown | Blood | 2006 | Asia | Southeast Asia | Indonesia | unknown | unknown | Wong et al, 2015 |
| A150 | ERR331215 | N/A | 3 | unknown | unknown | Blood | 2006 | Asia | Southeast Asia | Indonesia | unknown | unknown | Wong et al, 2015 |
| A153 | ERR331216 | N/A | 2.1.8 | unknown | unknown | Blood | 2010 | Asia | Southeast Asia | Indonesia | unknown | unknown | Wong et al, 2015 |
| A154 | ERR331217 | N/A | 2.1.9 | unknown | unknown | Blood | 2009 | Asia | Southeast Asia | Indonesia | unknown | unknown | Wong et al, 2015 |
| A155 | ERR331218 | N/A | 2.1.1 | unknown | unknown | Blood | 2006 | Asia | Southeast Asia | Indonesia | unknown | unknown | Wong et al, 2015 |
| A156 | ERR331219 | N/A | 3 | unknown | unknown | Blood | 2007 | Asia | Southeast Asia | Indonesia | unknown | unknown | Wong et al, 2015 |
| A158 | ERR331220 | N/A | 3.1.2 | unknown | unknown | Blood | 2009 | Asia | Southeast Asia | Indonesia | unknown | unknown | Wong et al, 2015 |
| A160 | ERR331221 | N/A | 2.1.1 | unknown | unknown | Blood | 2006 | Asia | Southeast Asia | Indonesia | unknown | unknown | Wong et al, 2015 |
| A173 | ERR331224 | N/A | 2.1.9 | unknown | unknown | Blood | 2009 | Asia | Southeast Asia | Indonesia | unknown | unknown | Wong et al, 2015 |
| A174 | ERR331225 | N/A | 2.1.3 | unknown | unknown | Blood | 2011 | Asia | Southeast Asia | Indonesia | unknown | unknown | Wong et al, 2015 |
| A178 | ERR331226 | N/A | 3.1.2 | unknown | unknown | Blood | 2009 | Asia | Southeast Asia | Indonesia | unknown | unknown | Wong et al, 2015 |
| A180 | ERR331227 | N/A | 4.1 | unknown | unknown | Blood | 2009 | Asia | Southeast Asia | Indonesia | unknown | unknown | Wong et al, 2015 |
| A235 | ERR331229 | N/A | 3 | unknown | unknown | Blood | 2009 | Asia | Southeast Asia | Indonesia | unknown | unknown | Wong et al, 2015 |
| A236 | ERR331230 | N/A | 4.1 | unknown | unknown | Blood | 2009 | Asia | Southeast Asia | Indonesia | unknown | unknown | Wong et al, 2015 |
| A237 | ERR331231 | N/A | 3 | unknown | unknown | Blood | 2010 | Asia | Southeast Asia | Indonesia | unknown | unknown | Wong et al, 2015 |
| A238 | ERR331232 | N/A | 4.1 | unknown | unknown | Blood | 2009 | Asia | Southeast Asia | Indonesia | unknown | unknown | Wong et al, 2015 |
| A239 | ERR331233 | N/A | 3.1.2 | unknown | unknown | Blood | 2009 | Asia | Southeast Asia | Indonesia | unknown | unknown | Wong et al, 2015 |
| A240 | ERR331234 | N/A | 4.1 | unknown | unknown | Blood | 2010 | Asia | Southeast Asia | Indonesia | unknown | unknown | Wong et al, 2015 |
| A243 | ERR331236 | N/A | 4.1 | unknown | unknown | Blood | 2011 | Asia | Southeast Asia | Indonesia | unknown | unknown | Wong et al, 2015 |
| A244 | ERR331237 | N/A | 3 | unknown | unknown | Blood | 2011 | Asia | Southeast Asia | Indonesia | unknown | unknown | Wong et al, 2015 |
| A245 | ERR331238 | N/A | 4.1 | unknown | unknown | Blood | 2009 | Asia | Southeast Asia | Indonesia | unknown | unknown | Wong et al, 2015 |
| A246 | ERR331239 | N/A | 4.1 | unknown | unknown | Blood | 2010 | Asia | Southeast Asia | Indonesia | unknown | unknown | Wong et al, 2015 |
| A247 | ERR331240 | N/A | 4.1 | unknown | unknown | Blood | 2010 | Asia | Southeast Asia | Indonesia | unknown | unknown | Wong et al, 2015 |
| A248 | ERR331241 | N/A | 3 | unknown | unknown | Blood | 2009 | Asia | Southeast Asia | Indonesia | unknown | unknown | Wong et al, 2015 |
| A249 | ERR331242 | N/A | 3.1.2 | unknown | unknown | Blood | 2010 | Asia | Southeast Asia | Indonesia | unknown | unknown | Wong et al, 2015 |
| A250 | ERR331243 | N/A | 4.1 | unknown | unknown | Blood | 2009 | Asia | Southeast Asia | Indonesia | unknown | unknown | Wong et al, 2015 |
| A252 | ERR331244 | N/A | 4.1 | unknown | unknown | Blood | 2011 | Asia | Southeast Asia | Indonesia | unknown | unknown | Wong et al, 2015 |
| A253 | ERR331245 | N/A | 2.1.6 | unknown | unknown | Blood | 2009 | Asia | Southeast Asia | Indonesia | unknown | unknown | Wong et al, 2015 |
| A255 | ERR331247 | N/A | 4.1 | unknown | unknown | Blood | 2011 | Asia | Southeast Asia | Indonesia | unknown | unknown | Wong et al, 2015 |
| A256 | ERR331248 | N/A | 3 | unknown | unknown | Blood | 2010 | Asia | Southeast Asia | Indonesia | unknown | unknown | Wong et al, 2015 |
| A257 | ERR331249 | N/A | 3 | unknown | unknown | Blood | 2010 | Asia | Southeast Asia | Indonesia | unknown | unknown | Wong et al, 2015 |
| A258 | ERR331250 | N/A | 4.1 | unknown | unknown | Blood | 2009 | Asia | Southeast Asia | Indonesia | unknown | unknown | Wong et al, 2015 |
| A259 | ERR331251 | N/A | 4.1 | unknown | unknown | Blood | 2010 | Asia | Southeast Asia | Indonesia | unknown | unknown | Wong et al, 2015 |
| A261 | ERR331252 | N/A | 2.1.6 | unknown | unknown | Blood | 2009 | Asia | Southeast Asia | Indonesia | unknown | unknown | Wong et al, 2015 |
| A262 | ERR331253 | N/A | 2.1.6 | unknown | unknown | Blood | 2010 | Asia | Southeast Asia | Indonesia | unknown | unknown | Wong et al, 2015 |
| A263 | ERR331254 | N/A | 4.1 | unknown | unknown | Blood | 2009 | Asia | Southeast Asia | Indonesia | unknown | unknown | Wong et al, 2015 |

| | | | | | | | | | | | | | |
|---|---|---|---|---|---|---|---|---|---|---|---|---|---|
| A265 | ERR331256 | N/A | 4.1 | unknown | unknown | Blood | 2010 | Asia | Southeast Asia | Indonesia | unknown | unknown | Wong et al, 2015 |
| A266 | ERR331257 | N/A | 4.1 | unknown | unknown | Blood | 2010 | Asia | Southeast Asia | Indonesia | unknown | unknown | Wong et al, 2015 |
| A267 | ERR331258 | N/A | 2.1.6 | unknown | unknown | Blood | 2008 | Asia | Southeast Asia | Indonesia | unknown | unknown | Wong et al, 2015 |
| 100949 | ERR331260 | N/A | 4.2.2 | unknown | unknown | Unknown | 2011 | Australia & Oceania | Oceania | Fiji | unknown | unknown | Wong et al, 2015 |
| 102196 | ERR331261 | N/A | 4.2.2 | unknown | unknown | Unknown | 2011 | Australia & Oceania | Oceania | Fiji | unknown | unknown | Wong et al, 2015 |
| 103643 | ERR331262 | N/A | 4.2.1 | unknown | unknown | Unknown | 2011 | Australia & Oceania | Oceania | Fiji | unknown | unknown | Wong et al, 2015 |
| 106774 | ERR331263 | N/A | 4.2.2 | unknown | unknown | Unknown | 2012 | Australia & Oceania | Oceania | Fiji | unknown | unknown | Wong et al, 2015 |
| 107123 | ERR331264 | N/A | 4.2.1 | unknown | unknown | Unknown | 2012 | Australia & Oceania | Oceania | Fiji | unknown | unknown | Wong et al, 2015 |
| 107364 | ERR331265 | N/A | 4.2.2 | unknown | unknown | Unknown | 2012 | Australia & Oceania | Oceania | Fiji | unknown | unknown | Wong et al, 2015 |
| 108891 | ERR331266 | N/A | 4.2.2 | unknown | unknown | Unknown | 2012 | Australia & Oceania | Oceania | Fiji | unknown | unknown | Wong et al, 2015 |
| 108910 | ERR331267 | N/A | 4.2.2 | unknown | unknown | Unknown | 2012 | Australia & Oceania | Oceania | Fiji | unknown | unknown | Wong et al, 2015 |
| 109520 | ERR331268 | N/A | 4.2.2 | unknown | unknown | Unknown | 2012 | Australia & Oceania | Oceania | Fiji | unknown | unknown | Wong et al, 2015 |
| 110829 | ERR331269 | N/A | 4.2.2 | unknown | unknown | Unknown | 2012 | Australia & Oceania | Oceania | Fiji | unknown | unknown | Wong et al, 2015 |
| B1271 | ERR331270 | N/A | 4.2.1 | unknown | unknown | Blood | 2012 | Australia & Oceania | Oceania | Fiji | unknown | unknown | Wong et al, 2015 |
| B1305 | ERR331271 | N/A | 4.2.2 | unknown | unknown | Blood | 2012 | Australia & Oceania | Oceania | Fiji | unknown | unknown | Wong et al, 2015 |
| B1357 | ERR331273 | N/A | 4.2.2 | unknown | unknown | Blood | 2012 | Australia & Oceania | Oceania | Fiji | unknown | unknown | Wong et al, 2015 |
| B1472 | ERR331274 | N/A | 4.2.2 | unknown | unknown | Blood | 2012 | Australia & Oceania | Oceania | Fiji | unknown | unknown | Wong et al, 2015 |
| B1501 | ERR331275 | N/A | 4.2.2 | unknown | unknown | Blood | 2012 | Australia & Oceania | Oceania | Fiji | unknown | unknown | Wong et al, 2015 |
| B1502 | ERR331276 | N/A | 4.2.2 | unknown | unknown | Blood | 2012 | Australia & Oceania | Oceania | Fiji | unknown | unknown | Wong et al, 2015 |
| B1542 | ERR331278 | N/A | 4.2.2 | unknown | unknown | Blood | 2012 | Australia & Oceania | Oceania | Fiji | unknown | unknown | Wong et al, 2015 |
| B1563 | ERR331279 | N/A | 4.2.2 | unknown | unknown | Blood | 2012 | Australia & Oceania | Oceania | Fiji | unknown | unknown | Wong et al, 2015 |
| B1596 | ERR331280 | N/A | 4.2.2 | unknown | unknown | Blood | 2012 | Australia & Oceania | Oceania | Fiji | unknown | unknown | Wong et al, 2015 |
| B1616 | ERR331281 | N/A | 4.2.2 | unknown | unknown | Blood | 2012 | Australia & Oceania | Oceania | Fiji | unknown | unknown | Wong et al, 2015 |
| B1628 | ERR331282 | N/A | 4.2.2 | unknown | unknown | Blood | 2012 | Australia & Oceania | Oceania | Fiji | unknown | unknown | Wong et al, 2015 |
| B2667 | ERR331283 | N/A | 4.2.2 | unknown | unknown | Unknown | 2008 | Australia & Oceania | Oceania | Fiji | unknown | unknown | Wong et al, 2015 |
| B4381 | ERR331284 | N/A | 4.2.2 | unknown | unknown | Unknown | 2008 | Australia & Oceania | Oceania | Fiji | unknown | unknown | Wong et al, 2015 |
| B4387 | ERR331285 | N/A | 4.2.2 | unknown | unknown | Unknown | 2008 | Australia & Oceania | Oceania | Fiji | unknown | unknown | Wong et al, 2015 |
| P749 | ERR331287 | N/A | 4.2.2 | unknown | unknown | Unknown | 2008 | Australia & Oceania | Oceania | Fiji | unknown | unknown | Wong et al, 2015 |
| W478 | ERR331288 | N/A | 4.2.2 | unknown | unknown | Rectal swab | 2012 | Australia & Oceania | Oceania | Fiji | unknown | unknown | Wong et al, 2015 |
| W794 | ERR331289 | N/A | 4.2.2 | unknown | unknown | Stool | 2012 | Australia & Oceania | Oceania | Fiji | unknown | unknown | Wong et al, 2015 |
| 1102 | ERR331290 | N/A | 4.3.1.1 | unknown | unknown | Blood | 2011 | Asia | South Asia | Bangladesh | unknown | unknown | Wong et al, 2015 |
| 1154 | ERR331291 | N/A | 4.3.1.3 | unknown | unknown | Blood | 2011 | Asia | South Asia | Bangladesh | unknown | unknown | Wong et al, 2015 |
| 1404 | ERR331292 | N/A | 4.3.1.1 | unknown | unknown | Blood | 2011 | Asia | South Asia | Bangladesh | unknown | unknown | Wong et al, 2015 |
| 1499 | ERR331293 | N/A | 4.3.1.1 | unknown | unknown | Blood | 2011 | Asia | South Asia | Bangladesh | unknown | unknown | Wong et al, 2015 |
| 1553 | ERR331294 | N/A | 4.3.1.3.Bdq | unknown | unknown | Blood | 2011 | Asia | South Asia | Bangladesh | unknown | unknown | Wong et al, 2015 |
| 1607 | ERR331295 | N/A | 4.3.1.3.Bdq | unknown | unknown | Blood | 2011 | Asia | South Asia | Bangladesh | unknown | unknown | Wong et al, 2015 |
| 1703 | ERR331296 | N/A | 4.3.1.1 | unknown | unknown | Blood | 2008 | Asia | South Asia | Bangladesh | unknown | unknown | Wong et al, 2015 |
| 1707 | ERR331297 | N/A | 4.3.1.1 | unknown | unknown | Blood | 2012 | Asia | South Asia | Bangladesh | unknown | unknown | Wong et al, 2015 |
| LNT757 | ERR331298 | N/A | 2.3.4 | unknown | unknown | Blood | 2009 | Asia | Southeast Asia | Laos | unknown | unknown | Wong et al, 2015 |
| LNT899 | ERR331300 | N/A | 3.4 | unknown | unknown | Blood | 2009 | Asia | Southeast Asia | Laos | unknown | unknown | Wong et al, 2015 |
| LPB866 | ERR331301 | N/A | 4.3.1.1 | unknown | unknown | Blood | 2008 | Asia | Southeast Asia | Laos | unknown | unknown | Wong et al, 2015 |
| ST716 | ERR331302 | N/A | 3.4 | unknown | unknown | Blood | 2008 | Asia | Southeast Asia | Laos | unknown | unknown | Wong et al, 2015 |
| SV108 | ERR331303 | N/A | 3.2.1 | unknown | unknown | Blood | 2008 | Asia | Southeast Asia | Laos | unknown | unknown | Wong et al, 2015 |
| SV170 | ERR331304 | N/A | 3.2.1 | unknown | unknown | Blood | 2009 | Asia | Southeast Asia | Laos | unknown | unknown | Wong et al, 2015 |
| SV171 | ERR331305 | N/A | 3.2.1 | unknown | unknown | Blood | 2009 | Asia | Southeast Asia | Laos | unknown | unknown | Wong et al, 2015 |
| SV179 | ERR331306 | N/A | 3.2.1 | unknown | unknown | Blood | 2009 | Asia | Southeast Asia | Laos | unknown | unknown | Wong et al, 2015 |
| SV200 | ERR331307 | N/A | 3.4 | unknown | unknown | Blood | 2009 | Asia | Southeast Asia | Laos | unknown | unknown | Wong et al, 2015 |
| SV211 | ERR331308 | N/A | 3.2.1 | unknown | unknown | Blood | 2009 | Asia | Southeast Asia | Laos | unknown | unknown | Wong et al, 2015 |
| SV357 | ERR331309 | N/A | 3.5.2 | unknown | unknown | Blood | 2009 | Asia | Southeast Asia | Laos | unknown | unknown | Wong et al, 2015 |
| SV430 | ERR331310 | N/A | 2.2.3,2.2.2 | unknown | unknown | Blood | 2009 | Asia | Southeast Asia | Laos | unknown | unknown | Wong et al, 2015 |

| | | | | | | | | | | | | | |
|---|---|---|---|---|---|---|---|---|---|---|---|---|---|
| SV431 | ERR331311 | N/A | 3.2.1 | unknown | unknown | Blood | 2009 | Asia | Southeast Asia | Laos | unknown | unknown | Wong et al, 2015 |
| SV500 | ERR331312 | N/A | 3.5.2 | unknown | unknown | Blood | 2010 | Asia | Southeast Asia | Laos | unknown | unknown | Wong et al, 2015 |
| SV547 | ERR331313 | N/A | 3.5.2 | unknown | unknown | Blood | 2010 | Asia | Southeast Asia | Laos | unknown | unknown | Wong et al, 2015 |
| SV552 | ERR331314 | N/A | 3.5.2 | unknown | unknown | Blood | 2010 | Asia | Southeast Asia | Laos | unknown | unknown | Wong et al, 2015 |
| SV610 | ERR331315 | N/A | 3.5.2 | unknown | unknown | Blood | 2010 | Asia | Southeast Asia | Laos | unknown | unknown | Wong et al, 2015 |
| UI10006 | ERR331316 | N/A | 2.3.4 | unknown | unknown | Blood | 2007 | Asia | Southeast Asia | Laos | unknown | unknown | Wong et al, 2015 |
| UI10788 | ERR331317 | N/A | 2.2.3,2.2.2 | unknown | unknown | Blood | 2007 | Asia | Southeast Asia | Laos | unknown | unknown | Wong et al, 2015 |
| UI11483 | ERR331318 | N/A | 3.4 | unknown | unknown | Blood | 2008 | Asia | Southeast Asia | Laos | unknown | unknown | Wong et al, 2015 |
| UI11562 | ERR331319 | N/A | 4.1 | unknown | unknown | Blood | 2008 | Asia | Southeast Asia | Laos | unknown | unknown | Wong et al, 2015 |
| UI11955 | ERR331320 | N/A | 3.4 | unknown | unknown | Blood | 2008 | Asia | Southeast Asia | Laos | unknown | unknown | Wong et al, 2015 |
| UI12162 | ERR331321 | N/A | 3.2.1 | unknown | unknown | Blood | 2008 | Asia | Southeast Asia | Laos | unknown | unknown | Wong et al, 2015 |
| UI13529-2 | ERR331322 | N/A | 3.4 | unknown | unknown | Blood | 2009 | Asia | Southeast Asia | Laos | unknown | unknown | Wong et al, 2015 |
| UI13599 | ERR331323 | N/A | 3.2.1 | unknown | unknown | Blood | 2009 | Asia | Southeast Asia | Laos | unknown | unknown | Wong et al, 2015 |
| UI13797 | ERR331324 | N/A | 3.2.1 | unknown | unknown | Blood | 2009 | Asia | Southeast Asia | Laos | unknown | unknown | Wong et al, 2015 |
| UI13823 | ERR331325 | N/A | 4.3.1.1 | unknown | unknown | Blood | 2009 | Asia | Southeast Asia | Laos | unknown | unknown | Wong et al, 2015 |
| UI14191/3 | ERR331326 | N/A | 3.4 | unknown | unknown | Blood | 2009 | Asia | Southeast Asia | Laos | unknown | unknown | Wong et al, 2015 |
| UI14598 | ERR331327 | N/A | 2.2.3,2.2.2 | unknown | unknown | Blood | 2009 | Asia | Southeast Asia | Laos | unknown | unknown | Wong et al, 2015 |
| UI15075 | ERR331328 | N/A | 4.3.1.1 | unknown | unknown | Blood | 2008 | Asia | Southeast Asia | Laos | unknown | unknown | Wong et al, 2015 |
| UI16161 | ERR331329 | N/A | 3.4 | unknown | unknown | Blood | 2010 | Asia | Southeast Asia | Laos | unknown | unknown | Wong et al, 2015 |
| UI16704 | ERR331330 | N/A | 4.3.1.1 | unknown | unknown | Blood | 2010 | Asia | Southeast Asia | Laos | unknown | unknown | Wong et al, 2015 |
| UI17187 | ERR331331 | N/A | 4.1 | unknown | unknown | Blood | 2010 | Asia | Southeast Asia | Laos | unknown | unknown | Wong et al, 2015 |
| UI17614 | ERR331332 | N/A | 3.2.1 | unknown | unknown | Blood | 2010 | Asia | Southeast Asia | Laos | unknown | unknown | Wong et al, 2015 |
| UI3398 | ERR331333 | N/A | 4.3.1.1 | unknown | unknown | Blood | 2003 | Asia | Southeast Asia | Laos | unknown | unknown | Wong et al, 2015 |
| UI3446 | ERR331334 | N/A | 3 | unknown | unknown | Blood | 2003 | Asia | Southeast Asia | Laos | unknown | unknown | Wong et al, 2015 |
| UI3452 | ERR331335 | N/A | 4.3.1.1 | unknown | unknown | Blood | 2003 | Asia | Southeast Asia | Laos | unknown | unknown | Wong et al, 2015 |
| UI3492 | ERR331336 | N/A | 4.3.1.1 | unknown | unknown | Blood | 2003 | Asia | Southeast Asia | Laos | unknown | unknown | Wong et al, 2015 |
| UI3564 | ERR331337 | N/A | 4.1 | unknown | unknown | Blood | 2003 | Asia | Southeast Asia | Laos | unknown | unknown | Wong et al, 2015 |
| UI3608 | ERR331339 | N/A | 2.2 | unknown | unknown | Blood | 2003 | Asia | Southeast Asia | Laos | unknown | unknown | Wong et al, 2015 |
| UI3744 | ERR331340 | N/A | 3.4 | unknown | unknown | Blood | 2003 | Asia | Southeast Asia | Laos | unknown | unknown | Wong et al, 2015 |
| UI3753 | ERR331341 | N/A | 3.2.1 | unknown | unknown | Blood | 2003 | Asia | Southeast Asia | Laos | unknown | unknown | Wong et al, 2015 |
| UI3816 | ERR331342 | N/A | 3.4 | unknown | unknown | Blood | 2003 | Asia | Southeast Asia | Laos | unknown | unknown | Wong et al, 2015 |
| UI3862 | ERR331343 | N/A | 3.4 | unknown | unknown | Blood | 2003 | Asia | Southeast Asia | Laos | unknown | unknown | Wong et al, 2015 |
| UI3915 | ERR331344 | N/A | 4.3.1.1 | unknown | unknown | Blood | 2003 | Asia | Southeast Asia | Laos | unknown | unknown | Wong et al, 2015 |
| UI3930 | ERR331345 | N/A | 3.4 | unknown | unknown | Blood | 2003 | Asia | Southeast Asia | Laos | unknown | unknown | Wong et al, 2015 |
| UI4389 | ERR331346 | N/A | 4.3.1.1 | unknown | unknown | Blood | 2003 | Asia | Southeast Asia | Laos | unknown | unknown | Wong et al, 2015 |
| UI4639 | ERR331347 | N/A | 4.3.1.1 | unknown | unknown | Blood | 2003 | Asia | Southeast Asia | Laos | unknown | unknown | Wong et al, 2015 |
| UI4692 | ERR331348 | N/A | 2.2.3,2.2.2 | unknown | unknown | Blood | 2004 | Asia | Southeast Asia | Laos | unknown | unknown | Wong et al, 2015 |
| UI4940 | ERR331349 | N/A | 4.1 | unknown | unknown | Blood | 2004 | Asia | Southeast Asia | Laos | unknown | unknown | Wong et al, 2015 |
| UI5026 | ERR331350 | N/A | 3.4 | unknown | unknown | Blood | 2004 | Asia | Southeast Asia | Laos | unknown | unknown | Wong et al, 2015 |
| UI5106 | ERR331351 | N/A | 4.3.1.1 | unknown | unknown | Blood | 2004 | Asia | Southeast Asia | Laos | unknown | unknown | Wong et al, 2015 |
| UI5154 | ERR331352 | N/A | 4.3.1.1 | unknown | unknown | Blood | 2004 | Asia | Southeast Asia | Laos | unknown | unknown | Wong et al, 2015 |
| UI5275 | ERR331353 | N/A | 3.2.1 | unknown | unknown | Blood | 2004 | Asia | Southeast Asia | Laos | unknown | unknown | Wong et al, 2015 |
| UI5312 | ERR331354 | N/A | 3.2.1 | unknown | unknown | Blood | 2004 | Asia | Southeast Asia | Laos | unknown | unknown | Wong et al, 2015 |
| UI5446 | ERR331355 | N/A | 3.4 | unknown | unknown | Blood | 2004 | Asia | Southeast Asia | Laos | unknown | unknown | Wong et al, 2015 |
| UI5557 | ERR331356 | N/A | 3.4 | unknown | unknown | Blood | 2004 | Asia | Southeast Asia | Laos | unknown | unknown | Wong et al, 2015 |
| UI5669 | ERR331357 | N/A | 4.1 | unknown | unknown | Blood | 2004 | Asia | Southeast Asia | Laos | unknown | unknown | Wong et al, 2015 |
| UI5685-2 | ERR331358 | N/A | 4.1 | unknown | unknown | Blood | 2004 | Asia | Southeast Asia | Laos | unknown | unknown | Wong et al, 2015 |
| UX229 | ERR331359 | N/A | 3.4 | unknown | unknown | Blood | 2009 | Asia | Southeast Asia | Laos | unknown | unknown | Wong et al, 2015 |
| UX305 | ERR331360 | N/A | 3.4 | unknown | unknown | Blood | 2009 | Asia | Southeast Asia | Laos | unknown | unknown | Wong et al, 2015 |
| UX309 | ERR331361 | N/A | 3.4 | unknown | unknown | Blood | 2009 | Asia | Southeast Asia | Laos | unknown | unknown | Wong et al, 2015 |

| | | | | | | | | | | | | | |
|---|---|---|---|---|---|---|---|---|---|---|---|---|---|
| XK1682 | ERR331363 | N/A | 3.2.1 | unknown | unknown | Blood | 2010 | Asia | Southeast Asia | Laos | unknown | unknown | Wong et al, 2015 |
| XK1684 | ERR331364 | N/A | 3.2.1 | unknown | unknown | Blood | 2010 | Asia | Southeast Asia | Laos | unknown | unknown | Wong et al, 2015 |
| XN211 | ERR331365 | N/A | 3.4 | unknown | unknown | Blood | 2010 | Asia | Southeast Asia | Laos | unknown | unknown | Wong et al, 2015 |
| 60434 | ERR331366 | N/A | 2.2.2 | unknown | unknown | Blood | 2006 | Africa | East Africa | Tanzania | unknown | unknown | Wong et al, 2015 |
| 129-0242-M | ERR331368 | N/A | 4.3.1.1.EA1 | unknown | unknown | Blood | 2008 | Africa | East Africa | Tanzania | unknown | unknown | Wong et al, 2015 |
| 129-0289-M | ERR331369 | N/A | 4.3.1.1.EA1 | unknown | unknown | Blood | 2008 | Africa | East Africa | Tanzania | unknown | unknown | Wong et al, 2015 |
| 61025 | ERR331370 | N/A | 2.2 | unknown | unknown | Blood | 2006 | Africa | East Africa | Tanzania | unknown | unknown | Wong et al, 2015 |
| 63714 | ERR331371 | N/A | 3.3.1 | unknown | unknown | Blood | 2007 | Africa | East Africa | Tanzania | unknown | unknown | Wong et al, 2015 |
| 129-0327-M | ERR331372 | N/A | 4.3.1.1.EA1 | unknown | unknown | Blood | 2008 | Africa | East Africa | Tanzania | unknown | unknown | Wong et al, 2015 |
| 129-0177-M | ERR331373 | N/A | 3.1 | unknown | unknown | Blood | 2007 | Africa | East Africa | Tanzania | unknown | unknown | Wong et al, 2015 |
| 129-0268-M | ERR331374 | N/A | 4.3.1.1.EA1 | unknown | unknown | Blood | 2008 | Africa | East Africa | Tanzania | unknown | unknown | Wong et al, 2015 |
| 129-0339-M | ERR331375 | N/A | 4.3.1.1.EA1 | unknown | unknown | Blood | 2008 | Africa | East Africa | Tanzania | unknown | unknown | Wong et al, 2015 |
| 129-0303-M | ERR331376 | N/A | 4.3.1.1.EA1 | unknown | unknown | Blood | 2008 | Africa | East Africa | Tanzania | unknown | unknown | Wong et al, 2015 |
| 129-0257-M | ERR331377 | N/A | 4.3.1.1.EA1 | unknown | unknown | Blood | 2008 | Africa | East Africa | Tanzania | unknown | unknown | Wong et al, 2015 |
| 129-0230-K | ERR331378 | N/A | 4.3.1.1.EA1 | unknown | unknown | Blood | 2008 | Africa | East Africa | Tanzania | unknown | unknown | Wong et al, 2015 |
| 129-0254-M | ERR331379 | N/A | 4.3.1.1.EA1 | unknown | unknown | Blood | 2008 | Africa | East Africa | Tanzania | unknown | unknown | Wong et al, 2015 |
| 129-0238-M | ERR331380 | N/A | 4.3.1.1.EA1 | unknown | unknown | Blood | 2008 | Africa | East Africa | Tanzania | unknown | unknown | Wong et al, 2015 |
| 62717 | ERR331381 | N/A | 2.2 | unknown | unknown | Blood | 2006 | Africa | East Africa | Tanzania | unknown | unknown | Wong et al, 2015 |
| H062640481 | ERR331382 | N/A | 4.3.1.1 | unknown | unknown | Blood | 2006 | Asia | Western Asia | Iraq | unknown | unknown | Wong et al, 2015 |
| H094620494 | ERR331383 | N/A | 4.3.1.1 | unknown | unknown | Blood | 2009 | Asia | Western Asia | Iraq | unknown | unknown | Wong et al, 2015 |
| H083520583 | ERR331384 | N/A | 4.3.1.1 | unknown | unknown | Blood | 2008 | Asia | Western Asia | Iraq | unknown | unknown | Wong et al, 2015 |
| H104280445 | ERR331385 | N/A | 4.3.1.1 | unknown | unknown | Blood | 2010 | Asia | Western Asia | Iraq | unknown | unknown | Wong et al, 2015 |
| H075080543 | ERR331386 | N/A | 4.3.1.1 | unknown | unknown | Blood | 2007 | Asia | Western Asia | Iraq | unknown | unknown | Wong et al, 2015 |
| H094420550 | ERR331387 | N/A | 4.3.1.1 | unknown | unknown | Stool | 2009 | Asia | Western Asia | Iraq | unknown | unknown | Wong et al, 2015 |
| H084140247 | ERR331388 | N/A | 4.3.1.1 | unknown | unknown | Blood | 2008 | Asia | Western Asia | Iraq | unknown | unknown | Wong et al, 2015 |
| H094320403 | ERR331389 | N/A | 4.3.1.1 | unknown | unknown | Blood | 2009 | Asia | Western Asia | Iraq | unknown | unknown | Wong et al, 2015 |
| H084260341 | ERR331390 | N/A | 4.3.1.1 | unknown | unknown | Blood | 2008 | Asia | Western Asia | Iraq | unknown | unknown | Wong et al, 2015 |
| 1738 | ERR337977 | N/A | 4.3.1.3.Bdq | unknown | unknown | Blood | 2012 | Asia | South Asia | Bangladesh | unknown | unknown | Wong et al, 2015 |
| 1877 | ERR337978 | N/A | 4.3.1.3 | unknown | unknown | Blood | 2012 | Asia | South Asia | Bangladesh | unknown | unknown | Wong et al, 2015 |
| 2181 | ERR337979 | N/A | 3.3.2 | unknown | unknown | Blood | 2012 | Asia | South Asia | Bangladesh | unknown | unknown | Wong et al, 2015 |
| TY019 | ERR337981 | N/A | 3.3.2 | unknown | unknown | Blood | 2012 | Asia | South Asia | Bangladesh | unknown | unknown | Wong et al, 2015 |
| TY029 | ERR337982 | N/A | 3.3.2 | unknown | unknown | Blood | 2012 | Asia | South Asia | Bangladesh | unknown | unknown | Wong et al, 2015 |
| TY032 | ERR337983 | N/A | 3.3.2 | unknown | unknown | Blood | 2012 | Asia | South Asia | Bangladesh | unknown | unknown | Wong et al, 2015 |
| TY053 | ERR337984 | N/A | 4.3.1.1 | unknown | unknown | Blood | 2012 | Asia | South Asia | Bangladesh | unknown | unknown | Wong et al, 2015 |
| TY063 | ERR337985 | N/A | 3.3.2 | unknown | unknown | Blood | 2012 | Asia | South Asia | Bangladesh | unknown | unknown | Wong et al, 2015 |
| TY068 | ERR337986 | N/A | 3.2.2 | unknown | unknown | Blood | 2012 | Asia | South Asia | Bangladesh | unknown | unknown | Wong et al, 2015 |
| TY107 | ERR337987 | N/A | 4.3.1.1 | unknown | unknown | Blood | 2012 | Asia | South Asia | Bangladesh | unknown | unknown | Wong et al, 2015 |
| TY108 | ERR337988 | N/A | 3.3.2 | unknown | unknown | Blood | 2012 | Asia | South Asia | Bangladesh | unknown | unknown | Wong et al, 2015 |
| TY121 | ERR337989 | N/A | 4.3.1.1 | unknown | unknown | Blood | 2012 | Asia | South Asia | Bangladesh | unknown | unknown | Wong et al, 2015 |
| TY132 | ERR337990 | N/A | 4.3.1.1 | unknown | unknown | Blood | 2012 | Asia | South Asia | Bangladesh | unknown | unknown | Wong et al, 2015 |
| TY138 | ERR337991 | N/A | 3.3.2 | unknown | unknown | Blood | 2012 | Asia | South Asia | Bangladesh | unknown | unknown | Wong et al, 2015 |
| TY524 | ERR337992 | N/A | 3.3.2 | unknown | unknown | Blood | 2012 | Asia | South Asia | Bangladesh | unknown | unknown | Wong et al, 2015 |
| TY529 | ERR337993 | N/A | 4.3.1.3 | unknown | unknown | Blood | 2012 | Asia | South Asia | Bangladesh | unknown | unknown | Wong et al, 2015 |
| TY585 | ERR337995 | N/A | 4.3.1.1 | unknown | unknown | Blood | 2012 | Asia | South Asia | Bangladesh | unknown | unknown | Wong et al, 2015 |
| TY589 | ERR337996 | N/A | 4.3.1.2 | unknown | unknown | Blood | 2012 | Asia | South Asia | Bangladesh | unknown | unknown | Wong et al, 2015 |
| TY647 | ERR337997 | N/A | 4.3.1.3 | unknown | unknown | Blood | 2012 | Asia | South Asia | Bangladesh | unknown | unknown | Wong et al, 2015 |
| 231697 | ERR337998 | N/A | 2.4.1 | unknown | unknown | Blood | 2007 | Africa | South Africa | South Africa | unknown | unknown | Wong et al, 2015 |
| 155749 | ERR337999 | N/A | 2.5 | unknown | unknown | Blood | 2006 | Africa | South Africa | South Africa | unknown | unknown | Wong et al, 2015 |
| 175806 | ERR338001 | N/A | 2.4 | unknown | unknown | Blood | 2006 | Africa | South Africa | South Africa | unknown | unknown | Wong et al, 2015 |
| 1647901 | ERR338002 | N/A | 4.3.1.1.EA1 | unknown | unknown | Blood | 2005 | Africa | South Africa | South Africa | unknown | unknown | Wong et al, 2015 |

| | | | | | | | | | | | | | |
|---|---|---|---|---|---|---|---|---|---|---|---|---|---|
| 1025041 | ERR338003 | N/A | 2.4.1 | unknown | unknown | Blood | 2004 | Africa | South Africa | South Africa | unknown | unknown | Wong et al, 2015 |
| 1025617 | ERR338004 | N/A | 4.1.1 | unknown | unknown | Blood | 2004 | Africa | South Africa | South Africa | unknown | unknown | Wong et al, 2015 |
| 1647624 | ERR338005 | N/A | 2.4 | unknown | unknown | Blood | 2005 | Africa | South Africa | South Africa | unknown | unknown | Wong et al, 2015 |
| 237091 | ERR338006 | N/A | 2.4.1 | unknown | unknown | Blood | 2007 | Africa | South Africa | South Africa | unknown | unknown | Wong et al, 2015 |
| 226195 | ERR338007 | N/A | 2.4.1 | unknown | unknown | Blood | 2007 | Africa | South Africa | South Africa | unknown | unknown | Wong et al, 2015 |
| 206926 | ERR338008 | N/A | 1.1.2 | unknown | unknown | Blood | 2007 | Africa | South Africa | South Africa | unknown | unknown | Wong et al, 2015 |
| 225555 | ERR338009 | N/A | 2.4 | unknown | unknown | Blood | 2007 | Africa | South Africa | South Africa | unknown | unknown | Wong et al, 2015 |
| 241451 | ERR338010 | N/A | 2.4.1 | unknown | unknown | Blood | 2008 | Africa | South Africa | South Africa | unknown | unknown | Wong et al, 2015 |
| 238675 | ERR338011 | N/A | 2.4 | unknown | unknown | Blood | 2008 | Africa | South Africa | South Africa | unknown | unknown | Wong et al, 2015 |
| 241454 | ERR338012 | N/A | 2.4.1 | unknown | unknown | Blood | 2008 | Africa | South Africa | South Africa | unknown | unknown | Wong et al, 2015 |
| 257401 | ERR338013 | N/A | 2.4 | unknown | unknown | Blood | 2008 | Africa | South Africa | South Africa | unknown | unknown | Wong et al, 2015 |
| 141461 | ERR338016 | N/A | 2.4.1 | unknown | unknown | Blood | 2006 | Africa | South Africa | South Africa | unknown | unknown | Wong et al, 2015 |
| 1650362 | ERR338019 | N/A | 2.4 | unknown | unknown | Blood | 2005 | Africa | South Africa | South Africa | unknown | unknown | Wong et al, 2015 |
| 1026656 | ERR338020 | N/A | 2.4 | unknown | unknown | Blood | 2004 | Africa | South Africa | South Africa | unknown | unknown | Wong et al, 2015 |
| 493790 | ERR338023 | N/A | 4.3.1.1.EA1 | unknown | unknown | Blood | 2010 | Africa | South Africa | South Africa | unknown | unknown | Wong et al, 2015 |
| 520772 | ERR338024 | N/A | 2.4.1 | unknown | unknown | Breast pus | 2010 | Africa | South Africa | South Africa | unknown | unknown | Wong et al, 2015 |
| 516981 | ERR338025 | N/A | 2.4 | unknown | unknown | Blood | 2010 | Africa | South Africa | South Africa | unknown | unknown | Wong et al, 2015 |
| 474899 | ERR338026 | N/A | 4.3.1.3.Bdq | unknown | unknown | Blood | 2010 | Africa | South Africa | South Africa | unknown | unknown | Wong et al, 2015 |
| 427489 | ERR338027 | N/A | 3.3.1 | unknown | unknown | Blood & bone marrow | 2010 | Africa | South Africa | South Africa | unknown | unknown | Wong et al, 2015 |
| 412921 | ERR338028 | N/A | 2.5 | unknown | unknown | Blood | 2009 | Africa | South Africa | South Africa | unknown | unknown | Wong et al, 2015 |
| 410378 | ERR338029 | N/A | 2.4 | unknown | unknown | Blood | 2009 | Africa | South Africa | South Africa | unknown | unknown | Wong et al, 2015 |
| 410323 | ERR338030 | N/A | 3.3.1 | unknown | unknown | Blood | 2009 | Africa | South Africa | South Africa | unknown | unknown | Wong et al, 2015 |
| 542825 | ERR338031 | N/A | 3.3.1 | unknown | unknown | Blood | 2011 | Africa | South Africa | South Africa | unknown | unknown | Wong et al, 2015 |
| 540614 | ERR338032 | N/A | 2.5 | unknown | unknown | Blood | 2011 | Africa | South Africa | South Africa | unknown | unknown | Wong et al, 2015 |
| 583236 | ERR338034 | N/A | 4.3.1.1.EA1 | unknown | unknown | Blood | 2011 | Africa | South Africa | South Africa | unknown | unknown | Wong et al, 2015 |
| 627334 | ERR338035 | N/A | 1.1.2 | unknown | unknown | Blood | 2012 | Africa | South Africa | South Africa | unknown | unknown | Wong et al, 2015 |
| 648212 | ERR338036 | N/A | 4.3.1.1.EA1 | unknown | unknown | Blood | 2012 | Africa | South Africa | South Africa | unknown | unknown | Wong et al, 2015 |
| 639624 | ERR338037 | N/A | 3.3.1 | unknown | unknown | Blood | 2012 | Africa | South Africa | South Africa | unknown | unknown | Wong et al, 2015 |
| 363192 | ERR338039 | N/A | 3.3.1 | unknown | unknown | Blood | 2009 | Africa | South Africa | South Africa | unknown | unknown | Wong et al, 2015 |
| 316083 | ERR338040 | N/A | 2.5 | unknown | unknown | Blood | 2009 | Africa | South Africa | South Africa | unknown | unknown | Wong et al, 2015 |
| 671445 | ERR338041 | N/A | 2.4.1 | unknown | unknown | Stool | 2012 | Africa | South Africa | South Africa | unknown | unknown | Wong et al, 2015 |
| 325681 | ERR338042 | N/A | 4.1.1 | unknown | unknown | CSF & Blood | 2009 | Africa | South Africa | South Africa | unknown | unknown | Wong et al, 2015 |
| 338900 | ERR338043 | N/A | 2.4.1 | unknown | unknown | Blood | 2009 | Africa | South Africa | South Africa | unknown | unknown | Wong et al, 2015 |
| 400746 | ERR338044 | N/A | 2.4 | unknown | unknown | Blood | 2009 | Africa | South Africa | South Africa | unknown | unknown | Wong et al, 2015 |
| 379905 | ERR338046 | N/A | 2.4.1 | unknown | unknown | Blood | 2009 | Africa | South Africa | South Africa | unknown | unknown | Wong et al, 2015 |
| 634660 | ERR338047 | N/A | 2.4 | unknown | unknown | Blood | 2012 | Africa | South Africa | South Africa | unknown | unknown | Wong et al, 2015 |
| 671076 | ERR338049 | N/A | 3.1 | unknown | unknown | Blood | 2012 | Africa | South Africa | South Africa | unknown | unknown | Wong et al, 2015 |
| A102 | ERR338050 | N/A | 2.1.9 | unknown | unknown | Blood | 2005 | Asia | Southeast Asia | Indonesia | unknown | unknown | Wong et al, 2015 |
| A105 | ERR338051 | N/A | 3.1.2 | unknown | unknown | Blood | 2012 | Asia | Southeast Asia | Indonesia | unknown | unknown | Wong et al, 2015 |
| A109 | ERR338052 | N/A | 2.1.3 | unknown | unknown | Blood | 2004 | Asia | Southeast Asia | Indonesia | unknown | unknown | Wong et al, 2015 |
| A122 | ERR338058 | N/A | 3.1.2 | unknown | unknown | Blood | 2005 | Asia | Southeast Asia | Indonesia | unknown | unknown | Wong et al, 2015 |
| A125 | ERR338060 | N/A | 3.1.2 | unknown | unknown | Blood | 2005 | Asia | Southeast Asia | Indonesia | unknown | unknown | Wong et al, 2015 |
| A127 | ERR338061 | N/A | 2.1.8 | unknown | unknown | Blood | 2011 | Asia | Southeast Asia | Indonesia | unknown | unknown | Wong et al, 2015 |
| A128 | ERR338062 | N/A | 3.1.2 | unknown | unknown | Blood | 2008 | Asia | Southeast Asia | Indonesia | unknown | unknown | Wong et al, 2015 |
| A131 | ERR338064 | N/A | 3 | unknown | unknown | Blood | 2004 | Asia | Southeast Asia | Indonesia | unknown | unknown | Wong et al, 2015 |
| A132 | ERR338065 | N/A | 3.1.2 | unknown | unknown | Blood | 2011 | Asia | Southeast Asia | Indonesia | unknown | unknown | Wong et al, 2015 |
| A135 | ERR338067 | N/A | 3.1.2 | unknown | unknown | Blood | 2004 | Asia | Southeast Asia | Indonesia | unknown | unknown | Wong et al, 2015 |
| A136 | ERR338068 | N/A | 3.1.2 | unknown | unknown | Blood | 2010 | Asia | Southeast Asia | Indonesia | unknown | unknown | Wong et al, 2015 |
| A137 | ERR338069 | N/A | 4.1 | unknown | unknown | Blood | 2008 | Asia | Southeast Asia | Indonesia | unknown | unknown | Wong et al, 2015 |
| Gen-0001 | ERR338070 | N/A | 3.5.4 | unknown | unknown | Not provided | 2012 | Australia & Oceania | Oceania | Samoa | unknown | unknown | Wong et al, 2015 |

| | | | | | | | | | | | | | |
|---|---|---|---|---|---|---|---|---|---|---|---|---|---|
| Gen-0002 | ERR338071 | N/A | 3.5.4 | unknown | unknown | Not provided | 2012 | Australia & Oceania | Oceania | Samoa | unknown | unknown | Wong et al, 2015 |
| Gen-0003 | ERR338072 | N/A | 3.5.4 | unknown | unknown | Not provided | 2012 | Australia & Oceania | Oceania | Samoa | unknown | unknown | Wong et al, 2015 |
| Gen-0004 | ERR338073 | N/A | 3.5.4 | unknown | unknown | Not provided | 2012 | Australia & Oceania | Oceania | Samoa | unknown | unknown | Wong et al, 2015 |
| Gen-0005 | ERR338074 | N/A | 3.5.4 | unknown | unknown | Not provided | 2012 | Australia & Oceania | Oceania | Samoa | unknown | unknown | Wong et al, 2015 |
| Gen-0006 | ERR338075 | N/A | 3.5.4 | unknown | unknown | Not provided | 2012 | Australia & Oceania | Oceania | Samoa | unknown | unknown | Wong et al, 2015 |
| Gen-0007 | ERR338076 | N/A | 3.5.4 | unknown | unknown | Not provided | 2012 | Australia & Oceania | Oceania | Samoa | unknown | unknown | Wong et al, 2015 |
| Gen-0008 | ERR338077 | N/A | 3.5.4 | unknown | unknown | Not provided | 2012 | Australia & Oceania | Oceania | Samoa | unknown | unknown | Wong et al, 2015 |
| Gen-0009 | ERR338078 | N/A | 3.5.4 | unknown | unknown | Not provided | 2012 | Australia & Oceania | Oceania | Samoa | unknown | unknown | Wong et al, 2015 |
| Gen-0010 | ERR338079 | N/A | 3.5.4 | unknown | unknown | Not provided | 2012 | Australia & Oceania | Oceania | Samoa | unknown | unknown | Wong et al, 2015 |
| Gen-0011 | ERR338080 | N/A | 3.5.4 | unknown | unknown | Not provided | 2012 | Australia & Oceania | Oceania | Samoa | unknown | unknown | Wong et al, 2015 |
| Gen-0012 | ERR338081 | N/A | 3.5.4 | unknown | unknown | Not provided | 2012 | Australia & Oceania | Oceania | Samoa | unknown | unknown | Wong et al, 2015 |
| Gen-0013 | ERR338082 | N/A | 3.5.4 | unknown | unknown | Not provided | 2012 | Australia & Oceania | Oceania | Samoa | unknown | unknown | Wong et al, 2015 |
| Gen-0014 | ERR338083 | N/A | 3.5.4 | unknown | unknown | Not provided | 2012 | Australia & Oceania | Oceania | Samoa | unknown | unknown | Wong et al, 2015 |
| Gen-0015 | ERR338084 | N/A | 3.5.4 | unknown | unknown | Not provided | 2012 | Australia & Oceania | Oceania | Samoa | unknown | unknown | Wong et al, 2015 |
| Gen-0016 | ERR338085 | N/A | 3.5.4 | unknown | unknown | Not provided | 2012 | Australia & Oceania | Oceania | Samoa | unknown | unknown | Wong et al, 2015 |
| Gen-0017 | ERR338086 | N/A | 3.5.4 | unknown | unknown | Not provided | 2012 | Australia & Oceania | Oceania | Samoa | unknown | unknown | Wong et al, 2015 |
| Gen-0018 | ERR338087 | N/A | 3.5.4 | unknown | unknown | Not provided | 2012 | Australia & Oceania | Oceania | Samoa | unknown | unknown | Wong et al, 2015 |
| Gen-0019 | ERR338088 | N/A | 3.5.4 | unknown | unknown | Not provided | 2012 | Australia & Oceania | Oceania | Samoa | unknown | unknown | Wong et al, 2015 |
| Gen-0020 | ERR338089 | N/A | 3.5.4 | unknown | unknown | Not provided | 2012 | Australia & Oceania | Oceania | Samoa | unknown | unknown | Wong et al, 2015 |
| Gen-0021 | ERR338090 | N/A | 3.5.4 | unknown | unknown | Not provided | 2012 | Australia & Oceania | Oceania | Samoa | unknown | unknown | Wong et al, 2015 |
| Gen-0022 | ERR338091 | N/A | 3.5.4 | unknown | unknown | Not provided | 2012 | Australia & Oceania | Oceania | Samoa | unknown | unknown | Wong et al, 2015 |
| Gen-0023 | ERR338092 | N/A | 3.5.4 | unknown | unknown | Not provided | 2012 | Australia & Oceania | Oceania | Samoa | unknown | unknown | Wong et al, 2015 |
| Gen-0024 | ERR338093 | N/A | 3.5.4 | unknown | unknown | Not provided | 2012 | Australia & Oceania | Oceania | Samoa | unknown | unknown | Wong et al, 2015 |
| Gen-0025 | ERR338094 | N/A | 3.5.4 | unknown | unknown | Not provided | 2012 | Australia & Oceania | Oceania | Samoa | unknown | unknown | Wong et al, 2015 |
| Gen-0026 | ERR338095 | N/A | 3.5.4 | unknown | unknown | Not provided | 2012 | Australia & Oceania | Oceania | Samoa | unknown | unknown | Wong et al, 2015 |
| Gen-0027 | ERR338096 | N/A | 3.5.4 | unknown | unknown | Not provided | 2012 | Australia & Oceania | Oceania | Samoa | unknown | unknown | Wong et al, 2015 |
| Gen-0028 | ERR338097 | N/A | 3.5.4 | unknown | unknown | Not provided | 2012 | Australia & Oceania | Oceania | Samoa | unknown | unknown | Wong et al, 2015 |
| Gen-0029 | ERR338098 | N/A | 3.5.4 | unknown | unknown | Not provided | 2012 | Australia & Oceania | Oceania | Samoa | unknown | unknown | Wong et al, 2015 |
| Gen-0030 | ERR338099 | N/A | 4.1 | unknown | unknown | Not provided | 2012 | Australia & Oceania | Oceania | Samoa | unknown | unknown | Wong et al, 2015 |
| Gen-0031 | ERR338100 | N/A | 3.5.4 | unknown | unknown | Not provided | 2012 | Australia & Oceania | Oceania | Samoa | unknown | unknown | Wong et al, 2015 |
| Gen-0032 | ERR338101 | N/A | 3.5.4 | unknown | unknown | Not provided | 2012 | Australia & Oceania | Oceania | Samoa | unknown | unknown | Wong et al, 2015 |
| Gen-0033 | ERR338102 | N/A | 3.5.4 | unknown | unknown | Not provided | 2012 | Australia & Oceania | Oceania | Samoa | unknown | unknown | Wong et al, 2015 |
| Gen-0034 | ERR338103 | N/A | 3.5.4 | unknown | unknown | Not provided | 2012 | Australia & Oceania | Oceania | Samoa | unknown | unknown | Wong et al, 2015 |
| Gen-0035 | ERR338104 | N/A | 3.5.4 | unknown | unknown | Not provided | 2012 | Australia & Oceania | Oceania | Samoa | unknown | unknown | Wong et al, 2015 |
| Gen-0036 | ERR338105 | N/A | 3.5.4 | unknown | unknown | Not provided | 2012 | Australia & Oceania | Oceania | Samoa | unknown | unknown | Wong et al, 2015 |
| Gen-0037 | ERR338106 | N/A | 3.5.4 | unknown | unknown | Not provided | 2012 | Australia & Oceania | Oceania | Samoa | unknown | unknown | Wong et al, 2015 |
| Gen-0038 | ERR338107 | N/A | 3.5.4 | unknown | unknown | Not provided | 2012 | Australia & Oceania | Oceania | Samoa | unknown | unknown | Wong et al, 2015 |
| Gen-0039 | ERR338108 | N/A | 3.5.4 | unknown | unknown | Not provided | 2012 | Australia & Oceania | Oceania | Samoa | unknown | unknown | Wong et al, 2015 |
| Gen-0040 | ERR338109 | N/A | 3.5.4 | unknown | unknown | Not provided | 2012 | Australia & Oceania | Oceania | Samoa | unknown | unknown | Wong et al, 2015 |
| Gen-0041 | ERR338110 | N/A | 3.5.4 | unknown | unknown | Not provided | 2012 | Australia & Oceania | Oceania | Samoa | unknown | unknown | Wong et al, 2015 |
| Gen-0042 | ERR338111 | N/A | 3.5.4 | unknown | unknown | Not provided | 2012 | Australia & Oceania | Oceania | Samoa | unknown | unknown | Wong et al, 2015 |
| Gen-0043 | ERR338112 | N/A | 3.5.4 | unknown | unknown | Not provided | 2012 | Australia & Oceania | Oceania | Samoa | unknown | unknown | Wong et al, 2015 |
| Gen-0044 | ERR338113 | N/A | 3.5.4 | unknown | unknown | Not provided | 2012 | Australia & Oceania | Oceania | Samoa | unknown | unknown | Wong et al, 2015 |
| Gen-0045 | ERR338114 | N/A | 3.5.4 | unknown | unknown | Not provided | 2012 | Australia & Oceania | Oceania | Samoa | unknown | unknown | Wong et al, 2015 |
| Gen-0046 | ERR338115 | N/A | 3.5.4 | unknown | unknown | Not provided | 2012 | Australia & Oceania | Oceania | Samoa | unknown | unknown | Wong et al, 2015 |
| Gen-0047 | ERR338116 | N/A | 3.5.4 | unknown | unknown | Not provided | 2012 | Australia & Oceania | Oceania | Samoa | unknown | unknown | Wong et al, 2015 |
| Gen-0048 | ERR338117 | N/A | 3.5.4 | unknown | unknown | Not provided | 2012 | Australia & Oceania | Oceania | Samoa | unknown | unknown | Wong et al, 2015 |
| Gen-0049 | ERR338118 | N/A | 3.5.4 | unknown | unknown | Not provided | 2012 | Australia & Oceania | Oceania | Samoa | unknown | unknown | Wong et al, 2015 |
| Gen-0050 | ERR338119 | N/A | 3.5.4 | unknown | unknown | Not provided | 2012 | Australia & Oceania | Oceania | Samoa | unknown | unknown | Wong et al, 2015 |
| Gen-0051 | ERR338120 | N/A | 3.5.4 | unknown | unknown | Not provided | 2012 | Australia & Oceania | Oceania | Samoa | unknown | unknown | Wong et al, 2015 |

| | | | | | | | | | | | | | |
|---|---|---|---|---|---|---|---|---|---|---|---|---|---|
| Gen-0053 | ERR338122 | N/A | 3.5.4 | unknown | unknown | Not provided | 2012 | Australia & Oceania | Oceania | Samoa | unknown | unknown | Wong et al, 2015 |
| Gen-0054 | ERR338123 | N/A | 3.5.4 | unknown | unknown | Not provided | 2012 | Australia & Oceania | Oceania | Samoa | unknown | unknown | Wong et al, 2015 |
| Gen-0055 | ERR338124 | N/A | 3.5.4 | unknown | unknown | Not provided | 2012 | Australia & Oceania | Oceania | Samoa | unknown | unknown | Wong et al, 2015 |
| Gen-0056 | ERR338125 | N/A | 3.5.4 | unknown | unknown | Not provided | 2012 | Australia & Oceania | Oceania | Samoa | unknown | unknown | Wong et al, 2015 |
| Gen-0058 | ERR338127 | N/A | 3.5.4 | unknown | unknown | Not provided | 2012 | Australia & Oceania | Oceania | Samoa | unknown | unknown | Wong et al, 2015 |
| Gen-0059 | ERR338128 | N/A | 4.1 | unknown | unknown | Not provided | 2012 | Australia & Oceania | Oceania | Samoa | unknown | unknown | Wong et al, 2015 |
| Gen-0060 | ERR338129 | N/A | 4.1 | unknown | unknown | Not provided | 2012 | Australia & Oceania | Oceania | Samoa | unknown | unknown | Wong et al, 2015 |
| 2010K-0515 | ERR338131 | N/A | 4.3.1.1.EA1 | unknown | unknown | Blood | 2007 | Africa | East Africa | Kenya | unknown | unknown | Wong et al, 2015 |
| 2010K-0552 | ERR338132 | N/A | 4.3.1.1.EA1 | unknown | unknown | Blood | 2008 | Africa | East Africa | Kenya | unknown | unknown | Wong et al, 2015 |
| 2010K-0527 | ERR338133 | N/A | 4.3.1.1.EA1 | unknown | unknown | Blood | 2007 | Africa | East Africa | Kenya | unknown | unknown | Wong et al, 2015 |
| 2010K-0507 | ERR338134 | N/A | 4.3.1.2.EA2 | unknown | unknown | Blood | 2007 | Africa | East Africa | Kenya | unknown | unknown | Wong et al, 2015 |
| 2010K-0519 | ERR338135 | N/A | 4.3.1.2.EA2 | unknown | unknown | Blood | 2007 | Africa | East Africa | Kenya | unknown | unknown | Wong et al, 2015 |
| 2010K-0533 | ERR338136 | N/A | 4.3.1.1.EA1 | unknown | unknown | Blood | 2008 | Africa | East Africa | Kenya | unknown | unknown | Wong et al, 2015 |
| 2010K-0512 | ERR338137 | N/A | 4.3.1.1.EA1 | unknown | unknown | Blood | 2007 | Africa | East Africa | Kenya | unknown | unknown | Wong et al, 2015 |
| 2010K-0530 | ERR338138 | N/A | 4.3.1.2.EA2 | unknown | unknown | Blood | 2007 | Africa | East Africa | Kenya | unknown | unknown | Wong et al, 2015 |
| 2010K-0553 | ERR338139 | N/A | 4.3.1.1.EA1 | unknown | unknown | Blood | 2008 | Africa | East Africa | Kenya | unknown | unknown | Wong et al, 2015 |
| 2010K-0550 | ERR338140 | N/A | 4.3.1.1.EA1 | unknown | unknown | Blood | 2009 | Africa | East Africa | Kenya | unknown | unknown | Wong et al, 2015 |
| 2010K-0511 | ERR338141 | N/A | 4.3.1.1.EA1 | unknown | unknown | Blood | 2007 | Africa | East Africa | Kenya | unknown | unknown | Wong et al, 2015 |
| 2010K-0539 | ERR338143 | N/A | 4.3.1.1.EA1 | unknown | unknown | Blood | 2008 | Africa | East Africa | Kenya | unknown | unknown | Wong et al, 2015 |
| 2010K-0522 | ERR338145 | N/A | 4.3.1.1.EA1 | unknown | unknown | Blood | 2007 | Africa | East Africa | Kenya | unknown | unknown | Wong et al, 2015 |
| 2010K-0517 | ERR338146 | N/A | 4.3.1.2.EA2 | unknown | unknown | Blood | 2007 | Africa | East Africa | Kenya | unknown | unknown | Wong et al, 2015 |
| 2010K-0548 | ERR338147 | N/A | 4.3.1.1.EA1 | unknown | unknown | Blood | 2008 | Africa | East Africa | Kenya | unknown | unknown | Wong et al, 2015 |
| 2010K-0509 | ERR338148 | N/A | 4.3.1.1.EA1 | unknown | unknown | Blood | 2007 | Africa | East Africa | Kenya | unknown | unknown | Wong et al, 2015 |
| 2010K-0500 | ERR338149 | N/A | 4.3.1.2.EA2 | unknown | unknown | Stool | 2007 | Africa | East Africa | Kenya | unknown | unknown | Wong et al, 2015 |
| 2010K-0506 | ERR338150 | N/A | 4.3.1.2.EA2 | unknown | unknown | Blood | 2007 | Africa | East Africa | Kenya | unknown | unknown | Wong et al, 2015 |
| 2010K-0554 | ERR338151 | N/A | 4.3.1.1.EA1 | unknown | unknown | Blood | 2008 | Africa | East Africa | Kenya | unknown | unknown | Wong et al, 2015 |
| 2010K-0508 | ERR338152 | N/A | 4.3.1.1.EA1 | unknown | unknown | Blood | 2007 | Africa | East Africa | Kenya | unknown | unknown | Wong et al, 2015 |
| 2010K-0626 | ERR338153 | N/A | 3.3.1 | unknown | unknown | Blood | 2009 | Africa | East Africa | Kenya | unknown | unknown | Wong et al, 2015 |
| 2010K-0524 | ERR338157 | N/A | 4.3.1.1.EA1 | unknown | unknown | Blood | 2007 | Africa | East Africa | Kenya | unknown | unknown | Wong et al, 2015 |
| 2010K-0523 | ERR338158 | N/A | 4.3.1.2.EA2 | unknown | unknown | Blood | 2007 | Africa | East Africa | Kenya | unknown | unknown | Wong et al, 2015 |
| 2010K-0544 | ERR338159 | N/A | 4.3.1.1.EA1 | unknown | unknown | Blood | 2009 | Africa | East Africa | Kenya | unknown | unknown | Wong et al, 2015 |
| 2010K-0545 | ERR338160 | N/A | 4.3.1.1.EA1 | unknown | unknown | Blood | 2008 | Africa | East Africa | Kenya | unknown | unknown | Wong et al, 2015 |
| 2010K-0538 | ERR338161 | N/A | 4.3.1.2.EA2 | unknown | unknown | Blood | 2008 | Africa | East Africa | Kenya | unknown | unknown | Wong et al, 2015 |
| 2010K-0535 | ERR338162 | N/A | 3.3.1 | unknown | unknown | Blood | 2009 | Africa | East Africa | Kenya | unknown | unknown | Wong et al, 2015 |
| 2010K-0534 | ERR338163 | N/A | 4.3.1.1.EA1 | unknown | unknown | Blood | 2008 | Africa | East Africa | Kenya | unknown | unknown | Wong et al, 2015 |
| 2010K-0514 | ERR338164 | N/A | 4.3.1.1.EA1 | unknown | unknown | Blood | 2007 | Africa | East Africa | Kenya | unknown | unknown | Wong et al, 2015 |
| 2007-001525 | ERR340757 | N/A | 4.3.1.1 | unknown | unknown | Blood | 2007 | Asia | Southeast Asia | Cambodia | unknown | unknown | Wong et al, 2015 |
| 2007-001619 | ERR340758 | N/A | 4.3.1.1 | unknown | unknown | Blood | 2007 | Asia | Southeast Asia | Cambodia | unknown | unknown | Wong et al, 2015 |
| 2001-102645 | ERR340759 | N/A | 4.3.1.1 | unknown | unknown | Blood | 2007 | Asia | Southeast Asia | Cambodia | unknown | unknown | Wong et al, 2015 |
| 2007-003270 | ERR340760 | N/A | 4.3.1.1 | unknown | unknown | Blood | 2007 | Asia | Southeast Asia | Cambodia | unknown | unknown | Wong et al, 2015 |
| 2007-006110 | ERR340761 | N/A | 4.3.1.1 | unknown | unknown | Blood | 2007 | Asia | Southeast Asia | Cambodia | unknown | unknown | Wong et al, 2015 |
| 2007-005516 | ERR340763 | N/A | 4.3.1.1 | unknown | unknown | Blood | 2007 | Asia | Southeast Asia | Cambodia | unknown | unknown | Wong et al, 2015 |
| 2007-007073 | ERR340765 | N/A | 3.2.1 | unknown | unknown | Blood | 2007 | Asia | Southeast Asia | Cambodia | unknown | unknown | Wong et al, 2015 |
| 2007-007506 | ERR340766 | N/A | 4.3.1.1 | unknown | unknown | Blood | 2007 | Asia | Southeast Asia | Cambodia | unknown | unknown | Wong et al, 2015 |
| 2008-006685 | ERR340767 | N/A | 4.3.1.1 | unknown | unknown | Blood | 2008 | Asia | Southeast Asia | Cambodia | unknown | unknown | Wong et al, 2015 |
| 2007-011815 | ERR340768 | N/A | 4.3.1.1 | unknown | unknown | Blood | 2007 | Asia | Southeast Asia | Cambodia | unknown | unknown | Wong et al, 2015 |
| 2001-100239 | ERR340769 | N/A | 3.2.1 | unknown | unknown | Blood | 2007 | Asia | Southeast Asia | Cambodia | unknown | unknown | Wong et al, 2015 |
| 2007-015823 | ERR340770 | N/A | 4.3.1.1 | unknown | unknown | Blood | 2007 | Asia | Southeast Asia | Cambodia | unknown | unknown | Wong et al, 2015 |
| 2007-023117 | ERR340771 | N/A | 4.3.1.1 | unknown | unknown | Blood | 2007 | Asia | Southeast Asia | Cambodia | unknown | unknown | Wong et al, 2015 |
| 2007-024445 | ERR340772 | N/A | 4.3.1.1 | unknown | unknown | Blood | 2007 | Asia | Southeast Asia | Cambodia | unknown | unknown | Wong et al, 2015 |

| | | | | | | | | | | | | | |
|---|---|---|---|---|---|---|---|---|---|---|---|---|---|
| 2007-023116 | ERR340773 | N/A | 4.3.1.1 | unknown | unknown | Blood | 2007 | Asia | Southeast Asia | Cambodia | unknown | unknown | Wong et al, 2015 |
| 2007-024758 | ERR340774 | N/A | 4.3.1.1 | unknown | unknown | Blood | 2007 | Asia | Southeast Asia | Cambodia | unknown | unknown | Wong et al, 2015 |
| 2007-007363 | ERR340775 | N/A | 4.3.1.1 | unknown | unknown | Blood | 2007 | Asia | Southeast Asia | Cambodia | unknown | unknown | Wong et al, 2015 |
| LNT 08 | ERR340777 | N/A | 3.4 | unknown | unknown | Blood | 2001 | Asia | Southeast Asia | Laos | unknown | unknown | Wong et al, 2015 |
| LNT1108 | ERR340778 | N/A | 3.4 | unknown | unknown | Blood | 2001 | Asia | Southeast Asia | Laos | unknown | unknown | Wong et al, 2015 |
| LNT1148 | ERR340779 | N/A | 3.4 | unknown | unknown | Blood | 2001 | Asia | Southeast Asia | Laos | unknown | unknown | Wong et al, 2015 |
| LNT22 | ERR340780 | N/A | 3.4 | unknown | unknown | Blood | 2001 | Asia | Southeast Asia | Laos | unknown | unknown | Wong et al, 2015 |
| LNT26 | ERR340781 | N/A | 3.4 | unknown | unknown | Blood | 2001 | Asia | Southeast Asia | Laos | unknown | unknown | Wong et al, 2015 |
| LNT358 | ERR340782 | N/A | 3.4 | unknown | unknown | Blood | 2000 | Asia | Southeast Asia | Laos | unknown | unknown | Wong et al, 2015 |
| UI1059 | ERR340783 | N/A | 4.3.1.1 | unknown | unknown | Blood | 2001 | Asia | Southeast Asia | Laos | unknown | unknown | Wong et al, 2015 |
| UI1151 | ERR340784 | N/A | 4.3.1.1 | unknown | unknown | Blood | 2001 | Asia | Southeast Asia | Laos | unknown | unknown | Wong et al, 2015 |
| UI1203 | ERR340785 | N/A | 3.2.1 | unknown | unknown | Blood | 2001 | Asia | Southeast Asia | Laos | unknown | unknown | Wong et al, 2015 |
| UI1236 | ERR340786 | N/A | 3.2.1 | unknown | unknown | Blood | 2001 | Asia | Southeast Asia | Laos | unknown | unknown | Wong et al, 2015 |
| UI1954-12/3 | ERR340787 | N/A | 4.3.1.1 | unknown | unknown | Blood | 2002 | Asia | Southeast Asia | Laos | unknown | unknown | Wong et al, 2015 |
| UI1954-2/3 | ERR340788 | N/A | 3.2.1 | unknown | unknown | Blood | 2002 | Asia | Southeast Asia | Laos | unknown | unknown | Wong et al, 2015 |
| UI1965 | ERR340789 | N/A | 4.3.1.1 | unknown | unknown | Blood | 2002 | Asia | Southeast Asia | Laos | unknown | unknown | Wong et al, 2015 |
| UI1998 | ERR340790 | N/A | 4.3.1.1 | unknown | unknown | Blood | 2002 | Asia | Southeast Asia | Laos | unknown | unknown | Wong et al, 2015 |
| UI2000 | ERR340791 | N/A | 4.3.1.1 | unknown | unknown | Blood | 2002 | Asia | Southeast Asia | Laos | unknown | unknown | Wong et al, 2015 |
| UI2001 | ERR340792 | N/A | 2.2.3,2.2.2 | unknown | unknown | Blood | 2002 | Asia | Southeast Asia | Laos | unknown | unknown | Wong et al, 2015 |
| UI2006 | ERR340793 | N/A | 3.2.1 | unknown | unknown | Blood | 2002 | Asia | Southeast Asia | Laos | unknown | unknown | Wong et al, 2015 |
| UI2063 | ERR340794 | N/A | 4.3.1.1 | unknown | unknown | Blood | 2002 | Asia | Southeast Asia | Laos | unknown | unknown | Wong et al, 2015 |
| UI2120 | ERR340795 | N/A | 4.3.1.1 | unknown | unknown | Blood | 2002 | Asia | Southeast Asia | Laos | unknown | unknown | Wong et al, 2015 |
| UI2144 | ERR340797 | N/A | 4.3.1.1 | unknown | unknown | Blood | 2002 | Asia | Southeast Asia | Laos | unknown | unknown | Wong et al, 2015 |
| UI2152 | ERR340798 | N/A | 4.1 | unknown | unknown | Blood | 2002 | Asia | Southeast Asia | Laos | unknown | unknown | Wong et al, 2015 |
| UI2155 | ERR340799 | N/A | 4.3.1.1 | unknown | unknown | Blood | 2002 | Asia | Southeast Asia | Laos | unknown | unknown | Wong et al, 2015 |
| UI2220 | ERR340800 | N/A | 4.3.1.1 | unknown | unknown | Blood | 2002 | Asia | Southeast Asia | Laos | unknown | unknown | Wong et al, 2015 |
| UI2351 | ERR340801 | N/A | 3.2.1 | unknown | unknown | Blood | 2002 | Asia | Southeast Asia | Laos | unknown | unknown | Wong et al, 2015 |
| UI247 | ERR340802 | N/A | 4.3.1.1 | unknown | unknown | Blood | 2000 | Asia | Southeast Asia | Laos | unknown | unknown | Wong et al, 2015 |
| UI2591 | ERR340803 | N/A | 3.4 | unknown | unknown | Blood | 2002 | Asia | Southeast Asia | Laos | unknown | unknown | Wong et al, 2015 |
| UI2658 | ERR340804 | N/A | 3.4 | unknown | unknown | Blood | 2002 | Asia | Southeast Asia | Laos | unknown | unknown | Wong et al, 2015 |
| UI2679 | ERR340805 | N/A | 3.2.1 | unknown | unknown | Blood | 2002 | Asia | Southeast Asia | Laos | unknown | unknown | Wong et al, 2015 |
| UI2824 | ERR340806 | N/A | 4.3.1.1 | unknown | unknown | Blood | 2002 | Asia | Southeast Asia | Laos | unknown | unknown | Wong et al, 2015 |
| UI286 | ERR340807 | N/A | 4.3.1.1 | unknown | unknown | Blood | 2000 | Asia | Southeast Asia | Laos | unknown | unknown | Wong et al, 2015 |
| UI3146 | ERR340808 | N/A | 2.3.4 | unknown | unknown | Blood | 2002 | Asia | Southeast Asia | Laos | unknown | unknown | Wong et al, 2015 |
| UI315 | ERR340809 | N/A | 4.3.1.1 | unknown | unknown | Blood | 2000 | Asia | Southeast Asia | Laos | unknown | unknown | Wong et al, 2015 |
| UI3151 | ERR340810 | N/A | 2.3.4 | unknown | unknown | Blood | 2002 | Asia | Southeast Asia | Laos | unknown | unknown | Wong et al, 2015 |
| UI3260 | ERR340811 | N/A | 3.2.1 | unknown | unknown | Blood | 2003 | Asia | Southeast Asia | Laos | unknown | unknown | Wong et al, 2015 |
| UI3275 | ERR340812 | N/A | 4.3.1.1 | unknown | unknown | Blood | 2003 | Asia | Southeast Asia | Laos | unknown | unknown | Wong et al, 2015 |
| UI3313 | ERR340813 | N/A | 3.4 | unknown | unknown | Blood | 2003 | Asia | Southeast Asia | Laos | unknown | unknown | Wong et al, 2015 |
| UI3396 | ERR340814 | N/A | 2.2.3,2.2.2 | unknown | unknown | Blood | 2003 | Asia | Southeast Asia | Laos | unknown | unknown | Wong et al, 2015 |
| UI473 | ERR340815 | N/A | 4.3.1.1 | unknown | unknown | Blood | 2000 | Asia | Southeast Asia | Laos | unknown | unknown | Wong et al, 2015 |
| UI54 | ERR340816 | N/A | 4.1 | unknown | unknown | Blood | 2000 | Asia | Southeast Asia | Laos | unknown | unknown | Wong et al, 2015 |
| ERL021174 | ERR343249 | N/A | 3.5.4 | unknown | unknown | Blood | 2002 | Australia & Oceania | Oceania | Samoa | unknown | unknown | Wong et al, 2015 |
| ERL022463 | ERR343250 | N/A | 2.1.4 | unknown | unknown | Stool | 2002 | Asia | Western Asia | Western Asia | unknown | unknown | Wong et al, 2015 |
| ERL022464 | ERR343251 | N/A | 2.1.2 | unknown | unknown | Blood | 2002 | Asia | Western Asia | Western Asia | unknown | unknown | Wong et al, 2015 |
| ERL024120 | ERR343252 | N/A | 2.1.2 | unknown | unknown | Stool | 2002 | Asia | Southeast Asia | Indonesia | unknown | unknown | Wong et al, 2015 |
| ERL024182 | ERR343253 | N/A | 2.5 | unknown | unknown | Blood | 2002 | Asia | South Asia | India | unknown | unknown | Wong et al, 2015 |
| ERL02425 | ERR343254 | N/A | 2.5 | unknown | unknown | Stool | 2002 | Asia | South Asia | India | unknown | unknown | Wong et al, 2015 |
| ERL024919 | ERR343255 | N/A | 3.5.4 | unknown | unknown | Blood | 2002 | Australia & Oceania | Oceania | Samoa | unknown | unknown | Wong et al, 2015 |
| ERL02732 | ERR343256 | N/A | 3.1.2 | unknown | unknown | Blood | 2002 | Asia | Southeast Asia | Indonesia | unknown | unknown | Wong et al, 2015 |

| | | | | | | | | | | | | | |
|---|---|---|---|---|---|---|---|---|---|---|---|---|---|
| ERL032200 | ERR343257 | N/A | 4.2 | unknown | unknown | Blood | 2003 | Australia & Oceania | Oceania | Tonga | unknown | unknown | Wong et al, 2015 |
| ERL032330 | ERR343258 | N/A | 4.3.1.2 | unknown | unknown | Blood | 2003 | Asia | South Asia | India | unknown | unknown | Wong et al, 2015 |
| ERL034151 | ERR343260 | N/A | 2.2.1 | unknown | unknown | Blood | 2003 | Asia | Southeast Asia | Indonesia | unknown | unknown | Wong et al, 2015 |
| ERL04140 | ERR343262 | N/A | 4.3.1.2 | unknown | unknown | Blood | 2004 | Asia | South Asia | India | unknown | unknown | Wong et al, 2015 |
| ERL041419 | ERR343263 | N/A | 4.1 | unknown | unknown | Blood | 2004 | Asia | South Asia | India | unknown | unknown | Wong et al, 2015 |
| ERL041834 | ERR343264 | N/A | 3.3 | unknown | unknown | Stool | 2004 | Asia | South Asia | India | unknown | unknown | Wong et al, 2015 |
| ERL041932 | ERR343265 | N/A | 4.1 | unknown | unknown | Blood | 2004 | Australia & Oceania | Oceania | Samoa | unknown | unknown | Wong et al, 2015 |
| ERL042857 | ERR343266 | N/A | 3.5.4 | unknown | unknown | Blood | 2004 | Australia & Oceania | Oceania | Samoa | unknown | unknown | Wong et al, 2015 |
| ERL043008 | ERR343267 | N/A | 3.5.4 | unknown | unknown | Stool | 2004 | Australia & Oceania | Oceania | Samoa | unknown | unknown | Wong et al, 2015 |
| ERL052042 | ERR343269 | N/A | 4.3.1.1 | unknown | unknown | Blood | 2005 | Asia | South Asia | India | unknown | unknown | Wong et al, 2015 |
| ERL061748 | ERR343270 | N/A | 4.3.1.1 | unknown | unknown | Blood | 2006 | Asia | Southeast Asia | South-east Asia | unknown | unknown | Wong et al, 2015 |
| ERL062282 | ERR343271 | N/A | 4.3.1.2 | unknown | unknown | Blood | 2006 | Asia | South Asia | India | unknown | unknown | Wong et al, 2015 |
| ERL063423 | ERR343272 | N/A | 4.3.1.1 | unknown | unknown | Blood | 2006 | Asia | South Asia | India | unknown | unknown | Wong et al, 2015 |
| ERL063424 | ERR343273 | N/A | 4.3.1.2 | unknown | unknown | Blood | 2006 | Asia | South Asia | India | unknown | unknown | Wong et al, 2015 |
| ERL064553 | ERR343274 | N/A | 3.5.4 | unknown | unknown | Blood | 2006 | Australia & Oceania | Oceania | Samoa | unknown | unknown | Wong et al, 2015 |
| ERL0661 | ERR343275 | N/A | 2.5 | unknown | unknown | Blood | 2006 | Asia | South Asia | India | unknown | unknown | Wong et al, 2015 |
| ERL07263 | ERR343276 | N/A | 3.5.4 | unknown | unknown | Blood | 2007 | Australia & Oceania | Oceania | Samoa | unknown | unknown | Wong et al, 2015 |
| ERL07264 | ERR343277 | N/A | 3.5.4 | unknown | unknown | Blood | 2007 | Australia & Oceania | Oceania | Samoa | unknown | unknown | Wong et al, 2015 |
| ERL072830 | ERR343278 | N/A | 3.5.4 | unknown | unknown | Blood | 2007 | Australia & Oceania | Oceania | Samoa | unknown | unknown | Wong et al, 2015 |
| ERL072973 | ERR343279 | N/A | 4.2.2 | unknown | unknown | Blood | 2007 | Australia & Oceania | Oceania | Fiji | unknown | unknown | Wong et al, 2015 |
| ERL07434 | ERR343280 | N/A | 3.5.4 | unknown | unknown | Blood | 2007 | Australia & Oceania | Oceania | Samoa | unknown | unknown | Wong et al, 2015 |
| ERL082325 | ERR343281 | N/A | 4.2.1 | unknown | unknown | Blood | 2008 | Australia & Oceania | Oceania | Fiji | unknown | unknown | Wong et al, 2015 |
| ERL082356 | ERR343282 | N/A | 4.3.1.2 | unknown | unknown | Stool | 2008 | Asia | South Asia | India | unknown | unknown | Wong et al, 2015 |
| ERL082408 | ERR343283 | N/A | 3.5.4 | unknown | unknown | Blood | 2008 | Australia & Oceania | Oceania | Samoa | unknown | unknown | Wong et al, 2015 |
| ERL082444 | ERR343284 | N/A | 3.5.4 | unknown | unknown | Blood | 2008 | Australia & Oceania | Oceania | Samoa | unknown | unknown | Wong et al, 2015 |
| ERL082759 | ERR343285 | N/A | 4.2.2 | unknown | unknown | Blood | 2008 | Australia & Oceania | Oceania | Fiji | unknown | unknown | Wong et al, 2015 |
| ERL084047 | ERR343286 | N/A | 2.2.2 | unknown | unknown | Stool | 2008 | Asia | South Asia | India | unknown | unknown | Wong et al, 2015 |
| ERL084170 | ERR343287 | N/A | 3.1.2 | unknown | unknown | Blood | 2008 | Asia | East Asia | China | unknown | unknown | Wong et al, 2015 |
| ERL08619 | ERR343288 | N/A | 4.3.1.2 | unknown | unknown | Unknown | 2008 | Asia | South Asia | India | unknown | unknown | Wong et al, 2015 |
| ERL08758 | ERR343289 | N/A | 2.1.7 | unknown | unknown | Blood | 2008 | Asia | South Asia | India | unknown | unknown | Wong et al, 2015 |
| ERL091092 | ERR343291 | N/A | 3.1.2 | unknown | unknown | Blood | 2009 | Asia | Southeast Asia | Indonesia | unknown | unknown | Wong et al, 2015 |
| ERL091300 | ERR343292 | N/A | 3.5.4 | unknown | unknown | Blood | 2009 | Australia & Oceania | Oceania | Samoa | unknown | unknown | Wong et al, 2015 |
| ERL091788 | ERR343293 | N/A | 3.2.2 | unknown | unknown | Blood | 2009 | Asia | Southeast Asia | South-east Asia | unknown | unknown | Wong et al, 2015 |
| ERL091797 | ERR343294 | N/A | 4.3.1.2 | unknown | unknown | Blood | 2009 | Asia | South Asia | India | unknown | unknown | Wong et al, 2015 |
| ERL09383 | ERR343295 | N/A | 3.1 | unknown | unknown | Stool | 2009 | Asia | South Asia | India | unknown | unknown | Wong et al, 2015 |
| ERL094053 | ERR343296 | N/A | 3.5.4 | unknown | unknown | Blood | 2009 | Australia & Oceania | Oceania | Samoa | unknown | unknown | Wong et al, 2015 |
| ERL0982 | ERR343297 | N/A | 3.5.4 | unknown | unknown | Blood | 2009 | Australia & Oceania | Oceania | Samoa | unknown | unknown | Wong et al, 2015 |
| ERL0983 | ERR343298 | N/A | 3.5.4 | unknown | unknown | Blood | 2009 | Australia & Oceania | Oceania | Samoa | unknown | unknown | Wong et al, 2015 |
| ERL09892 | ERR343299 | N/A | 3.5.4 | unknown | unknown | Unknown | 2009 | Australia & Oceania | Oceania | Samoa | unknown | unknown | Wong et al, 2015 |
| ERL101104 | ERR343300 | N/A | 4.3.1.2 | unknown | unknown | Blood | 2010 | Asia | South Asia | India | unknown | unknown | Wong et al, 2015 |
| ERL101621 | ERR343301 | N/A | 2.0.1 | unknown | unknown | Blood | 2010 | Asia | South Asia | India | unknown | unknown | Wong et al, 2015 |
| ERL102156 | ERR343302 | N/A | 4.3.1.1 | unknown | unknown | Blood | 2010 | Asia | South Asia | India | unknown | unknown | Wong et al, 2015 |
| ERL102275 | ERR343303 | N/A | 4.3.1.2 | unknown | unknown | Blood | 2010 | Asia | South Asia | India | unknown | unknown | Wong et al, 2015 |
| ERL102461 | ERR343304 | N/A | 3.5.4 | unknown | unknown | Blood | 2010 | Australia & Oceania | Oceania | Samoa | unknown | unknown | Wong et al, 2015 |
| ERL10320 | ERR343305 | N/A | 3.5.4 | unknown | unknown | Stool | 2010 | Australia & Oceania | Oceania | Samoa | unknown | unknown | Wong et al, 2015 |
| ERL10338 | ERR343306 | N/A | 4.3.1 | unknown | unknown | Blood | 2010 | Asia | Southeast Asia | Thailand | unknown | unknown | Wong et al, 2015 |
| ERL103534 | ERR343307 | N/A | 4.3.1.2 | unknown | unknown | Blood | 2010 | Asia | South Asia | India | unknown | unknown | Wong et al, 2015 |
| ERL103914 | ERR343308 | N/A | 2.3.2 | unknown | unknown | Blood | 2010 | South America | South America | South America | unknown | unknown | Wong et al, 2015 |
| ERL1048 | ERR343309 | N/A | 4.3.1.2 | unknown | unknown | Blood | 2010 | Asia | South Asia | India | unknown | unknown | Wong et al, 2015 |
| ERL10492 | ERR343310 | N/A | 3.5.4 | unknown | unknown | Blood | 2010 | Australia & Oceania | Oceania | Samoa | unknown | unknown | Wong et al, 2015 |

| | | | | | | | | | | | | | |
|---|---|---|---|---|---|---|---|---|---|---|---|---|---|
| ERL10504 | ERR343311 | N/A | 3.5.4 | unknown | unknown | Blood | 2010 | Australia & Oceania | Oceania | Samoa | unknown | unknown | Wong et al, 2015 |
| ERL111572 | ERR343312 | N/A | 3.5.4 | unknown | unknown | Blood | 2011 | Australia & Oceania | Oceania | Samoa | unknown | unknown | Wong et al, 2015 |
| ERL111998 | ERR343313 | N/A | 3.5.4 | unknown | unknown | Blood | 2011 | Australia & Oceania | Oceania | Samoa | unknown | unknown | Wong et al, 2015 |
| ERL11299 | ERR343314 | N/A | 2.0.1 | unknown | unknown | Stool | 2011 | Asia | South Asia | Pakistan | unknown | unknown | Wong et al, 2015 |
| ERL113095 | ERR343315 | N/A | 4.3.1.2 | unknown | unknown | Blood | 2011 | Asia | South Asia | India | unknown | unknown | Wong et al, 2015 |
| ERL113434 | ERR343316 | N/A | 3.5.4 | unknown | unknown | Blood | 2011 | Australia & Oceania | Oceania | Samoa | unknown | unknown | Wong et al, 2015 |
| ERL114000 | ERR343317 | N/A | 2.2 | unknown | unknown | Blood | 2011 | Asia | South Asia | Nepal | unknown | unknown | Wong et al, 2015 |
| ERL114070 | ERR343318 | N/A | 2.2 | unknown | unknown | Blood | 2011 | South America | South America | South America | unknown | unknown | Wong et al, 2015 |
| ERL114224 | ERR343319 | N/A | 4.3.1.2 | unknown | unknown | Blood | 2011 | Asia | South Asia | India | unknown | unknown | Wong et al, 2015 |
| ERL11877 | ERR343320 | N/A | 4.3.1.2 | unknown | unknown | Stool | 2011 | Asia | South Asia | India | unknown | unknown | Wong et al, 2015 |
| ERL11909 | ERR343321 | N/A | 2.0.2 | unknown | unknown | Stool | 2011 | North America | North America | Mexico | unknown | unknown | Wong et al, 2015 |
| ERL12148 | ERR343322 | N/A | 4.3.1.1 | unknown | unknown | Stool | 2012 | Asia | South Asia | India | unknown | unknown | Wong et al, 2015 |
| ERL12375 | ERR343323 | N/A | 4.3.1.2 | unknown | unknown | Blood | 2012 | Asia | South Asia | India | unknown | unknown | Wong et al, 2015 |
| ERL12590 | ERR343324 | N/A | 4.1 | unknown | unknown | Stool | 2012 | Asia | South Asia | India | unknown | unknown | Wong et al, 2015 |
| ERL12680 | ERR343325 | N/A | 2.2.4 | unknown | unknown | Stool | 2012 | Asia | South Asia | India | unknown | unknown | Wong et al, 2015 |
| ERL12959 | ERR343326 | N/A | 3.1.2 | unknown | unknown | Blood | 2012 | Asia | South Asia | India | unknown | unknown | Wong et al, 2015 |
| ERL12960 | ERR343327 | N/A | 4.3.1.1 | unknown | unknown | Unknown | 2012 | Asia | South Asia | India | unknown | unknown | Wong et al, 2015 |
| H12ESR00755-001A | ERR343328 | N/A | 3 | unknown | unknown | Blood | 2012 | Asia | Southeast Asia | Phillipines | unknown | unknown | Wong et al, 2015 |
| H12ESR02737-001A | ERR343329 | N/A | 2.4 | unknown | unknown | Blood | 2012 | Asia | South Asia | India | unknown | unknown | Wong et al, 2015 |
| H12ESR04734-001A | ERR343330 | N/A | 3.3 | unknown | unknown | Blood | 2012 | Asia | South Asia | India | unknown | unknown | Wong et al, 2015 |
| dtc8 | ERR349331 | N/A | 4.3.1.1 | unknown | unknown | Blood | 1994 | Asia | Southeast Asia | Vietnam | unknown | unknown | Wong et al, 2015 |
| np69 | ERR349332 | N/A | 4.3.1.2 | unknown | unknown | Blood | 2011 | Asia | South Asia | Nepal | unknown | unknown | Wong et al, 2015 |
| np74 | ERR349333 | N/A | 4.3.1.2 | unknown | unknown | Blood | 2011 | Asia | South Asia | Nepal | unknown | unknown | Wong et al, 2015 |
| dtc86 | ERR349334 | N/A | 4.3.1.1 | unknown | unknown | Blood | 1994 | Asia | Southeast Asia | Vietnam | unknown | unknown | Wong et al, 2015 |
| np45 | ERR349335 | N/A | 3.3.2 | unknown | unknown | Blood | 2011 | Asia | South Asia | Nepal | unknown | unknown | Wong et al, 2015 |
| np80 | ERR349336 | N/A | 4.3.1.2 | unknown | unknown | Blood | 2011 | Asia | South Asia | Nepal | unknown | unknown | Wong et al, 2015 |
| dtc105 | ERR349337 | N/A | 4.3.1.1 | unknown | unknown | Blood | 1994 | Asia | Southeast Asia | Vietnam | unknown | unknown | Wong et al, 2015 |
| dtc153 | ERR349338 | N/A | 4.3.1.1 | unknown | unknown | Blood | 1995 | Asia | Southeast Asia | Vietnam | unknown | unknown | Wong et al, 2015 |
| np27 | ERR349339 | N/A | 4.3.1.2 | unknown | unknown | Blood | 2011 | Asia | South Asia | Nepal | unknown | unknown | Wong et al, 2015 |
| dtc174 | ERR349341 | N/A | 4.3.1.1 | unknown | unknown | Blood | 1995 | Asia | Southeast Asia | Vietnam | unknown | unknown | Wong et al, 2015 |
| BRD948 | ERR349343 | N/A | 4.1 | unknown | unknown | Not provided | 1996 | Europe | Eastern Europe | Russia | unknown | unknown | Wong et al, 2015 |
| dn191 | ERR349345 | N/A | 4.3.1.1 | unknown | unknown | Blood | 1996 | Asia | Southeast Asia | Vietnam | unknown | unknown | Wong et al, 2015 |
| dn14 | ERR349346 | N/A | 4.3.1.1 | unknown | unknown | Blood | 1995 | Asia | Southeast Asia | Vietnam | unknown | unknown | Wong et al, 2015 |
| dtc79 | ERR349347 | N/A | 4.3.1.1 | unknown | unknown | Blood | 1994 | Asia | Southeast Asia | Vietnam | unknown | unknown | Wong et al, 2015 |
| dn63 | ERR349348 | N/A | 4.3.1.1 | unknown | unknown | Blood | 1995 | Asia | Southeast Asia | Vietnam | unknown | unknown | Wong et al, 2015 |
| dn121 | ERR349349 | N/A | 4.3.1.1 | unknown | unknown | Blood | 1996 | Asia | Southeast Asia | Vietnam | unknown | unknown | Wong et al, 2015 |
| dn136 | ERR349350 | N/A | 4.3.1.1 | unknown | unknown | Blood | 1996 | Asia | Southeast Asia | Vietnam | unknown | unknown | Wong et al, 2015 |
| dtc116 | ERR349351 | N/A | 3.4 | unknown | unknown | Blood | 1995 | Asia | Southeast Asia | Vietnam | unknown | unknown | Wong et al, 2015 |
| dn86 | ERR349353 | N/A | 4.3.1.1 | unknown | unknown | Blood | 1995 | Asia | Southeast Asia | Vietnam | unknown | unknown | Wong et al, 2015 |
| dn93 | ERR349354 | N/A | 4.3.1.1 | unknown | unknown | Blood | 1995 | Asia | Southeast Asia | Vietnam | unknown | unknown | Wong et al, 2015 |
| dn162 | ERR349356 | N/A | 4.3.1.1 | unknown | unknown | Blood | 1996 | Asia | Southeast Asia | Vietnam | unknown | unknown | Wong et al, 2015 |
| dn18 | ERR349357 | N/A | 4.3.1.1 | unknown | unknown | Blood | 1995 | Asia | Southeast Asia | Vietnam | unknown | unknown | Wong et al, 2015 |
| dtc103 | ERR349359 | N/A | 4.3.1.1 | unknown | unknown | Blood | 1994 | Asia | Southeast Asia | Vietnam | unknown | unknown | Wong et al, 2015 |
| dn19 | ERR349360 | N/A | 4.3.1.1 | unknown | unknown | Blood | 1995 | Asia | Southeast Asia | Vietnam | unknown | unknown | Wong et al, 2015 |
| dn160 | ERR349361 | N/A | 4.3.1.1 | unknown | unknown | Blood | 1996 | Asia | Southeast Asia | Vietnam | unknown | unknown | Wong et al, 2015 |
| dtc111 | ERR349362 | N/A | 4.1 | unknown | unknown | Blood | 1994 | Asia | Southeast Asia | Vietnam | unknown | unknown | Wong et al, 2015 |
| dn189 | ERR349363 | N/A | 4.3.1.1 | unknown | unknown | Blood | 1996 | Asia | Southeast Asia | Vietnam | unknown | unknown | Wong et al, 2015 |
| dtc93 | ERR349364 | N/A | 4.3.1.1 | unknown | unknown | Blood | 1994 | Asia | Southeast Asia | Vietnam | unknown | unknown | Wong et al, 2015 |
| dn61 | ERR349367 | N/A | 4.3.1.1 | unknown | unknown | Blood | 1995 | Asia | Southeast Asia | Vietnam | unknown | unknown | Wong et al, 2015 |
| ct1-7 | ERR349370 | N/A | 3.2.1 | unknown | unknown | Blood | 1993 | Asia | Southeast Asia | Vietnam | unknown | unknown | Wong et al, 2015 |

| | | | | | | | | | | | | | |
|---|---|---|---|---|---|---|---|---|---|---|---|---|---|
| ct1-40 | ERR349372 | N/A | 4.3.1.1 | unknown | unknown | Blood | 1994 | Asia | Southeast Asia | Vietnam | unknown | unknown | Wong et al, 2015 |
| dty1-73 | ERR349373 | N/A | 4.3.1.1 | unknown | unknown | Blood | 1997 | Asia | Southeast Asia | Vietnam | unknown | unknown | Wong et al, 2015 |
| ct1-17 | ERR349375 | N/A | 4.3.1.1 | unknown | unknown | Blood | 1993 | Asia | Southeast Asia | Vietnam | unknown | unknown | Wong et al, 2015 |
| ipt57 | ERR349376 | N/A | 2.3.4 | unknown | unknown | Blood | 1997 | Asia | Southeast Asia | Vietnam | unknown | unknown | Wong et al, 2015 |
| ty3-193 | ERR349378 | N/A | 4.3.1.1 | unknown | unknown | Blood | 1997 | Asia | Southeast Asia | Vietnam | unknown | unknown | Wong et al, 2015 |
| ct1-3 | ERR349381 | N/A | 4.3.1.1 | unknown | unknown | Blood | 1993 | Asia | Southeast Asia | Vietnam | unknown | unknown | Wong et al, 2015 |
| ty3-214 | ERR349384 | N/A | 4.1 | unknown | unknown | Blood | 1997 | Asia | Southeast Asia | Vietnam | unknown | unknown | Wong et al, 2015 |
| ct1-102 | ERR349385 | N/A | 4.3.1.1 | unknown | unknown | Blood | 1994 | Asia | Southeast Asia | Vietnam | unknown | unknown | Wong et al, 2015 |
| ct1-34 | ERR349387 | N/A | 4.3.1.1 | unknown | unknown | Blood | 1994 | Asia | Southeast Asia | Vietnam | unknown | unknown | Wong et al, 2015 |
| ipt41 | ERR349388 | N/A | 4.3.1.1 | unknown | unknown | Blood | 1997 | Asia | Southeast Asia | Vietnam | unknown | unknown | Wong et al, 2015 |
| ty1-35 | ERR349390 | N/A | 3.5 | unknown | unknown | Blood | 1993 | Asia | Southeast Asia | Vietnam | unknown | unknown | Wong et al, 2015 |
| ty2-120 | ERR349392 | N/A | 4.3.1.1 | unknown | unknown | Blood | 1994 | Asia | Southeast Asia | Vietnam | unknown | unknown | Wong et al, 2015 |
| ty2-91 | ERR349397 | N/A | 3.2.1 | unknown | unknown | Blood | 1994 | Asia | Southeast Asia | Vietnam | unknown | unknown | Wong et al, 2015 |
| ty2-107 | ERR349398 | N/A | 2.3.2 | unknown | unknown | Blood | 1994 | Asia | Southeast Asia | Vietnam | unknown | unknown | Wong et al, 2015 |
| ty2-32 | ERR349400 | N/A | 3.2.1 | unknown | unknown | Blood | 1993 | Asia | Southeast Asia | Vietnam | unknown | unknown | Wong et al, 2015 |
| ty2-93 | ERR349402 | N/A | 4.1 | unknown | unknown | Blood | 1994 | Asia | Southeast Asia | Vietnam | unknown | unknown | Wong et al, 2015 |
| ty2-108 | ERR349403 | N/A | 4.3.1.1 | unknown | unknown | Blood | 1994 | Asia | Southeast Asia | Vietnam | unknown | unknown | Wong et al, 2015 |
| ty1-16 | ERR349405 | N/A | 3.1 | unknown | unknown | Blood | 1993 | Asia | Southeast Asia | Vietnam | unknown | unknown | Wong et al, 2015 |
| ty2-80 | ERR349406 | N/A | 3.4 | unknown | unknown | Blood | 1994 | Asia | Southeast Asia | Vietnam | unknown | unknown | Wong et al, 2015 |
| ty2-98 | ERR349411 | N/A | 3.2.1 | unknown | unknown | Blood | 1994 | Asia | Southeast Asia | Vietnam | unknown | unknown | Wong et al, 2015 |
| ty2-111 | ERR349412 | N/A | 2.1.7 | unknown | unknown | Blood | 1994 | Asia | Southeast Asia | Vietnam | unknown | unknown | Wong et al, 2015 |
| ty2-99 | ERR349415 | N/A | 3.2.1 | unknown | unknown | Blood | 1994 | Asia | Southeast Asia | Vietnam | unknown | unknown | Wong et al, 2015 |
| ty2-86 | ERR349423 | N/A | 3.2.1 | unknown | unknown | Blood | 1994 | Asia | Southeast Asia | Vietnam | unknown | unknown | Wong et al, 2015 |
| ty1-34 | ERR349424 | N/A | 2.3.3 | unknown | unknown | Blood | 1993 | Asia | Southeast Asia | Vietnam | unknown | unknown | Wong et al, 2015 |
| ty2-75 | ERR349426 | N/A | 3.2.1 | unknown | unknown | Blood | 1994 | Asia | Southeast Asia | Vietnam | unknown | unknown | Wong et al, 2015 |
| dtc95 | ERR349523 | N/A | 4.3.1.1 | unknown | unknown | Blood | 1994 | Asia | Southeast Asia | Vietnam | unknown | unknown | Wong et al, 2015 |
| dtc110 | ERR349524 | N/A | 4.3.1.1 | unknown | unknown | Blood | 1994 | Asia | Southeast Asia | Vietnam | unknown | unknown | Wong et al, 2015 |
| 3525/3 | ERR349525 | N/A | 2.5.1 | unknown | unknown | Blood | 2011 | Africa | Central Africa | DRC | unknown | unknown | Wong et al, 2015 |
| 3080/3 | ERR349526 | N/A | 2.5.1 | unknown | unknown | Blood | 2010 | Africa | Central Africa | DRC | unknown | unknown | Wong et al, 2015 |
| 3069/3 | ERR349527 | N/A | 2.5.1 | unknown | unknown | Blood | 2010 | Africa | Central Africa | DRC | unknown | unknown | Wong et al, 2015 |
| np61 | ERR349528 | N/A | 4.3.1.2 | unknown | unknown | Blood | 2011 | Asia | South Asia | Nepal | unknown | unknown | Wong et al, 2015 |
| dtc139 | ERR349531 | N/A | 4.3.1.1 | unknown | unknown | Blood | 1995 | Asia | Southeast Asia | Vietnam | unknown | unknown | Wong et al, 2015 |
| dtc102 | ERR349532 | N/A | 4.3.1.1 | unknown | unknown | Blood | 1994 | Asia | Southeast Asia | Vietnam | unknown | unknown | Wong et al, 2015 |
| dn163 | ERR349533 | N/A | 4.3.1.1 | unknown | unknown | Blood | 1996 | Asia | Southeast Asia | Vietnam | unknown | unknown | Wong et al, 2015 |
| dn45 | ERR349534 | N/A | 4.3.1.1 | unknown | unknown | Blood | 1995 | Asia | Southeast Asia | Vietnam | unknown | unknown | Wong et al, 2015 |
| dtc3 | ERR349535 | N/A | 4.3.1.1 | unknown | unknown | Blood | 1994 | Asia | Southeast Asia | Vietnam | unknown | unknown | Wong et al, 2015 |
| dn15 | ERR349536 | N/A | 4.3.1.1 | unknown | unknown | Blood | 1995 | Asia | Southeast Asia | Vietnam | unknown | unknown | Wong et al, 2015 |
| dn110 | ERR349537 | N/A | 3.4 | unknown | unknown | Blood | 1996 | Asia | Southeast Asia | Vietnam | unknown | unknown | Wong et al, 2015 |
| dtc150 | ERR349538 | N/A | 4.3.1.1 | unknown | unknown | Blood | 1995 | Asia | Southeast Asia | Vietnam | unknown | unknown | Wong et al, 2015 |
| dtc92 | ERR349539 | N/A | 4.3.1.1 | unknown | unknown | Blood | 1994 | Asia | Southeast Asia | Vietnam | unknown | unknown | Wong et al, 2015 |
| dn10 | ERR349541 | N/A | 4.3.1.1 | unknown | unknown | Blood | 1995 | Asia | Southeast Asia | Vietnam | unknown | unknown | Wong et al, 2015 |
| dn109 | ERR349543 | N/A | 4.3.1.1 | unknown | unknown | Blood | 1996 | Asia | Southeast Asia | Vietnam | unknown | unknown | Wong et al, 2015 |
| dtc138 | ERR349545 | N/A | 4.3.1.1 | unknown | unknown | Blood | 1995 | Asia | Southeast Asia | Vietnam | unknown | unknown | Wong et al, 2015 |
| dtc69 | ERR349546 | N/A | 4.3.1.1 | unknown | unknown | Blood | 1994 | Asia | Southeast Asia | Vietnam | unknown | unknown | Wong et al, 2015 |
| dn12 | ERR349547 | N/A | 4.3.1.1 | unknown | unknown | Blood | 1995 | Asia | Southeast Asia | Vietnam | unknown | unknown | Wong et al, 2015 |
| dn108 | ERR349548 | N/A | 4.3.1.1 | unknown | unknown | Blood | 1996 | Asia | Southeast Asia | Vietnam | unknown | unknown | Wong et al, 2015 |
| dn57 | ERR349549 | N/A | 4.3.1.1 | unknown | unknown | Blood | 1995 | Asia | Southeast Asia | Vietnam | unknown | unknown | Wong et al, 2015 |
| dtc71 | ERR349550 | N/A | 4.3.1.1 | unknown | unknown | Blood | 1994 | Asia | Southeast Asia | Vietnam | unknown | unknown | Wong et al, 2015 |
| dn20 | ERR349551 | N/A | 4.3.1.1 | unknown | unknown | Blood | 1995 | Asia | Southeast Asia | Vietnam | unknown | unknown | Wong et al, 2015 |
| dtc21 | ERR349552 | N/A | 4.3.1.1 | unknown | unknown | Blood | 1994 | Asia | Southeast Asia | Vietnam | unknown | unknown | Wong et al, 2015 |

| | | | | | | | | | | | | | |
|---|---|---|---|---|---|---|---|---|---|---|---|---|---|
| dn28 | ERR349553 | N/A | 4.3.1.1 | unknown | unknown | Blood | 1995 | Asia | Southeast Asia | Vietnam | unknown | unknown | Wong et al, 2015 |
| ipt43 | ERR349556 | N/A | 4.3.1.1 | unknown | unknown | Blood | 1997 | Asia | Southeast Asia | Vietnam | unknown | unknown | Wong et al, 2015 |
| ty2-160 | ERR349557 | N/A | 4.3.1.1 | unknown | unknown | Blood | 1994 | Asia | Southeast Asia | Vietnam | unknown | unknown | Wong et al, 2015 |
| ipt47 | ERR349558 | N/A | 4.3.1.1 | unknown | unknown | Blood | 1997 | Asia | Southeast Asia | Vietnam | unknown | unknown | Wong et al, 2015 |
| ty2-166 | ERR349559 | N/A | 2.2 | unknown | unknown | Blood | 1994 | Asia | Southeast Asia | Vietnam | unknown | unknown | Wong et al, 2015 |
| ct1-45 | ERR349560 | N/A | 4.3.1.1 | unknown | unknown | Blood | 1994 | Asia | Southeast Asia | Vietnam | unknown | unknown | Wong et al, 2015 |
| dty1-75 | ERR349562 | N/A | 4.3.1.1 | unknown | unknown | Blood | 1997 | Asia | Southeast Asia | Vietnam | unknown | unknown | Wong et al, 2015 |
| ty2-170 | ERR349563 | N/A | 4.3.1.1 | unknown | unknown | Blood | 1995 | Asia | Southeast Asia | Vietnam | unknown | unknown | Wong et al, 2015 |
| ty3-253 | ERR349564 | N/A | 4.3.1.1 | unknown | unknown | Blood | 1998 | Asia | Southeast Asia | Vietnam | unknown | unknown | Wong et al, 2015 |
| as16 | ERR349565 | N/A | 2.3.2 | unknown | unknown | Blood | 1998 | Asia | Southeast Asia | Vietnam | unknown | unknown | Wong et al, 2015 |
| ipt79 | ERR349568 | N/A | 4.3.1.1 | unknown | unknown | Blood | 1997 | Asia | Southeast Asia | Vietnam | unknown | unknown | Wong et al, 2015 |
| dty1-86 | ERR349569 | N/A | 4.3.1.1 | unknown | unknown | Blood | 1997 | Asia | Southeast Asia | Vietnam | unknown | unknown | Wong et al, 2015 |
| ct1-23 | ERR349570 | N/A | 4.3.1.1 | unknown | unknown | Blood | 1993 | Asia | Southeast Asia | Vietnam | unknown | unknown | Wong et al, 2015 |
| ipt76 | ERR349571 | N/A | 4.3.1.1 | unknown | unknown | Blood | 1997 | Asia | Southeast Asia | Vietnam | unknown | unknown | Wong et al, 2015 |
| dty1-88 | ERR349572 | N/A | 4.3.1.1 | unknown | unknown | Blood | 1997 | Asia | Southeast Asia | Vietnam | unknown | unknown | Wong et al, 2015 |
| ty3-212 | ERR349573 | N/A | 4.3.1.1 | unknown | unknown | Blood | 1997 | Asia | Southeast Asia | Vietnam | unknown | unknown | Wong et al, 2015 |
| as257 | ERR349575 | N/A | 3.2.1 | unknown | unknown | Blood | 1999 | Asia | Southeast Asia | Vietnam | unknown | unknown | Wong et al, 2015 |
| dty1-89 | ERR349576 | N/A | 4.3.1.1 | unknown | unknown | Blood | 1997 | Asia | Southeast Asia | Vietnam | unknown | unknown | Wong et al, 2015 |
| ct1-65 | ERR349578 | N/A | 4.3.1.1 | unknown | unknown | Blood | 1994 | Asia | Southeast Asia | Vietnam | unknown | unknown | Wong et al, 2015 |
| dty1-46 | ERR349579 | N/A | 4.3.1.1 | unknown | unknown | Blood | 1995 | Asia | Southeast Asia | Vietnam | unknown | unknown | Wong et al, 2015 |
| dty1-91 | ERR349580 | N/A | 4.3.1.1 | unknown | unknown | Blood | 1997 | Asia | Southeast Asia | Vietnam | unknown | unknown | Wong et al, 2015 |
| dty1-58 | ERR349581 | N/A | 4.3.1.1 | unknown | unknown | Blood | 1995 | Asia | Southeast Asia | Vietnam | unknown | unknown | Wong et al, 2015 |
| dty1-93 | ERR349582 | N/A | 4.3.1.1 | unknown | unknown | Blood | 1997 | Asia | Southeast Asia | Vietnam | unknown | unknown | Wong et al, 2015 |
| ct1-33 | ERR349583 | N/A | 4.3.1.1 | unknown | unknown | Blood | 1994 | Asia | Southeast Asia | Vietnam | unknown | unknown | Wong et al, 2015 |
| dty1-67 | ERR349584 | N/A | 4.3.1.1 | unknown | unknown | Blood | 1995 | Asia | Southeast Asia | Vietnam | unknown | unknown | Wong et al, 2015 |
| ty3-215 | ERR349585 | N/A | 4.3.1.1 | unknown | unknown | Blood | 1997 | Asia | Southeast Asia | Vietnam | unknown | unknown | Wong et al, 2015 |
| ty2-76 | ERR349588 | N/A | 4.3.1.1 | unknown | unknown | Blood | 1994 | Asia | Southeast Asia | Vietnam | unknown | unknown | Wong et al, 2015 |
| ty2-17 | ERR349590 | N/A | 4.3.1.1 | unknown | unknown | Blood | 1993 | Asia | Southeast Asia | Vietnam | unknown | unknown | Wong et al, 2015 |
| ty2-90 | ERR349591 | N/A | 4.3.1.1 | unknown | unknown | Blood | 1994 | Asia | Southeast Asia | Vietnam | unknown | unknown | Wong et al, 2015 |
| ty2-139 | ERR349592 | N/A | 4.3.1.1 | unknown | unknown | Blood | 1994 | Asia | Southeast Asia | Vietnam | unknown | unknown | Wong et al, 2015 |
| ty2-123 | ERR349593 | N/A | 4.3.1.1 | unknown | unknown | Blood | 1994 | Asia | Southeast Asia | Vietnam | unknown | unknown | Wong et al, 2015 |
| ty2-144 | ERR349595 | N/A | 4.3.1.1 | unknown | unknown | Blood | 1994 | Asia | Southeast Asia | Vietnam | unknown | unknown | Wong et al, 2015 |
| ty2-146 | ERR349597 | N/A | 4.3.1.1 | unknown | unknown | Blood | 1994 | Asia | Southeast Asia | Vietnam | unknown | unknown | Wong et al, 2015 |
| ty2-100 | ERR349599 | N/A | 4.3.1.1 | unknown | unknown | Blood | 1994 | Asia | Southeast Asia | Vietnam | unknown | unknown | Wong et al, 2015 |
| ty2-114 | ERR349600 | N/A | 4.3.1.1 | unknown | unknown | Blood | 1994 | Asia | Southeast Asia | Vietnam | unknown | unknown | Wong et al, 2015 |
| ty2-131 | ERR349601 | N/A | 4.3.1.1 | unknown | unknown | Blood | 1994 | Asia | Southeast Asia | Vietnam | unknown | unknown | Wong et al, 2015 |
| ty2-71 | ERR349602 | N/A | 4.3.1.1 | unknown | unknown | Blood | 1994 | Asia | Southeast Asia | Vietnam | unknown | unknown | Wong et al, 2015 |
| ty2-101 | ERR349603 | N/A | 4.3.1.1 | unknown | unknown | Blood | 1994 | Asia | Southeast Asia | Vietnam | unknown | unknown | Wong et al, 2015 |
| ty2-115 | ERR349604 | N/A | 4.1 | unknown | unknown | Blood | 1994 | Asia | Southeast Asia | Vietnam | unknown | unknown | Wong et al, 2015 |
| ty2-132 | ERR349605 | N/A | 4.3.1.1 | unknown | unknown | Blood | 1994 | Asia | Southeast Asia | Vietnam | unknown | unknown | Wong et al, 2015 |
| ty2-150 | ERR349606 | N/A | 4.3.1.1 | unknown | unknown | Blood | 1994 | Asia | Southeast Asia | Vietnam | unknown | unknown | Wong et al, 2015 |
| ty2-153 | ERR349609 | N/A | 4.3.1.1 | unknown | unknown | Blood | 1994 | Asia | Southeast Asia | Vietnam | unknown | unknown | Wong et al, 2015 |
| ty2-103 | ERR349613 | N/A | 4.3.1.1 | unknown | unknown | Blood | 1994 | Asia | Southeast Asia | Vietnam | unknown | unknown | Wong et al, 2015 |
| ty2-134 | ERR349615 | N/A | 4.3.1.1 | unknown | unknown | Blood | 1994 | Asia | Southeast Asia | Vietnam | unknown | unknown | Wong et al, 2015 |
| ty2-104 | ERR349616 | N/A | 4.3.1.1 | unknown | unknown | Blood | 1994 | Asia | Southeast Asia | Vietnam | unknown | unknown | Wong et al, 2015 |
| ty2-155 | ERR349617 | N/A | 4.3.1.1 | unknown | unknown | Blood | 1994 | Asia | Southeast Asia | Vietnam | unknown | unknown | Wong et al, 2015 |
| UI 3257 | ERR352253 | N/A | 3.2.1 | unknown | unknown | Blood | 2003 | Asia | Southeast Asia | Laos | unknown | unknown | Wong et al, 2015 |
| MDUST106 | ERR352254 | N/A | 3.1 | unknown | unknown | Blood | 2006 | Asia | Western Asia | Lebanon | unknown | unknown | Wong et al, 2015 |
| MDUST115 | ERR352255 | N/A | 2.1.7.1 | unknown | unknown | Stool | 2009 | Australia & Oceania | Oceania | Papua New Guinea | unknown | unknown | Wong et al, 2015 |
| MDUST120 | ERR352256 | N/A | 3.1 | unknown | unknown | Stool | 2009 | Europe | Southern Europe | Malta | unknown | unknown | Wong et al, 2015 |

| | | | | | | | | | | | | | |
|---|---|---|---|---|---|---|---|---|---|---|---|---|---|
| MDUST121 | ERR352257 | N/A | 2.2.1 | unknown | unknown | Blood | 2010 | Australia & Oceania | Australia | Australia | unknown | unknown | Wong et al, 2015 |
| MDUST123 | ERR352258 | N/A | 3.5.4 | unknown | unknown | Blood | 2010 | Australia & Oceania | Oceania | Samoa | unknown | unknown | Wong et al, 2015 |
| MDUST127 | ERR352259 | N/A | 4.3.1.2 | unknown | unknown | Blood | 2011 | Asia | South Asia | India | unknown | unknown | Wong et al, 2015 |
| MDUST130 | ERR352260 | N/A | 3 | unknown | unknown | Stool | 2011 | Asia | Southeast Asia | Malaysia | unknown | unknown | Wong et al, 2015 |
| MDUST133 | ERR352261 | N/A | 3.5.4 | unknown | unknown | Blood | 2011 | Australia & Oceania | Oceania | Samoa | unknown | unknown | Wong et al, 2015 |
| MDUST135 | ERR352262 | N/A | 2.3.5 | unknown | unknown | Blood | 2011 | Australia & Oceania | Oceania | Tonga | unknown | unknown | Wong et al, 2015 |
| MDUST139 | ERR352263 | N/A | 3.3.1 | unknown | unknown | Stool | 2012 | Asia | Southeast Asia | Myanmar | unknown | unknown | Wong et al, 2015 |
| MDUST141 | ERR352264 | N/A | 4.3.1.1 | unknown | unknown | Blood | 2012 | Asia | South Asia | India | unknown | unknown | Wong et al, 2015 |
| MDUST145 | ERR352265 | N/A | 4.3.1.2 | unknown | unknown | Blood | 2012 | Asia | South Asia | India | unknown | unknown | Wong et al, 2015 |
| MDUST147 | ERR352266 | N/A | 4.3.1.2 | unknown | unknown | Blood | 2012 | Asia | South Asia | India | unknown | unknown | Wong et al, 2015 |
| MDUST149 | ERR352267 | N/A | 3.0.2 | unknown | unknown | Blood | 2012 | Asia | South Asia | India | unknown | unknown | Wong et al, 2015 |
| MDUST151 | ERR352268 | N/A | 4.3.1.2 | unknown | unknown | Blood | 2012 | Unknown | Unknown | Unknown | unknown | unknown | Wong et al, 2015 |
| MDUST154 | ERR352269 | N/A | 2.3.4 | unknown | unknown | Stool | 2011 | Asia | East Asia | China | unknown | unknown | Wong et al, 2015 |
| MDUST158 | ERR352270 | N/A | 4.3.1.2 | unknown | unknown | Blood | 2012 | Asia | South Asia | India | unknown | unknown | Wong et al, 2015 |
| MDUST161 | ERR352271 | N/A | 4.3.1.2 | unknown | unknown | Blood | 2011 | Asia | South Asia | India | unknown | unknown | Wong et al, 2015 |
| MDUST166 | ERR352272 | N/A | 4.3.1.1 | unknown | unknown | Blood | 2011 | Asia | South Asia | India | unknown | unknown | Wong et al, 2015 |
| MDUST169 | ERR352273 | N/A | 4.3.1.2 | unknown | unknown | Blood | 2011 | Asia | South Asia | India | unknown | unknown | Wong et al, 2015 |
| MDUST182 | ERR352274 | N/A | 4.3.1.1 | unknown | unknown | Blood | 2011 | Asia | South Asia | India | unknown | unknown | Wong et al, 2015 |
| MDUST185 | ERR352275 | N/A | 4.3.1 | unknown | unknown | Blood | 2011 | Asia | South Asia | India | unknown | unknown | Wong et al, 2015 |
| MDUST187 | ERR352276 | N/A | 4.3.1.2 | unknown | unknown | Blood | 2011 | Unknown | Unknown | Unknown | unknown | unknown | Wong et al, 2015 |
| MDUST196 | ERR352277 | N/A | 4.3.1.3.Bdq | unknown | unknown | Blood | 2011 | Asia | South Asia | Bangladesh | unknown | unknown | Wong et al, 2015 |
| MDUST197 | ERR352278 | N/A | 4.3.1.2 | unknown | unknown | Blood | 2010 | Asia | South Asia | India | unknown | unknown | Wong et al, 2015 |
| MDUST198 | ERR352279 | N/A | 4.3.1.1 | unknown | unknown | Blood | 2010 | Asia | Southeast Asia | Myanmar | unknown | unknown | Wong et al, 2015 |
| MDUST199 | ERR352280 | N/A | 4.3.1.1 | unknown | unknown | Blood | 2011 | Asia | South Asia | India | unknown | unknown | Wong et al, 2015 |
| MDUST200 | ERR352281 | N/A | 4.3.1 | unknown | unknown | Blood | 2011 | Asia | South Asia | India | unknown | unknown | Wong et al, 2015 |
| MDUST206 | ERR352282 | N/A | 3 | unknown | unknown | Blood | 2010 | Asia | Southeast Asia | Indonesia | unknown | unknown | Wong et al, 2015 |
| MDUST211 | ERR352283 | N/A | 4.3.1.2 | unknown | unknown | Blood | 2011 | Asia | South Asia | India | unknown | unknown | Wong et al, 2015 |
| MDUST216 | ERR352284 | N/A | 4.3.1.2 | unknown | unknown | Blood | 2011 | Asia | South Asia | India | unknown | unknown | Wong et al, 2015 |
| MDUST217 | ERR352285 | N/A | 3.2.1 | unknown | unknown | Blood | 2011 | Asia | South Asia | India | unknown | unknown | Wong et al, 2015 |
| MDUST223 | ERR352286 | N/A | 4.3.1.2 | unknown | unknown | Blood | 2011 | Unknown | Unknown | Unknown | unknown | unknown | Wong et al, 2015 |
| MDUST226 | ERR352287 | N/A | 2.1.7.1 | unknown | unknown | Blood | 2010 | Australia & Oceania | Oceania | Papua New Guinea | unknown | unknown | Wong et al, 2015 |
| MDUST229 | ERR352288 | N/A | 2.1.7.1 | unknown | unknown | Blood | 2010 | Australia & Oceania | Oceania | Papua New Guinea | unknown | unknown | Wong et al, 2015 |
| MDUST231 | ERR352289 | N/A | 4.3.1.2 | unknown | unknown | Blood | 2010 | Asia | Western Asia | Lebanon | unknown | unknown | Wong et al, 2015 |
| MDUST234 | ERR352290 | N/A | 3.3.2.Bd2 | unknown | unknown | Blood | 2010 | Asia | South Asia | Bangladesh | unknown | unknown | Wong et al, 2015 |
| MDUST237 | ERR352291 | N/A | 4.3.1.2 | unknown | unknown | Stool | 2010 | Asia | South Asia | Nepal | unknown | unknown | Wong et al, 2015 |
| MDUST239 | ERR352292 | N/A | 3.5.4 | unknown | unknown | Blood | 2011 | Unknown | Unknown | Unknown | unknown | unknown | Wong et al, 2015 |
| MDUST241 | ERR352293 | N/A | 4.3.1.3.Bdq | unknown | unknown | Blood | 2010 | Asia | South Asia | Bangladesh | unknown | unknown | Wong et al, 2015 |
| MDUST242 | ERR352294 | N/A | 4.3.1.1 | unknown | unknown | Blood | 2011 | Asia | South Asia | India | unknown | unknown | Wong et al, 2015 |
| MDUST243 | ERR352295 | N/A | 4.3.1.2 | unknown | unknown | Blood | 2011 | Asia | South Asia | India | unknown | unknown | Wong et al, 2015 |
| MDUST248 | ERR352296 | N/A | 4.3.1.2 | unknown | unknown | Blood | 2011 | Asia | South Asia | India | unknown | unknown | Wong et al, 2015 |
| MDUST249 | ERR352297 | N/A | 4.3.1.2 | unknown | unknown | Blood | 2011 | Asia | South Asia | India | unknown | unknown | Wong et al, 2015 |
| MDUST252 | ERR352298 | N/A | 3.3.2.Bd1 | unknown | unknown | Blood | 2010 | Asia | South Asia | Bangladesh | unknown | unknown | Wong et al, 2015 |
| MDUST253 | ERR352299 | N/A | 3.1.2 | unknown | unknown | Blood | 2010 | Asia | Southeast Asia | Indonesia | unknown | unknown | Wong et al, 2015 |
| MDUST255 | ERR352300 | N/A | 2.1.7.1 | unknown | unknown | Blood | 2010 | Australia & Oceania | Oceania | Papua New Guinea | unknown | unknown | Wong et al, 2015 |
| MDUST256 | ERR352301 | N/A | 2.1.7.1 | unknown | unknown | Blood | 2010 | Australia & Oceania | Oceania | Papua New Guinea | unknown | unknown | Wong et al, 2015 |
| MDUST257 | ERR352302 | N/A | 2.1.5 | unknown | unknown | Blood | 2010 | Asia | Southeast Asia | Indonesia | unknown | unknown | Wong et al, 2015 |
| MDUST258 | ERR352303 | N/A | 3.2.2 | unknown | unknown | Stool | 2010 | Asia | South Asia | Bangladesh | unknown | unknown | Wong et al, 2015 |
| MDUST259 | ERR352304 | N/A | 4.3.1.1 | unknown | unknown | Blood | 2010 | Asia | South Asia | Bangladesh | unknown | unknown | Wong et al, 2015 |
| MDUST265 | ERR352305 | N/A | 4.3.1.3.Bdq | unknown | unknown | Blood | 2011 | Unknown | Unknown | Unknown | unknown | unknown | Wong et al, 2015 |
| MDUST267 | ERR352306 | N/A | 2.5 | unknown | unknown | Blood | 2011 | Asia | South Asia | India | unknown | unknown | Wong et al, 2015 |

| | | | | | | | | | | | | | |
|---|---|---|---|---|---|---|---|---|---|---|---|---|---|
| MDUST269 | ERR352307 | N/A | 4.3.1.1 | unknown | unknown | Blood | 2011 | Asia | South Asia | Pakistan | unknown | unknown | Wong et al, 2015 |
| MDUST270 | ERR352308 | N/A | 4.3.1.1 | unknown | unknown | Stool | 2011 | Asia | South Asia | Bangladesh | unknown | unknown | Wong et al, 2015 |
| MDUST274 | ERR352309 | N/A | 2.1.7.2 | unknown | unknown | Blood | 1996 | Australia & Oceania | Oceania | Papua New Guinea | unknown | unknown | Wong et al, 2015 |
| MDUST295 | ERR352310 | N/A | 2.1 | unknown | unknown | Blood | 2010 | Asia | Southeast Asia | Indonesia | unknown | unknown | Wong et al, 2015 |
| MDUST310 | ERR352311 | N/A | 3.5 | unknown | unknown | Stool | 1991 | Australia & Oceania | Oceania | Fiji | unknown | unknown | Wong et al, 2015 |
| MDUST319 | ERR352312 | N/A | 4.2.1 | unknown | unknown | Blood | 1983 | Australia & Oceania | Oceania | Fiji | unknown | unknown | Wong et al, 2015 |
| MDUST326 | ERR352313 | N/A | 4.3.1.1 | unknown | unknown | Stool | 1992 | Australia & Oceania | Oceania | Fiji | unknown | unknown | Wong et al, 2015 |
| MDUST328 | ERR352314 | N/A | 2.1.7.1 | unknown | unknown | Blood | 1992 | Australia & Oceania | Oceania | Papua New Guinea | unknown | unknown | Wong et al, 2015 |
| MDUST335 | ERR352315 | N/A | 4.3.1.1 | unknown | unknown | Stool | 1993 | Australia & Oceania | Oceania | Fiji | unknown | unknown | Wong et al, 2015 |
| MDUST337 | ERR352316 | N/A | 4.2.1 | unknown | unknown | Blood | 1984 | Australia & Oceania | Oceania | Fiji | unknown | unknown | Wong et al, 2015 |
| MDUST354 | ERR352317 | N/A | 4.2.3 | unknown | unknown | Blood | 1993 | Australia & Oceania | Oceania | Fiji | unknown | unknown | Wong et al, 2015 |
| MDUST355 | ERR352318 | N/A | 2.1.7 | unknown | unknown | Blood | 1994 | Australia & Oceania | Oceania | Papua New Guinea | unknown | unknown | Wong et al, 2015 |
| MDUST361 | ERR352319 | N/A | 2.1.7.1 | unknown | unknown | Stool | 1998 | Australia & Oceania | Oceania | Papua New Guinea | unknown | unknown | Wong et al, 2015 |
| MDUST362 | ERR352320 | N/A | 2.1.7.1 | unknown | unknown | Blood | 1998 | Australia & Oceania | Oceania | Papua New Guinea | unknown | unknown | Wong et al, 2015 |
| MDUST363 | ERR352321 | N/A | 2.1.7.1 | unknown | unknown | Blood | 1998 | Australia & Oceania | Oceania | Papua New Guinea | unknown | unknown | Wong et al, 2015 |
| MDUST364 | ERR352322 | N/A | 2.1.7.1 | unknown | unknown | Stool | 1999 | Australia & Oceania | Oceania | Papua New Guinea | unknown | unknown | Wong et al, 2015 |
| MDUST366 | ERR352323 | N/A | 3.3.1 | unknown | unknown | Blood | 2011 | Asia | Western Asia | Iraq | unknown | unknown | Wong et al, 2015 |
| MDUST382 | ERR352324 | N/A | 3.3.1 | unknown | unknown | Stool | 2011 | Asia | South Asia | India | unknown | unknown | Wong et al, 2015 |
| MDUST385 | ERR352325 | N/A | 4.3.1.2 | unknown | unknown | Blood | 2011 | Asia | South Asia | India | unknown | unknown | Wong et al, 2015 |
| MDUST386 | ERR352326 | N/A | 4.3.1.1 | unknown | unknown | Blood | 2011 | Asia | Western Asia | Iraq | unknown | unknown | Wong et al, 2015 |
| MDUST387 | ERR352327 | N/A | 4.3.1 | unknown | unknown | Blood | 2011 | Asia | South Asia | India | unknown | unknown | Wong et al, 2015 |
| MDUST391 | ERR352328 | N/A | 2.2 | unknown | unknown | Blood | 2011 | Asia | South Asia | India | unknown | unknown | Wong et al, 2015 |
| MDUST393 | ERR352329 | N/A | 4.3.1.2 | unknown | unknown | Stool | 2011 | Asia | South Asia | India | unknown | unknown | Wong et al, 2015 |
| MDUST396 | ERR352330 | N/A | 2.3.2 | unknown | unknown | Stool | 2012 | South America | South America | El Salvador | unknown | unknown | Wong et al, 2015 |
| MDUST397 | ERR352331 | N/A | 3 | unknown | unknown | Blood | 2011 | Asia | Southeast Asia | Indonesia | unknown | unknown | Wong et al, 2015 |
| MDUST403 | ERR352332 | N/A | 4.3.1.1 | unknown | unknown | Blood | 2012 | Asia | Southeast Asia | Cambodia | unknown | unknown | Wong et al, 2015 |
| MDUST404 | ERR352333 | N/A | 4.2.2 | unknown | unknown | Blood | 2012 | Australia & Oceania | Oceania | Fiji | unknown | unknown | Wong et al, 2015 |
| MDUST406 | ERR352334 | N/A | 3.5.4 | unknown | unknown | Blood | 2012 | Australia & Oceania | Oceania | Samoa | unknown | unknown | Wong et al, 2015 |
| MDUST408 | ERR352335 | N/A | 4.3.1.2 | unknown | unknown | Blood | 2012 | Asia | South Asia | India | unknown | unknown | Wong et al, 2015 |
| MDUST410 | ERR352336 | N/A | 4.3.1 | unknown | unknown | Blood | 2011 | Asia | South Asia | India | unknown | unknown | Wong et al, 2015 |
| MDUST411 | ERR352337 | N/A | 4.1 | unknown | unknown | Blood | 2011 | Unknown | Unknown | Unknown | unknown | unknown | Wong et al, 2015 |
| MDUST412 | ERR352338 | N/A | 4.3.1.2 | unknown | unknown | Stool | 2011 | Asia | South Asia | Nepal | unknown | unknown | Wong et al, 2015 |
| MDUST418 | ERR352339 | N/A | 4.3.1 | unknown | unknown | Blood | 2011 | Asia | South Asia | Pakistan | unknown | unknown | Wong et al, 2015 |
| MDUST108 | ERR352426 | N/A | 3.1 | unknown | unknown | Stool | 2007 | Africa | West Africa | Liberia | unknown | unknown | Wong et al, 2015 |
| MDUST109 | ERR352427 | N/A | 2.1.7.1 | unknown | unknown | Blood | 2007 | Australia & Oceania | Oceania | Papua New Guinea | unknown | unknown | Wong et al, 2015 |
| MDUST113 | ERR352428 | N/A | 2.2.1 | unknown | unknown | Blood | 2009 | Asia | Western Asia | Lebanon | unknown | unknown | Wong et al, 2015 |
| MDUST116 | ERR352429 | N/A | 3.5.4 | unknown | unknown | Blood | 2009 | Australia & Oceania | Oceania | Samoa | unknown | unknown | Wong et al, 2015 |
| MDUST117 | ERR352430 | N/A | 3.5.4 | unknown | unknown | Blood | 2009 | Australia & Oceania | Oceania | Samoa | unknown | unknown | Wong et al, 2015 |
| MDUST118 | ERR352431 | N/A | 3.5.4 | unknown | unknown | Blood | 2009 | Australia & Oceania | Oceania | Samoa | unknown | unknown | Wong et al, 2015 |
| MDUST125 | ERR352432 | N/A | 3.5.4 | unknown | unknown | Blood | 2011 | Australia & Oceania | Oceania | Samoa | unknown | unknown | Wong et al, 2015 |
| MDUST126 | ERR352433 | N/A | 2.2.1 | unknown | unknown | Blood | 2010 | Asia | Southeast Asia | Malaysia | unknown | unknown | Wong et al, 2015 |
| MDUST128 | ERR352434 | N/A | 4.3.1.2 | unknown | unknown | Blood | 2011 | Asia | South Asia | India | unknown | unknown | Wong et al, 2015 |
| MDUST129 | ERR352435 | N/A | 3 | unknown | unknown | Blood | 2011 | Asia | Southeast Asia | Phillipines | unknown | unknown | Wong et al, 2015 |
| MDUST131 | ERR352436 | N/A | 3.3 | unknown | unknown | Blood | 2011 | Unknown | Unknown | Unknown | unknown | unknown | Wong et al, 2015 |
| MDUST132 | ERR352437 | N/A | 3.5.4 | unknown | unknown | Blood | 2011 | Australia & Oceania | Oceania | Samoa | unknown | unknown | Wong et al, 2015 |
| MDUST136 | ERR352438 | N/A | 3.3.1 | unknown | unknown | Blood | 2011 | Asia | South Asia | India | unknown | unknown | Wong et al, 2015 |
| MDUST140 | ERR352439 | N/A | 4.3.1.2 | unknown | unknown | Blood | 2011 | Asia | South Asia | India | unknown | unknown | Wong et al, 2015 |
| MDUST143 | ERR352440 | N/A | 2.2.1 | unknown | unknown | Stool | 2012 | Asia | South Asia | India | unknown | unknown | Wong et al, 2015 |
| MDUST150 | ERR352441 | N/A | 3 | unknown | unknown | Blood | 2012 | Asia | Southeast Asia | Indonesia | unknown | unknown | Wong et al, 2015 |
| MDUST152 | ERR352442 | N/A | 4.3.1.1.EA1 | unknown | unknown | Blood | 2012 | Africa | Africa | Africa | unknown | unknown | Wong et al, 2015 |

| | | | | | | | | | | | | | |
|---|---|---|---|---|---|---|---|---|---|---|---|---|---|
| MDUST153 | ERR352443 | N/A | 4.3.1 | unknown | unknown | Blood | 2012 | Asia | South Asia | India | unknown | unknown | Wong et al, 2015 |
| MDUST156 | ERR352444 | N/A | 4.3.1.1 | unknown | unknown | Blood | 2012 | Asia | South Asia | Pakistan | unknown | unknown | Wong et al, 2015 |
| MDUST157 | ERR352445 | N/A | 4.3.1.2 | unknown | unknown | Stool | 2012 | Asia | South Asia | Nepal | unknown | unknown | Wong et al, 2015 |
| MDUST159 | ERR352446 | N/A | 4.3.1 | unknown | unknown | Blood | 2012 | Asia | South Asia | India | unknown | unknown | Wong et al, 2015 |
| MDUST162 | ERR352447 | N/A | 4.3.1.2 | unknown | unknown | Blood | 2011 | Unknown | Unknown | Unknown | unknown | unknown | Wong et al, 2015 |
| MDUST163 | ERR352448 | N/A | 4.3.1.2 | unknown | unknown | Blood | 2011 | Asia | South Asia | India | unknown | unknown | Wong et al, 2015 |
| MDUST168 | ERR352449 | N/A | 4.3.1 | unknown | unknown | Blood | 2011 | Asia | South Asia | India | unknown | unknown | Wong et al, 2015 |
| MDUST174 | ERR352451 | N/A | 4.3.1 | unknown | unknown | Blood | 2011 | Asia | South Asia | Sri Lanka | unknown | unknown | Wong et al, 2015 |
| MDUST175 | ERR352452 | N/A | 4.3.1 | unknown | unknown | Blood | 2011 | Asia | South Asia | Pakistan | unknown | unknown | Wong et al, 2015 |
| MDUST177 | ERR352453 | N/A | 1.1.4 | unknown | unknown | Urine | 2011 | Unknown | Unknown | Unknown | unknown | unknown | Wong et al, 2015 |
| MDUST179 | ERR352454 | N/A | 3.2.2 | unknown | unknown | Blood | 2011 | Asia | South Asia | India | unknown | unknown | Wong et al, 2015 |
| MDUST183 | ERR352455 | N/A | 4.3.1.2 | unknown | unknown | Blood | 2011 | Asia | South Asia | India | unknown | unknown | Wong et al, 2015 |
| MDUST184 | ERR352456 | N/A | 4.3.1.1 | unknown | unknown | Blood | 2011 | Asia | South Asia | India | unknown | unknown | Wong et al, 2015 |
| MDUST194 | ERR352457 | N/A | 3.2.2 | unknown | unknown | Blood | 2011 | Asia | South Asia | India | unknown | unknown | Wong et al, 2015 |
| MDUST201 | ERR352458 | N/A | 4.3.1.2 | unknown | unknown | Blood | 2011 | Asia | South Asia | India | unknown | unknown | Wong et al, 2015 |
| MDUST202 | ERR352459 | N/A | 2 | unknown | unknown | Blood | 2010 | Asia | South Asia | Bangladesh | unknown | unknown | Wong et al, 2015 |
| MDUST203 | ERR352460 | N/A | 2 | unknown | unknown | Blood | 2010 | Asia | South Asia | Bangladesh | unknown | unknown | Wong et al, 2015 |
| MDUST204 | ERR352461 | N/A | 2 | unknown | unknown | Blood | 2010 | Asia | South Asia | Bangladesh | unknown | unknown | Wong et al, 2015 |
| MDUST205 | ERR352462 | N/A | 2 | unknown | unknown | Blood | 2010 | Asia | South Asia | Bangladesh | unknown | unknown | Wong et al, 2015 |
| MDUST207 | ERR352463 | N/A | 4.3.1.1 | unknown | unknown | Blood | 2010 | Asia | South Asia | Pakistan | unknown | unknown | Wong et al, 2015 |
| MDUST214 | ERR352464 | N/A | 4.3.1.2 | unknown | unknown | Blood | 2011 | Asia | South Asia | India | unknown | unknown | Wong et al, 2015 |
| MDUST215 | ERR352465 | N/A | 3 | unknown | unknown | Stool | 2011 | Asia | South Asia | India | unknown | unknown | Wong et al, 2015 |
| MDUST220 | ERR352466 | N/A | 4.3.1.1 | unknown | unknown | Blood | 2011 | Asia | South Asia | India | unknown | unknown | Wong et al, 2015 |
| MDUST222 | ERR352467 | N/A | 2.1.5 | unknown | unknown | Blood | 2011 | Asia | Southeast Asia | Indonesia | unknown | unknown | Wong et al, 2015 |
| MDUST232 | ERR352468 | N/A | 4.3.1.1 | unknown | unknown | Blood | 2010 | Asia | South Asia | Bangladesh | unknown | unknown | Wong et al, 2015 |
| MDUST235 | ERR352469 | N/A | 3.2.1 | unknown | unknown | Stool | 2010 | Asia | Southeast Asia | Vietnam | unknown | unknown | Wong et al, 2015 |
| MDUST236 | ERR352470 | N/A | 4.3.1.1 | unknown | unknown | Blood | 2010 | Asia | Southeast Asia | Thailand | unknown | unknown | Wong et al, 2015 |
| MDUST244 | ERR352471 | N/A | 4.3.1.2 | unknown | unknown | Blood | 2011 | Asia | South Asia | India | unknown | unknown | Wong et al, 2015 |
| MDUST246 | ERR352472 | N/A | 2.1 | unknown | unknown | Blood | 2011 | Asia | Southeast Asia | Indonesia | unknown | unknown | Wong et al, 2015 |
| MDUST250 | ERR352473 | N/A | 0.0.2 | unknown | unknown | Blood | 2010 | Asia | Southeast Asia | Indonesia | unknown | unknown | Wong et al, 2015 |
| MDUST261 | ERR352474 | N/A | 4.3.1 | unknown | unknown | Blood | 2011 | Unknown | Unknown | Unknown | unknown | unknown | Wong et al, 2015 |
| MDUST268 | ERR352475 | N/A | 2.5 | unknown | unknown | Blood | 2011 | Unknown | Unknown | Unknown | unknown | unknown | Wong et al, 2015 |
| MDUST276 | ERR352476 | N/A | 2.1.7.2 | unknown | unknown | Stool | 1994 | Australia & Oceania | Oceania | Papua New Guinea | unknown | unknown | Wong et al, 2015 |
| MDUST278 | ERR352477 | N/A | 2.1.7.1 | unknown | unknown | Blood | 2001 | Australia & Oceania | Oceania | Papua New Guinea | unknown | unknown | Wong et al, 2015 |
| MDUST279 | ERR352478 | N/A | 2.1.7.1 | unknown | unknown | Blood | 2002 | Australia & Oceania | Oceania | Papua New Guinea | unknown | unknown | Wong et al, 2015 |
| MDUST282 | ERR352479 | N/A | 3.5.4 | unknown | unknown | Blood | 2004 | Australia & Oceania | Oceania | Samoa | unknown | unknown | Wong et al, 2015 |
| MDUST283 | ERR352480 | N/A | 3.5.4 | unknown | unknown | Blood | 2004 | Australia & Oceania | Oceania | Samoa | unknown | unknown | Wong et al, 2015 |
| MDUST286 | ERR352481 | N/A | 3.5.4 | unknown | unknown | Blood | 2005 | Australia & Oceania | Oceania | Samoa | unknown | unknown | Wong et al, 2015 |
| MDUST287 | ERR352482 | N/A | 2.1.7.1 | unknown | unknown | Blood | 2007 | Australia & Oceania | Oceania | Papua New Guinea | unknown | unknown | Wong et al, 2015 |
| MDUST288 | ERR352483 | N/A | 2.1.7.1 | unknown | unknown | Blood | 2007 | Australia & Oceania | Oceania | Papua New Guinea | unknown | unknown | Wong et al, 2015 |
| MDUST289 | ERR352484 | N/A | 3.5.4 | unknown | unknown | Blood | 2008 | Australia & Oceania | Oceania | Samoa | unknown | unknown | Wong et al, 2015 |
| MDUST291 | ERR352485 | N/A | 3.5.4 | unknown | unknown | Stool | 2008 | Australia & Oceania | Oceania | Samoa | unknown | unknown | Wong et al, 2015 |
| MDUST292 | ERR352486 | N/A | 3.5.4 | unknown | unknown | Blood | 2008 | Australia & Oceania | Oceania | Samoa | unknown | unknown | Wong et al, 2015 |
| MDUST296 | ERR352487 | N/A | 2.1.7.1 | unknown | unknown | Blood | 2010 | Australia & Oceania | Oceania | Papua New Guinea | unknown | unknown | Wong et al, 2015 |
| MDUST297 | ERR352488 | N/A | 3.5.4 | unknown | unknown | Blood | 2011 | Unknown | Unknown | Unknown | unknown | unknown | Wong et al, 2015 |
| MDUST298 | ERR352489 | N/A | 2.1.7.2 | unknown | unknown | Blood | 1998 | Australia & Oceania | Oceania | Papua New Guinea | unknown | unknown | Wong et al, 2015 |
| MDUST299 | ERR352490 | N/A | 2.1.7.2 | unknown | unknown | Blood | 1998 | Australia & Oceania | Oceania | Papua New Guinea | unknown | unknown | Wong et al, 2015 |
| MDUST300 | ERR352491 | N/A | 2.1.7.1 | unknown | unknown | Stool | 1998 | Australia & Oceania | Oceania | Papua New Guinea | unknown | unknown | Wong et al, 2015 |
| MDUST301 | ERR352492 | N/A | 2.1.7.1 | unknown | unknown | Stool | 1988 | Asia | South Asia | India | unknown | unknown | Wong et al, 2015 |
| MDUST313 | ERR352493 | N/A | 2.1.7.1 | unknown | unknown | Blood | 1994 | Australia & Oceania | Oceania | Papua New Guinea | unknown | unknown | Wong et al, 2015 |

| | | | | | | | | | | | | | |
|---|---|---|---|---|---|---|---|---|---|---|---|---|---|
| MDUST321 | ERR352494 | N/A | 4.2.1 | unknown | unknown | Blood | 1983 | Australia & Oceania | Oceania | Fiji | unknown | unknown | Wong et al, 2015 |
| MDUST322 | ERR352495 | N/A | 2.3.5 | unknown | unknown | Blood | 1983 | Australia & Oceania | Oceania | Fiji | unknown | unknown | Wong et al, 2015 |
| MDUST330 | ERR352496 | N/A | 2.1.7.2 | unknown | unknown | Blood | 1992 | Australia & Oceania | Oceania | Papua New Guinea | unknown | unknown | Wong et al, 2015 |
| MDUST331 | ERR352497 | N/A | 2.1.7.1 | unknown | unknown | Blood | 1992 | Australia & Oceania | Oceania | Papua New Guinea | unknown | unknown | Wong et al, 2015 |
| MDUST336 | ERR352498 | N/A | 4.3.1.1 | unknown | unknown | Stool | 1993 | Australia & Oceania | Oceania | Fiji | unknown | unknown | Wong et al, 2015 |
| MDUST339 | ERR352499 | N/A | 2.3.5 | unknown | unknown | Stool | 1984 | Australia & Oceania | Oceania | Fiji | unknown | unknown | Wong et al, 2015 |
| MDUST348 | ERR352500 | N/A | 2.1.7.2 | unknown | unknown | Unknown | 1985 | Australia & Oceania | Oceania | Papua New Guinea | unknown | unknown | Wong et al, 2015 |
| MDUST351 | ERR352501 | N/A | 3.5.1 | unknown | unknown | Stool | 1985 | Australia & Oceania | Oceania | Fiji | unknown | unknown | Wong et al, 2015 |
| MDUST352 | ERR352502 | N/A | 2.3.5 | unknown | unknown | Blood | 1986 | Australia & Oceania | Oceania | Fiji | unknown | unknown | Wong et al, 2015 |
| MDUST358 | ERR352503 | N/A | 2.1.7.2 | unknown | unknown | Stool | 1996 | Australia & Oceania | Oceania | Papua New Guinea | unknown | unknown | Wong et al, 2015 |
| MDUST368 | ERR352504 | N/A | 4.3.1.2 | unknown | unknown | Blood | 2011 | Asia | South Asia | India | unknown | unknown | Wong et al, 2015 |
| MDUST383 | ERR352505 | N/A | 4.3.1.1 | unknown | unknown | Blood | 2011 | Asia | South Asia | India | unknown | unknown | Wong et al, 2015 |
| MDUST388 | ERR352506 | N/A | 4.3.1.1 | unknown | unknown | Blood | 2011 | Asia | South Asia | India | unknown | unknown | Wong et al, 2015 |
| MDUST389 | ERR352507 | N/A | 4.3.1.2 | unknown | unknown | Blood | 2011 | Asia | Southeast Asia | Indonesia | unknown | unknown | Wong et al, 2015 |
| MDUST401 | ERR352508 | N/A | 2.0.1 | unknown | unknown | Blood | 2012 | Asia | South Asia | Bangladesh | unknown | unknown | Wong et al, 2015 |
| MDUST402 | ERR352509 | N/A | 4.3.1.1 | unknown | unknown | Blood | 2012 | Asia | South Asia | Afghanistan | unknown | unknown | Wong et al, 2015 |
| MDUST407 | ERR352510 | N/A | 2.2.2 | unknown | unknown | Stool | 2007 | Asia | Western Asia | Lebanon | unknown | unknown | Wong et al, 2015 |
| MDUST415 | ERR352511 | N/A | 3 | unknown | unknown | Blood | 2012 | Unknown | Unknown | Unknown | unknown | unknown | Wong et al, 2015 |
| 3592/3 | ERR352599 | N/A | 2.5.1 | unknown | unknown | Blood | 2011 | Africa | Central Africa | DRC | unknown | unknown | Wong et al, 2015 |
| 3306/3 | ERR352602 | N/A | 2.5.1 | unknown | unknown | Blood | 2011 | Africa | Central Africa | DRC | unknown | unknown | Wong et al, 2015 |
| 3322/3 | ERR352604 | N/A | 2.5.1 | unknown | unknown | Blood | 2011 | Africa | Central Africa | DRC | unknown | unknown | Wong et al, 2015 |
| dn51 | ERR352605 | N/A | 4.3.1.1 | unknown | unknown | Blood | 1995 | Asia | Southeast Asia | Vietnam | unknown | unknown | Wong et al, 2015 |
| dn17 | ERR352606 | N/A | 4.3.1.1 | unknown | unknown | Blood | 1995 | Asia | Southeast Asia | Vietnam | unknown | unknown | Wong et al, 2015 |
| dn40 | ERR352607 | N/A | 4.3.1.1 | unknown | unknown | Blood | 1995 | Asia | Southeast Asia | Vietnam | unknown | unknown | Wong et al, 2015 |
| dtc84 | ERR352608 | N/A | 4.3.1.1 | unknown | unknown | Blood | 1994 | Asia | Southeast Asia | Vietnam | unknown | unknown | Wong et al, 2015 |
| dn126 | ERR352609 | N/A | 4.3.1.1 | unknown | unknown | Blood | 1996 | Asia | Southeast Asia | Vietnam | unknown | unknown | Wong et al, 2015 |
| dtc81 | ERR352611 | N/A | 4.3.1.1 | unknown | unknown | Blood | 1994 | Asia | Southeast Asia | Vietnam | unknown | unknown | Wong et al, 2015 |
| dn192 | ERR352612 | N/A | 4.3.1.1 | unknown | unknown | Blood | 1996 | Asia | Southeast Asia | Vietnam | unknown | unknown | Wong et al, 2015 |
| dn152 | ERR352613 | N/A | 4.3.1.1 | unknown | unknown | Blood | 1996 | Asia | Southeast Asia | Vietnam | unknown | unknown | Wong et al, 2015 |
| dtc76 | ERR352614 | N/A | 4.3.1.1 | unknown | unknown | Blood | 1994 | Asia | Southeast Asia | Vietnam | unknown | unknown | Wong et al, 2015 |
| dn95 | ERR352615 | N/A | 4.3.1.1 | unknown | unknown | Blood | 1995 | Asia | Southeast Asia | Vietnam | unknown | unknown | Wong et al, 2015 |
| dn182 | ERR352616 | N/A | 4.3.1.1 | unknown | unknown | Blood | 1996 | Asia | Southeast Asia | Vietnam | unknown | unknown | Wong et al, 2015 |
| dtc109 | ERR352617 | N/A | 4.3.1.1 | unknown | unknown | Blood | 1994 | Asia | Southeast Asia | Vietnam | unknown | unknown | Wong et al, 2015 |
| dn38 | ERR352618 | N/A | 4.3.1.1 | unknown | unknown | Blood | 1995 | Asia | Southeast Asia | Vietnam | unknown | unknown | Wong et al, 2015 |
| dn122 | ERR352619 | N/A | 4.3.1.1 | unknown | unknown | Blood | 1996 | Asia | Southeast Asia | Vietnam | unknown | unknown | Wong et al, 2015 |
| dn154 | ERR352620 | N/A | 4.3.1.1 | unknown | unknown | Blood | 1996 | Asia | Southeast Asia | Vietnam | unknown | unknown | Wong et al, 2015 |
| dn67 | ERR352621 | N/A | 4.3.1.1 | unknown | unknown | Blood | 1995 | Asia | Southeast Asia | Vietnam | unknown | unknown | Wong et al, 2015 |
| dn174 | ERR352623 | N/A | 4.3.1.1 | unknown | unknown | Blood | 1996 | Asia | Southeast Asia | Vietnam | unknown | unknown | Wong et al, 2015 |
| dn145 | ERR352624 | N/A | 4.3.1.1 | unknown | unknown | Blood | 1996 | Asia | Southeast Asia | Vietnam | unknown | unknown | Wong et al, 2015 |
| dn56 | ERR352625 | N/A | 4.3.1.1 | unknown | unknown | Blood | 1995 | Asia | Southeast Asia | Vietnam | unknown | unknown | Wong et al, 2015 |
| dtc115 | ERR352626 | N/A | 4.3.1.1 | unknown | unknown | Blood | 1995 | Asia | Southeast Asia | Vietnam | unknown | unknown | Wong et al, 2015 |
| dn113 | ERR352627 | N/A | 4.3.1.1 | unknown | unknown | Blood | 1996 | Asia | Southeast Asia | Vietnam | unknown | unknown | Wong et al, 2015 |
| dn26 | ERR352628 | N/A | 4.3.1.1 | unknown | unknown | Blood | 1995 | Asia | Southeast Asia | Vietnam | unknown | unknown | Wong et al, 2015 |
| dn137 | ERR352629 | N/A | 4.3.1.1 | unknown | unknown | Blood | 1996 | Asia | Southeast Asia | Vietnam | unknown | unknown | Wong et al, 2015 |
| dn178 | ERR352631 | N/A | 4.3.1.1 | unknown | unknown | Blood | 1996 | Asia | Southeast Asia | Vietnam | unknown | unknown | Wong et al, 2015 |
| dn22 | ERR352632 | N/A | 4.3.1.1 | unknown | unknown | Blood | 1995 | Asia | Southeast Asia | Vietnam | unknown | unknown | Wong et al, 2015 |
| dn89 | ERR352633 | N/A | 4.3.1.1 | unknown | unknown | Blood | 1995 | Asia | Southeast Asia | Vietnam | unknown | unknown | Wong et al, 2015 |
| dtc80 | ERR352635 | N/A | 4.3.1.1 | unknown | unknown | Blood | 1994 | Asia | Southeast Asia | Vietnam | unknown | unknown | Wong et al, 2015 |
| dn21 | ERR352636 | N/A | 4.3.1.1 | unknown | unknown | Blood | 1995 | Asia | Southeast Asia | Vietnam | unknown | unknown | Wong et al, 2015 |
| dn176 | ERR352637 | N/A | 4.3.1.1 | unknown | unknown | Blood | 1996 | Asia | Southeast Asia | Vietnam | unknown | unknown | Wong et al, 2015 |

| | | | | | | | | | | | | | |
|---|---|---|---|---|---|---|---|---|---|---|---|---|---|
| dn148 | ERR352638 | N/A | 4.3.1.1 | unknown | unknown | Blood | 1996 | Asia | Southeast Asia | Vietnam | unknown | unknown | Wong et al, 2015 |
| dn149 | ERR352639 | N/A | 4.3.1.1 | unknown | unknown | Blood | 1996 | Asia | Southeast Asia | Vietnam | unknown | unknown | Wong et al, 2015 |
| dn123 | ERR352640 | N/A | 4.3.1.1 | unknown | unknown | Blood | 1996 | Asia | Southeast Asia | Vietnam | unknown | unknown | Wong et al, 2015 |
| dtc57 | ERR352641 | N/A | 4.3.1.1 | unknown | unknown | Blood | 1994 | Asia | Southeast Asia | Vietnam | unknown | unknown | Wong et al, 2015 |
| dtc90 | ERR352642 | N/A | 4.3.1.1 | unknown | unknown | Blood | 1994 | Asia | Southeast Asia | Vietnam | unknown | unknown | Wong et al, 2015 |
| ty3-201 | ERR352643 | N/A | 4.3.1.1 | unknown | unknown | Blood | 1997 | Asia | Southeast Asia | Vietnam | unknown | unknown | Wong et al, 2015 |
| ty3-224 | ERR352644 | N/A | 4.3.1.1 | unknown | unknown | Blood | 1997 | Asia | Southeast Asia | Vietnam | unknown | unknown | Wong et al, 2015 |
| ct1-105 | ERR352645 | N/A | 4.3.1.1 | unknown | unknown | Blood | 1994 | Asia | Southeast Asia | Vietnam | unknown | unknown | Wong et al, 2015 |
| ty3-202 | ERR352646 | N/A | 4.3.1.1 | unknown | unknown | Blood | 1997 | Asia | Southeast Asia | Vietnam | unknown | unknown | Wong et al, 2015 |
| ty3-226 | ERR352647 | N/A | 4.3.1.1 | unknown | unknown | Blood | 1997 | Asia | Southeast Asia | Vietnam | unknown | unknown | Wong et al, 2015 |
| ct1-107 | ERR352648 | N/A | 4.3.1.1 | unknown | unknown | Blood | 1994 | Asia | Southeast Asia | Vietnam | unknown | unknown | Wong et al, 2015 |
| ty3-205 | ERR352649 | N/A | 4.3.1.1 | unknown | unknown | Blood | 1997 | Asia | Southeast Asia | Vietnam | unknown | unknown | Wong et al, 2015 |
| ty3-243 | ERR352650 | N/A | 4.3.1.1 | unknown | unknown | Blood | 1998 | Asia | Southeast Asia | Vietnam | unknown | unknown | Wong et al, 2015 |
| ipt51 | ERR352651 | N/A | 4.3.1.1 | unknown | unknown | Blood | 1997 | Asia | Southeast Asia | Vietnam | unknown | unknown | Wong et al, 2015 |
| ty3-206 | ERR352652 | N/A | 4.3.1.1 | unknown | unknown | Blood | 1997 | Asia | Southeast Asia | Vietnam | unknown | unknown | Wong et al, 2015 |
| dty1-80 | ERR352654 | N/A | 4.3.1.1 | unknown | unknown | Blood | 1997 | Asia | Southeast Asia | Vietnam | unknown | unknown | Wong et al, 2015 |
| ty2-173 | ERR352655 | N/A | 4.3.1.1 | unknown | unknown | Blood | 1995 | Asia | Southeast Asia | Vietnam | unknown | unknown | Wong et al, 2015 |
| ty3-207 | ERR352656 | N/A | 4.3.1.1 | unknown | unknown | Blood | 1997 | Asia | Southeast Asia | Vietnam | unknown | unknown | Wong et al, 2015 |
| ty3-254 | ERR352657 | N/A | 4.3.1.1 | unknown | unknown | Blood | 1998 | Asia | Southeast Asia | Vietnam | unknown | unknown | Wong et al, 2015 |
| ct1-66 | ERR352658 | N/A | 4.3.1.1 | unknown | unknown | Blood | 1994 | Asia | Southeast Asia | Vietnam | unknown | unknown | Wong et al, 2015 |
| as118 | ERR352659 | N/A | 4.3.1.1 | unknown | unknown | Blood | 1998 | Asia | Southeast Asia | Vietnam | unknown | unknown | Wong et al, 2015 |
| ipt61 | ERR352661 | N/A | 4.3.1.1 | unknown | unknown | Blood | 1997 | Asia | Southeast Asia | Vietnam | unknown | unknown | Wong et al, 2015 |
| ty3-191 | ERR352662 | N/A | 4.3.1.1 | unknown | unknown | Blood | 1997 | Asia | Southeast Asia | Vietnam | unknown | unknown | Wong et al, 2015 |
| ty3-269 | ERR352663 | N/A | 4.3.1.1 | unknown | unknown | Blood | 1998 | Asia | Southeast Asia | Vietnam | unknown | unknown | Wong et al, 2015 |
| as250 | ERR352665 | N/A | 4.3.1.1 | unknown | unknown | Blood | 1999 | Asia | Southeast Asia | Vietnam | unknown | unknown | Wong et al, 2015 |
| ty3-211 | ERR352667 | N/A | 4.3.1.1 | unknown | unknown | Blood | 1997 | Asia | Southeast Asia | Vietnam | unknown | unknown | Wong et al, 2015 |
| ct1-70 | ERR352669 | N/A | 4.3.1.1 | unknown | unknown | Blood | 1994 | Asia | Southeast Asia | Vietnam | unknown | unknown | Wong et al, 2015 |
| as252 | ERR352671 | N/A | 4.3.1.1 | unknown | unknown | Blood | 1999 | Asia | Southeast Asia | Vietnam | unknown | unknown | Wong et al, 2015 |
| ty3-194 | ERR352673 | N/A | 4.3.1.1 | unknown | unknown | Blood | 1997 | Asia | Southeast Asia | Vietnam | unknown | unknown | Wong et al, 2015 |
| ipt83 | ERR352677 | N/A | 4.3.1.1 | unknown | unknown | Blood | 1998 | Asia | Southeast Asia | Vietnam | unknown | unknown | Wong et al, 2015 |
| ty3-196 | ERR352679 | N/A | 4.3.1.1 | unknown | unknown | Blood | 1997 | Asia | Southeast Asia | Vietnam | unknown | unknown | Wong et al, 2015 |
| ty3-222 | ERR352681 | N/A | 4.3.1.1 | unknown | unknown | Blood | 1997 | Asia | Southeast Asia | Vietnam | unknown | unknown | Wong et al, 2015 |
| ty3-197 | ERR352685 | N/A | 4.3.1.1 | unknown | unknown | Blood | 1997 | Asia | Southeast Asia | Vietnam | unknown | unknown | Wong et al, 2015 |
| ty3-213 | ERR352687 | N/A | 3.2.1 | unknown | unknown | Blood | 1997 | Asia | Southeast Asia | Vietnam | unknown | unknown | Wong et al, 2015 |
| ct1-69 | ERR352689 | N/A | 4.3.1.1 | unknown | unknown | Blood | 1994 | Asia | Southeast Asia | Vietnam | unknown | unknown | Wong et al, 2015 |
| ipt16 | ERR352691 | N/A | 4.3.1.1 | unknown | unknown | Blood | 1995 | Asia | Southeast Asia | Vietnam | unknown | unknown | Wong et al, 2015 |
| ipt19 | ERR352697 | N/A | 4.3.1.1 | unknown | unknown | Blood | 1995 | Asia | Southeast Asia | Vietnam | unknown | unknown | Wong et al, 2015 |
| ty3-199 | ERR352699 | N/A | 3.2.1 | unknown | unknown | Blood | 1997 | Asia | Southeast Asia | Vietnam | unknown | unknown | Wong et al, 2015 |
| ty2-105 | ERR352703 | N/A | 4.3.1.1 | unknown | unknown | Blood | 1994 | Asia | Southeast Asia | Vietnam | unknown | unknown | Wong et al, 2015 |
| ty2-138 | ERR352705 | N/A | 4.1 | unknown | unknown | Blood | 1994 | Asia | Southeast Asia | Vietnam | unknown | unknown | Wong et al, 2015 |
| ty2-106 | ERR352709 | N/A | 4.3.1.1 | unknown | unknown | Blood | 1994 | Asia | Southeast Asia | Vietnam | unknown | unknown | Wong et al, 2015 |
| ty2-141 | ERR352711 | N/A | 4.3.1.1 | unknown | unknown | Blood | 1994 | Asia | Southeast Asia | Vietnam | unknown | unknown | Wong et al, 2015 |
| ty2-143 | ERR352713 | N/A | 4.3.1.1 | unknown | unknown | Blood | 1994 | Asia | Southeast Asia | Vietnam | unknown | unknown | Wong et al, 2015 |
| ty2-113 | ERR352715 | N/A | 4.3.1.1 | unknown | unknown | Blood | 1994 | Asia | Southeast Asia | Vietnam | unknown | unknown | Wong et al, 2015 |
| ty2-116 | ERR352717 | N/A | 4.3.1.1 | unknown | unknown | Blood | 1994 | Asia | Southeast Asia | Vietnam | unknown | unknown | Wong et al, 2015 |
| ty2-133 | ERR352719 | N/A | 4.3.1.1 | unknown | unknown | Blood | 1994 | Asia | Southeast Asia | Vietnam | unknown | unknown | Wong et al, 2015 |
| ty2-154 | ERR352721 | N/A | 4.3.1.1 | unknown | unknown | Blood | 1994 | Asia | Southeast Asia | Vietnam | unknown | unknown | Wong et al, 2015 |
| ty2-119 | ERR352725 | N/A | 4.3.1.1 | unknown | unknown | Blood | 1994 | Asia | Southeast Asia | Vietnam | unknown | unknown | Wong et al, 2015 |
| MDUST107 | ERR352941 | N/A | 2.1.7.1 | unknown | unknown | Blood | 2007 | Australia & Oceania | Oceania | Papua New Guinea | unknown | unknown | Wong et al, 2015 |
| MDUST111 | ERR352942 | N/A | 3.3 | unknown | unknown | Urine | 2008 | Africa | North Africa | Sudan | unknown | unknown | Wong et al, 2015 |

| | | | | | | | | | | | | | |
|---|---|---|---|---|---|---|---|---|---|---|---|---|---|
| MDUST181 | ERR352943 | N/A | 4.3.1.2 | unknown | unknown | Blood | 2011 | Unknown | Unknown | Unknown | unknown | unknown | Wong et al, 2015 |
| MDUST186 | ERR352944 | N/A | 4.3.1.2 | unknown | unknown | Blood | 2011 | Asia | South Asia | India | unknown | unknown | Wong et al, 2015 |
| MDUST188 | ERR352945 | N/A | 3 | unknown | unknown | Blood | 2011 | Asia | Southeast Asia | Indonesia | unknown | unknown | Wong et al, 2015 |
| MDUST189 | ERR352946 | N/A | 4.3.1.2 | unknown | unknown | Blood | 2011 | Asia | South Asia | India | unknown | unknown | Wong et al, 2015 |
| MDUST208 | ERR352947 | N/A | 2.2.1 | unknown | unknown | Blood | 2010 | Australia & Oceania | Australia | Australia | unknown | unknown | Wong et al, 2015 |
| MDUST209 | ERR352948 | N/A | 4.3.1.1 | unknown | unknown | Blood | 2011 | Asia | South Asia | India | unknown | unknown | Wong et al, 2015 |
| MDUST210 | ERR352949 | N/A | 4.3.1.2 | unknown | unknown | Blood | 2011 | Asia | South Asia | India | unknown | unknown | Wong et al, 2015 |
| MDUST212 | ERR352950 | N/A | 2.2.2 | unknown | unknown | Blood | 2011 | Asia | South Asia | India | unknown | unknown | Wong et al, 2015 |
| MDUST213 | ERR352951 | N/A | 4.3.1 | unknown | unknown | Blood | 2011 | Asia | South Asia | India | unknown | unknown | Wong et al, 2015 |
| MDUST219 | ERR352952 | N/A | 2.2.2 | unknown | unknown | Stool | 2011 | Asia | Western Asia | Lebanon | unknown | unknown | Wong et al, 2015 |
| MDUST224 | ERR352953 | N/A | 4.3.1.2 | unknown | unknown | Blood | 2011 | Asia | South Asia | India | unknown | unknown | Wong et al, 2015 |
| MDUST225 | ERR352954 | N/A | 4.3.1.2 | unknown | unknown | Blood | 2011 | Asia | South Asia | India | unknown | unknown | Wong et al, 2015 |
| ST1134/01 | ERR353330 | N/A | 2 | unknown | unknown | Stool | 2001 | South America | South America | Argentina | unknown | unknown | Wong et al, 2015 |
| ST1197/88 | ERR353331 | N/A | 2.3.3 | unknown | unknown | Urine | 1988 | South America | South America | Argentina | unknown | unknown | Wong et al, 2015 |
| ST1309/04 | ERR353332 | N/A | 2.3.3 | unknown | unknown | Blood | 2004 | South America | South America | Argentina | unknown | unknown | Wong et al, 2015 |
| ST1625/88 | ERR353334 | N/A | 4.1 | unknown | unknown | Stool | 1988 | South America | South America | Argentina | unknown | unknown | Wong et al, 2015 |
| ST1921/06 | ERR353335 | N/A | 2.3.3 | unknown | unknown | Stool | 2006 | South America | South America | Argentina | unknown | unknown | Wong et al, 2015 |
| ST2338/98 | ERR353336 | N/A | 2.3.3 | unknown | unknown | Blood | 1905 | South America | South America | Argentina | unknown | unknown | Wong et al, 2015 |
| ST3090/99 | ERR353338 | N/A | 2 | unknown | unknown | Stool | 1999 | South America | South America | Argentina | unknown | unknown | Wong et al, 2015 |
| ST472/01 | ERR353339 | N/A | 4.1 | unknown | unknown | Stool | 1905 | South America | South America | Argentina | unknown | unknown | Wong et al, 2015 |
| ST805/02 | ERR353340 | N/A | 2.3.2 | unknown | unknown | Stool | 2002 | South America | South America | Argentina | unknown | unknown | Wong et al, 2015 |
| ST821/98 | ERR353341 | N/A | 4.1 | unknown | unknown | Stool | 1905 | South America | South America | Argentina | unknown | unknown | Wong et al, 2015 |
| ST860/95 | ERR353342 | N/A | 2.3.2 | unknown | unknown | Stool | 1905 | South America | South America | Argentina | unknown | unknown | Wong et al, 2015 |
| MDUST114 | ERR357440 | N/A | 3.5.4 | unknown | unknown | Blood | 2009 | Australia & Oceania | Oceania | Samoa | unknown | unknown | Wong et al, 2015 |
| MDUST138 | ERR357441 | N/A | 3.3.1 | unknown | unknown | Stool | 2012 | Asia | Southeast Asia | Myanmar | unknown | unknown | Wong et al, 2015 |
| MDUST155 | ERR357442 | N/A | 4.3.1.1 | unknown | unknown | Blood | 2012 | Asia | South Asia | India | unknown | unknown | Wong et al, 2015 |
| MDUST160 | ERR357443 | N/A | 3.0.2 | unknown | unknown | Stool | 2012 | Asia | South Asia | India | unknown | unknown | Wong et al, 2015 |
| MDUST164 | ERR357444 | N/A | 4.3.1.2 | unknown | unknown | Blood | 2011 | Asia | South Asia | India | unknown | unknown | Wong et al, 2015 |
| MDUST178 | ERR357445 | N/A | 2.1 | unknown | unknown | Stool | 2011 | Asia | Southeast Asia | EastTimor | unknown | unknown | Wong et al, 2015 |
| MDUST180 | ERR357446 | N/A | 4.3.1.2 | unknown | unknown | Blood | 2011 | Asia | South Asia | India | unknown | unknown | Wong et al, 2015 |
| MDUST277 | ERR357447 | N/A | 2.1.7.1 | unknown | unknown | Blood | 2001 | Australia & Oceania | Oceania | Papua New Guinea | unknown | unknown | Wong et al, 2015 |
| MDUST281 | ERR357449 | N/A | 2.1.7.1 | unknown | unknown | Blood | 2002 | Australia & Oceania | Oceania | Papua New Guinea | unknown | unknown | Wong et al, 2015 |
| MDUST284 | ERR357450 | N/A | 4.1 | unknown | unknown | Blood | 2004 | Australia & Oceania | Oceania | Samoa | unknown | unknown | Wong et al, 2015 |
| MDUST285 | ERR357451 | N/A | 4.1 | unknown | unknown | Blood | 2004 | Australia & Oceania | Oceania | Samoa | unknown | unknown | Wong et al, 2015 |
| MDUST293 | ERR357452 | N/A | 2.1.7.1 | unknown | unknown | Stool | 2009 | Australia & Oceania | Oceania | Papua New Guinea | unknown | unknown | Wong et al, 2015 |
| MDUST294 | ERR357453 | N/A | 4.3.1.2 | unknown | unknown | Blood | 2010 | Asia | South Asia | Nepal | unknown | unknown | Wong et al, 2015 |
| MDUST303 | ERR357454 | N/A | 4.2 | unknown | unknown | Unknown | 1980 | Australia & Oceania | Oceania | Tonga | unknown | unknown | Wong et al, 2015 |
| MDUST304 | ERR357455 | N/A | 2.1.7.1 | unknown | unknown | Stool | 1990 | Australia & Oceania | Oceania | Papua New Guinea | unknown | unknown | Wong et al, 2015 |
| MDUST305 | ERR357456 | N/A | 2.1.7.1 | unknown | unknown | Blood | 1990 | Australia & Oceania | Oceania | Papua New Guinea | unknown | unknown | Wong et al, 2015 |
| MDUST306 | ERR357457 | N/A | 4.2.1 | unknown | unknown | Unknown | 1981 | Australia & Oceania | Oceania | Fiji | unknown | unknown | Wong et al, 2015 |
| MDUST309 | ERR357459 | N/A | 4.2 | unknown | unknown | Unknown | 1981 | Australia & Oceania | Oceania | Fiji | unknown | unknown | Wong et al, 2015 |
| MDUST311 | ERR357460 | N/A | 4.2 | unknown | unknown | Stool | 1982 | Australia & Oceania | Oceania | Fiji | unknown | unknown | Wong et al, 2015 |
| MDUST312 | ERR357461 | N/A | 4.2 | unknown | unknown | Stool | 1982 | Australia & Oceania | Oceania | Fiji | unknown | unknown | Wong et al, 2015 |
| MDUST315 | ERR357462 | N/A | 4.1 | unknown | unknown | Stool | 1991 | Australia & Oceania | Oceania | Vanuatu | unknown | unknown | Wong et al, 2015 |
| MDUST316 | ERR357463 | N/A | 4.2 | unknown | unknown | Unknown | 1982 | Australia & Oceania | Oceania | Fiji | unknown | unknown | Wong et al, 2015 |
| MDUST317 | ERR357464 | N/A | 4.2.1 | unknown | unknown | Knee aspirate | 1982 | Australia & Oceania | Oceania | Fiji | unknown | unknown | Wong et al, 2015 |
| MDUST318 | ERR357465 | N/A | 4.2 | unknown | unknown | Unknown | 1982 | Australia & Oceania | Oceania | Fiji | unknown | unknown | Wong et al, 2015 |
| MDUST320 | ERR357466 | N/A | 4.2.1 | unknown | unknown | Stool | 1983 | Australia & Oceania | Oceania | Fiji | unknown | unknown | Wong et al, 2015 |
| MDUST323 | ERR357467 | N/A | 4.2.1 | unknown | unknown | Blood | 1983 | Australia & Oceania | Oceania | Fiji | unknown | unknown | Wong et al, 2015 |
| MDUST325 | ERR357468 | N/A | 3.5.4 | unknown | unknown | Blood | 1992 | Australia & Oceania | Oceania | Samoa | unknown | unknown | Wong et al, 2015 |

| | | | | | | | | | | | | | |
|---|---|---|---|---|---|---|---|---|---|---|---|---|---|
| MDUST327 | ERR357469 | N/A | 4.3.1.1 | unknown | unknown | Blood | 1992 | Australia & Oceania | Oceania | Fiji | unknown | unknown | Wong et al, 2015 |
| MDUST329 | ERR357470 | N/A | 2.1.7.1 | unknown | unknown | Blood | 1992 | Australia & Oceania | Oceania | Papua New Guinea | unknown | unknown | Wong et al, 2015 |
| MDUST332 | ERR357471 | N/A | 2.1.7.2 | unknown | unknown | Blood | 1992 | Australia & Oceania | Oceania | Papua New Guinea | unknown | unknown | Wong et al, 2015 |
| MDUST333 | ERR357472 | N/A | 2.1.7.1 | unknown | unknown | Blood | 1992 | Australia & Oceania | Oceania | Papua New Guinea | unknown | unknown | Wong et al, 2015 |
| MDUST334 | ERR357473 | N/A | 3 | unknown | unknown | Blood | 1984 | Australia & Oceania | Oceania | Fiji | unknown | unknown | Wong et al, 2015 |
| MDUST340 | ERR357474 | N/A | 4.3.1.1 | unknown | unknown | Stool | 1993 | Australia & Oceania | Oceania | Fiji | unknown | unknown | Wong et al, 2015 |
| MDUST341 | ERR357475 | N/A | 4.3.1.1 | unknown | unknown | Stool | 1993 | Australia & Oceania | Oceania | Fiji | unknown | unknown | Wong et al, 2015 |
| MDUST342 | ERR357476 | N/A | 4.3.1.1 | unknown | unknown | Stool | 1993 | Australia & Oceania | Oceania | Fiji | unknown | unknown | Wong et al, 2015 |
| MDUST344 | ERR357477 | N/A | 4.2.1 | unknown | unknown | Stool | 1985 | Australia & Oceania | Oceania | Fiji | unknown | unknown | Wong et al, 2015 |
| MDUST345 | ERR357478 | N/A | 2.1.7.2 | unknown | unknown | Unknown | 1985 | Australia & Oceania | Oceania | Papua New Guinea | unknown | unknown | Wong et al, 2015 |
| MDUST349 | ERR357479 | N/A | 2.3.5 | unknown | unknown | Stool | 1985 | Australia & Oceania | Oceania | Fiji | unknown | unknown | Wong et al, 2015 |
| MDUST356 | ERR357480 | N/A | 2.1.7.1 | unknown | unknown | Blood | 1994 | Australia & Oceania | Oceania | Papua New Guinea | unknown | unknown | Wong et al, 2015 |
| MDUST360 | ERR357481 | N/A | 2.1.7.1 | unknown | unknown | Stool | 1998 | Australia & Oceania | Oceania | Papua New Guinea | unknown | unknown | Wong et al, 2015 |
| np4 | ERR357576 | N/A | 4.3.1.2 | unknown | unknown | Blood | 2011 | Asia | South Asia | Nepal | unknown | unknown | Wong et al, 2015 |
| np41 | ERR357577 | N/A | 3.3.2 | unknown | unknown | Blood | 2011 | Asia | South Asia | Nepal | unknown | unknown | Wong et al, 2015 |
| np94 | ERR357578 | N/A | 4.3.1.2 | unknown | unknown | Blood | 2011 | Asia | South Asia | Nepal | unknown | unknown | Wong et al, 2015 |
| 3671/3 | ERR357579 | N/A | 2.5.1 | unknown | unknown | Blood | 2011 | Africa | Central Africa | DRC | unknown | unknown | Wong et al, 2015 |
| 3332/3 | ERR357580 | N/A | 2.5.1 | unknown | unknown | Blood | 2011 | Africa | Central Africa | DRC | unknown | unknown | Wong et al, 2015 |
| ERL113717 | ERR357581 | N/A | 4.3.1 | unknown | unknown | Blood | 2011 | Asia | South Asia | India | unknown | unknown | Wong et al, 2015 |
| np5 | ERR357583 | N/A | 4.3.1.2 | unknown | unknown | Blood | 2011 | Asia | South Asia | Nepal | unknown | unknown | Wong et al, 2015 |
| np43 | ERR357584 | N/A | 4.3.1.2 | unknown | unknown | Blood | 2011 | Asia | South Asia | Nepal | unknown | unknown | Wong et al, 2015 |
| np95 | ERR357585 | N/A | 4.3.1.2 | unknown | unknown | Blood | 2011 | Asia | South Asia | Nepal | unknown | unknown | Wong et al, 2015 |
| np8 | ERR357587 | N/A | 4.3.1.2 | unknown | unknown | Blood | 2011 | Asia | South Asia | Nepal | unknown | unknown | Wong et al, 2015 |
| np44 | ERR357588 | N/A | 4.3.1.2 | unknown | unknown | Blood | 2011 | Asia | South Asia | Nepal | unknown | unknown | Wong et al, 2015 |
| np75 | ERR357589 | N/A | 4.3.1.2 | unknown | unknown | Blood | 2011 | Asia | South Asia | Nepal | unknown | unknown | Wong et al, 2015 |
| np97 | ERR357590 | N/A | 4.3.1.2 | unknown | unknown | Blood | 2011 | Asia | South Asia | Nepal | unknown | unknown | Wong et al, 2015 |
| 3632/3 | ERR357591 | N/A | 2.5.1 | unknown | unknown | Blood | 2011 | Africa | Central Africa | DRC | unknown | unknown | Wong et al, 2015 |
| 3139/3 | ERR357592 | N/A | 2.5.1 | unknown | unknown | Blood | 2010 | Africa | Central Africa | DRC | unknown | unknown | Wong et al, 2015 |
| H12ESR01052-001A | ERR357593 | N/A | 3.5.4 | unknown | unknown | Blood | 2012 | Australia & Oceania | Oceania | Samoa | unknown | unknown | Wong et al, 2015 |
| np11 | ERR357594 | N/A | 4.3.1.2 | unknown | unknown | Blood | 2011 | Asia | South Asia | Nepal | unknown | unknown | Wong et al, 2015 |
| np99 | ERR357595 | N/A | 3.3.2 | unknown | unknown | Blood | 2011 | Asia | South Asia | Nepal | unknown | unknown | Wong et al, 2015 |
| 3182/3 | ERR357596 | N/A | 2.5.1 | unknown | unknown | Blood | 2010 | Africa | Central Africa | DRC | unknown | unknown | Wong et al, 2015 |
| H12ESR01946-001A | ERR357598 | N/A | 3.5.4 | unknown | unknown | Unknown | 2012 | Australia & Oceania | Oceania | Samoa | unknown | unknown | Wong et al, 2015 |
| dtc97 | ERR357599 | N/A | 4.3.1.1 | unknown | unknown | Blood | 1994 | Asia | Southeast Asia | Vietnam | unknown | unknown | Wong et al, 2015 |
| np12 | ERR357600 | N/A | 3.3.2 | unknown | unknown | Blood | 2011 | Asia | South Asia | Nepal | unknown | unknown | Wong et al, 2015 |
| np49 | ERR357601 | N/A | 4.3.1.2 | unknown | unknown | Blood | 2011 | Asia | South Asia | Nepal | unknown | unknown | Wong et al, 2015 |
| np81 | ERR357602 | N/A | 4.3.1.2 | unknown | unknown | Blood | 2011 | Asia | South Asia | Nepal | unknown | unknown | Wong et al, 2015 |
| np101 | ERR357603 | N/A | 4.3.1.1 | unknown | unknown | Blood | 2011 | Asia | South Asia | Nepal | unknown | unknown | Wong et al, 2015 |
| 3157/3 | ERR357604 | N/A | 2.5.1 | unknown | unknown | Blood | 2010 | Africa | Central Africa | DRC | unknown | unknown | Wong et al, 2015 |
| H12ESR00394-001A | ERR357605 | N/A | 3.5.4 | unknown | unknown | Blood | 2012 | Australia & Oceania | Oceania | Samoa | unknown | unknown | Wong et al, 2015 |
| np13 | ERR357606 | N/A | 4.3.1.2 | unknown | unknown | Blood | 2011 | Asia | South Asia | Nepal | unknown | unknown | Wong et al, 2015 |
| np50 | ERR357607 | N/A | 3.2.2 | unknown | unknown | Blood | 2011 | Asia | South Asia | Nepal | unknown | unknown | Wong et al, 2015 |
| np83 | ERR357608 | N/A | 4.3.1.2 | unknown | unknown | Blood | 2011 | Asia | South Asia | Nepal | unknown | unknown | Wong et al, 2015 |
| 3653/3 | ERR357610 | N/A | 2.5.1 | unknown | unknown | Blood | 2011 | Africa | Central Africa | DRC | unknown | unknown | Wong et al, 2015 |
| H12ESR04893-001A | ERR357611 | N/A | 3.5.4 | unknown | unknown | Stool | 2012 | Australia & Oceania | Oceania | Samoa | unknown | unknown | Wong et al, 2015 |
| np16 | ERR357612 | N/A | 3.3.2 | unknown | unknown | Blood | 2011 | Asia | South Asia | Nepal | unknown | unknown | Wong et al, 2015 |
| np51 | ERR357613 | N/A | 4.3.1.2 | unknown | unknown | Blood | 2011 | Asia | South Asia | Nepal | unknown | unknown | Wong et al, 2015 |
| np87 | ERR357614 | N/A | 4.3.1.2 | unknown | unknown | Blood | 2011 | Asia | South Asia | Nepal | unknown | unknown | Wong et al, 2015 |
| ERL022368 | ERR357615 | N/A | 4.3.1.1 | unknown | unknown | Blood | 2002 | Asia | Western Asia | Western Asia | unknown | unknown | Wong et al, 2015 |
| H12ESR04732-001A | ERR357617 | N/A | 3.5.4 | unknown | unknown | Blood | 2012 | Australia & Oceania | Oceania | Samoa | unknown | unknown | Wong et al, 2015 |

| | | | | | | | | | | | | | |
|---|---|---|---|---|---|---|---|---|---|---|---|---|---|
| dtc122 | ERR357618 | N/A | 4.3.1.1 | unknown | unknown | Blood | 1995 | Asia | Southeast Asia | Vietnam | unknown | unknown | Wong et al, 2015 |
| np22 | ERR357619 | N/A | 4.3.1.2 | unknown | unknown | Blood | 2011 | Asia | South Asia | Nepal | unknown | unknown | Wong et al, 2015 |
| np57 | ERR357620 | N/A | 4.3.1.2 | unknown | unknown | Blood | 2011 | Asia | South Asia | Nepal | unknown | unknown | Wong et al, 2015 |
| np88 | ERR357621 | N/A | 4.3.1.2 | unknown | unknown | Blood | 2011 | Asia | South Asia | Nepal | unknown | unknown | Wong et al, 2015 |
| 3673/3 | ERR357623 | N/A | 2.5.1 | unknown | unknown | Blood | 2011 | Africa | Central Africa | DRC | unknown | unknown | Wong et al, 2015 |
| H12ESR00753-001A | ERR357624 | N/A | 3.5.4 | unknown | unknown | Stool | 2012 | Australia & Oceania | Oceania | Samoa | unknown | unknown | Wong et al, 2015 |
| np60 | ERR357625 | N/A | 4.3.1.2 | unknown | unknown | Blood | 2011 | Asia | South Asia | Nepal | unknown | unknown | Wong et al, 2015 |
| np89 | ERR357626 | N/A | 4.3.1.2 | unknown | unknown | Blood | 2011 | Asia | South Asia | Nepal | unknown | unknown | Wong et al, 2015 |
| H12ESR04835-001A | ERR357627 | N/A | 3.5.4 | unknown | unknown | Blood | 2012 | Australia & Oceania | Oceania | Samoa | unknown | unknown | Wong et al, 2015 |
| np31 | ERR357628 | N/A | 4.3.1.2 | unknown | unknown | Blood | 2011 | Asia | South Asia | Nepal | unknown | unknown | Wong et al, 2015 |
| np90 | ERR357629 | N/A | 4.3.1.2 | unknown | unknown | Blood | 2011 | Asia | South Asia | Nepal | unknown | unknown | Wong et al, 2015 |
| ERL09896 | ERR357630 | N/A | 4.3.1.2 | unknown | unknown | Blood | 2009 | Asia | South Asia | India | unknown | unknown | Wong et al, 2015 |
| H12ESR04928-001A | ERR357631 | N/A | 2.2.1 | unknown | unknown | Stool | 2012 | Australia & Oceania | Oceania | Samoa | unknown | unknown | Wong et al, 2015 |
| np39 | ERR357632 | N/A | 4.3.1.2 | unknown | unknown | Blood | 2011 | Asia | South Asia | Nepal | unknown | unknown | Wong et al, 2015 |
| np65 | ERR357633 | N/A | 4.3.1.2 | unknown | unknown | Blood | 2011 | Asia | South Asia | Nepal | unknown | unknown | Wong et al, 2015 |
| np92 | ERR357634 | N/A | 4.3.1.2 | unknown | unknown | Blood | 2011 | Asia | South Asia | Nepal | unknown | unknown | Wong et al, 2015 |
| 3143/3 | ERR357635 | N/A | 2.5.1 | unknown | unknown | Blood | 2010 | Africa | Central Africa | DRC | unknown | unknown | Wong et al, 2015 |
| ERL102292 | ERR357636 | N/A | 4.3.1 | unknown | unknown | Blood | 2010 | Asia | South Asia | India | unknown | unknown | Wong et al, 2015 |
| np40 | ERR357637 | N/A | 4.3.1.2 | unknown | unknown | Blood | 2011 | Asia | South Asia | Nepal | unknown | unknown | Wong et al, 2015 |
| np67 | ERR357638 | N/A | 4.3.1.2 | unknown | unknown | Blood | 2011 | Asia | South Asia | Nepal | unknown | unknown | Wong et al, 2015 |
| np93 | ERR357639 | N/A | 4.3.1.2 | unknown | unknown | Blood | 2011 | Asia | South Asia | Nepal | unknown | unknown | Wong et al, 2015 |
| ERL101102 | ERR357641 | N/A | 4.3.1.2 | unknown | unknown | Stool | 2010 | Asia | South Asia | India | unknown | unknown | Wong et al, 2015 |
| dtc99 | ERR357642 | N/A | 4.3.1.1 | unknown | unknown | Blood | 1994 | Asia | Southeast Asia | Vietnam | unknown | unknown | Wong et al, 2015 |
| dtc98 | ERR357643 | N/A | 4.3.1.1 | unknown | unknown | Blood | 1994 | Asia | Southeast Asia | Vietnam | unknown | unknown | Wong et al, 2015 |
| dn88 | ERR357644 | N/A | 4.3.1.1 | unknown | unknown | Blood | 1995 | Asia | Southeast Asia | Vietnam | unknown | unknown | Wong et al, 2015 |
| dtc20 | ERR357645 | N/A | 4.3.1.1 | unknown | unknown | Blood | 1994 | Asia | Southeast Asia | Vietnam | unknown | unknown | Wong et al, 2015 |
| dtc131 | ERR357646 | N/A | 3.4 | unknown | unknown | Blood | 1995 | Asia | Southeast Asia | Vietnam | unknown | unknown | Wong et al, 2015 |
| dtc180 | ERR357647 | N/A | 4.3.1.1 | unknown | unknown | Blood | 1995 | Asia | Southeast Asia | Vietnam | unknown | unknown | Wong et al, 2015 |
| dtc5 | ERR357648 | N/A | 3.2.1 | unknown | unknown | Blood | 1994 | Asia | Southeast Asia | Vietnam | unknown | unknown | Wong et al, 2015 |
| dtc107 | ERR357649 | N/A | 4.3.1.1 | unknown | unknown | Blood | 1994 | Asia | Southeast Asia | Vietnam | unknown | unknown | Wong et al, 2015 |
| ct1-13 | ERR357651 | N/A | 4.3.1.1 | unknown | unknown | Blood | 1993 | Asia | Southeast Asia | Vietnam | unknown | unknown | Wong et al, 2015 |
| ct1-19 | ERR357652 | N/A | 1.2.1 | unknown | unknown | Blood | 1993 | Asia | Southeast Asia | Vietnam | unknown | unknown | Wong et al, 2015 |
| dty1-99 | ERR357653 | N/A | 4.3.1.1 | unknown | unknown | Blood | 1997 | Asia | Southeast Asia | Vietnam | unknown | unknown | Wong et al, 2015 |
| dty1-71 | ERR357654 | N/A | 4.3.1.1 | unknown | unknown | Blood | 1997 | Asia | Southeast Asia | Vietnam | unknown | unknown | Wong et al, 2015 |
| ty2-121 | ERR357655 | N/A | 4.3.1.1 | unknown | unknown | Blood | 1994 | Asia | Southeast Asia | Vietnam | unknown | unknown | Wong et al, 2015 |
| ty2-122 | ERR357657 | N/A | 4.3.1.1 | unknown | unknown | Blood | 1994 | Asia | Southeast Asia | Vietnam | unknown | unknown | Wong et al, 2015 |
| ty1-48 | ERR357659 | N/A | 4.3.1.1 | unknown | unknown | Blood | 1993 | Asia | Southeast Asia | Vietnam | unknown | unknown | Wong et al, 2015 |
| ty2-124 | ERR357661 | N/A | 4.3.1.1 | unknown | unknown | Blood | 1994 | Asia | Southeast Asia | Vietnam | unknown | unknown | Wong et al, 2015 |
| ty2-68 | ERR357663 | N/A | 3 | unknown | unknown | Blood | 1994 | Asia | Southeast Asia | Vietnam | unknown | unknown | Wong et al, 2015 |
| ty2-69 | ERR357665 | N/A | 4.1 | unknown | unknown | Blood | 1994 | Asia | Southeast Asia | Vietnam | unknown | unknown | Wong et al, 2015 |
| ty1-30 | ERR357667 | N/A | 3.2.1 | unknown | unknown | Blood | 1993 | Asia | Southeast Asia | Vietnam | unknown | unknown | Wong et al, 2015 |
| MDUST110 | ERR357756 | N/A | 2.2.1 | unknown | unknown | Blood | 2007 | Africa | North Africa | Egypt | unknown | unknown | Wong et al, 2015 |
| MDUST112 | ERR357757 | N/A | 3.3 | unknown | unknown | Urine | 2008 | Africa | North Africa | Sudan | unknown | unknown | Wong et al, 2015 |
| MDUST119 | ERR357758 | N/A | 3.5.4 | unknown | unknown | Blood | 2009 | Australia & Oceania | Oceania | Samoa | unknown | unknown | Wong et al, 2015 |
| MDUST122 | ERR357759 | N/A | 3.5.4 | unknown | unknown | Urine | 2010 | Australia & Oceania | Oceania | Samoa | unknown | unknown | Wong et al, 2015 |
| MDUST124 | ERR357760 | N/A | 3.5.4 | unknown | unknown | Blood | 2011 | Australia & Oceania | Oceania | Samoa | unknown | unknown | Wong et al, 2015 |
| MDUST134 | ERR357761 | N/A | 3 | unknown | unknown | Stool | 2011 | Asia | Southeast Asia | Phillipines | unknown | unknown | Wong et al, 2015 |
| MDUST137 | ERR357762 | N/A | 3.3.1 | unknown | unknown | Blood | 2011 | Asia | South Asia | India | unknown | unknown | Wong et al, 2015 |
| MDUST142 | ERR357763 | N/A | 4.3.1.1 | unknown | unknown | Blood | 2012 | Asia | South Asia | India | unknown | unknown | Wong et al, 2015 |
| MDUST144 | ERR357764 | N/A | 4.3.1.1 | unknown | unknown | Blood | 2012 | Australia & Oceania | Australia | Australia | unknown | unknown | Wong et al, 2015 |

| | | | | | | | | | | | | | |
|---|---|---|---|---|---|---|---|---|---|---|---|---|---|
| MDUST146 | ERR357765 | N/A | 4.3.1.2 | unknown | unknown | Blood | 2012 | Unknown | Unknown | Unknown | unknown | unknown | Wong et al, 2015 |
| MDUST148 | ERR357766 | N/A | 4.3.1.2 | unknown | unknown | Blood | 2012 | Asia | South Asia | India | unknown | unknown | Wong et al, 2015 |
| MDUST165 | ERR357767 | N/A | 2.2.2 | unknown | unknown | Blood | 2011 | Asia | South Asia | India | unknown | unknown | Wong et al, 2015 |
| MDUST167 | ERR357768 | N/A | 4.3.1.2 | unknown | unknown | Blood | 2011 | Asia | South Asia | India | unknown | unknown | Wong et al, 2015 |
| MDUST170 | ERR357769 | N/A | 4.3.1 | unknown | unknown | Stool | 2011 | Asia | Southeast Asia | South-east Asia | unknown | unknown | Wong et al, 2015 |
| MDUST171 | ERR357770 | N/A | 3.3 | unknown | unknown | Blood | 2011 | Asia | South Asia | India | unknown | unknown | Wong et al, 2015 |
| MDUST172 | ERR357771 | N/A | 4.3.1.1 | unknown | unknown | Blood | 2011 | Asia | Southeast Asia | Thailand | unknown | unknown | Wong et al, 2015 |
| MDUST176 | ERR357772 | N/A | 3.3.2.Bd2 | unknown | unknown | Blood | 2011 | Unknown | Unknown | Unknown | unknown | unknown | Wong et al, 2015 |
| MDUST190 | ERR357773 | N/A | 4.3.1.2 | unknown | unknown | Blood | 2011 | Unknown | Unknown | Unknown | unknown | unknown | Wong et al, 2015 |
| MDUST191 | ERR357774 | N/A | 3.3.2.Bd1 | unknown | unknown | Blood | 2011 | Asia | South Asia | Bangladesh | unknown | unknown | Wong et al, 2015 |
| MDUST192 | ERR357775 | N/A | 0.0.2 | unknown | unknown | Blood | 2011 | Asia | Southeast Asia | Indonesia | unknown | unknown | Wong et al, 2015 |
| MDUST193 | ERR357776 | N/A | 4.3.1.1 | unknown | unknown | Blood | 2011 | Asia | South Asia | Bangladesh | unknown | unknown | Wong et al, 2015 |
| MDUST195 | ERR357777 | N/A | 3.3 | unknown | unknown | Blood | 2011 | Asia | South Asia | Bangladesh | unknown | unknown | Wong et al, 2015 |
| MDUST218 | ERR357778 | N/A | 2.2.2 | unknown | unknown | Blood | 2011 | Asia | Western Asia | Lebanon | unknown | unknown | Wong et al, 2015 |
| MDUST221 | ERR357779 | N/A | 3.3 | unknown | unknown | Blood | 2011 | Asia | South Asia | India | unknown | unknown | Wong et al, 2015 |
| MDUST227 | ERR357780 | N/A | 3.2.2 | unknown | unknown | Blood | 2010 | Asia | South Asia | Bangladesh | unknown | unknown | Wong et al, 2015 |
| MDUST228 | ERR357781 | N/A | 3.5.4 | unknown | unknown | Urine | 2010 | Australia & Oceania | Oceania | Samoa | unknown | unknown | Wong et al, 2015 |
| MDUST230 | ERR357782 | N/A | 4.3.1.1 | unknown | unknown | Blood | 2010 | Asia | South Asia | Bangladesh | unknown | unknown | Wong et al, 2015 |
| MDUST233 | ERR357783 | N/A | 4.3.1.2 | unknown | unknown | Blood | 2010 | Asia | South Asia | India | unknown | unknown | Wong et al, 2015 |
| MDUST238 | ERR357784 | N/A | 4.3.1.2 | unknown | unknown | Blood | 2011 | Asia | South Asia | India | unknown | unknown | Wong et al, 2015 |
| MDUST240 | ERR357785 | N/A | 4.3.1.2 | unknown | unknown | Blood | 2011 | Asia | South Asia | India | unknown | unknown | Wong et al, 2015 |
| MDUST245 | ERR357786 | N/A | 4.3.1.2 | unknown | unknown | Blood | 2011 | Asia | South Asia | India | unknown | unknown | Wong et al, 2015 |
| MDUST247 | ERR357787 | N/A | 4.3.1.2 | unknown | unknown | Stool | 2011 | Asia | South Asia | India | unknown | unknown | Wong et al, 2015 |
| MDUST251 | ERR357788 | N/A | 4.3.1.1 | unknown | unknown | Blood | 2010 | Asia | South Asia | Pakistan | unknown | unknown | Wong et al, 2015 |
| MDUST254 | ERR357789 | N/A | 4.3.1.2 | unknown | unknown | Blood | 2010 | Asia | South Asia | India | unknown | unknown | Wong et al, 2015 |
| MDUST260 | ERR357790 | N/A | 2.2.2 | unknown | unknown | Blood | 2011 | Unknown | Unknown | Unknown | unknown | unknown | Wong et al, 2015 |
| MDUST262 | ERR357791 | N/A | 4.3.1.1 | unknown | unknown | Blood | 2011 | Asia | South Asia | India | unknown | unknown | Wong et al, 2015 |
| MDUST263 | ERR357792 | N/A | 4.3.1.1 | unknown | unknown | Blood | 2011 | Asia | South Asia | Bangladesh | unknown | unknown | Wong et al, 2015 |
| MDUST264 | ERR357793 | N/A | 4.3.1.1 | unknown | unknown | Blood | 2011 | Asia | South Asia | India | unknown | unknown | Wong et al, 2015 |
| MDUST266 | ERR357794 | N/A | 4.3.1 | unknown | unknown | Stool | 2011 | Asia | South Asia | Pakistan | unknown | unknown | Wong et al, 2015 |
| MDUST271 | ERR357795 | N/A | 4.3.1.1 | unknown | unknown | Blood | 2011 | Asia | South Asia | India | unknown | unknown | Wong et al, 2015 |
| MDUST272 | ERR357796 | N/A | 4.3.1.1 | unknown | unknown | Blood | 2011 | Asia | South Asia | India | unknown | unknown | Wong et al, 2015 |
| MDUST273 | ERR357797 | N/A | 4.3.1.2 | unknown | unknown | Stool | 2011 | Asia | South Asia | India | unknown | unknown | Wong et al, 2015 |
| MDUST275 | ERR357798 | N/A | 2.1.7.1 | unknown | unknown | Stool | 1992 | Australia & Oceania | Oceania | Papua New Guinea | unknown | unknown | Wong et al, 2015 |
| MDUST290 | ERR357799 | N/A | 3.5.4 | unknown | unknown | Blood | 2008 | Australia & Oceania | Oceania | Samoa | unknown | unknown | Wong et al, 2015 |
| MDUST302 | ERR357800 | N/A | 4.1 | unknown | unknown | Unknown | 1980 | Australia & Oceania | Oceania | Papua New Guinea | unknown | unknown | Wong et al, 2015 |
| MDUST307 | ERR357801 | N/A | 4.2.1 | unknown | unknown | Unknown | 1981 | Australia & Oceania | Oceania | Fiji | unknown | unknown | Wong et al, 2015 |
| MDUST314 | ERR357802 | N/A | 4.1 | unknown | unknown | Blood | 1991 | Australia & Oceania | Oceania | Vanuatu | unknown | unknown | Wong et al, 2015 |
| MDUST324 | ERR357803 | N/A | 4.3.1.1 | unknown | unknown | Stool | 1992 | Australia & Oceania | Oceania | Fiji | unknown | unknown | Wong et al, 2015 |
| MDUST338 | ERR357804 | N/A | 2.1.7.1 | unknown | unknown | Blood | 1993 | Australia & Oceania | Oceania | Papua New Guinea | unknown | unknown | Wong et al, 2015 |
| MDUST343 | ERR357805 | N/A | 2.3.5 | unknown | unknown | Blood | 1993 | Australia & Oceania | Oceania | Fiji | unknown | unknown | Wong et al, 2015 |
| MDUST346 | ERR357806 | N/A | 4.2.1 | unknown | unknown | Blood | 1984 | Australia & Oceania | Oceania | Fiji | unknown | unknown | Wong et al, 2015 |
| MDUST347 | ERR357807 | N/A | 4.3.1.1 | unknown | unknown | Stool | 1993 | Australia & Oceania | Oceania | Fiji | unknown | unknown | Wong et al, 2015 |
| MDUST350 | ERR357808 | N/A | 2.3.5 | unknown | unknown | Blood | 1985 | Australia & Oceania | Oceania | Fiji | unknown | unknown | Wong et al, 2015 |
| MDUST353 | ERR357809 | N/A | 4.1 | unknown | unknown | Blood | 1986 | Australia & Oceania | Oceania | Papua New Guinea | unknown | unknown | Wong et al, 2015 |
| MDUST357 | ERR357810 | N/A | 2.1.7.2 | unknown | unknown | Blood | 1994 | Australia & Oceania | Oceania | Papua New Guinea | unknown | unknown | Wong et al, 2015 |
| MDUST359 | ERR357811 | N/A | 2.1.7.1 | unknown | unknown | Blood | 1996 | Australia & Oceania | Oceania | Papua New Guinea | unknown | unknown | Wong et al, 2015 |
| MDUST367 | ERR357813 | N/A | 4.3.1.2 | unknown | unknown | Stool | 2011 | Asia | South Asia | India | unknown | unknown | Wong et al, 2015 |
| MDUST369 | ERR357814 | N/A | 4.3.1.2 | unknown | unknown | Stool | 2011 | Asia | South Asia | India | unknown | unknown | Wong et al, 2015 |
| MDUST370 | ERR357815 | N/A | 4.3.1.1 | unknown | unknown | Blood | 2011 | Asia | South Asia | India | unknown | unknown | Wong et al, 2015 |

| | | | | | | | | | | | | | |
|---|---|---|---|---|---|---|---|---|---|---|---|---|---|
| MDUST371 | ERR357816 | N/A | 3.5.4 | unknown | unknown | Blood | 2011 | Australia & Oceania | Oceania | Samoa | unknown | unknown | Wong et al, 2015 |
| MDUST372 | ERR357817 | N/A | 4.3.1 | unknown | unknown | Stool | 2011 | Asia | South Asia | Pakistan | unknown | unknown | Wong et al, 2015 |
| MDUST373 | ERR357818 | N/A | 2.5 | unknown | unknown | Blood | 2011 | Asia | South Asia | India | unknown | unknown | Wong et al, 2015 |
| MDUST374 | ERR357819 | N/A | 4.3.1.1 | unknown | unknown | Stool | 2011 | Asia | South Asia | Pakistan | unknown | unknown | Wong et al, 2015 |
| MDUST375 | ERR357820 | N/A | 4.3.1.1 | unknown | unknown | Blood | 2011 | Asia | South Asia | India | unknown | unknown | Wong et al, 2015 |
| MDUST376 | ERR357821 | N/A | 4.3.1.1 | unknown | unknown | Blood | 2011 | Asia | South Asia | India | unknown | unknown | Wong et al, 2015 |
| MDUST377 | ERR357822 | N/A | 4.3.1.2 | unknown | unknown | Stool | 2011 | Asia | South Asia | India | unknown | unknown | Wong et al, 2015 |
| MDUST378 | ERR357823 | N/A | 3 | unknown | unknown | Stool | 2011 | Asia | South Asia | India | unknown | unknown | Wong et al, 2015 |
| MDUST379 | ERR357824 | N/A | 4.3.1.1 | unknown | unknown | Stool | 2011 | Asia | Southeast Asia | Myanmar | unknown | unknown | Wong et al, 2015 |
| MDUST380 | ERR357825 | N/A | 3.2.1 | unknown | unknown | Blood | 2011 | Asia | South Asia | India | unknown | unknown | Wong et al, 2015 |
| MDUST381 | ERR357826 | N/A | 3.3.1 | unknown | unknown | Stool | 2011 | Asia | South Asia | India | unknown | unknown | Wong et al, 2015 |
| MDUST384 | ERR357827 | N/A | 2.1 | unknown | unknown | Stool | 2011 | Asia | Southeast Asia | Indonesia | unknown | unknown | Wong et al, 2015 |
| MDUST390 | ERR357828 | N/A | 4.3.1.3 | unknown | unknown | Stool | 2011 | Asia | South Asia | Bangladesh | unknown | unknown | Wong et al, 2015 |
| MDUST392 | ERR357829 | N/A | 4.3.1 | unknown | unknown | Unknown | 2011 | Asia | South Asia | India | unknown | unknown | Wong et al, 2015 |
| MDUST394 | ERR357830 | N/A | 4.3.1.1 | unknown | unknown | Blood | 2011 | Asia | South Asia | India | unknown | unknown | Wong et al, 2015 |
| MDUST395 | ERR357831 | N/A | 2.1.6 | unknown | unknown | Blood | 2012 | Asia | Southeast Asia | Indonesia | unknown | unknown | Wong et al, 2015 |
| MDUST398 | ERR357832 | N/A | 4.3.1.2 | unknown | unknown | Blood | 2011 | Asia | Southeast Asia | Indonesia | unknown | unknown | Wong et al, 2015 |
| MDUST399 | ERR357833 | N/A | 4.3.1.1 | unknown | unknown | Blood | 2011 | Asia | Southeast Asia | Vietnam | unknown | unknown | Wong et al, 2015 |
| MDUST400 | ERR357834 | N/A | 4.3.1.2 | unknown | unknown | Blood | 2012 | Asia | South Asia | India | unknown | unknown | Wong et al, 2015 |
| MDUST405 | ERR357835 | N/A | 3.5.4 | unknown | unknown | Blood | 2012 | Australia & Oceania | Oceania | Samoa | unknown | unknown | Wong et al, 2015 |
| MDUST409 | ERR357836 | N/A | 4.3.1.2 | unknown | unknown | Blood | 2011 | Asia | South Asia | India | unknown | unknown | Wong et al, 2015 |
| MDUST413 | ERR357837 | N/A | 2.2.1 | unknown | unknown | Blood | 2011 | Asia | South Asia | India | unknown | unknown | Wong et al, 2015 |
| MDUST414 | ERR357838 | N/A | 4.3.1.2 | unknown | unknown | Blood | 2011 | Asia | South Asia | India | unknown | unknown | Wong et al, 2015 |
| MDUST416 | ERR357839 | N/A | 3.5.4 | unknown | unknown | Blood | 2012 | Australia & Oceania | Oceania | Samoa | unknown | unknown | Wong et al, 2015 |
| MDUST417 | ERR357840 | N/A | 3.5.4 | unknown | unknown | Blood | 2012 | Australia & Oceania | Oceania | Samoa | unknown | unknown | Wong et al, 2015 |
| MDUST419 | ERR357841 | N/A | 3.3.1 | unknown | unknown | Stool | 2011 | Australia & Oceania | Oceania | Fiji | unknown | unknown | Wong et al, 2015 |
| 820 | ERR360449 | N/A | 2.0.1 | unknown | unknown | Blood | 2003 | Asia | South Asia | Pakistan | unknown | unknown | Wong et al, 2015 |
| 5504 | ERR360450 | N/A | 2.0.1 | unknown | unknown | Blood | 2003 | Asia | South Asia | Pakistan | unknown | unknown | Wong et al, 2015 |
| 3657 | ERR360451 | N/A | 2.0.1 | unknown | unknown | Blood | 2003 | Asia | South Asia | Pakistan | unknown | unknown | Wong et al, 2015 |
| 1093 | ERR360452 | N/A | 4.3.1.1 | unknown | unknown | Blood | 2003 | Asia | South Asia | Pakistan | unknown | unknown | Wong et al, 2015 |
| 17311 | ERR360453 | N/A | 2 | unknown | unknown | Blood | 2003 | Asia | South Asia | Pakistan | unknown | unknown | Wong et al, 2015 |
| 18763 | ERR360454 | N/A | 3.3.1 | unknown | unknown | Blood | 2003 | Asia | South Asia | Pakistan | unknown | unknown | Wong et al, 2015 |
| 3802 | ERR360455 | N/A | 2 | unknown | unknown | Blood | 2003 | Asia | South Asia | Pakistan | unknown | unknown | Wong et al, 2015 |
| 16637 | ERR360456 | N/A | 2.0.1 | unknown | unknown | Blood | 2003 | Asia | South Asia | Pakistan | unknown | unknown | Wong et al, 2015 |
| 3723 | ERR360457 | N/A | 4.3.1 | unknown | unknown | Blood | 2003 | Asia | South Asia | Pakistan | unknown | unknown | Wong et al, 2015 |
| 3484 | ERR360458 | N/A | 4.3.1.1 | unknown | unknown | Blood | 2003 | Asia | South Asia | Pakistan | unknown | unknown | Wong et al, 2015 |
| 16599 | ERR360459 | N/A | 4.3.1 | unknown | unknown | Blood | 2003 | Asia | South Asia | Pakistan | unknown | unknown | Wong et al, 2015 |
| 2439 | ERR360460 | N/A | 2.0.1 | unknown | unknown | Blood | 2003 | Asia | South Asia | Pakistan | unknown | unknown | Wong et al, 2015 |
| 1382 | ERR360461 | N/A | 4.3.1.2 | unknown | unknown | Blood | 2003 | Asia | South Asia | Pakistan | unknown | unknown | Wong et al, 2015 |
| 200 | ERR360462 | N/A | 2 | unknown | unknown | Blood | 2003 | Asia | South Asia | Pakistan | unknown | unknown | Wong et al, 2015 |
| 20753 | ERR360463 | N/A | 3.2.2 | unknown | unknown | Blood | 2003 | Asia | South Asia | Pakistan | unknown | unknown | Wong et al, 2015 |
| 3163 | ERR360464 | N/A | 3.3.1 | unknown | unknown | Blood | 2003 | Asia | South Asia | Pakistan | unknown | unknown | Wong et al, 2015 |
| 11193 | ERR360465 | N/A | 3.3 | unknown | unknown | Blood | 2003 | Asia | South Asia | Pakistan | unknown | unknown | Wong et al, 2015 |
| 6289 | ERR360466 | N/A | 2 | unknown | unknown | Blood | 2003 | Asia | South Asia | Pakistan | unknown | unknown | Wong et al, 2015 |
| 2064 | ERR360467 | N/A | 2.0.1 | unknown | unknown | Blood | 2003 | Asia | South Asia | Pakistan | unknown | unknown | Wong et al, 2015 |
| 1041 | ERR360468 | N/A | 4.3.1.1 | unknown | unknown | Blood | 2003 | Asia | South Asia | Pakistan | unknown | unknown | Wong et al, 2015 |
| 4180 | ERR360469 | N/A | 2.3.3 | unknown | unknown | Blood | 2003 | Asia | South Asia | Pakistan | unknown | unknown | Wong et al, 2015 |
| 21801 | ERR360470 | N/A | 4.3.1.2 | unknown | unknown | Blood | 2003 | Asia | South Asia | Pakistan | unknown | unknown | Wong et al, 2015 |
| 3283 | ERR360471 | N/A | 4.3.1 | unknown | unknown | Blood | 2003 | Asia | South Asia | Pakistan | unknown | unknown | Wong et al, 2015 |
| 10109 | ERR360472 | N/A | 4.3.1 | unknown | unknown | Blood | 2003 | Asia | South Asia | Pakistan | unknown | unknown | Wong et al, 2015 |

| | | | | | | | | | | | | | |
|---|---|---|---|---|---|---|---|---|---|---|---|---|---|
| 20911 | ERR360473 | N/A | 3.2.2 | unknown | unknown | Blood | 2003 | Asia | South Asia | Pakistan | unknown | unknown | Wong et al, 2015 |
| 21095 | ERR360474 | N/A | 4.3.1 | unknown | unknown | Blood | 2003 | Asia | South Asia | Pakistan | unknown | unknown | Wong et al, 2015 |
| 21669 | ERR360475 | N/A | 3.2.2 | unknown | unknown | Blood | 2003 | Asia | South Asia | Pakistan | unknown | unknown | Wong et al, 2015 |
| 2176 | ERR360476 | N/A | 4.3.1.1 | unknown | unknown | Blood | 2003 | Asia | South Asia | Pakistan | unknown | unknown | Wong et al, 2015 |
| 11110 | ERR360477 | N/A | 3.2.2 | unknown | unknown | Blood | 2003 | Asia | South Asia | Pakistan | unknown | unknown | Wong et al, 2015 |
| 2513 | ERR360478 | N/A | 4.3.1 | unknown | unknown | Blood | 2003 | Asia | South Asia | Pakistan | unknown | unknown | Wong et al, 2015 |
| 3164 | ERR360479 | N/A | 2.3.3 | unknown | unknown | Blood | 2003 | Asia | South Asia | Pakistan | unknown | unknown | Wong et al, 2015 |
| 42 | ERR360480 | N/A | 4.3.1.1 | unknown | unknown | Blood | 2003 | Asia | South Asia | Pakistan | unknown | unknown | Wong et al, 2015 |
| 3769 | ERR360481 | N/A | 4.3.1 | unknown | unknown | Blood | 2003 | Asia | South Asia | Pakistan | unknown | unknown | Wong et al, 2015 |
| 7415 | ERR360482 | N/A | 3.0.1 | unknown | unknown | Blood | 2003 | Asia | South Asia | Pakistan | unknown | unknown | Wong et al, 2015 |
| E99-6646 | ERR360483 | N/A | 3.1 | unknown | unknown | Not provided | 1999 | Africa | North Africa | Algeria | unknown | unknown | Wong et al, 2015 |
| Sep-18 | ERR360484 | N/A | 0.1 | unknown | unknown | Not provided | 2009 | Africa | North Africa | Algeria | unknown | unknown | Wong et al, 2015 |
| 77-303 | ERR360485 | N/A | 2.5 | unknown | unknown | Not provided | 1977 | Asia | South Asia | India | unknown | unknown | Wong et al, 2015 |
| E98-8119 | ERR360486 | N/A | 0.1.3 | unknown | unknown | Not provided | 1998 | South America | South America | Peru | unknown | unknown | Wong et al, 2015 |
| E99-8013 | ERR360487 | N/A | 2.2 | unknown | unknown | Not provided | 1999 | Africa | North Africa | Morocco | unknown | unknown | Wong et al, 2015 |
| Jul-65 | ERR360488 | N/A | 3.1.1 | unknown | unknown | Not provided | 2007 | Africa | West Africa | IvoryCoast | unknown | unknown | Wong et al, 2015 |
| E98-2107 | ERR360489 | N/A | 2.3.1 | unknown | unknown | Not provided | 1998 | Africa | West Africa | Senegal | unknown | unknown | Wong et al, 2015 |
| E98-2601 | ERR360490 | N/A | 3.3 | unknown | unknown | Not provided | 1998 | Africa | West Africa | Gabon | unknown | unknown | Wong et al, 2015 |
| E99-9794 | ERR360491 | N/A | 3 | unknown | unknown | Not provided | 1999 | Africa | East Africa | Comoros | unknown | unknown | Wong et al, 2015 |
| E00-1382 | ERR360492 | N/A | 2.0.2 | unknown | unknown | Not provided | 2000 | Africa | North Africa | Algeria | unknown | unknown | Wong et al, 2015 |
| E97-2364 | ERR360493 | N/A | 2.5 | unknown | unknown | Not provided | 1997 | Asia | South Asia | India | unknown | unknown | Wong et al, 2015 |
| IPCU | ERR360494 | N/A | 2.3.1 | unknown | unknown | Not provided | 2007 | Africa | Central Africa | Cameroon | unknown | unknown | Wong et al, 2015 |
| Sep-21 | ERR360495 | N/A | 3.1.1 | unknown | unknown | Not provided | 2009 | Africa | West Africa | Guinea | unknown | unknown | Wong et al, 2015 |
| E97-3246 | ERR360496 | N/A | 2.5.2 | unknown | unknown | Not provided | 1997 | Africa | East Africa | Madagascar | unknown | unknown | Wong et al, 2015 |
| E02-2364 | ERR360497 | N/A | 2.5 | unknown | unknown | Not provided | 2002 | Africa | East Africa | Comoros | unknown | unknown | Wong et al, 2015 |
| E98-4364 | ERR360498 | N/A | 2.3.2 | unknown | unknown | Not provided | 1998 | North America | North America | Mexico | unknown | unknown | Wong et al, 2015 |
| E99-4879 | ERR360499 | N/A | 2 | unknown | unknown | Not provided | 1999 | Africa | North Africa | Morocco | unknown | unknown | Wong et al, 2015 |
| Sep-84 | ERR360500 | N/A | 3.1.1 | unknown | unknown | Not provided | 2009 | Africa | Africa | Africa | unknown | unknown | Wong et al, 2015 |
| Sep-14 | ERR360501 | N/A | 3.1.1 | unknown | unknown | Not provided | 2009 | Europe | Western Europe | (mother-child, African | unknown | unknown | Wong et al, 2015 |
| E00-6657 | ERR360502 | N/A | 4.1 | unknown | unknown | Not provided | 2000 | Africa | North Africa | Morocco | unknown | unknown | Wong et al, 2015 |
| E98-11555 | ERR360503 | N/A | 2 | unknown | unknown | Not provided | 1998 | Asia | Western Asia | Armenia | unknown | unknown | Wong et al, 2015 |
| E02-1963 | ERR360504 | N/A | 4.3.1.1 | unknown | unknown | Blood | 2002 | Asia | Southeast Asia | Laos | unknown | unknown | Wong et al, 2015 |
| E01-1747 | ERR360505 | N/A | 0.1.1 | unknown | unknown | Not provided | 2001 | Africa | Central Africa | Cameroon | unknown | unknown | Wong et al, 2015 |
| Sep-06 | ERR360506 | N/A | 2.5.1 | unknown | unknown | Not provided | 2009 | Africa | Central Africa | CAR | unknown | unknown | Wong et al, 2015 |
| E99-6359 | ERR360507 | N/A | 2.3.2 | unknown | unknown | Not provided | 1999 | Africa | West Africa | Mali | unknown | unknown | Wong et al, 2015 |
| 80-2002 | ERR360509 | N/A | 2.2 | unknown | unknown | Not provided | 1980 | Africa | East Africa | Madagascar | unknown | unknown | Wong et al, 2015 |
| E99-8635 | ERR360510 | N/A | 4.3.1.3 | unknown | unknown | Blood | 1999 | Asia | South Asia | Nepal | unknown | unknown | Wong et al, 2015 |
| 76-1292 | ERR360511 | N/A | 1.1.3 | unknown | unknown | Blood | 1976 | Africa | Central Africa | DRC | unknown | unknown | Wong et al, 2015 |
| Sep-24 | ERR360513 | N/A | 3.1.1 | unknown | unknown | Not provided | 2009 | Africa | Central Africa | Cameroon | unknown | unknown | Wong et al, 2015 |
| E96-12081 | ERR360514 | N/A | 4.1 | unknown | unknown | Not provided | 1996 | Africa | East Africa | Madagascar | unknown | unknown | Wong et al, 2015 |
| E97-2307 | ERR360515 | N/A | 3.3.2 | unknown | unknown | Not provided | 1997 | Asia | South Asia | India | unknown | unknown | Wong et al, 2015 |
| E99-1028 | ERR360516 | N/A | 0.1 | unknown | unknown | Not provided | 1999 | Africa | West Africa | Senegal | unknown | unknown | Wong et al, 2015 |
| E02-0530 | ERR360517 | N/A | 2.3.2 | unknown | unknown | Not provided | 2002 | Africa | West Africa | Nigeria | unknown | unknown | Wong et al, 2015 |
| E98-6926 | ERR360518 | N/A | 4.1.1 | unknown | unknown | Not provided | 1998 | Africa | West Africa | Mauritania | unknown | unknown | Wong et al, 2015 |
| IPCC | ERR360519 | N/A | 2.3.1 | unknown | unknown | Not provided | 2002 | Africa | Central Africa | Cameroon | unknown | unknown | Wong et al, 2015 |
| IPCG | ERR360520 | N/A | 2.3.1 | unknown | unknown | Not provided | 2003 | Africa | Central Africa | Cameroon | unknown | unknown | Wong et al, 2015 |
| E01-8716 | ERR360521 | N/A | 3 | unknown | unknown | Not provided | 2001 | Asia | South Asia | Sri Lanka | unknown | unknown | Wong et al, 2015 |
| E99-7012 | ERR360522 | N/A | 3.1 | unknown | unknown | Not provided | 1999 | Africa | North Africa | Morocco | unknown | unknown | Wong et al, 2015 |
| Sep-32 | ERR360523 | N/A | 3.1.1 | unknown | unknown | Not provided | 2009 | Africa | West Africa | Mauritania | unknown | unknown | Wong et al, 2015 |
| IPCO | ERR360524 | N/A | 2.3.1 | unknown | unknown | Not provided | 2006 | Africa | Central Africa | Cameroon | unknown | unknown | Wong et al, 2015 |

| | | | | | | | | | | | | | |
|---|---|---|---|---|---|---|---|---|---|---|---|---|---|
| Jun-73 | ERR360525 | N/A | 3.1.1 | unknown | unknown | Not provided | 2006 | Africa | West Africa | Ivory Coast | unknown | unknown | Wong et al, 2015 |
| Jul-20 | ERR360526 | N/A | 4.3.1.1 | unknown | unknown | Not provided | 2007 | Asia | South Asia | Bangladesh | unknown | unknown | Wong et al, 2015 |
| Jun-55 | ERR360527 | N/A | 3.1.1 | unknown | unknown | Not provided | 2006 | Africa | West Africa | Burkina Faso | unknown | unknown | Wong et al, 2015 |
| E03-6418 | ERR360528 | N/A | 4.3.1.3 | unknown | unknown | Not provided | 2003 | Asia | South Asia | Bangladesh | unknown | unknown | Wong et al, 2015 |
| Sep-13 | ERR360529 | N/A | 3.1.1 | unknown | unknown | Not provided | 2009 | Europe | Western Europe | (mother-child, African | unknown | unknown | Wong et al, 2015 |
| Sep-93 | ERR360530 | N/A | 3.1.1 | unknown | unknown | Not provided | 2009 | Africa | West Africa | Nigeria | unknown | unknown | Wong et al, 2015 |
| E00-3459 | ERR360614 | N/A | 2.2 | unknown | unknown | Not provided | 2000 | Africa | East Africa | Madagascar | unknown | unknown | Wong et al, 2015 |
| 72-1258 | ERR360615 | N/A | 2.3.2 | unknown | unknown | Not provided | 1972 | North America | North America | Mexico | unknown | unknown | Wong et al, 2015 |
| IPCK | ERR360616 | N/A | 2.3.1 | unknown | unknown | Not provided | 2004 | Africa | Central Africa | Cameroon | unknown | unknown | Wong et al, 2015 |
| E00-2388 | ERR360617 | N/A | 2.5 | unknown | unknown | Not provided | 2000 | Africa | North Africa | Egypt | unknown | unknown | Wong et al, 2015 |
| E00-2756 | ERR360618 | N/A | 2 | unknown | unknown | Not provided | 2000 | Asia | South Asia | India | unknown | unknown | Wong et al, 2015 |
| E00-3370 | ERR360619 | N/A | 0.1 | unknown | unknown | Not provided | 2000 | Africa | North Africa | Morocco | unknown | unknown | Wong et al, 2015 |
| E02-0945 | ERR360620 | N/A | 2.3.1 | unknown | unknown | Not provided | 2002 | Africa | Central Africa | Cameroon | unknown | unknown | Wong et al, 2015 |
| 04-0339 | ERR360621 | N/A | 3.1.1 | unknown | unknown | Not provided | 2004 | Africa | West Africa | Togo | unknown | unknown | Wong et al, 2015 |
| E99-6785 | ERR360622 | N/A | 0.1 | unknown | unknown | Not provided | 1999 | Africa | North Africa | Morocco | unknown | unknown | Wong et al, 2015 |
| E02-2159 | ERR360623 | N/A | 4.3.1.2 | unknown | unknown | Not provided | 2002 | Asia | South Asia | SriLanka | unknown | unknown | Wong et al, 2015 |
| E02-1687 | ERR360624 | N/A | 2.3.2 | unknown | unknown | Not provided | 2002 | Asia | Southeast Asia | Thailand | unknown | unknown | Wong et al, 2015 |
| Sep-92 | ERR360625 | N/A | 3.1.1 | unknown | unknown | Not provided | 2009 | Africa | Africa | Africa | unknown | unknown | Wong et al, 2015 |
| E99-9082 | ERR360626 | N/A | 2.3.2 | unknown | unknown | Not provided | 1999 | Africa | West Africa | Niger | unknown | unknown | Wong et al, 2015 |
| May-83 | ERR360627 | N/A | 1.1.1 | unknown | unknown | Not provided | 2005 | Africa | North Africa | Algeria | unknown | unknown | Wong et al, 2015 |
| Jul-08 | ERR360628 | N/A | 2.3.1 | unknown | unknown | Not provided | 2007 | Africa | Central Africa | Cameroon | unknown | unknown | Wong et al, 2015 |
| 69-61 | ERR360629 | N/A | 3.1 | unknown | unknown | Not provided | 1961 | Africa | North Africa | Tunisia | unknown | unknown | Wong et al, 2015 |
| Apr-67 | ERR360630 | N/A | 3.1.1 | unknown | unknown | Not provided | 2004 | Africa | West Africa | Benin | unknown | unknown | Wong et al, 2015 |
| E02-5919 | ERR360631 | N/A | 3.1 | unknown | unknown | Not provided | 2002 | Asia | East Asia | China | unknown | unknown | Wong et al, 2015 |
| 06-1510 | ERR360632 | N/A | 3.1.1 | unknown | unknown | Not provided | 2006 | Africa | West Africa | Burkina Faso | unknown | unknown | Wong et al, 2015 |
| Jul-94 | ERR360633 | N/A | 3.1.1 | unknown | unknown | Not provided | 2007 | Africa | West Africa | Ivory Coast | unknown | unknown | Wong et al, 2015 |
| 14-58 | ERR360634 | N/A | 2.3.1 | unknown | unknown | Not provided | 1958 | Africa | Central Africa | Cameroon | unknown | unknown | Wong et al, 2015 |
| E01-1811 | ERR360635 | N/A | 2.2 | unknown | unknown | Not provided | 2001 | Africa | West Africa | Mali | unknown | unknown | Wong et al, 2015 |
| IPCE | ERR360636 | N/A | 2.3.1 | unknown | unknown | Not provided | 2002 | Africa | Central Africa | Cameroon | unknown | unknown | Wong et al, 2015 |
| E99-2862 | ERR360637 | N/A | 2.2 | unknown | unknown | Not provided | 1999 | Africa | North Africa | Egypt | unknown | unknown | Wong et al, 2015 |
| E00-9345 | ERR360639 | N/A | 4.3.1.2 | unknown | unknown | Not provided | 2000 | Asia | South Asia | India | unknown | unknown | Wong et al, 2015 |
| Apr-20 | ERR360640 | N/A | 2.3.1 | unknown | unknown | Not provided | 2004 | Africa | Central Africa | Cameroon | unknown | unknown | Wong et al, 2015 |
| Sep-74 | ERR360641 | N/A | 3.1.1 | unknown | unknown | Not provided | 2009 | Africa | West Africa | Benin | unknown | unknown | Wong et al, 2015 |
| 76-1406 | ERR360642 | N/A | 3.2.1 | unknown | unknown | Blood | 1976 | Asia | Southeast Asia | Indonesia | unknown | unknown | Wong et al, 2015 |
| E02-1739 | ERR360643 | N/A | 3.1.1 | unknown | unknown | Not provided | 2002 | Africa | West Africa | Benin | unknown | unknown | Wong et al, 2015 |
| Dec-58 | ERR360644 | N/A | 0.1.1 | unknown | unknown | Not provided | 1958 | Africa | Central Africa | Cameroon | unknown | unknown | Wong et al, 2015 |
| Sep-19 | ERR360645 | N/A | 3.1.1 | unknown | unknown | Not provided | 2009 | Europe | Western Europe | France (African meal) | unknown | unknown | Wong et al, 2015 |
| May-83 | ERR360646 | N/A | 2.5.1 | unknown | unknown | Not provided | 2005 | Africa | Central Africa | Angola | unknown | unknown | Wong et al, 2015 |
| IPCI | ERR360647 | N/A | 2.3.1 | unknown | unknown | Not provided | 2004 | Africa | Central Africa | Cameroon | unknown | unknown | Wong et al, 2015 |
| IPCL | ERR360648 | N/A | 2.3.1 | unknown | unknown | Not provided | 2004 | Africa | Central Africa | Cameroon | unknown | unknown | Wong et al, 2015 |
| 72-1907 | ERR360649 | N/A | 3.4 | unknown | unknown | Not provided | 1972 | Asia | Southeast Asia | Vietnam | unknown | unknown | Wong et al, 2015 |
| E00-6999 | ERR360650 | N/A | 2.3.3 | unknown | unknown | Not provided | 2000 | South America | South America | Peru | unknown | unknown | Wong et al, 2015 |
| 06-467 | ERR360651 | N/A | 2.3.1 | unknown | unknown | Not provided | 2006 | Africa | Central Africa | Cameroon | unknown | unknown | Wong et al, 2015 |
| E02-0232 | ERR360652 | N/A | 2.5 | unknown | unknown | Not provided | 2002 | South America | South America | French Guiana | unknown | unknown | Wong et al, 2015 |
| E97-9141 | ERR360653 | N/A | 2.3.2 | unknown | unknown | Not provided | 1997 | Asia | Western Asia | Turkey | unknown | unknown | Wong et al, 2015 |
| E00-6599 | ERR360654 | N/A | 3.3.1 | unknown | unknown | Not provided | 2000 | Africa | West Africa | Cape Verde | unknown | unknown | Wong et al, 2015 |
| IPCA | ERR360655 | N/A | 0.0.1 | unknown | unknown | Not provided | 2000 | Africa | Central Africa | Cameroon | unknown | unknown | Wong et al, 2015 |
| E00-7878 | ERR360656 | N/A | 2 | unknown | unknown | Not provided | 2000 | Africa | North Africa | Morocco | unknown | unknown | Wong et al, 2015 |
| E98-8120 | ERR360657 | N/A | 0.0.1 | unknown | unknown | Not provided | 1998 | Africa | Central Africa | Cameroon | unknown | unknown | Wong et al, 2015 |
| E00-7463 | ERR360658 | N/A | 3.0.1 | unknown | unknown | Not provided | 2000 | Africa | North Africa | Morocco | unknown | unknown | Wong et al, 2015 |

| | | | | | | | | | | | | | |
|---|---|---|---|---|---|---|---|---|---|---|---|---|---|
| E99-6478 | ERR360659 | N/A | 3.1 | unknown | unknown | Not provided | 1999 | Africa | West Africa | Guinea | unknown | unknown | Wong et al, 2015 |
| IPCR | ERR360660 | N/A | 2.3.1 | unknown | unknown | Not provided | 2006 | Africa | Central Africa | Cameroon | unknown | unknown | Wong et al, 2015 |
| 73-1102 | ERR360661 | N/A | 4.1 | unknown | unknown | Not provided | 1973 | Asia | Southeast Asia | Vietnam | unknown | unknown | Wong et al, 2015 |
| E00-5869 | ERR360662 | N/A | 2.3.3 | unknown | unknown | Not provided | 2000 | Asia | South Asia | Bangladesh | unknown | unknown | Wong et al, 2015 |
| IPCT | ERR360663 | N/A | 2.3.1 | unknown | unknown | Not provided | 2007 | Africa | Central Africa | Cameroon | unknown | unknown | Wong et al, 2015 |
| E02-2612 | ERR360664 | N/A | 2.2 | unknown | unknown | Not provided | 2002 | Asia | South Asia | India | unknown | unknown | Wong et al, 2015 |
| IPCQ | ERR360665 | N/A | 2.3.1 | unknown | unknown | Not provided | 2006 | Africa | Central Africa | Cameroon | unknown | unknown | Wong et al, 2015 |
| 07-291 | ERR360666 | N/A | 2.3.1 | unknown | unknown | Not provided | 2007 | Africa | Central Africa | Cameroon | unknown | unknown | Wong et al, 2015 |
| E99-5920 | ERR360667 | N/A | 2.0.2 | unknown | unknown | Not provided | 1999 | Africa | North Africa | Tunisia | unknown | unknown | Wong et al, 2015 |
| E99-8067 | ERR360668 | N/A | 0.1.2 | unknown | unknown | Not provided | 1999 | Africa | North Africa | Algeria | unknown | unknown | Wong et al, 2015 |
| 07-702 | ERR360669 | N/A | 2.3.1 | unknown | unknown | Not provided | 2007 | Africa | Central Africa | Cameroon | unknown | unknown | Wong et al, 2015 |
| 08-10571 | ERR360670 | N/A | 3.1.1 | unknown | unknown | Not provided | 2008 | Africa | West Africa | Mali | unknown | unknown | Wong et al, 2015 |
| Apr-45 | ERR360671 | N/A | 3.1.1 | unknown | unknown | Not provided | 2004 | Africa | West Africa | Benin | unknown | unknown | Wong et al, 2015 |
| E00-3201 | ERR360672 | N/A | 2.3.2 | unknown | unknown | Not provided | 2000 | Africa | West Africa | Mali | unknown | unknown | Wong et al, 2015 |
| E01-0407 | ERR360673 | N/A | 4.1 | unknown | unknown | Not provided | 2001 | Asia | Western Asia | Turkey | unknown | unknown | Wong et al, 2015 |
| IPCP | ERR360674 | N/A | 2.3.1 | unknown | unknown | Not provided | 2006 | Africa | Central Africa | Cameroon | unknown | unknown | Wong et al, 2015 |
| 06-1513 | ERR360675 | N/A | 3.1.1 | unknown | unknown | Not provided | 2006 | Africa | West Africa | Burkina Faso | unknown | unknown | Wong et al, 2015 |
| Sep-77 | ERR360676 | N/A | 0.1 | unknown | unknown | Not provided | 2009 | Africa | North Africa | Algeria | unknown | unknown | Wong et al, 2015 |
| Aug-14 | ERR360677 | N/A | 2.5.1 | unknown | unknown | Blood | 2008 | Africa | Central Africa | DRC | unknown | unknown | Wong et al, 2015 |
| 07-977 | ERR360678 | N/A | 2.5.1 | unknown | unknown | Not provided | 2007 | Africa | Central Africa | CAR | unknown | unknown | Wong et al, 2015 |
| Jun-29 | ERR360679 | N/A | 2.3.1 | unknown | unknown | Not provided | 2006 | Africa | Central Africa | Cameroon | unknown | unknown | Wong et al, 2015 |
| E02-1536 | ERR360680 | N/A | 2.5 | unknown | unknown | Not provided | 2002 | South America | South America | French Guiana | unknown | unknown | Wong et al, 2015 |
| E02-0937 | ERR360681 | N/A | 2.5 | unknown | unknown | Not provided | 2002 | Africa | East Africa | Comoros | unknown | unknown | Wong et al, 2015 |
| Jun-90 | ERR360682 | N/A | 3.1.1 | unknown | unknown | Not provided | 2006 | Africa | West Africa | Togo | unknown | unknown | Wong et al, 2015 |
| Q0904130 | ERR360685 | N/A | 2.5.1 | unknown | unknown | Not provided | 2006 | Africa | Central Africa | CAR | unknown | unknown | Wong et al, 2015 |
| Aug-27 | ERR360686 | N/A | 3.1.1 | unknown | unknown | Not provided | 2008 | Africa | West Africa | Ivory Coast | unknown | unknown | Wong et al, 2015 |
| Sep-97 | ERR360687 | N/A | 3.1.1 | unknown | unknown | Not provided | 2009 | Europe | Western Europe | France (African meal) | unknown | unknown | Wong et al, 2015 |
| E99-8095 | ERR360688 | N/A | 4.1 | unknown | unknown | Not provided | 1999 | Africa | North Africa | Algeria | unknown | unknown | Wong et al, 2015 |
| IPCS | ERR360689 | N/A | 4.1.1 | unknown | unknown | Not provided | 2006 | Africa | Central Africa | Cameroon | unknown | unknown | Wong et al, 2015 |
| IPCN | ERR360690 | N/A | 3.1.1 | unknown | unknown | Not provided | 2005 | Africa | Central Africa | Cameroon | unknown | unknown | Wong et al, 2015 |
| E01-7006 | ERR360691 | N/A | 4.1 | unknown | unknown | Not provided | 2001 | Asia | Western Asia | Lebanon | unknown | unknown | Wong et al, 2015 |
| E01-7101 | ERR360692 | N/A | 2.3.2 | unknown | unknown | Not provided | 2001 | Africa | West Africa | Togo | unknown | unknown | Wong et al, 2015 |
| Aug-02 | ERR360693 | N/A | 4.3.1.1 | unknown | unknown | Not provided | 2008 | Asia | Western Asia | Palestine | unknown | unknown | Wong et al, 2015 |
| E00-7666 | ERR360694 | N/A | 3.1 | unknown | unknown | Not provided | 2000 | Africa | North Africa | Tunisia | unknown | unknown | Wong et al, 2015 |
| E01-5741 | ERR360695 | N/A | 1.1.3 | unknown | unknown | Not provided | 2001 | Africa | Central Africa | Angola | unknown | unknown | Wong et al, 2015 |
| E00-6172 | ERR360696 | N/A | 3.5 | unknown | unknown | Blood | 2000 | Asia | Southeast Asia | Indonesia | unknown | unknown | Wong et al, 2015 |
| 2003-013044 | ERR360738 | N/A | 4.3.1.1 | unknown | unknown | Blood | 2008 | Asia | Southeast Asia | Cambodia | unknown | unknown | Wong et al, 2015 |
| 2008-006720 | ERR360739 | N/A | 4.3.1.1 | unknown | unknown | Blood | 2008 | Asia | Southeast Asia | Cambodia | unknown | unknown | Wong et al, 2015 |
| 2008-007824 | ERR360740 | N/A | 4.3.1.1 | unknown | unknown | Blood | 2008 | Asia | Southeast Asia | Cambodia | unknown | unknown | Wong et al, 2015 |
| 2008-010439 | ERR360741 | N/A | 4.3.1.1 | unknown | unknown | Blood | 2008 | Asia | Southeast Asia | Cambodia | unknown | unknown | Wong et al, 2015 |
| 2008-011433 | ERR360742 | N/A | 4.3.1.1 | unknown | unknown | Blood | 2008 | Asia | Southeast Asia | Cambodia | unknown | unknown | Wong et al, 2015 |
| 2005-005326 | ERR360743 | N/A | 4.3.1.1 | unknown | unknown | Blood | 2008 | Asia | Southeast Asia | Cambodia | unknown | unknown | Wong et al, 2015 |
| 2009-019983 | ERR360744 | N/A | 4.3.1.1 | unknown | unknown | Blood | 2009 | Asia | Southeast Asia | Cambodia | unknown | unknown | Wong et al, 2015 |
| 2009-020666 | ERR360745 | N/A | 4.3.1.1 | unknown | unknown | Thigh pus | 2009 | Asia | Southeast Asia | Cambodia | unknown | unknown | Wong et al, 2015 |
| 2003-011175 | ERR360746 | N/A | 4.3.1.1 | unknown | unknown | Blood | 2009 | Asia | Southeast Asia | Cambodia | unknown | unknown | Wong et al, 2015 |
| 2010-002168 | ERR360747 | N/A | 3.4 | unknown | unknown | Blood | 2010 | Asia | Southeast Asia | Cambodia | unknown | unknown | Wong et al, 2015 |
| 2010-012339 | ERR360748 | N/A | 4.3.1.1 | unknown | unknown | Blood | 2010 | Asia | Southeast Asia | Cambodia | unknown | unknown | Wong et al, 2015 |
| 2009-008387 | ERR360754 | N/A | 4.3.1.1 | unknown | unknown | Pleural fluid | 2010 | Asia | Southeast Asia | Cambodia | unknown | unknown | Wong et al, 2015 |
| 2011-002088 | ERR360755 | N/A | 4.3.1.1 | unknown | unknown | Blood | 2011 | Asia | Southeast Asia | Cambodia | unknown | unknown | Wong et al, 2015 |
| 01-2010-000934 | ERR360756 | N/A | 4.3.1.1 | unknown | unknown | Blood | 2011 | Asia | Southeast Asia | Cambodia | unknown | unknown | Wong et al, 2015 |

| | | | | | | | | | | | | | |
|---|---|---|---|---|---|---|---|---|---|---|---|---|---|
| 2011-007188 | ERR360758 | N/A | 4.3.1.1 | unknown | unknown | Gallbladder fluid | 2011 | Asia | Southeast Asia | Cambodia | unknown | unknown | Wong et al, 2015 |
| 2002-214318 | ERR360759 | N/A | 4.3.1.1 | unknown | unknown | Blood | 2011 | Asia | Southeast Asia | Cambodia | unknown | unknown | Wong et al, 2015 |
| 2011-008491 | ERR360760 | N/A | 4.3.1.1 | unknown | unknown | Blood | 2011 | Asia | Southeast Asia | Cambodia | unknown | unknown | Wong et al, 2015 |
| 2010-006459 | ERR360761 | N/A | 4.3.1.1 | unknown | unknown | Blood | 2011 | Asia | Southeast Asia | Cambodia | unknown | unknown | Wong et al, 2015 |
| 2002-205957 | ERR360762 | N/A | 4.3.1.1 | unknown | unknown | Blood | 2011 | Asia | Southeast Asia | Cambodia | unknown | unknown | Wong et al, 2015 |
| 2007-017523 | ERR360763 | N/A | 4.3.1.1 | unknown | unknown | Blood | 2011 | Asia | Southeast Asia | Cambodia | unknown | unknown | Wong et al, 2015 |
| 2011-009404 | ERR360765 | N/A | 4.3.1.1 | unknown | unknown | Blood | 2011 | Asia | Southeast Asia | Cambodia | unknown | unknown | Wong et al, 2015 |
| 2009-004677 | ERR360766 | N/A | 4.3.1.1 | unknown | unknown | Blood | 2011 | Asia | Southeast Asia | Cambodia | unknown | unknown | Wong et al, 2015 |
| 2011-015914 | ERR360769 | N/A | 4.3.1.1 | unknown | unknown | Blood | 2011 | Asia | Southeast Asia | Cambodia | unknown | unknown | Wong et al, 2015 |
| 2011-017747 | ERR360770 | N/A | 4.3.1.1 | unknown | unknown | Blood | 2011 | Asia | Southeast Asia | Cambodia | unknown | unknown | Wong et al, 2015 |
| 2011-017563 | ERR360771 | N/A | 4.3.1.1 | unknown | unknown | Blood | 2011 | Asia | Southeast Asia | Cambodia | unknown | unknown | Wong et al, 2015 |
| 2011-019378 | ERR360772 | N/A | 4.3.1.1 | unknown | unknown | Blood | 2011 | Asia | Southeast Asia | Cambodia | unknown | unknown | Wong et al, 2015 |
| 2011-019409 | ERR360773 | N/A | 4.3.1.1 | unknown | unknown | Hip joint pus | 2011 | Asia | Southeast Asia | Cambodia | unknown | unknown | Wong et al, 2015 |
| 01-2011-005712 | ERR360774 | N/A | 4.3.1.1 | unknown | unknown | Blood | 2011 | Asia | Southeast Asia | Cambodia | unknown | unknown | Wong et al, 2015 |
| 2012-010218 | ERR360775 | N/A | 4.3.1.1 | unknown | unknown | Blood | 2012 | Asia | Southeast Asia | Cambodia | unknown | unknown | Wong et al, 2015 |
| 2012-011302 | ERR360776 | N/A | 4.3.1.1 | unknown | unknown | Stool | 2012 | Asia | Southeast Asia | Cambodia | unknown | unknown | Wong et al, 2015 |
| 2012-011333 | ERR360777 | N/A | 4.3.1.1 | unknown | unknown | Blood | 2012 | Asia | Southeast Asia | Cambodia | unknown | unknown | Wong et al, 2015 |
| 2010-011024 | ERR360778 | N/A | 4.3.1.1 | unknown | unknown | Blood | 2012 | Asia | Southeast Asia | Cambodia | unknown | unknown | Wong et al, 2015 |
| 2011-019645 | ERR360779 | N/A | 4.3.1.1 | unknown | unknown | Blood | 2012 | Asia | Southeast Asia | Cambodia | unknown | unknown | Wong et al, 2015 |
| 2009-019909 | ERR360780 | N/A | 4.3.1.1 | unknown | unknown | Blood | 2012 | Asia | Southeast Asia | Cambodia | unknown | unknown | Wong et al, 2015 |
| 2012-011344 | ERR360781 | N/A | 4.3.1.1 | unknown | unknown | Blood | 2012 | Asia | Southeast Asia | Cambodia | unknown | unknown | Wong et al, 2015 |
| 2008-011089 | ERR360782 | N/A | 4.3.1.1 | unknown | unknown | Blood | 2012 | Asia | Southeast Asia | Cambodia | unknown | unknown | Wong et al, 2015 |
| 2012-010551 | ERR360783 | N/A | 4.3.1.1 | unknown | unknown | Blood | 2012 | Asia | Southeast Asia | Cambodia | unknown | unknown | Wong et al, 2015 |
| 2012-010616 | ERR360785 | N/A | 4.3.1.1 | unknown | unknown | Blood | 2012 | Asia | Southeast Asia | Cambodia | unknown | unknown | Wong et al, 2015 |
| 01-2010-000073 | ERR360786 | N/A | 4.3.1.1 | unknown | unknown | Blood | 2012 | Asia | Southeast Asia | Cambodia | unknown | unknown | Wong et al, 2015 |
| 2012-011809 | ERR360787 | N/A | 4.3.1.1 | unknown | unknown | Blood | 2012 | Asia | Southeast Asia | Cambodia | unknown | unknown | Wong et al, 2015 |
| 2012-010204 | ERR360788 | N/A | 4.3.1.1 | unknown | unknown | Blood | 2012 | Asia | Southeast Asia | Cambodia | unknown | unknown | Wong et al, 2015 |
| 2005-012355 | ERR360789 | N/A | 4.3.1.1 | unknown | unknown | Blood | 2012 | Asia | Southeast Asia | Cambodia | unknown | unknown | Wong et al, 2015 |
| 2012-015100 | ERR360791 | N/A | 4.3.1.1 | unknown | unknown | Blood | 2012 | Asia | Southeast Asia | Cambodia | unknown | unknown | Wong et al, 2015 |
| 2012-017695 | ERR360792 | N/A | 4.3.1.1 | unknown | unknown | Blood | 2012 | Asia | Southeast Asia | Cambodia | unknown | unknown | Wong et al, 2015 |
| 1035759 | ERR360793 | N/A | 4.3.1.1.EA1 | unknown | unknown | Blood | 2012 | Africa | Southern Africa | Malawi | unknown | unknown | Wong et al, 2015 |
| BKQ4CF | ERR360794 | N/A | 4.3.1.1.EA1 | unknown | unknown | Blood | 2013 | Africa | Southern Africa | Malawi | unknown | unknown | Wong et al, 2015 |
| A57501 | ERR360795 | N/A | 4.1.1 | unknown | unknown | Blood | 2010 | Africa | Southern Africa | Malawi | unknown | unknown | Wong et al, 2015 |
| A58372 | ERR360796 | N/A | 4.1.1 | unknown | unknown | Blood | 2010 | Africa | Southern Africa | Malawi | unknown | unknown | Wong et al, 2015 |
| 1005624 | ERR360797 | N/A | 4.3.1.1.EA1 | unknown | unknown | Blood | 2010 | Africa | Southern Africa | Malawi | unknown | unknown | Wong et al, 2015 |
| 1003108 | ERR360798 | N/A | 4.3.1.1.EA1 | unknown | unknown | Blood | 2010 | Africa | Southern Africa | Malawi | unknown | unknown | Wong et al, 2015 |
| BKQ22S | ERR360799 | N/A | 4.3.1.1.EA1 | unknown | unknown | Blood | 2012 | Africa | Southern Africa | Malawi | unknown | unknown | Wong et al, 2015 |
| 1004971 | ERR360800 | N/A | 2.4.1 | unknown | unknown | Blood | 2010 | Africa | Southern Africa | Malawi | unknown | unknown | Wong et al, 2015 |
| 1006099 | ERR360801 | N/A | 4.3.1.1.EA1 | unknown | unknown | Blood | 2010 | Africa | Southern Africa | Malawi | unknown | unknown | Wong et al, 2015 |
| 1003117 | ERR360802 | N/A | 4.3.1.1.EA1 | unknown | unknown | Blood | 2010 | Africa | Southern Africa | Malawi | unknown | unknown | Wong et al, 2015 |
| BKQ1GA | ERR360803 | N/A | 4.1.1 | unknown | unknown | Blood | 2012 | Africa | Southern Africa | Malawi | unknown | unknown | Wong et al, 2015 |
| 1007000 | ERR360804 | N/A | 2.2.2 | unknown | unknown | Blood | 2010 | Africa | Southern Africa | Malawi | unknown | unknown | Wong et al, 2015 |
| BHA1P5 | ERR360805 | N/A | 4.3.1.1.EA1 | unknown | unknown | Blood | 2012 | Africa | Southern Africa | Malawi | unknown | unknown | Wong et al, 2015 |
| BKQ2KZ | ERR360806 | N/A | 4.3.1.1.EA1 | unknown | unknown | Blood | 2012 | Africa | Southern Africa | Malawi | unknown | unknown | Wong et al, 2015 |
| BKQ4SJ | ERR360807 | N/A | 4.3.1.1.EA1 | unknown | unknown | Blood | 2013 | Africa | Southern Africa | Malawi | unknown | unknown | Wong et al, 2015 |
| BKQ4NZ | ERR360809 | N/A | 4.3.1.1.EA1 | unknown | unknown | Blood | 2013 | Africa | Southern Africa | Malawi | unknown | unknown | Wong et al, 2015 |
| BKQ2H5 | ERR360810 | N/A | 4.3.1.1.EA1 | unknown | unknown | Blood | 2012 | Africa | Southern Africa | Malawi | unknown | unknown | Wong et al, 2015 |
| BHA2HH | ERR360811 | N/A | 4.3.1.1.EA1 | unknown | unknown | Blood | 2012 | Africa | Southern Africa | Malawi | unknown | unknown | Wong et al, 2015 |
| 1003078 | ERR360812 | N/A | 4.3.1.1 | unknown | unknown | Blood | 2010 | Africa | Southern Africa | Malawi | unknown | unknown | Wong et al, 2015 |
| D56675 | ERR360813 | N/A | 2.4.1 | unknown | unknown | Blood | 2010 | Africa | Southern Africa | Malawi | unknown | unknown | Wong et al, 2015 |

| | | | | | | | | | | | | | |
|---|---|---|---|---|---|---|---|---|---|---|---|---|---|
| A59204 | ERR360814 | N/A | 4.1.1 | unknown | unknown | Blood | 2010 | Africa | Southern Africa | Malawi | unknown | unknown | Wong et al, 2015 |
| BHA2QL | ERR360815 | N/A | 4.3.1.1.EA1 | unknown | unknown | Blood | 2013 | Africa | Southern Africa | Malawi | unknown | unknown | Wong et al, 2015 |
| BKQ4H1 | ERR360816 | N/A | 4.3.1.1.EA1 | unknown | unknown | Blood | 2013 | Africa | Southern Africa | Malawi | unknown | unknown | Wong et al, 2015 |
| 1038377 | ERR360817 | N/A | 4.3.1.1.EA1 | unknown | unknown | Blood | 2012 | Africa | Southern Africa | Malawi | unknown | unknown | Wong et al, 2015 |
| 1037921 | ERR360818 | N/A | 4.3.1.1.EA1 | unknown | unknown | Blood | 2012 | Africa | Southern Africa | Malawi | unknown | unknown | Wong et al, 2015 |
| BKQ4E3 | ERR360819 | N/A | 4.3.1.1.EA1 | unknown | unknown | Blood | 2013 | Africa | Southern Africa | Malawi | unknown | unknown | Wong et al, 2015 |
| D55055 | ERR360820 | N/A | 2.2 | unknown | unknown | Blood | 2010 | Africa | Southern Africa | Malawi | unknown | unknown | Wong et al, 2015 |
| D56393 | ERR360821 | N/A | 4.1.1 | unknown | unknown | Blood | 2010 | Africa | Southern Africa | Malawi | unknown | unknown | Wong et al, 2015 |
| A58452 | ERR360822 | N/A | 3.3.1 | unknown | unknown | Blood | 2010 | Africa | Southern Africa | Malawi | unknown | unknown | Wong et al, 2015 |
| A59217 | ERR360824 | N/A | 4.3.1.2 | unknown | unknown | Blood | 2010 | Africa | Southern Africa | Malawi | unknown | unknown | Wong et al, 2015 |
| A58450 | ERR360825 | N/A | 4.1.1 | unknown | unknown | Blood | 2010 | Africa | Southern Africa | Malawi | unknown | unknown | Wong et al, 2015 |
| D54782 | ERR360826 | N/A | 2.4.1 | unknown | unknown | Blood | 2010 | Africa | Southern Africa | Malawi | unknown | unknown | Wong et al, 2015 |
| BKQ4JU | ERR360827 | N/A | 4.3.1.1.EA1 | unknown | unknown | Blood | 2013 | Africa | Southern Africa | Malawi | unknown | unknown | Wong et al, 2015 |
| 1036491 | ERR360828 | N/A | 4.3.1.1.EA1 | unknown | unknown | Blood | 2012 | Africa | Southern Africa | Malawi | unknown | unknown | Wong et al, 2015 |
| 1036143 | ERR360829 | N/A | 4.3.1.1.EA1 | unknown | unknown | Blood | 2012 | Africa | Southern Africa | Malawi | unknown | unknown | Wong et al, 2015 |
| BKQ4D9 | ERR360830 | N/A | 4.3.1.1.EA1 | unknown | unknown | Blood | 2013 | Africa | Southern Africa | Malawi | unknown | unknown | Wong et al, 2015 |
| A58390 | ERR360832 | N/A | 4.3.1.2 | unknown | unknown | Blood | 2010 | Africa | Southern Africa | Malawi | unknown | unknown | Wong et al, 2015 |
| A59307 | ERR360833 | N/A | 4.1.1 | unknown | unknown | Blood | 2010 | Africa | Southern Africa | Malawi | unknown | unknown | Wong et al, 2015 |
| 2010-013700 | ERR360845 | N/A | 4.3.1.1 | unknown | unknown | Blood | 2010 | Asia | Southeast Asia | Cambodia | unknown | unknown | Wong et al, 2015 |
| 2004-007692 | ERR360846 | N/A | 4.3.1.1 | unknown | unknown | Blood | 2010 | Asia | Southeast Asia | Cambodia | unknown | unknown | Wong et al, 2015 |
| 2008-000234 | ERR360847 | N/A | 4.3.1.1 | unknown | unknown | Blood | 2010 | Asia | Southeast Asia | Cambodia | unknown | unknown | Wong et al, 2015 |
| 2009-017391 | ERR360848 | N/A | 4.3.1.1 | unknown | unknown | Blood | 2010 | Asia | Southeast Asia | Cambodia | unknown | unknown | Wong et al, 2015 |
| 2008-003275 | ERR360889 | N/A | 4.3.1.1 | unknown | unknown | Blood | 2008 | Asia | Southeast Asia | Cambodia | unknown | unknown | Wong et al, 2015 |
| 2007-025892 | ERR360890 | N/A | 4.3.1.1 | unknown | unknown | Blood | 2008 | Asia | Southeast Asia | Cambodia | unknown | unknown | Wong et al, 2015 |
| 2008-007203 | ERR360891 | N/A | 4.3.1.1 | unknown | unknown | Blood | 2008 | Asia | Southeast Asia | Cambodia | unknown | unknown | Wong et al, 2015 |
| 2008-007991 | ERR360948 | N/A | 4.3.1.1 | unknown | unknown | Blood | 2008 | Asia | Southeast Asia | Cambodia | unknown | unknown | Wong et al, 2015 |
| 2007-016977 | ERR360949 | N/A | 4.3.1.1 | unknown | unknown | Blood | 2008 | Asia | Southeast Asia | Cambodia | unknown | unknown | Wong et al, 2015 |
| 2008-012004 | ERR360950 | N/A | 4.3.1.1 | unknown | unknown | Blood | 2008 | Asia | Southeast Asia | Cambodia | unknown | unknown | Wong et al, 2015 |
| 2010-012486 | ERR360952 | N/A | 4.3.1.1 | unknown | unknown | Blood | 2010 | Asia | Southeast Asia | Cambodia | unknown | unknown | Wong et al, 2015 |
| 2010-015926 | ERR360954 | N/A | 4.3.1.1 | unknown | unknown | Blood | 2010 | Asia | Southeast Asia | Cambodia | unknown | unknown | Wong et al, 2015 |
| 2010-018118 | ERR360955 | N/A | 4.3.1.1 | unknown | unknown | Blood | 2010 | Asia | Southeast Asia | Cambodia | unknown | unknown | Wong et al, 2015 |
| 2010-018908 | ERR360956 | N/A | 4.3.1.1 | unknown | unknown | Blood | 2010 | Asia | Southeast Asia | Cambodia | unknown | unknown | Wong et al, 2015 |
| 01-2010-005821 | ERR360958 | N/A | 4.3.1.1 | unknown | unknown | Blood | 2010 | Asia | Southeast Asia | Cambodia | unknown | unknown | Wong et al, 2015 |
| 2010-020905 | ERR360959 | N/A | 4.3.1.1 | unknown | unknown | Blood | 2010 | Asia | Southeast Asia | Cambodia | unknown | unknown | Wong et al, 2015 |
| 2006-024861 | ERR360960 | N/A | 4.3.1.1 | unknown | unknown | Blood | 2010 | Asia | Southeast Asia | Cambodia | unknown | unknown | Wong et al, 2015 |
| 01-2011-000664 | ERR360961 | N/A | 4.3.1.1 | unknown | unknown | Blood | 2011 | Asia | Southeast Asia | Cambodia | unknown | unknown | Wong et al, 2015 |
| 2011-024146 | ERR360962 | N/A | 4.3.1.1 | unknown | unknown | Blood | 2011 | Asia | Southeast Asia | Cambodia | unknown | unknown | Wong et al, 2015 |
| 2012-010535 | ERR360963 | N/A | 4.3.1.1 | unknown | unknown | Blood | 2012 | Asia | Southeast Asia | Cambodia | unknown | unknown | Wong et al, 2015 |
| 2012-010742 | ERR360964 | N/A | 4.3.1.1 | unknown | unknown | Blood | 2012 | Asia | Southeast Asia | Cambodia | unknown | unknown | Wong et al, 2015 |
| 2012-009188 | ERR360965 | N/A | 4.3.1.1 | unknown | unknown | Blood | 2012 | Asia | Southeast Asia | Cambodia | unknown | unknown | Wong et al, 2015 |
| 2009-018790 | ERR360966 | N/A | 4.3.1.1 | unknown | unknown | Blood | 2012 | Asia | Southeast Asia | Cambodia | unknown | unknown | Wong et al, 2015 |
| 01-2012-002891 | ERR360967 | N/A | 4.3.1.1 | unknown | unknown | Blood | 2012 | Asia | Southeast Asia | Cambodia | unknown | unknown | Wong et al, 2015 |
| 2008-020523 | ERR360968 | N/A | 4.3.1.1 | unknown | unknown | Blood | 2012 | Asia | Southeast Asia | Cambodia | unknown | unknown | Wong et al, 2015 |
| 2003-000735 | ERR360970 | N/A | 4.3.1.1 | unknown | unknown | Blood | 2012 | Asia | Southeast Asia | Cambodia | unknown | unknown | Wong et al, 2015 |
| 2012-011897 | ERR360971 | N/A | 4.3.1.1 | unknown | unknown | Blood | 2012 | Asia | Southeast Asia | Cambodia | unknown | unknown | Wong et al, 2015 |
| 2012-011885 | ERR360972 | N/A | 4.3.1.1 | unknown | unknown | Blood | 2012 | Asia | Southeast Asia | Cambodia | unknown | unknown | Wong et al, 2015 |
| 2012-011284 | ERR360973 | N/A | 4.3.1.1 | unknown | unknown | Blood | 2012 | Asia | Southeast Asia | Cambodia | unknown | unknown | Wong et al, 2015 |
| 01-2012-003910 | ERR360975 | N/A | 4.3.1.1 | unknown | unknown | Stool | 2012 | Asia | Southeast Asia | Cambodia | unknown | unknown | Wong et al, 2015 |
| 2012-004687 | ERR360976 | N/A | 4.3.1.1 | unknown | unknown | Blood | 2012 | Asia | Southeast Asia | Cambodia | unknown | unknown | Wong et al, 2015 |
| 2012-016786 | ERR360977 | N/A | 4.3.1.1 | unknown | unknown | Blood | 2012 | Asia | Southeast Asia | Cambodia | unknown | unknown | Wong et al, 2015 |

| | | | | | | | | | | | | | |
|---|---|---|---|---|---|---|---|---|---|---|---|---|---|
| 01-2010-001056 | ERR360978 | N/A | 4.3.1.1 | unknown | unknown | Blood | 2012 | Asia | Southeast Asia | Cambodia | unknown | unknown | Wong et al, 2015 |
| 2012-010759 | ERR360979 | N/A | 4.3.1.1 | unknown | unknown | Blood | 2012 | Asia | Southeast Asia | Cambodia | unknown | unknown | Wong et al, 2015 |
| 2012-000742 | ERR360980 | N/A | 4.3.1.1 | unknown | unknown | Blood | 2012 | Asia | Southeast Asia | Cambodia | unknown | unknown | Wong et al, 2015 |
| 2012-010559 | ERR360981 | N/A | 4.3.1.1 | unknown | unknown | Blood | 2012 | Asia | Southeast Asia | Cambodia | unknown | unknown | Wong et al, 2015 |
| 01-2012-002876 | ERR360982 | N/A | 4.3.1.1 | unknown | unknown | Blood | 2012 | Asia | Southeast Asia | Cambodia | unknown | unknown | Wong et al, 2015 |
| 2009-012656 | ERR360983 | N/A | 4.3.1.1 | unknown | unknown | Blood | 2012 | Asia | Southeast Asia | Cambodia | unknown | unknown | Wong et al, 2015 |
| 2010-019292 | ERR360984 | N/A | 4.3.1.1 | unknown | unknown | Blood | 2012 | Asia | Southeast Asia | Cambodia | unknown | unknown | Wong et al, 2015 |
| 01-2010-005164 | ERR360985 | N/A | 4.3.1.1 | unknown | unknown | Blood | 2012 | Asia | Southeast Asia | Cambodia | unknown | unknown | Wong et al, 2015 |
| 2012-012029 | ERR360986 | N/A | 4.3.1.1 | unknown | unknown | Blood | 2012 | Asia | Southeast Asia | Cambodia | unknown | unknown | Wong et al, 2015 |
| 2012-011846 | ERR360987 | N/A | 4.3.1.1 | unknown | unknown | Blood | 2012 | Asia | Southeast Asia | Cambodia | unknown | unknown | Wong et al, 2015 |
| 2012-012131 | ERR360988 | N/A | 4.3.1.1 | unknown | unknown | Blood | 2012 | Asia | Southeast Asia | Cambodia | unknown | unknown | Wong et al, 2015 |
| 2004-006350 | ERR360990 | N/A | 4.3.1.1 | unknown | unknown | Blood | 2012 | Asia | Southeast Asia | Cambodia | unknown | unknown | Wong et al, 2015 |
| 2002-213689 | ERR360991 | N/A | 4.3.1.1 | unknown | unknown | Blood | 2012 | Asia | Southeast Asia | Cambodia | unknown | unknown | Wong et al, 2015 |
| 2002-215815 | ERR360992 | N/A | 4.3.1.1 | unknown | unknown | Blood | 2012 | Asia | Southeast Asia | Cambodia | unknown | unknown | Wong et al, 2015 |
| 01-2011-005568 | ERR360993 | N/A | 4.3.1.1 | unknown | unknown | Blood | 2012 | Asia | Southeast Asia | Cambodia | unknown | unknown | Wong et al, 2015 |
| 01-2011-004759 | ERR360994 | N/A | 4.3.1.1 | unknown | unknown | Blood | 2012 | Asia | Southeast Asia | Cambodia | unknown | unknown | Wong et al, 2015 |
| 2012-014737 | ERR360995 | N/A | 4.1 | unknown | unknown | Blood | 2012 | Asia | Southeast Asia | Cambodia | unknown | unknown | Wong et al, 2015 |
| 2012-015146 | ERR360996 | N/A | 4.3.1.1 | unknown | unknown | Blood | 2012 | Asia | Southeast Asia | Cambodia | unknown | unknown | Wong et al, 2015 |
| 2012-019308 | ERR360997 | N/A | 4.3.1.1 | unknown | unknown | Blood | 2012 | Asia | Southeast Asia | Cambodia | unknown | unknown | Wong et al, 2015 |
| 2012-020127 | ERR360998 | N/A | 4.3.1.1 | unknown | unknown | Blood | 2012 | Asia | Southeast Asia | Cambodia | unknown | unknown | Wong et al, 2015 |
| 2011-008969 | ERR360999 | N/A | 4.3.1.1 | unknown | unknown | Blood | 2012 | Asia | Southeast Asia | Cambodia | unknown | unknown | Wong et al, 2015 |
| 2012-022646 | ERR361000 | N/A | 4.3.1.1 | unknown | unknown | Blood | 2012 | Asia | Southeast Asia | Cambodia | unknown | unknown | Wong et al, 2015 |
| BCR43 | ERR420413 | N/A | 4.3.1.2 | unknown | unknown | Blood | 2009 | Asia | South Asia | India | unknown | unknown | Wong et al, 2015 |
| BCR48 | ERR420414 | N/A | 4.3.1.2 | unknown | unknown | Blood | 2009 | Asia | South Asia | India | unknown | unknown | Wong et al, 2015 |
| BCR49 | ERR420415 | N/A | 4.3.1.1 | unknown | unknown | Blood | 2009 | Asia | South Asia | India | unknown | unknown | Wong et al, 2015 |
| BCR52 | ERR420416 | N/A | 2.2.1 | unknown | unknown | Blood | 2009 | Asia | South Asia | India | unknown | unknown | Wong et al, 2015 |
| BCR62 | ERR420417 | N/A | 4.3.1.2 | unknown | unknown | Blood | 2009 | Asia | South Asia | India | unknown | unknown | Wong et al, 2015 |
| BCR89 | ERR420418 | N/A | 2.2.1 | unknown | unknown | Blood | 2010 | Asia | South Asia | India | unknown | unknown | Wong et al, 2015 |
| BCR108 | ERR420420 | N/A | 4.3.1.2 | unknown | unknown | Blood | 2010 | Asia | South Asia | India | unknown | unknown | Wong et al, 2015 |
| BCR110 | ERR420421 | N/A | 4.3.1.2 | unknown | unknown | Blood | 2010 | Asia | South Asia | India | unknown | unknown | Wong et al, 2015 |
| BCR162 | ERR420422 | N/A | 4.3.1.2 | unknown | unknown | Blood | 2011 | Asia | South Asia | India | unknown | unknown | Wong et al, 2015 |
| BCR170 | ERR420423 | N/A | 4.3.1.2 | unknown | unknown | Blood | 2011 | Asia | South Asia | India | unknown | unknown | Wong et al, 2015 |
| BCR175 | ERR420424 | N/A | 4.3.1.2 | unknown | unknown | Blood | 2011 | Asia | South Asia | India | unknown | unknown | Wong et al, 2015 |
| BCR177 | ERR420425 | N/A | 4.3.1.1 | unknown | unknown | Blood | 2011 | Asia | South Asia | India | unknown | unknown | Wong et al, 2015 |
| BCR191 | ERR420426 | N/A | 4.3.1.2 | unknown | unknown | Blood | 2011 | Asia | South Asia | India | unknown | unknown | Wong et al, 2015 |
| BCR211 | ERR420427 | N/A | 4.3.1.2 | unknown | unknown | Blood | 2011 | Asia | South Asia | India | unknown | unknown | Wong et al, 2015 |
| BCR232 | ERR420428 | N/A | 2 | unknown | unknown | Blood | 2011 | Asia | South Asia | India | unknown | unknown | Wong et al, 2015 |
| SP2 | ERR420429 | N/A | 4.3.1.1 | unknown | unknown | Blood | 2009 | Asia | South Asia | India | unknown | unknown | Wong et al, 2015 |
| SP9 | ERR420430 | N/A | 4.3.1.2 | unknown | unknown | Blood | 2009 | Asia | South Asia | India | unknown | unknown | Wong et al, 2015 |
| SP11 | ERR420431 | N/A | 4.3.1.2 | unknown | unknown | Blood | 2009 | Asia | South Asia | India | unknown | unknown | Wong et al, 2015 |
| SP47 | ERR420432 | N/A | 4.3.1.1 | unknown | unknown | Blood | 2009 | Asia | South Asia | India | unknown | unknown | Wong et al, 2015 |
| SP66 | ERR420433 | N/A | 3.3.2 | unknown | unknown | Blood | 2009 | Asia | South Asia | India | unknown | unknown | Wong et al, 2015 |
| SP80 | ERR420434 | N/A | 4.3.1.2 | unknown | unknown | Blood | 2011 | Asia | South Asia | India | unknown | unknown | Wong et al, 2015 |
| H09176224 | ERR422751 | N/A | 4.3.1.2 | unknown | unknown | Blood | 2009 | Asia | South Asia | India | unknown | unknown | Wong et al, 2016 |
| H09188293 | ERR422752 | N/A | 4.3.1.1 | unknown | unknown | Blood | 2009 | Asia | South Asia | Bangladesh | unknown | unknown | Wong et al, 2016 |
| H0920834 | ERR422754 | N/A | 3.3.2.Bd1 | unknown | unknown | Blood | 2009 | Asia | South Asia | Bangladesh | unknown | unknown | Wong et al, 2016 |
| H09266336 | ERR422757 | N/A | 4.3.1 | unknown | unknown | Blood | 2009 | Unknown | Unknown | Unknown | unknown | unknown | Wong et al, 2016 |
| H09354568 | ERR422759 | N/A | 4.3.1.1 | unknown | unknown | Blood | 2009 | Asia | South Asia | India | unknown | unknown | Wong et al, 2016 |
| H10044399 | ERR422762 | N/A | 4.3.1.2 | unknown | unknown | Blood | 2010 | Asia | South Asia | India | unknown | unknown | Wong et al, 2016 |
| H10046338 | ERR422764 | N/A | 4.3.1.3 | unknown | unknown | Blood | 2010 | Unknown | Unknown | Unknown | unknown | unknown | Wong et al, 2016 |

| | | | | | | | | | | | | | |
|---|---|---|---|---|---|---|---|---|---|---|---|---|---|
| H10048267 | ERR422765 | N/A | 4.3.1.2 | unknown | unknown | Blood | 2010 | Asia | South Asia | India | unknown | unknown | Wong et al, 2016 |
| H10084431 | ERR422766 | N/A | 4.3.1.1 | unknown | unknown | Blood | 2010 | Asia | South Asia | Bangladesh | unknown | unknown | Wong et al, 2016 |
| H10090344 | ERR422767 | N/A | 3.3.2 | unknown | unknown | Blood | 2010 | Asia | South Asia | Bangladesh, India | unknown | unknown | Wong et al, 2016 |
| H10092333 | ERR422768 | N/A | 3.3.2 | unknown | unknown | Blood | 2010 | Asia | South Asia | Bangladesh, India | unknown | unknown | Wong et al, 2016 |
| H10100338 | ERR422769 | N/A | 3.3.2.Bd2 | unknown | unknown | Blood | 2010 | Asia | South Asia | Bangladesh | unknown | unknown | Wong et al, 2016 |
| H10106250 | ERR422770 | N/A | 4.3.1.2 | unknown | unknown | Blood | 2010 | Asia | South Asia | India | unknown | unknown | Wong et al, 2016 |
| H10108318 | ERR422771 | N/A | 3.3.2.Bd2 | unknown | unknown | Blood | 2010 | Asia | South Asia | Bangladesh | unknown | unknown | Wong et al, 2016 |
| H1011698 | ERR422772 | N/A | 3.3.2.Bd2 | unknown | unknown | Blood | 2010 | Asia | South Asia | Bangladesh | unknown | unknown | Wong et al, 2016 |
| H10122493 | ERR422773 | N/A | 3.3.2.Bd2 | unknown | unknown | Blood | 2010 | Asia | South Asia | Bangladesh | unknown | unknown | Wong et al, 2016 |
| H10134171 | ERR422774 | N/A | 3.3.2.Bd2 | unknown | unknown | Blood | 2010 | Asia | South Asia | Bangladesh | unknown | unknown | Wong et al, 2016 |
| H10148201 | ERR422775 | N/A | 4.3.1.1 | unknown | unknown | Blood | 2010 | Asia | South Asia | Bangladesh | unknown | unknown | Wong et al, 2016 |
| H1015074 | ERR422776 | N/A | 4.3.1.1 | unknown | unknown | Blood | 2010 | Asia | South Asia | Bangladesh | unknown | unknown | Wong et al, 2016 |
| H10182335 | ERR422778 | N/A | 3.3.1 | unknown | unknown | Blood | 2010 | Unknown | Unknown | Unknown | unknown | unknown | Wong et al, 2016 |
| H10184197 | ERR422780 | N/A | 4.3.1 | unknown | unknown | Blood | 2010 | Asia | South Asia | India | unknown | unknown | Wong et al, 2016 |
| H10202409 | ERR422781 | N/A | 4.3.1.1 | unknown | unknown | Blood | 2010 | Asia | South Asia | Pakistan | unknown | unknown | Wong et al, 2016 |
| H10294193 | ERR422787 | N/A | 4.3.1.2 | unknown | unknown | Blood | 2010 | Asia | South Asia | India | unknown | unknown | Wong et al, 2016 |
| H10310204 | ERR422788 | N/A | 2.3.3 | unknown | unknown | Blood | 2010 | Asia | South Asia | Bangladesh | unknown | unknown | Wong et al, 2016 |
| H10334602 | ERR422790 | N/A | 2.1.7 | unknown | unknown | Blood | 2010 | Asia | South Asia | India | unknown | unknown | Wong et al, 2016 |
| H10340496 | ERR422792 | N/A | 2.1.7 | unknown | unknown | Blood | 2010 | Asia | South Asia | India | unknown | unknown | Wong et al, 2016 |
| H10382491 | ERR422796 | N/A | 4.3.1 | unknown | unknown | Blood | 2010 | Unknown | Unknown | Unknown | unknown | unknown | Wong et al, 2016 |
| H10394694 | ERR422798 | N/A | 4.3.1 | unknown | unknown | Blood | 2010 | Unknown | Unknown | Unknown | unknown | unknown | Wong et al, 2016 |
| H10432531 | ERR422800 | N/A | 4.3.1.1 | unknown | unknown | Blood | 2010 | Asia | South Asia | Bangladesh | unknown | unknown | Wong et al, 2016 |
| H10462591 | ERR422801 | N/A | 4.3.1.1 | unknown | unknown | Blood | 2010 | Asia | South Asia | Bangladesh | unknown | unknown | Wong et al, 2016 |
| H11044442 | ERR422803 | N/A | 4.3.1 | unknown | unknown | Blood | 2011 | Asia | South Asia | Pakistan | unknown | unknown | Wong et al, 2016 |
| H11054345 | ERR422804 | N/A | 4.3.1.2 | unknown | unknown | Blood | 2011 | Asia | South Asia | India | unknown | unknown | Wong et al, 2016 |
| H11096403 | ERR422805 | N/A | 4.3.1.1 | unknown | unknown | Blood | 2011 | Asia | South Asia | Pakistan | unknown | unknown | Wong et al, 2016 |
| H11150244 | ERR422807 | N/A | 4.3.1.3 | unknown | unknown | Blood | 2011 | Asia | South Asia | Bangladesh | unknown | unknown | Wong et al, 2016 |
| H11194353 | ERR422810 | N/A | 4.3.1.1 | unknown | unknown | Blood | 2011 | Asia | South Asia | Bangladesh | unknown | unknown | Wong et al, 2016 |
| H11244554 | ERR422811 | N/A | 2.3.3 | unknown | unknown | Blood | 2011 | Asia | South Asia | Bangladesh | unknown | unknown | Wong et al, 2016 |
| H11254619 | ERR422812 | N/A | 4.3.1.1 | unknown | unknown | Blood | 2011 | Asia | South Asia | Bangladesh | unknown | unknown | Wong et al, 2016 |
| H11344588 | ERR422813 | N/A | 4.3.1.1 | unknown | unknown | Blood | 2011 | Asia | South Asia | Bangladesh | unknown | unknown | Wong et al, 2016 |
| H11354680 | ERR422814 | N/A | 4.3.1.2 | unknown | unknown | Blood | 2011 | Asia | South Asia | India | unknown | unknown | Wong et al, 2016 |
| H11372597 | ERR422815 | N/A | 4.3.1.1 | unknown | unknown | Blood | 2011 | Asia | South Asia | Bangladesh | unknown | unknown | Wong et al, 2016 |
| H11372598 | ERR422816 | N/A | 4.3.1 | unknown | unknown | Blood | 2011 | Unknown | Unknown | Unknown | unknown | unknown | Wong et al, 2016 |
| H11374579 | ERR422817 | N/A | 4.3.1.1 | unknown | unknown | Blood | 2011 | Asia | South Asia | India | unknown | unknown | Wong et al, 2016 |
| H11388492 | ERR422818 | N/A | 4.3.1.2 | unknown | unknown | Blood | 2011 | Asia | South Asia | Bangladesh | unknown | unknown | Wong et al, 2016 |
| H11420359 | ERR422819 | N/A | 4.3.1 | unknown | unknown | Blood | 2011 | Asia | South Asia | India | unknown | unknown | Wong et al, 2016 |
| H11424491 | ERR422820 | N/A | 3.2.2 | unknown | unknown | Blood | 2011 | Asia | South Asia | Bangladesh | unknown | unknown | Wong et al, 2016 |
| H11436323 | ERR422822 | N/A | 3.3.2.Bd1 | unknown | unknown | Blood | 2011 | Asia | South Asia | Bangladesh | unknown | unknown | Wong et al, 2016 |
| H12014687 | ERR422824 | N/A | 4.3.1.1 | unknown | unknown | Blood | 2011 | Asia | South Asia | Bangladesh | unknown | unknown | Wong et al, 2016 |
| H12054632 | ERR422826 | N/A | 4.3.1.3.Bdq | unknown | unknown | Blood | 2012 | Asia | South Asia | Bangladesh | unknown | unknown | Wong et al, 2016 |
| H121541053 | ERR422828 | N/A | 4.3.1 | unknown | unknown | Blood | 2012 | Asia | South Asia | India | unknown | unknown | Wong et al, 2016 |
| H12164448 | ERR422829 | N/A | 4.3.1.1 | unknown | unknown | Blood | 2012 | Asia | South Asia | Bangladesh | unknown | unknown | Wong et al, 2016 |
| H12200559 | ERR422830 | N/A | 2.0.1 | unknown | unknown | Blood | 2012 | Asia | South Asia | Bangladesh | unknown | unknown | Wong et al, 2016 |
| H12276630 | ERR422831 | N/A | 4.3.1.1 | unknown | unknown | Blood | 2012 | Asia | South Asia | Bangladesh | unknown | unknown | Wong et al, 2016 |
| H12282824 | ERR422832 | N/A | 3.2.2 | unknown | unknown | Blood | 2012 | Asia | South Asia | Pakistan | unknown | unknown | Wong et al, 2016 |
| H12382595 | ERR422833 | N/A | 4.3.1 | unknown | unknown | Blood | 2012 | Asia | South Asia | Pakistan | unknown | unknown | Wong et al, 2016 |
| H12414817 | ERR422834 | N/A | 4.3.1 | unknown | unknown | Blood | 2012 | Asia | South Asia | Pakistan | unknown | unknown | Wong et al, 2016 |
| H05196408 | ERR485127 | N/A | 4.3.1 | unknown | unknown | Blood | 2005 | Unknown | Unknown | Unknown | unknown | unknown | Wong et al, 2016 |
| H05212226 | ERR485129 | N/A | 4.3.1 | unknown | unknown | Blood | 2005 | Unknown | Unknown | Unknown | unknown | unknown | Wong et al, 2016 |

| | | | | | | | | | | | | | |
|---|---|---|---|---|---|---|---|---|---|---|---|---|---|
| H06136379 | ERR485132 | N/A | 4.3.1.3 | unknown | unknown | Blood | 2006 | Unknown | Unknown | Unknown | unknown | unknown | Wong et al, 2016 |
| H06136380 | ERR485133 | N/A | 4.3.1.3 | unknown | unknown | Blood | 2006 | Unknown | Unknown | Unknown | unknown | unknown | Wong et al, 2016 |
| H06414500 | ERR485138 | N/A | 4.3.1.1 | unknown | unknown | Blood | 2006 | Asia | South Asia | Bangladesh | unknown | unknown | Wong et al, 2016 |
| H06448472 | ERR485139 | N/A | 4.1 | unknown | unknown | Blood | 2006 | Asia | South Asia | India | unknown | unknown | Wong et al, 2016 |
| H07188291 | ERR485140 | N/A | 3.3.2.Bd2 | unknown | unknown | Blood | 2007 | Asia | South Asia | Bangladesh | unknown | unknown | Wong et al, 2016 |
| H08248234 | ERR485145 | N/A | 3.3.2.Bd2 | unknown | unknown | Blood | 2008 | Asia | South Asia | Bangladesh | unknown | unknown | Wong et al, 2016 |
| H09176223 | ERR485147 | N/A | 3.3.2.Bd2 | unknown | unknown | Blood | 2009 | Unknown | Unknown | Unknown | unknown | unknown | Wong et al, 2016 |
