## Supplementary Table 1 for "Population structure and evolution of *Salmonella enterica* serotype Typhi in Zimbabwe before a typhoid conjugate vaccine immunization campaign"

| tree | figure | number isolates | number bootstrap | outgroup |
|---|---|---|---|---|
| Global tree | Figure 5 | 1,999 | 350 | Paratyphi A |
| genotyphi clade 4.3 | Figure 6 | 1,006 | 450 | S. Typhi CT18 |
| 4.3.1.1EA1 tree | Figure 7 | | from genotyphi clade 4.3 tree) | from genotyphi clade 4.3 tree) |
| Zimbabwe isolates | Figure 2 / Figure 3 | 95 | 550 | ERR360627, ERR360499 and ERR360655 |
| Clade 3.3.1 | Suppl Figure 1 | 41 | NA (extracted from global tree) | NA (extracted from global tree) |
